## Supplementary material for "Dynamic Compression Flows for Neuroscience Data": Python code: paper_plot_ball_dim3_lag0.ipynb

```
# %% [code]
import numpy as np
import matplotlib.pyplot as plt
from matplotlib.collections import LineCollection
from matplotlib.colors import ListedColormap, BoundaryNorm, LinearSegmentedColormap, Normalize
from mpl_toolkits.mplot3d.art3d import Line3DCollection
from matplotlib.cm import ScalarMappable
from matplotlib.colors import LogNorm
import matplotlib as mpl
from mpl_toolkits.axes_grid1.inset_locator import inset_axes

import torch
import dnnlib.util_v7 as util
from tqdm import tqdm
import pickle
from matplotlib.ticker import FuncFormatter
import pickle, types
from torch_utils import persistence

from PIL import Image

import numpy as np
import pickle
import os
import re 
import json 
import click
import torch
from torch_utils import distributed as dist
from training import toy_training_loop_vfm_noDataGen
from tqdm import tqdm
import matplotlib.pyplot as plt
from matplotlib import colors
from plot_neural_v7 import *
from plot_sim_results_v7 import *
from plot_sim_results_img_v7 import * 


import pandas as pd

import warnings
warnings.filterwarnings('ignore', 'Grad strides do not match bucket view strides') # False warning 
device = torch.device("cuda" if torch.cuda.is_available() else "cpu")
torch.set_default_device(device)
colorgrad = "PuOr"

# %% [markdown]
# 0. Some functions...

# %% [code]
def plot_tau(T, out_imgs_tau, dset_samples, trial_idx, tau, title=None):
    # Plot True
    plot_frame(dset_samples, T, trial_idx)
    plt.suptitle(f'true, trial = {trial_idx}')
    plt.subplots_adjust(bottom=0.2)  # Make room for horizontal colorbar

    # Plot Simulated
    plot_frame(out_imgs_tau, T, trial_idx)
    if title is None:
        plt.suptitle(f'simulated- tau = {tau}')
    else:
        plt.suptitle(title)
    plt.subplots_adjust(bottom=0.2)

# %% [code]
def recon_check_x0(encoder, trial_check, T_check, dset_samples):
    encoder = encoder.to(device).eval()
    X1_ori = dset_samples[trial_check][T_check,0,:,:]
    data_flattent = [np.reshape(a, (a.shape[0], -1), order='C') for a in dset_samples]
    with torch.inference_mode():
        X1 = torch.from_numpy(data_flattent[trial_check][T_check,:]).to(device=device, dtype=torch.float32)
        X0 = encoder.rsample(X1)
    X0_back = X0.reshape(X0.shape[0], C, H, W)
    return X0_back[0,0,:,:].detach().cpu().numpy()

# %% [code]
def recon_check_x0_allT(encoder, trial_check, dset_samples):
    encoder = encoder.to(device).eval()
    X1_ori = dset_samples[trial_check][T_check,0,:,:]
    data_flattent = [np.reshape(a, (a.shape[0], -1), order='C') for a in dset_samples]
    with torch.inference_mode():
        X1 = torch.from_numpy(data_flattent[trial_check]).to(device=device, dtype=torch.float32)
        X0 = encoder.rsample(X1)
    X0_back = X0.reshape(X0.shape[0], C, H, W)
    return X0_back.detach().cpu().numpy()

# %% [code]
def mu_group_recon(bias,
                   loading_np: np.ndarray,
                   mu_all: np.ndarray,
                   mu_idx=None,             # int | list[int] | np.ndarray | slice | None
                   H: int = 28,
                   W: int = 28,
                   mean_img: np.ndarray = None,
                   clip=None):
    D = loading_np.shape[0]
    assert loading_np.ndim == 2
    K_load = loading_np.shape[1]          # number of latent dims in loading
    N, K_mu = mu_all.shape               # number of latent dims in mu_all
    assert H * W == D

    # normalize mu_idx to an integer index array IN LATENT SPACE
    if mu_idx is None:
        idx = np.arange(min(K_load, K_mu))
    elif isinstance(mu_idx, slice):
        idx = np.arange(min(K_load, K_mu))[mu_idx]
    else:
        arr = np.asarray(mu_idx)
        if arr.dtype == bool:
            idx = np.flatnonzero(arr)
        else:
            idx = arr.astype(int, copy=False).ravel()

    # sanity: all requested dims must exist in both loading and mu_all
    max_idx = int(idx.max()) if idx.size > 0 else -1
    assert max_idx < K_load, f"mu_idx uses dim {max_idx}, but loading has only {K_load} cols"
    assert max_idx < K_mu,   f"mu_idx uses dim {max_idx}, but mu_all has only {K_mu} cols"

    A_sub  = loading_np[:, idx]     # (D, K_sel)
    mu_sub = mu_all[:, idx]         # (N, K_sel)
    recon_flat = mu_sub @ A_sub.T   # (N, D)

    if mean_img is not None:
        mean_flat = mean_img.reshape(-1)
        assert mean_flat.size == D
        recon_flat = recon_flat + mean_flat

    recon_flat = recon_flat + bias
    recon = recon_flat.reshape(N, H, W)
    if clip is not None:
        lo, hi = clip
        recon = np.clip(recon, lo, hi)
    return recon


# %% [code]
def plot_mu_velocity_2d(
    mu,
    v_mu,
    trajectories=None,
    dims_2d=(0, 1),
    title=None,
    arrow_frac=0.05,
    arrow_size=1.2,
    linewidth=1.5,
    traj_alpha=0.4,
    traj_width=0.8,
    traj_color="gray",
    max_trajs=300,
    quiver_alpha=0.6,
    max_points=3000,
    figsize=(7, 6),
    speed_color=None,                      # NEW
    colorbar_label=r"$\|\dot{\mu}\|$",     # NEW
):
    fig, ax = plt.subplots(figsize=figsize, constrained_layout=True)
    ax.grid(False)

    # optional color source (full norm, e.g. 784D)
    if speed_color is not None:
        speed_color = speed_color.detach().cpu().numpy() if hasattr(speed_color, "detach") else np.asarray(speed_color)
        speed_color = np.asarray(speed_color).reshape(-1)
        assert speed_color.shape[0] == mu.shape[0]
        vmin_c = float(speed_color.min())
        vmax_c = float(speed_color.max())

    traj_list = []
    if trajectories is not None:
        traj_source = trajectories
        for i, traj in enumerate(traj_source):
            if i >= max_trajs:
                break
            t_np = traj.detach().cpu().numpy() if hasattr(traj, "detach") else np.array(traj)
            traj_list.append(t_np)

    d1, d2 = dims_2d
    for t_np in traj_list:
        ax.plot(
            t_np[:, d1], t_np[:, d2],
            color=traj_color, alpha=traj_alpha, linewidth=traj_width, zorder=1
        )

    coords = mu[:, [d1, d2]]
    vecs   = v_mu[:, [d1, d2]]
    N = coords.shape[0]

    if max_points is not None and N > max_points:
        idx = np.random.choice(N, size=max_points, replace=False)
        coords_2d = coords[idx]
        vecs_2d   = vecs[idx]
        if speed_color is not None:
            speed_c = speed_color[idx]
    else:
        coords_2d = coords
        vecs_2d   = vecs
        if speed_color is not None:
            speed_c = speed_color

    # directions from projected velocity
    speed_proj = np.linalg.norm(vecs_2d, axis=1)
    eps = 1e-8
    dirs = vecs_2d / (speed_proj[:, None] + eps)

    # color values
    if speed_color is None:
        speed_c = speed_proj
        vmin_c = float(speed_c.min())
        vmax_c = float(speed_c.max())

    x_min, x_max = coords_2d[:, 0].min(), coords_2d[:, 0].max()
    y_min, y_max = coords_2d[:, 1].min(), coords_2d[:, 1].max()
    span = max(x_max - x_min, y_max - y_min) + eps
    L = arrow_frac * span
    U = dirs[:, 0] * L
    V = dirs[:, 1] * L

    base_width = 0.004
    base_headwidth = 4.0
    base_headlength = 6.0
    base_headaxislength = 5.0
    width = base_width * arrow_size

    norm = Normalize(vmin=vmin_c, vmax=vmax_c)
    q = ax.quiver(
        coords_2d[:, 0], coords_2d[:, 1], U, V, speed_c,
        cmap=cm.viridis, norm=norm, angles="xy", scale_units="xy", scale=1.0,
        width=width,
        headwidth=base_headwidth * arrow_size,
        headlength=base_headlength * arrow_size,
        headaxislength=base_headaxislength * arrow_size,
        linewidth=linewidth, alpha=quiver_alpha, zorder=2
    )
    cb = fig.colorbar(q, ax=ax, fraction=0.046, pad=0.04)
    cb.set_label(colorbar_label)

    ax.set_xlabel(fr"$\mu_{{{d1+1}}}$")
    ax.set_ylabel(fr"$\mu_{{{d2+1}}}$")
    if title:
        ax.set_title(title)
    ax.axis("equal")

    return fig, ax, q


def plot_mu_velocity_3d(
    mu,
    v_mu,
    trajectories=None,
    dims_3d=(0, 1, 2),
    title=None,
    arrow_frac=0.1,
    arrow_size=1.5,
    linewidth=1.5,
    base_arrow_length_ratio=0.3,
    traj_alpha=0.4,
    traj_width=0.8,
    quiver_alpha=0.6,
    traj_color="gray",
    max_trajs=300,
    max_points=3000,
    figsize=(10, 7),
    speed_color=None,                      # NEW
    colorbar_label=r"$\|\dot{\mu}\|$",     # NEW
):
    fig = plt.figure(figsize=figsize, constrained_layout=True)
    ax = fig.add_subplot(1, 1, 1, projection="3d")
    ax.set_box_aspect((1, 1, 1))
    ax.grid(False)
    for axis in (ax.xaxis, ax.yaxis, ax.zaxis):
        axis.pane.fill = False
        axis.pane.set_edgecolor((1, 1, 1, 0))
        axis._axinfo["grid"]["color"] = (1, 1, 1, 0)
        axis._axinfo["grid"]["linewidth"] = 0

    # optional color source (full norm, e.g. 784D)
    if speed_color is not None:
        speed_color = speed_color.detach().cpu().numpy() if hasattr(speed_color, "detach") else np.asarray(speed_color)
        speed_color = np.asarray(speed_color).reshape(-1)
        assert speed_color.shape[0] == mu.shape[0]
        vmin_c = float(speed_color.min())
        vmax_c = float(speed_color.max())

    traj_list = []
    if trajectories is not None:
        traj_source = trajectories
        for i, traj in enumerate(traj_source):
            if i >= max_trajs:
                break
            t_np = traj.detach().cpu().numpy() if hasattr(traj, "detach") else np.array(traj)
            traj_list.append(t_np)

    d1, d2, d3 = dims_3d
    for t_np in traj_list:
        ax.plot(
            t_np[:, d1], t_np[:, d2], t_np[:, d3],
            color=traj_color, alpha=traj_alpha, linewidth=traj_width, zorder=1
        )

    coords = mu[:, [d1, d2, d3]]
    vecs   = v_mu[:, [d1, d2, d3]]
    N = coords.shape[0]

    if max_points is not None and N > max_points:
        idx = np.random.choice(N, size=max_points, replace=False)
        coords_3d = coords[idx]
        vecs_3d   = vecs[idx]
        if speed_color is not None:
            speed_c = speed_color[idx]
    else:
        coords_3d = coords
        vecs_3d   = vecs
        if speed_color is not None:
            speed_c = speed_color

    # directions from projected velocity
    speed_proj = np.linalg.norm(vecs_3d, axis=1)
    eps = 1e-8
    dirs = vecs_3d / (speed_proj[:, None] + eps)

    # color values
    if speed_color is None:
        speed_c = speed_proj
        vmin_c = float(speed_c.min())
        vmax_c = float(speed_c.max())

    mins = coords_3d.min(axis=0)
    maxs = coords_3d.max(axis=0)
    span = float(np.max(maxs - mins) + eps)

    L = arrow_frac * span
    U, V, W = dirs[:, 0], dirs[:, 1], dirs[:, 2]

    norm = Normalize(vmin=vmin_c, vmax=vmax_c)
    colors = cm.viridis(norm(speed_c))
    colors[:, 3] = quiver_alpha

    ax.quiver(
        coords_3d[:, 0], coords_3d[:, 1], coords_3d[:, 2],
        U, V, W,
        length=L, normalize=False, colors=colors,
        linewidth=linewidth * arrow_size,
        arrow_length_ratio=base_arrow_length_ratio * arrow_size,
        zorder=2
    )

    mappable = cm.ScalarMappable(norm=norm, cmap="viridis")
    mappable.set_array(speed_c)
    cb = fig.colorbar(mappable, ax=ax, fraction=0.035, pad=0.08)
    cb.set_label(colorbar_label)

    ax.set_xlabel(fr"$\mu_{{{d1+1}}}$")
    ax.set_ylabel(fr"$\mu_{{{d2+1}}}$")
    ax.set_zlabel(fr"$\mu_{{{d3+1}}}$")
    if title:
        ax.set_title(title)

    return fig, ax

# %% [code]
def plot_frame_compact(
    rows,                 # list like [dset_samples, out_imgs_tau0_flow, ...]
    trial_idx,
    idxs,                 # 1D int array, columns
    row_labels=None,      # list of strings, one per row
    mean_img=None,
    figsize=(6, 2.6),
    img_cmap="gray",
    vmin=None,
    vmax=None,
    border_from=None,
    border_lw=3.0,
    left=0.10, right=0.99, bottom=0.05, top=0.95,
    wspace=0.02, hspace=0.02,
):
    import numpy as np
    import matplotlib.pyplot as plt

    idxs = np.asarray(idxs).astype(int)
    n_rows = len(rows)
    n_cols = len(idxs)

    # --- global vmin/vmax (consistent across rows) ---
    if vmin is None or vmax is None:
        mn, mx = np.inf, -np.inf
        for data in rows:
            X = data[trial_idx]  # [T, 1, H, W] (torch or np)
            for i in idxs:
                img = X[i, 0]
                if hasattr(img, "detach"):
                    img = img.detach().cpu().numpy()
                else:
                    img = np.asarray(img)
                if mean_img is not None:
                    img = img + mean_img
                mn = min(mn, float(img.min()))
                mx = max(mx, float(img.max()))
        if vmin is None: vmin = mn
        if vmax is None: vmax = mx

    if border_from is not None:
        cmap = border_from.get_cmap()
        norm = border_from.norm
        col_colors = cmap(norm(idxs.astype(float)))
    else:
        col_colors = ["k"] * n_cols

    fig, axes = plt.subplots(n_rows, n_cols, figsize=figsize, squeeze=False)
    fig.subplots_adjust(left=left, right=right, bottom=bottom, top=top, wspace=wspace, hspace=hspace)

    for r, data in enumerate(rows):
        X = data[trial_idx]
        for c, i in enumerate(idxs):
            ax = axes[r, c]

            img = X[i, 0]
            if hasattr(img, "detach"):
                img = img.detach().cpu().numpy()
            else:
                img = np.asarray(img)
            if mean_img is not None:
                img = img + mean_img

            ax.imshow(img, cmap=img_cmap, vmin=vmin, vmax=vmax)

            # keep spines for colored border, so don't ax.axis("off")
            ax.set_xticks([]); ax.set_yticks([])

            # titles only on first row
            if r == 0:
                ax.set_title(f"t={i}")
            else:
                ax.set_title("")

            # colored border per column
            for sp in ax.spines.values():
                sp.set_visible(True)
                sp.set_linewidth(border_lw)
                sp.set_color(col_colors[c])

    # row labels on the left, centered per row (compact)
    if row_labels is not None:
        for r, lab in enumerate(row_labels):
            pos = axes[r, 0].get_position()
            y = 0.5 * (pos.y0 + pos.y1)
            x = pos.x0 - 0.02
            fig.text(x, y, lab, ha="right", va="center")

    return fig, axes

# %% [markdown]
# 1. Read Data

# %% [code]
folder_data = 'out/ball/t_detach/data'
useCov = False

dataset_samples = np.load(folder_data + "/dataset_samples.npz", allow_pickle=True)
cov_static_samples = np.load(folder_data + "/cov_static_trunc_only.npz", allow_pickle=True)
dset_samples = to_list_of_arrays(dataset_samples['samples'])
cov_stat_samples = to_list_of_arrays(cov_static_samples['samples'])

# %% [code]
is_image = (np.asarray(dset_samples[0]).ndim == 4)
if is_image:
    _, C, H, W = np.asarray(dset_samples[0]).shape
T_min = min(arr.shape[0] for arr in dset_samples)
T_max = max(arr.shape[0] for arr in dset_samples)

# %% [code]
eps = 1e-1
padded = [np.pad(a, ((0, T_max-a.shape[0]), (0,0), (0,0), (0,0)), mode='constant', constant_values=np.nan)
          for a in dset_samples]                                              # -> (n_trial, T_max, C, H, W)
stacked = np.stack(padded, axis=0)
min_img = np.nanmin(stacked, axis=(0,1))                                      # shape (C,H,W)
max_img = np.nanmax(stacked, axis=(0,1))                                      # shape (C,H,W)
clamp_range = np.stack([min_img - eps, max_img + eps], axis=0).astype(np.float32)  # shape (2,C,H,W)

# %% [code]
TRIAL_IDX = 0
T = T_max

# %% [code]
data_flattent = [np.reshape(a, (a.shape[0], -1), order='C') for a in dset_samples]
data_flattent_all = np.concatenate(data_flattent, axis=0)

# %% [markdown]
# 2. Read Models

# %% [code]
folder_ref = 'out/ball/t_detach/00000-ball_dim50_lag0_none-prr-uncond-regularFM-gpus1-batch256-fp32'
folder = 'out/ball/t_detach/dim3_lag0/00004-ball_dim3_lag0_none-prr-uncond-regularFM-gpus1-batch256-fp32'

_, _, encoder_ref, _ = read_model(folder_ref, '/network-snapshot-004264.pkl')
flow_net4, dyn_net4, encoder4, lag = read_model(folder, '/network-snapshot-004766.pkl')

apply_posthoc_alignment(encoder4, encoder_ref)

# %% [code]
print("k_max =", encoder4.k_max, "use_pop_stats =", encoder4.use_pop_stats)

# %% [code]
# check reconstruction
trial_check = 0
T_check = 0
X0_back4 = recon_check_x0(encoder4, trial_check, T_check, dset_samples = dset_samples)

# %% [code]
fig, axs = plt.subplots(1, 2, figsize=(6, 5))

im1 = axs[0].imshow(X0_back4)
fig.colorbar(im1, ax=axs[0], fraction=0.046, pad=0.04)  # smaller, aligned
axs[0].set_title('encoder recon')

im2 = axs[1].imshow(dset_samples[trial_check][T_check,0,:,:])
fig.colorbar(im2, ax=axs[1], fraction=0.046, pad=0.04)  # same settings
axs[1].set_title('training data')

plt.tight_layout()
plt.show()

# %% [markdown]
# 3. Simulate Trajectories

# %% [code]
TRIAL_IDX = 0
n_step = T_max

# %% [markdown]
## 3.1 Data space (tau = 1.0)

# %% [code]
tau = 1.0
traj_sim_tau1_flattent, lag_cov_list = generate_traj_image_onthefly(dyn_net4.to(device),
                                                                    dset_samples, n_step,
                                                                    tau, device, lag,
                                                                    include_x0_tau=False,
                                                                    oracle=False,
                                                                    flow_net=None,
                                                                    clamp_range=clamp_range
                                                                    )

# %% [code]
out_imgs_tau1 = [arr.reshape(arr.shape[0], C, H, W) for arr in traj_sim_tau1_flattent]

# %% [code]
plot_tau(n_step, out_imgs_tau1, dset_samples, trial_idx = TRIAL_IDX, tau = 1.0)

# %% [markdown]
## 3.2 Latent space (tau = 0.0)

# %% [code]
out_imgs_tau0_flow_flatten = proj_to_latent(out_imgs_tau1, flow_net4.to(device), 0.0, device)
out_imgs_tau0_flow = [arr.reshape(arr.shape[0], C, H, W) for arr in out_imgs_tau0_flow_flatten ]

# %% [code]
tau = 0.0
traj_sim_tau0_flattent, _ = generate_traj_image_onthefly(dyn_net4.to(device),
                                                         dset_samples, n_step,
                                                         tau, device, lag,
                                                         include_x0_tau=False,
                                                         oracle=False,
                                                         flow_net=flow_net4.to(device),
                                                         clamp_range=clamp_range
                                                         )
out_imgs_tau0 = [arr.reshape(arr.shape[0], C, H, W) for arr in traj_sim_tau0_flattent ]

# %% [code]
plot_tau(T, out_imgs_tau0_flow, dset_samples, trial_idx = TRIAL_IDX, tau = 0.0, title = 'porjected, tau = 0.0; flow from tau = 1.0 directly')

# %% [code]
plot_tau(T, out_imgs_tau0, dset_samples, trial_idx = TRIAL_IDX, tau = 0.0, title = 'simulated, tau = 0.0')

# %% [markdown]
# 4. Latent space

# %% [code]
data_flattent = [np.reshape(a, (a.shape[0], -1), order='C') for a in dset_samples]

# %% [markdown]
## 4.1 directly from encoder: mu(x_data)

# %% [code]
dim_to_keep = 3
mu_keep4 = get_mu_from_encoder(encoder4, data_flattent, dim_to_keep, TRIAL_IDX)
plot_latent_time_series([mu_keep4], alpha = 0.4, dims=np.arange(dim_to_keep), 
                        dt=0.01, labels=None)

# %% [code]
plot_traj_3d(mu_keep4)

# %% [code]
mu_tmp, d, L, loading, bias = latent_mu(dset_samples, encoder4.to(device), dim_to_keep, device, fullRes = True)
L_np = L.detach().cpu().numpy()
d_np = d.detach().cpu().numpy()
loading_np = loading.detach().cpu().numpy()
mu_all = get_mu_from_encoder(encoder4, data_flattent, 28*28, TRIAL_IDX)

# %% [code]
k_max = 200
k_target = 3
loading_norm_all = torch.norm(loading[:,0:k_max], dim= 0).detach().cpu().numpy()
energy = loading_norm_all ** 2 
cum_energy = np.cumsum(energy)
K_eff = 1 + np.searchsorted(cum_energy, 0.95 * cum_energy[-1])

# %% [code]
fig = plt.figure(figsize=(3, 3))
plt.plot(np.sqrt(d_np[0:k_max]))
plt.axvline(x=k_target-1, color='red', label = 'k_target')
plt.axvline(x=K_eff-1, color='orange', label = 'k_eff = ' + str(K_eff))
plt.title('sqrt{d}');
plt.legend()

# %% [markdown]
Basically, $\sqrt{d_i}$ is loading column norm

# %% [code]
k_max = 200
dim_check_most = 100
k_target = 3
rows = loading.shape[0]
cols = dim_check_most
dim_check = np.arange(dim_check_most)
rec = mu_group_recon(bias.detach().cpu().numpy(), loading_np, mu_all, dim_check, H=28, W=28)
ball_idx = 0

fig, axs = plt.subplots(2, 2, figsize=(6, 5))
axs[0, 0].plot(loading_norm_all[:dim_check_most])
axs[0, 0].axvline(x=k_target-1, color='red', label = 'k_target')
axs[0, 0].axvline(x=K_eff-1, color='orange', label = 'k_eff = ' + str(K_eff))
axs[0, 0].set_title('loading norm')
axs[0, 0].legend()

im1 = axs[0, 1].imshow(np.abs(loading_np[:, 0:dim_check_most]), cmap='gray_r', aspect=cols / rows)
axs[0, 1].set_title('|loading|')
fig.colorbar(im1, ax=axs[0, 1])

axs[1, 0].plot(mu_all[:, :dim_check_most])
axs[1, 0].set_title('first ' + str(dim_check_most) + " dim of mu")

# Plot 4: cumulative contribution
im2 = axs[1, 1].imshow(rec[ball_idx], cmap='gray')
axs[1, 1].set_title('cumulative contribution from ' + str(dim_check[0]) + ' to ' + str(dim_check[-1]))
fig.colorbar(im2, ax=axs[1, 1])

fig.suptitle('Encoder Diagnostic', fontsize=16)
plt.tight_layout()
plt.show()

# %% [code]
dim_to_keep = 3
latent_traj_list_enc = []
for ii in range(len(dset_samples)):
    latent_traj_list_enc.append(recon_check_x0_allT(encoder4, ii, dset_samples))
mu_traj_sim_enc, d, L, loading, bias = latent_mu(latent_traj_list_enc, encoder4.to(device),
                                                 dim_to_keep, device,
                                                 fullRes = True, resid_tol=1e-15)

# %% [code]
mu_pts_encoder, v_mu_encoder, v_mu_full_encoder = compute_mu_velocity_field(
    latent_traj_list=latent_traj_list_enc,
    dyn_net=dyn_net4,
    encoder=encoder4,
    device=device,
    dim_preserve=dim_to_keep,
    lag=lag,
    t_plot=0.0,
    return_full=True
)

# %% [code]
plot_mu_velocity_panel(mu_pts_encoder, v_mu_encoder,
                       trajectories=mu_traj_sim_enc,
                       title_2d = "encoder: mu1–mu2 field (tau =0.0)",
                       title_3d = "encoder: mu1–mu2-mu3 field (tau =0.0)",
                       speed_color=v_mu_full_encoder,
                       arrow_frac_2d=0.05,
                       arrow_size_2d=1.2,
                       linewidth_2d=1.5,
                       
                       arrow_frac_3d=0.1,
                       arrow_size_3d=1.5,
                       linewidth_3d=1.5,
                       base_arrow_length_ratio_3d=0.3,
                       )

# %% [markdown]
## 4.2 flowed/ project

# %% [code]
dim_to_keep = 3
latent_traj_list = proj_to_latent(dset_samples, flow_net4.to(device), 0.0, device)

# %% [code]
latent_traj_list_enc = []
for ii in range(len(dset_samples)):
    latent_traj_list_enc.append(recon_check_x0_allT(encoder4, ii, dset_samples))

# %% [code]
# mu_traj_sim, d, L, loading, bias = latent_mu(latent_traj_list_enc, encoder4.to(device), dim_to_keep, device, fullRes = True)
mu_traj_sim, d, L, loading, bias = latent_mu(latent_traj_list, encoder4.to(device), dim_to_keep, device,
                                             fullRes = True, resid_tol=1e-15)

# %% [code]
data_flow = [arr.reshape(arr.shape[0], C, H, W) for arr in latent_traj_list ]
data_enc = [arr.reshape(arr.shape[0], C, H, W) for arr in latent_traj_list_enc ]

# %% [code]
trial_check = 0
T_check = 0
fig, axs = plt.subplots(1, 3, figsize=(12, 10))

im1 = axs[0].imshow(data_flow[trial_check][T_check,0])
fig.colorbar(im1, ax=axs[0], fraction=0.046, pad=0.04)  # smaller, aligned
axs[0].set_title('flow(X)')

im2 = axs[1].imshow(data_enc[trial_check][T_check,0])
fig.colorbar(im2, ax=axs[1], fraction=0.046, pad=0.04)  # smaller, aligned
axs[1].set_title('encoder(X)')

im3 = axs[2].imshow(data_flow[trial_check][T_check,0] - data_enc[trial_check][T_check,0])
fig.colorbar(im3, ax=axs[2], fraction=0.046, pad=0.04)  # smaller, aligned
axs[2].set_title('flow(X) - encoder(X)')

plt.tight_layout()
plt.show()

# %% [code]
mu_traj_flow_np = [mu_traj_sim[j].cpu().numpy() for j in range(len(mu_traj_sim))]
plot_latent_time_series(mu_traj_flow_np, alpha = 0.4, dims=np.arange(dim_to_keep), 
                        dt=0.01, labels=None)
plt.suptitle('flowed latent trajc for each trial')

# %% [code]
plot_traj_3d(mu_traj_flow_np[TRIAL_IDX][:,0:3])

# %% [code]
latent_traj_list = proj_to_latent(dset_samples, flow_net4.to(device), 0.0, device)
mu_pts_flow, v_mu_flow, v_mu_full_flow = compute_mu_velocity_field(
    latent_traj_list=latent_traj_list,
    dyn_net=dyn_net4,
    encoder=encoder4,
    device=device,
    dim_preserve=dim_to_keep,
    lag=lag,
    t_plot=0.0,
    return_full=True
)

plot_mu_velocity_panel(mu_pts_flow, v_mu_flow,
                       trajectories = mu_traj_flow_np,
                       title_2d = "flowed: mu1–mu2 field (tau =0.0)",
                       title_3d = "flowed: mu1–mu2-mu3 field (tau =0.0)",
                       speed_color=v_mu_full_flow,
                       arrow_frac_2d=0.05,
                       arrow_size_2d=1.2,
                       linewidth_2d=1.5,
                       
                       arrow_frac_3d=0.1,
                       arrow_size_3d=1.5,
                       linewidth_3d=1.5,
                       base_arrow_length_ratio_3d=0.3,
                       )

# %% [markdown]
### simulated

# %% [code]
mu_traj_sim, d, L, loading, bias = latent_mu(traj_sim_tau0_flattent, encoder4.to(device), dim_to_keep, device, fullRes = True)
mu_traj_sim_np = [mu_traj_sim[j].cpu().numpy() for j in range(len(mu_traj_sim))]

# %% [code]
mu_pts_sim, v_mu_sim, v_mu_full_sim = compute_mu_velocity_field(
    latent_traj_list=traj_sim_tau0_flattent,
    dyn_net=dyn_net4,
    encoder=encoder4,
    device=device,
    dim_preserve=dim_to_keep,
    lag=lag,
    t_plot=0.0,
    return_full=True
)

plot_mu_velocity_panel(mu_pts_sim, v_mu_sim,
                       trajectories=mu_traj_sim_np,
                       title_2d = "simulated: mu1–mu2 field (tau =0.0)",
                       title_3d = "simulated: mu1–mu2-mu3 field (tau =0.0)",
                       speed_color=v_mu_full_sim,
                       arrow_frac_2d=0.05,
                       arrow_size_2d=1.2,
                       linewidth_2d=1.5,
                       
                       arrow_frac_3d=0.1,
                       arrow_size_3d=1.5,
                       linewidth_3d=1.5,
                       base_arrow_length_ratio_3d=0.3,
                       )

# %% [markdown]
## 4.3 simulated

# %% [code]
dim_to_keep = 3
mu_traj_sim, d, L, loading, bias = latent_mu(traj_sim_tau0_flattent, encoder4.to(device), dim_to_keep, device, fullRes = True)

# %% [code]
mu_traj_sim_np = [mu_traj_sim[j].cpu().numpy() for j in range(len(mu_traj_sim))]
mu_traj_sim_np_plot = [mu_traj_sim_np[ii][10:,:] for ii in range(len(mu_traj_sim_np))]
plot_latent_time_series(mu_traj_sim_np_plot, alpha = 0.4, dims=np.arange(dim_to_keep), 
                        dt=0.01, labels=None)
plt.suptitle('simulated latent trajc for each trial')

# %% [code]
plot_traj_3d(mu_traj_sim_np_plot[TRIAL_IDX][:,0:3])

# %% [code]
# latent_traj_list = proj_to_latent(dset_samples, flow_net4.to(device), 0.0, device)
# mu_traj_sim, d, L, loading, bias = latent_mu(latent_traj_list, encoder4.to(device), dim_to_keep, device,
#                                              fullRes = True, resid_tol=1e-15)

mu_pts_flow, v_mu_flow, v_mu_full_flow = compute_mu_velocity_field(
    latent_traj_list=latent_traj_list,
    dyn_net=dyn_net4,
    encoder=encoder4,
    device=device,
    dim_preserve=dim_to_keep,
    lag=lag,
    t_plot=0.0,
    return_full=True
)

plot_mu_velocity_panel(mu_pts_flow, v_mu_flow,
                       trajectories=mu_traj_flow_np, # mu_traj_sim
                       title_2d = "flowed: mu1–mu2 field (tau =0.0)",
                       title_3d = "flowed: mu1–mu2-mu3 field (tau =0.0)",
                       speed_color=v_mu_full_flow,
                       arrow_frac_2d=0.05,
                       arrow_size_2d=1.2,
                       linewidth_2d=1.5,
                       
                       arrow_frac_3d=0.1,
                       arrow_size_3d=1.5,
                       linewidth_3d=1.5,
                       base_arrow_length_ratio_3d=0.3,
                       )

# %% [markdown]
# 5. Plots

# %% [code]
plotFolder = "paper_fig/ball/app_dim3_lag0"

# %% [code]
# 1. encoder vs. flow
trial_check = 0
T_check = 0

A = dset_samples[trial_check][T_check, 0]
B = data_enc[trial_check][T_check, 0]
C = data_flow[trial_check][T_check, 0]

diff_enc_abs  = np.abs(A - B)
diff_flow_abs = np.abs(A - C)

eps = 1e-12
den = float(np.mean(np.abs(A))) + eps
rmae_enc  = float(np.mean(np.abs(A - B))) / den
rmae_flow = float(np.mean(np.abs(A - C))) / den

vmin = float(min(diff_enc_abs.min(), diff_flow_abs.min()))
vmax = float(max(diff_enc_abs.max(), diff_flow_abs.max()))

fig, axs = plt.subplots(1, 2, figsize=(5, 5), constrained_layout=True)

im0 = axs[0].imshow(diff_enc_abs, vmin=vmin, vmax=vmax)
axs[0].set_title(f"|X - encoder(X)|\nrMAE={rmae_enc:.3g}")

im1 = axs[1].imshow(diff_flow_abs, vmin=vmin, vmax=vmax)
axs[1].set_title(f"|X - flow(X)|\nrMAE={rmae_flow:.3g}")

for ax in axs.ravel():
    ax.set_xticks([]); ax.set_yticks([])

cbar = fig.colorbar(im1, ax=axs[1], fraction=0.046, pad=0.04)
cbar.set_label("abs diff")

plt.savefig(plotFolder + "/1_encoder_vs_flow.pdf")
plt.savefig(plotFolder + "/1_encoder_vs_flow.svg")
plt.savefig(plotFolder + "/1_encoder_vs_flow.png")

# %% [code]
elev=20
azim=-140

view = dict(elev=elev, azim=azim)

# %% [code]
# 2.2 compressed flow field in mu (3D)
fig, ax = plot_mu_velocity_3d(mu_pts_flow, v_mu_flow, trajectories=mu_traj_flow_np,
                    speed_color=v_mu_full_flow,
                    dims_3d=(0,1,2),
                    title="flowed: mu1–mu2-mu3 field (tau =0.0)", figsize=(5, 5))

ax.view_init(**view)

plt.savefig(plotFolder + "/2_flow_v_3d.pdf")
plt.savefig(plotFolder + "/2_flow_v_3d.svg")
plt.savefig(plotFolder + "/2_flow_v_3d.png")

# %% [code]
# 2.2 compressed flow field in mu (3D)
fig, ax = plot_mu_velocity_3d(mu_pts_sim, v_mu_sim, trajectories=mu_traj_sim_np, dims_3d=(0,1,2),
                    speed_color=v_mu_full_sim,
                    title="simulated: mu1–mu2-mu3 field (tau =0.0)", figsize=(5, 5))

ax.view_init(**view)

plt.savefig(plotFolder + "/3_sim_v_3d.pdf")
plt.savefig(plotFolder + "/3_sim_v_3d.svg")
plt.savefig(plotFolder + "/3_sim_v_3d.png")

# %% [code]
idx_select = np.arange(0, dset_samples[0].shape[0]//2, 5, dtype=int)

# %% [code]
trial_idx = 0
fig = plt.figure(figsize=(5, 5), constrained_layout=True)
ax3d = fig.add_subplot(111, projection='3d')
ax3d.set_box_aspect((1, 1, 1))
ax3d.grid(False)
for axis in (ax3d.xaxis, ax3d.yaxis, ax3d.zaxis):
    axis.pane.fill = False
    axis.pane.set_edgecolor((1, 1, 1, 0))
    axis._axinfo["grid"]["color"] = (1, 1, 1, 0)
    axis._axinfo["grid"]["linewidth"] = 0
    
# ax3d.scatter(mu_pts_flow[:, 0], mu_pts_flow[:, 1], mu_pts_flow[:, 2],
#              c="k", s=2, alpha=0.2)
for traj in mu_traj_flow_np:
    ax3d.plot(traj[:, 0], traj[:, 1], traj[:, 2],
              color="k", lw=1.2, alpha=0.1)

idx_sel = np.asarray(idx_select)
order = np.argsort(idx_sel)
idx_sel = idx_sel[order]

vals = idx_sel.astype(float)

sc = ax3d.scatter(mu_pts_flow[idx_sel, 0], mu_pts_flow[idx_sel, 1], mu_pts_flow[idx_sel, 2],
                  c=vals, cmap="plasma", s=50, alpha=1.0)


cbar = fig.colorbar(sc, ax=ax3d, fraction=0.03, pad=0.08)
cbar.set_label("selection order")
fig.subplots_adjust(right=0.86)
ax3d.set_title("flowed mu") 

ax3d.view_init(**view)

plt.savefig(plotFolder + "/4_1_sel_flow_v_3d.pdf")
plt.savefig(plotFolder + "/4_1_sel_flow_v_3d.svg")
plt.savefig(plotFolder + "/4_1_sel_flow_v_3d.png")

# %% [code]
out_imgs_tau0_flow_flatten = proj_to_latent(dset_samples, flow_net4.to(device), 0.0, device)
# out_imgs_tau0_flow = [arr.reshape(arr.shape[0], C, H, W) for arr in out_imgs_tau0_flow_flatten ]

# %% [code]
height1 = 1.3
plot_frame_compact(
    rows=[dset_samples, out_imgs_tau0_flow],
    row_labels=["data", "tau = 0.0"],
    trial_idx=trial_idx,
    idxs=idx_select,
    border_from=sc,         
    figsize=(6.8, height1*2),
    mean_img=None,
);

plt.savefig(plotFolder + "/4_2_sel_flow_img.pdf")
plt.savefig(plotFolder + "/4_2_sel_flow_img.svg")
plt.savefig(plotFolder + "/4_2_sel_flow_img.png")

# %% [code]
fig = plt.figure(figsize=(5, 5), constrained_layout=True)
ax3d = fig.add_subplot(111, projection='3d')
ax3d.set_box_aspect((1, 1, 1))
ax3d.grid(False)
for axis in (ax3d.xaxis, ax3d.yaxis, ax3d.zaxis):
    axis.pane.fill = False
    axis.pane.set_edgecolor((1, 1, 1, 0))
    axis._axinfo["grid"]["color"] = (1, 1, 1, 0)
    axis._axinfo["grid"]["linewidth"] = 0

# ax3d.scatter(mu_pts_sim[:, 0], mu_pts_sim[:, 1], mu_pts_sim[:, 2],
#              c="k", s=2, alpha=0.2)

for traj in mu_traj_sim_np:
    ax3d.plot(traj[:, 0], traj[:, 1], traj[:, 2],
              color="k", lw=1.2, alpha=0.1)

idx_sel = np.asarray(idx_select)
order = np.argsort(idx_sel)
idx_sel = idx_sel[order]

vals = idx_sel.astype(float)

sc = ax3d.scatter(mu_pts_sim[idx_sel, 0], mu_pts_sim[idx_sel, 1], mu_pts_sim[idx_sel, 2],
                  c=vals, cmap="plasma", s=50, alpha=1.0)

cbar = fig.colorbar(sc, ax=ax3d, fraction=0.03, pad=0.08)
cbar.set_label("selection order")
fig.subplots_adjust(right=0.86)
ax3d.set_title('simulated mu')

ax3d.view_init(**view)

plt.savefig(plotFolder + "/5_1_sel_sim_v_3d.pdf")
plt.savefig(plotFolder + "/5_1_sel_sim_v_3d.svg")
plt.savefig(plotFolder + "/5_1_sel_sim_v_3d.png")

# %% [code]
plot_frame_compact(
    rows=[dset_samples, out_imgs_tau1, out_imgs_tau0],
    row_labels=["data", "tau = 1.0", "tau = 0.0"],
    trial_idx=trial_idx,
    idxs=idx_select,
    border_from=sc,         
    figsize=(6.8, height1*3),
    mean_img=None,
);

plt.savefig(plotFolder + "/5_2_sel_sim_img.pdf")
plt.savefig(plotFolder + "/5_2_sel_sim_img.svg")
plt.savefig(plotFolder + "/5_2_sel_sim_img.png")

# %% [code]


# %% [code]


# %% [code]


# %% [code]


# %% [code]
```
