## Supplementary material for "Dynamic Compression Flows for Neuroscience Data": Python code: paper_plot_ball_penalties.ipynb

```
# %% [code]
import numpy as np
import matplotlib.pyplot as plt
from matplotlib.collections import LineCollection
from matplotlib.colors import ListedColormap, BoundaryNorm, LinearSegmentedColormap, Normalize
from mpl_toolkits.mplot3d.art3d import Line3DCollection
from matplotlib.cm import ScalarMappable
from matplotlib.colors import LogNorm
import matplotlib as mpl

import torch
import dnnlib.util_v7 as util
from tqdm import tqdm
import pickle
from matplotlib.ticker import FuncFormatter
import pickle, types
from torch_utils import persistence

from PIL import Image

import numpy as np
import pickle
import os
import re 
import json 
import click
import torch
from torch_utils import distributed as dist
from training import toy_training_loop_vfm_noDataGen
from tqdm import tqdm
import matplotlib.pyplot as plt
from matplotlib import colors
from plot_neural_v7 import *
from plot_sim_results_v7 import *
from plot_sim_results_img_v7 import * 


import pandas as pd

import warnings
warnings.filterwarnings('ignore', 'Grad strides do not match bucket view strides') # False warning 
device = torch.device("cuda" if torch.cuda.is_available() else "cpu")
torch.set_default_device(device)
colorgrad = "PuOr"


# import matplotlib as mpl

# mpl.rcParams.update({
#     "font.family": "serif",
#     "font.serif": ["Palatino", "Palatino Linotype", "Palatino LT STD", "TeX Gyre Pagella"],
#     "mathtext.fontset": "custom",
#     "mathtext.rm": "Palatino",
#     "mathtext.it": "Palatino:italic",
#     "mathtext.bf": "Palatino:bold",
# })


# %% [markdown]
# 0. Some functions...

# %% [code]
def plot_tau(T, out_imgs_tau, dset_samples, trial_idx, tau, title=None):
    # Plot True
    plot_frame(dset_samples, T, trial_idx)
    plt.suptitle(f'true, trial = {trial_idx}')
    plt.subplots_adjust(bottom=0.2)  # Make room for horizontal colorbar

    # Plot Simulated
    plot_frame(out_imgs_tau, T, trial_idx)
    if title is None:
        plt.suptitle(f'simulated- tau = {tau}')
    else:
        plt.suptitle(title)
    plt.subplots_adjust(bottom=0.2)

# %% [code]
def recon_check_x0(encoder, trial_check, T_check, dset_samples):
    encoder = encoder.to(device).eval()
    X1_ori = dset_samples[trial_check][T_check,0,:,:]
    data_flattent = [np.reshape(a, (a.shape[0], -1), order='C') for a in dset_samples]
    with torch.inference_mode():
        X1 = torch.from_numpy(data_flattent[trial_check][T_check,:]).to(device=device, dtype=torch.float32)
        X0 = encoder.rsample(X1)
    X0_back = X0.reshape(X0.shape[0], C, H, W)
    return X0_back[0,0,:,:].detach().cpu().numpy()

# %% [code]
def recon_check_x0_allT(encoder, trial_check, dset_samples):
    encoder = encoder.to(device).eval()
    X1_ori = dset_samples[trial_check][T_check,0,:,:]
    data_flattent = [np.reshape(a, (a.shape[0], -1), order='C') for a in dset_samples]
    with torch.inference_mode():
        X1 = torch.from_numpy(data_flattent[trial_check]).to(device=device, dtype=torch.float32)
        X0 = encoder.rsample(X1)
    X0_back = X0.reshape(X0.shape[0], C, H, W)
    return X0_back.detach().cpu().numpy()

# %% [code]
def mu_group_recon(bias,
                   loading_np: np.ndarray,
                   mu_all: np.ndarray,
                   mu_idx=None,             # int | list[int] | np.ndarray | slice | None
                   H: int = 28,
                   W: int = 28,
                   mean_img: np.ndarray = None,
                   clip=None):
    D = loading_np.shape[0]
    assert loading_np.ndim == 2
    K_load = loading_np.shape[1]          # number of latent dims in loading
    N, K_mu = mu_all.shape               # number of latent dims in mu_all
    assert H * W == D

    # normalize mu_idx to an integer index array IN LATENT SPACE
    if mu_idx is None:
        idx = np.arange(min(K_load, K_mu))
    elif isinstance(mu_idx, slice):
        idx = np.arange(min(K_load, K_mu))[mu_idx]
    else:
        arr = np.asarray(mu_idx)
        if arr.dtype == bool:
            idx = np.flatnonzero(arr)
        else:
            idx = arr.astype(int, copy=False).ravel()

    # sanity: all requested dims must exist in both loading and mu_all
    max_idx = int(idx.max()) if idx.size > 0 else -1
    assert max_idx < K_load, f"mu_idx uses dim {max_idx}, but loading has only {K_load} cols"
    assert max_idx < K_mu,   f"mu_idx uses dim {max_idx}, but mu_all has only {K_mu} cols"

    A_sub  = loading_np[:, idx]     # (D, K_sel)
    mu_sub = mu_all[:, idx]         # (N, K_sel)
    recon_flat = mu_sub @ A_sub.T   # (N, D)

    if mean_img is not None:
        mean_flat = mean_img.reshape(-1)
        assert mean_flat.size == D
        recon_flat = recon_flat + mean_flat

    recon_flat = recon_flat + bias
    recon = recon_flat.reshape(N, H, W)
    if clip is not None:
        lo, hi = clip
        recon = np.clip(recon, lo, hi)
    return recon


# %% [code]
def plot_mu_velocity_2d(
    mu,
    v_mu,
    trajectories=None,
    dims_2d=(0, 1),
    title=None,
    arrow_frac=0.05,
    arrow_size=1.2,
    linewidth=1.5,
    traj_alpha=0.4,
    traj_width=0.8,
    traj_color="gray",
    max_trajs=300,
    quiver_alpha=0.6,
    max_points=3000,
    figsize=(7, 6),
    speed_color=None,                      # NEW
    colorbar_label=r"$\|\dot{\mu}\|$",     # NEW
):
    fig, ax = plt.subplots(figsize=figsize, constrained_layout=True)
    ax.grid(False)

    traj_list = []
    if trajectories is not None:
        traj_source = trajectories
        for i, traj in enumerate(traj_source):
            if i >= max_trajs:
                break
            t_np = traj.detach().cpu().numpy() if hasattr(traj, "detach") else np.array(traj)
            traj_list.append(t_np)

    d1, d2 = dims_2d
    for t_np in traj_list:
        ax.plot(
            t_np[:, d1], t_np[:, d2],
            color=traj_color, alpha=traj_alpha, linewidth=traj_width, zorder=1
        )

    coords = mu[:, [d1, d2]]
    vecs   = v_mu[:, [d1, d2]]
    N = coords.shape[0]

    # speed_color preprocessing
    if speed_color is not None:
        speed_color = speed_color.detach().cpu().numpy() if hasattr(speed_color, "detach") else np.asarray(speed_color)
        speed_color = np.asarray(speed_color).reshape(-1)
        assert speed_color.shape[0] == mu.shape[0]

    if max_points is not None and N > max_points:
        idx = np.random.choice(N, size=max_points, replace=False)
        coords_2d = coords[idx]
        vecs_2d   = vecs[idx]
        speed_c   = speed_color[idx] if speed_color is not None else None
    else:
        coords_2d = coords
        vecs_2d   = vecs
        speed_c   = speed_color if speed_color is not None else None

    speed_proj = np.linalg.norm(vecs_2d, axis=1)
    eps = 1e-8
    dirs = vecs_2d / (speed_proj[:, None] + eps)

    x_min, x_max = coords_2d[:, 0].min(), coords_2d[:, 0].max()
    y_min, y_max = coords_2d[:, 1].min(), coords_2d[:, 1].max()
    span = max(x_max - x_min, y_max - y_min) + eps
    L = arrow_frac * span
    U = dirs[:, 0] * L
    V = dirs[:, 1] * L

    base_width = 0.004
    base_headwidth = 4.0
    base_headlength = 6.0
    base_headaxislength = 5.0
    width = base_width * arrow_size

    # coloring values
    if speed_c is None:
        speed_c = speed_proj

    norm = Normalize(vmin=float(speed_c.min()), vmax=float(speed_c.max()))
    q = ax.quiver(
        coords_2d[:, 0], coords_2d[:, 1], U, V, speed_c,
        cmap=cm.viridis, norm=norm, angles="xy", scale_units="xy", scale=1.0,
        width=width,
        headwidth=base_headwidth * arrow_size,
        headlength=base_headlength * arrow_size,
        headaxislength=base_headaxislength * arrow_size,
        linewidth=linewidth, alpha=quiver_alpha, zorder=2
    )
    cb = fig.colorbar(q, ax=ax, fraction=0.046, pad=0.04)
    cb.set_label(colorbar_label)

    ax.set_xlabel(fr"$\mu_{{{d1+1}}}$")
    ax.set_ylabel(fr"$\mu_{{{d2+1}}}$")
    if title:
        ax.set_title(title)
    ax.axis("equal")

    return fig, ax, q


def plot_mu_velocity_3d(
    mu,
    v_mu,
    trajectories=None,
    dims_3d=(0, 1, 2),
    title=None,
    arrow_frac=0.1,
    arrow_size=1.5,
    linewidth=1.5,
    base_arrow_length_ratio=0.3,
    traj_alpha=0.4,
    traj_width=0.8,
    quiver_alpha=0.6,
    traj_color="gray",
    max_trajs=300,
    max_points=3000,
    figsize=(10, 7),
    elev=None,
    azim=None,
    roll=None,
    speed_norm=None,
    speed_color=None,                      # NEW
    colorbar_label=r"$\|\dot{\mu}\|$",     # NEW
):
    fig = plt.figure(figsize=figsize, constrained_layout=True)
    ax = fig.add_subplot(1, 1, 1, projection="3d")
    ax.set_box_aspect((1, 1, 1))
    ax.grid(False)
    for axis in (ax.xaxis, ax.yaxis, ax.zaxis):
        axis.pane.fill = False
        axis.pane.set_edgecolor((1, 1, 1, 0))
        axis._axinfo["grid"]["color"] = (1, 1, 1, 0)
        axis._axinfo["grid"]["linewidth"] = 0

    traj_list = []
    if trajectories is not None:
        traj_source = trajectories
        for i, traj in enumerate(traj_source):
            if i >= max_trajs:
                break
            t_np = traj.detach().cpu().numpy() if hasattr(traj, "detach") else np.array(traj)
            traj_list.append(t_np)

    d1, d2, d3 = dims_3d
    for t_np in traj_list:
        ax.plot(
            t_np[:, d1], t_np[:, d2], t_np[:, d3],
            color=traj_color, alpha=traj_alpha, linewidth=traj_width, zorder=1
        )

    coords = mu[:, [d1, d2, d3]]
    vecs   = v_mu[:, [d1, d2, d3]]
    N = coords.shape[0]

    # speed_color preprocessing
    if speed_color is not None:
        speed_color = speed_color.detach().cpu().numpy() if hasattr(speed_color, "detach") else np.asarray(speed_color)
        speed_color = np.asarray(speed_color).reshape(-1)
        assert speed_color.shape[0] == mu.shape[0]

    if max_points is not None and N > max_points:
        idx = np.random.choice(N, size=max_points, replace=False)
        coords_3d = coords[idx]
        vecs_3d   = vecs[idx]
        speed_c   = speed_color[idx] if speed_color is not None else None
    else:
        coords_3d = coords
        vecs_3d   = vecs
        speed_c   = speed_color if speed_color is not None else None

    speed_proj = np.linalg.norm(vecs_3d, axis=1)
    eps = 1e-8
    dirs = vecs_3d / (speed_proj[:, None] + eps)

    mins = coords_3d.min(axis=0)
    maxs = coords_3d.max(axis=0)
    span = float(np.max(maxs - mins) + eps)

    L = arrow_frac * span
    U, V, W = dirs[:, 0], dirs[:, 1], dirs[:, 2]

    # coloring values
    if speed_c is None:
        speed_c = speed_proj

    norm = speed_norm if speed_norm is not None else Normalize(
        vmin=float(speed_c.min()),
        vmax=float(speed_c.max()),
    )
    colors = cm.viridis(norm(speed_c))
    colors[:, 3] = quiver_alpha

    ax.quiver(
        coords_3d[:, 0], coords_3d[:, 1], coords_3d[:, 2],
        U, V, W,
        length=L, normalize=False, colors=colors,
        linewidth=linewidth * arrow_size,
        arrow_length_ratio=base_arrow_length_ratio * arrow_size,
        zorder=2
    )

    mappable = cm.ScalarMappable(norm=norm, cmap="viridis")
    mappable.set_array(speed_c)
    cb = fig.colorbar(mappable, ax=ax, fraction=0.035, pad=0.08)
    cb.set_label(colorbar_label)

    ax.set_xlabel(fr"$\mu_{{{d1+1}}}$")
    ax.set_ylabel(fr"$\mu_{{{d2+1}}}$")
    ax.set_zlabel(fr"$\mu_{{{d3+1}}}$")
    if title:
        ax.set_title(title)

    if (elev is not None) or (azim is not None) or (roll is not None):
        try:
            ax.view_init(elev=elev, azim=azim, roll=roll)
        except TypeError:
            ax.view_init(elev=elev, azim=azim)

    return fig, ax

# %% [code]
def plot_frame_compact(
    rows,                 # list like [dset_samples, out_imgs_tau0_flow, ...]
    trial_idx,
    idxs,                 # 1D int array, columns
    row_labels=None,      # list of strings, one per row
    mean_img=None,
    figsize=(6, 2.6),
    img_cmap="gray",
    vmin=None,
    vmax=None,
    border_from=None,
    border_lw=3.0,
    left=0.10, right=0.99, bottom=0.05, top=0.95,
    wspace=0.02, hspace=0.02,
):
    import numpy as np
    import matplotlib.pyplot as plt

    idxs = np.asarray(idxs).astype(int)
    n_rows = len(rows)
    n_cols = len(idxs)

    # --- global vmin/vmax (consistent across rows) ---
    if vmin is None or vmax is None:
        mn, mx = np.inf, -np.inf
        for data in rows:
            X = data[trial_idx]  # [T, 1, H, W] (torch or np)
            for i in idxs:
                img = X[i, 0]
                if hasattr(img, "detach"):
                    img = img.detach().cpu().numpy()
                else:
                    img = np.asarray(img)
                if mean_img is not None:
                    img = img + mean_img
                mn = min(mn, float(img.min()))
                mx = max(mx, float(img.max()))
        if vmin is None: vmin = mn
        if vmax is None: vmax = mx

    if border_from is not None:
        cmap = border_from.get_cmap()
        norm = border_from.norm
        col_colors = cmap(norm(idxs.astype(float)))
    else:
        col_colors = ["k"] * n_cols

    fig, axes = plt.subplots(n_rows, n_cols, figsize=figsize, squeeze=False)
    fig.subplots_adjust(left=left, right=right, bottom=bottom, top=top, wspace=wspace, hspace=hspace)

    for r, data in enumerate(rows):
        X = data[trial_idx]
        for c, i in enumerate(idxs):
            ax = axes[r, c]

            img = X[i, 0]
            if hasattr(img, "detach"):
                img = img.detach().cpu().numpy()
            else:
                img = np.asarray(img)
            if mean_img is not None:
                img = img + mean_img

            ax.imshow(img, cmap=img_cmap, vmin=vmin, vmax=vmax)

            # keep spines for colored border, so don't ax.axis("off")
            ax.set_xticks([]); ax.set_yticks([])

            # titles only on first row
            if r == 0:
                ax.set_title(f"t={i}")
            else:
                ax.set_title("")

            # colored border per column
            for sp in ax.spines.values():
                sp.set_visible(True)
                sp.set_linewidth(border_lw)
                sp.set_color(col_colors[c])

    # row labels on the left, centered per row (compact)
    if row_labels is not None:
        for r, lab in enumerate(row_labels):
            pos = axes[r, 0].get_position()
            y = 0.5 * (pos.y0 + pos.y1)
            x = pos.x0 - 0.02
            fig.text(x, y, lab, ha="right", va="center")

    return fig, axes

# %% [markdown]
# 1. Read Data

# %% [code]
folder_data = 'out/ball/t_detach/data'
useCov = False

dataset_samples = np.load(folder_data + "/dataset_samples.npz", allow_pickle=True)
cov_static_samples = np.load(folder_data + "/cov_static_trunc_only.npz", allow_pickle=True)
dset_samples = to_list_of_arrays(dataset_samples['samples'])
cov_stat_samples = to_list_of_arrays(cov_static_samples['samples'])

# %% [code]
is_image = (np.asarray(dset_samples[0]).ndim == 4)
if is_image:
    _, C, H, W = np.asarray(dset_samples[0]).shape
T_min = min(arr.shape[0] for arr in dset_samples)
T_max = max(arr.shape[0] for arr in dset_samples)

# %% [code]
eps = 1e-2
padded = [np.pad(a, ((0, T_max-a.shape[0]), (0,0), (0,0), (0,0)), mode='constant', constant_values=np.nan)
          for a in dset_samples]                                              # -> (n_trial, T_max, C, H, W)
stacked = np.stack(padded, axis=0)
min_img = np.nanmin(stacked, axis=(0,1))                                      # shape (C,H,W)
max_img = np.nanmax(stacked, axis=(0,1))                                      # shape (C,H,W)
clamp_range = np.stack([min_img - eps, max_img + eps], axis=0).astype(np.float32)  # shape (2,C,H,W)

# %% [code]
TRIAL_IDX = 0
T = T_max

# %% [code]
data_flattent = [np.reshape(a, (a.shape[0], -1), order='C') for a in dset_samples]
data_flattent_all = np.concatenate(data_flattent, axis=0)

# %% [markdown]
# 2. Read Models

# %% [code]
folder_ref = 'out/ball/t_detach/00000-ball_dim50_lag0_none-prr-uncond-regularFM-gpus1-batch256-fp32'
folder1 = 'out/ball/t_detach/00000-ball_dim50_lag0_none-prr-uncond-regularFM-gpus1-batch256-fp32'
folder2 = 'out/ball/t_detach/00001-ball_dim50_lag0_ridge-prr-uncond-regularFM-gpus1-batch256-fp32'
folder3 = 'out/ball/t_detach/00013-ball_dim50_lag0_lasso-prr-uncond-regularFM-gpus1-batch256-fp32'
folder4 = 'out/ball/t_detach/00003-ball_dim50_lag0_horseshoe-prr-uncond-regularFM-gpus1-batch256-fp32'

_, _, encoder_ref, _ = read_model(folder_ref, '/network-snapshot-004264.pkl')

flow_net1, dyn_net1, encoder1, lag = read_model(folder1, '/network-snapshot-004264.pkl')
flow_net2, dyn_net2, encoder2, lag = read_model(folder2, '/network-snapshot-004264.pkl')
flow_net3, dyn_net3, encoder3, lag = read_model(folder3, '/network-snapshot-004264.pkl')
flow_net4, dyn_net4, encoder4, lag = read_model(folder4, '/network-snapshot-004264.pkl')

apply_posthoc_alignment(encoder1, encoder_ref)
apply_posthoc_alignment(encoder2, encoder_ref)
apply_posthoc_alignment(encoder3, encoder_ref)
apply_posthoc_alignment(encoder4, encoder_ref)

# %% [code]
dim_to_keep = 3
_, d1, _, _, _ = latent_mu(dset_samples, encoder1.to(device), dim_to_keep, device, fullRes = True); d1_np = d1.detach().cpu().numpy()
_, d2, _, _, _ = latent_mu(dset_samples, encoder2.to(device), dim_to_keep, device, fullRes = True); d2_np = d2.detach().cpu().numpy()
_, d3, _, _, _ = latent_mu(dset_samples, encoder3.to(device), dim_to_keep, device, fullRes = True); d3_np = d3.detach().cpu().numpy()
_, d4, _, _, _ = latent_mu(dset_samples, encoder4.to(device), dim_to_keep, device, fullRes = True); d4_np = d4.detach().cpu().numpy()


# %% [code]
k_max = 200
k_target = 50

cum_energy1 = np.cumsum(d1_np); K_eff1 = 1 + np.searchsorted(cum_energy1, 0.95 * cum_energy1[-1])
cum_energy2 = np.cumsum(d2_np); K_eff2 = 1 + np.searchsorted(cum_energy2, 0.95 * cum_energy2[-1])
cum_energy3 = np.cumsum(d3_np); K_eff3 = 1 + np.searchsorted(cum_energy3, 0.95 * cum_energy3[-1])
cum_energy4 = np.cumsum(d4_np); K_eff4 = 1 + np.searchsorted(cum_energy4, 0.95 * cum_energy4[-1])

# %% [code]
print([K_eff1, K_eff2, K_eff3, K_eff4])

# %% [code]
fig = plt.figure(figsize=(3, 3))
plt.plot(np.sqrt(d1_np[0:k_max]))
plt.axvline(x=k_target-1, color='red', label = 'k_target')
plt.axvline(x=K_eff1-1, color='orange', label = 'k_eff = ' + str(K_eff1))
plt.title('None');
plt.legend()

# %% [code]
tau = 0.0
n_step = T_max

traj_sim_tau0_flattent1, _ = generate_traj_image_onthefly(dyn_net1.to(device),
                                                         dset_samples, n_step,
                                                         tau, device, lag,
                                                         include_x0_tau=False,
                                                         oracle=False,
                                                         flow_net=flow_net1.to(device),
                                                         clamp_range=clamp_range
                                                         )

traj_sim_tau0_flattent2, _ = generate_traj_image_onthefly(dyn_net2.to(device),
                                                         dset_samples, n_step,
                                                         tau, device, lag,
                                                         include_x0_tau=False,
                                                         oracle=False,
                                                         flow_net=flow_net2.to(device),
                                                         clamp_range=clamp_range
                                                         )

traj_sim_tau0_flattent3, _ = generate_traj_image_onthefly(dyn_net3.to(device),
                                                         dset_samples, n_step,
                                                         tau, device, lag,
                                                         include_x0_tau=False,
                                                         oracle=False,
                                                         flow_net=flow_net3.to(device),
                                                         clamp_range=clamp_range
                                                         )

traj_sim_tau0_flattent4, _ = generate_traj_image_onthefly(dyn_net4.to(device),
                                                         dset_samples, n_step,
                                                         tau, device, lag,
                                                         include_x0_tau=False,
                                                         oracle=False,
                                                         flow_net=flow_net4.to(device),
                                                         clamp_range=clamp_range
                                                         )

# %% [markdown]
# Plot

# %% [code]
plotFolder = "paper_fig/ball/app_penalties"

# %% [code]
fig, axs = plt.subplots(1, 4, figsize=(12, 3), constrained_layout=True, sharey=True)


axs[0].plot(np.sqrt(d1_np[0:k_max]))
axs[0].axvline(x=k_target-1, color="red", label="k_target")
axs[0].axvline(x=K_eff1-1, color="orange", label=f"k_eff = {K_eff1}")
axs[0].set_title("none")
axs[0].legend()

axs[1].plot(np.sqrt(d2_np[0:k_max]))
axs[1].axvline(x=k_target-1, color="red")
axs[1].axvline(x=K_eff2-1, color="orange", label=f"k_eff = {K_eff2}")
axs[1].set_title("ridge")
axs[1].legend()

axs[2].plot(np.sqrt(d3_np[0:k_max]))
axs[2].axvline(x=k_target-1, color="red")
axs[2].axvline(x=K_eff3-1, color="orange", label=f"k_eff = {K_eff3}")
axs[2].set_title("LASSO")
axs[2].legend()

axs[3].plot(np.sqrt(d4_np[0:k_max]))
axs[3].axvline(x=k_target-1, color="red")
axs[3].axvline(x=K_eff4-1, color="orange", label=f"k_eff = {K_eff4}")
axs[3].set_title("horseshoe")
axs[3].legend()

ymin = min(ax.get_ylim()[0] for ax in axs)
ymax = max(ax.get_ylim()[1] for ax in axs)
for ax in axs:
    ax.set_ylim(ymin, ymax)

plt.savefig(plotFolder + "/1_sqrt_d.pdf")
plt.savefig(plotFolder + "/1_sqrt_d.png")

# %% [code]
mu_traj_sim1, _, _, _, _ = latent_mu(traj_sim_tau0_flattent1,
                                    encoder1.to(device), dim_to_keep, device, fullRes = True)
mu_traj_sim2, _, _, _, _ = latent_mu(traj_sim_tau0_flattent2,
                                    encoder2.to(device), dim_to_keep, device, fullRes = True)
mu_traj_sim3, _, _, _, _ = latent_mu(traj_sim_tau0_flattent3,
                                    encoder3.to(device), dim_to_keep, device, fullRes = True)
mu_traj_sim4, _, _, _, _ = latent_mu(traj_sim_tau0_flattent4,
                                    encoder4.to(device), dim_to_keep, device, fullRes = True)

mu_traj_sim1_np = [mu_traj_sim1[j].cpu().numpy() for j in range(len(mu_traj_sim1))]
mu_traj_sim2_np = [mu_traj_sim2[j].cpu().numpy() for j in range(len(mu_traj_sim2))]
mu_traj_sim3_np = [mu_traj_sim3[j].cpu().numpy() for j in range(len(mu_traj_sim3))]
mu_traj_sim4_np = [mu_traj_sim4[j].cpu().numpy() for j in range(len(mu_traj_sim4))]

# %% [code]


# %% [code]
mu_pts_sim1, v_mu_sim1, v_mu_full_sim1 = compute_mu_velocity_field(
    latent_traj_list=traj_sim_tau0_flattent1,
    dyn_net=dyn_net1,
    encoder=encoder1,
    device=device,
    dim_preserve=dim_to_keep,
    lag=lag,
    t_plot=0.0,
    return_full=True
)

# %% [code]
mu_pts_sim2, v_mu_sim2, v_mu_full_sim2 = compute_mu_velocity_field(
    latent_traj_list=traj_sim_tau0_flattent2,
    dyn_net=dyn_net2,
    encoder=encoder2,
    device=device,
    dim_preserve=dim_to_keep,
    lag=lag,
    t_plot=0.0,
    return_full=True
)

# %% [code]
mu_pts_sim3, v_mu_sim3, v_mu_full_sim3 = compute_mu_velocity_field(
    latent_traj_list=traj_sim_tau0_flattent3,
    dyn_net=dyn_net3,
    encoder=encoder3,
    device=device,
    dim_preserve=dim_to_keep,
    lag=lag,
    t_plot=0.0,
    return_full=True
)

# %% [code]
mu_pts_sim4, v_mu_sim4, v_mu_full_sim4 = compute_mu_velocity_field(
    latent_traj_list=traj_sim_tau0_flattent4,
    dyn_net=dyn_net4,
    encoder=encoder4,
    device=device,
    dim_preserve=dim_to_keep,
    lag=lag,
    t_plot=0.0,
    return_full=True
)

# %% [code]
dim_to_keep = 3
latent_traj_list_1 = proj_to_latent(dset_samples, flow_net1.to(device), 0.0, device)
latent_traj_list_2 = proj_to_latent(dset_samples, flow_net2.to(device), 0.0, device)
latent_traj_list_3 = proj_to_latent(dset_samples, flow_net3.to(device), 0.0, device)
latent_traj_list_4 = proj_to_latent(dset_samples, flow_net4.to(device), 0.0, device)

mu_traj_flow1, _, _, _, _ = latent_mu(latent_traj_list_1, encoder1.to(device),
                                      dim_to_keep, device,
                                      fullRes = True, resid_tol=1e-15)
mu_traj_flow2, _, _, _, _ = latent_mu(latent_traj_list_2, encoder2.to(device),
                                      dim_to_keep, device,
                                      fullRes = True, resid_tol=1e-15)
mu_traj_flow3, _, _, _, _ = latent_mu(latent_traj_list_3, encoder3.to(device),
                                      dim_to_keep, device,
                                      fullRes = True, resid_tol=1e-15)
mu_traj_flow4, _, _, _, _ = latent_mu(latent_traj_list_4, encoder4.to(device),
                                      dim_to_keep, device,
                                      fullRes = True, resid_tol=1e-15)


mu_traj_flow1_np = [mu_traj_flow1[j].cpu().numpy() for j in range(len(mu_traj_flow1))]
mu_traj_flow2_np = [mu_traj_flow2[j].cpu().numpy() for j in range(len(mu_traj_flow2))]
mu_traj_flow3_np = [mu_traj_flow3[j].cpu().numpy() for j in range(len(mu_traj_flow3))]
mu_traj_flow4_np = [mu_traj_flow4[j].cpu().numpy() for j in range(len(mu_traj_flow4))]

# %% [code]
mu_pts_flow1, v_mu_flow1, v_mu_full_flow1 = compute_mu_velocity_field(
    latent_traj_list=latent_traj_list_1,
    dyn_net=dyn_net1,
    encoder=encoder1,
    device=device,
    dim_preserve=dim_to_keep,
    lag=lag,
    t_plot=0.0,
    return_full=True
)

mu_pts_flow2, v_mu_flow2, v_mu_full_flow2 = compute_mu_velocity_field(
    latent_traj_list=latent_traj_list_2,
    dyn_net=dyn_net2,
    encoder=encoder2,
    device=device,
    dim_preserve=dim_to_keep,
    lag=lag,
    t_plot=0.0,
    return_full=True
)

mu_pts_flow3, v_mu_flow3, v_mu_full_flow3 = compute_mu_velocity_field(
    latent_traj_list=latent_traj_list_3,
    dyn_net=dyn_net3,
    encoder=encoder3,
    device=device,
    dim_preserve=dim_to_keep,
    lag=lag,
    t_plot=0.0,
    return_full=True
)

mu_pts_flow4, v_mu_flow4, v_mu_full_flow4 = compute_mu_velocity_field(
    latent_traj_list=latent_traj_list_4,
    dyn_net=dyn_net4,
    encoder=encoder4,
    device=device,
    dim_preserve=dim_to_keep,
    lag=lag,
    t_plot=0.0,
    return_full=True
)

# %% [code]
d1, d2, d3 = (0, 1, 2)

all_mu = np.concatenate([
    np.asarray(mu_pts_sim1)[:, [d1, d2, d3]],
    np.asarray(mu_pts_sim2)[:, [d1, d2, d3]],
    np.asarray(mu_pts_sim3)[:, [d1, d2, d3]],
    np.asarray(mu_pts_sim4)[:, [d1, d2, d3]],
], axis=0)

mins = all_mu.min(axis=0)
maxs = all_mu.max(axis=0)
pad  = 0.02 * (maxs - mins + 1e-8)

xlim = (mins[0] - pad[0], maxs[0] + pad[0])
ylim = (mins[1] - pad[1], maxs[1] + pad[1])
zlim = (mins[2] - pad[2], maxs[2] + pad[2])


elev=20
azim=-140

view = dict(elev=elev, azim=azim)  # pick whatever you like, but keep it fixed


# d1, d2, d3 = dims

all_speed_colors_raw = [
    v_mu_full_sim1, v_mu_full_sim2, v_mu_full_sim3, v_mu_full_sim4,
    v_mu_full_flow1, v_mu_full_flow2, v_mu_full_flow3, v_mu_full_flow4,
]

all_speed_colors = []
for sc in all_speed_colors_raw:
    sc = sc.detach().cpu().numpy() if hasattr(sc, "detach") else np.asarray(sc)

    # if user accidentally passes full velocity vectors (N, D_full), convert to norm (N,)
    if sc.ndim == 2:
        sc = np.linalg.norm(sc, axis=1)

    sc = np.asarray(sc).reshape(-1)
    all_speed_colors.append(sc)

vmin = float(min(s.min() for s in all_speed_colors))
vmax = float(max(s.max() for s in all_speed_colors))
shared_norm = Normalize(vmin=vmin, vmax=vmax)


# %% [code]
fig, ax = plot_mu_velocity_3d(mu_pts_sim1, v_mu_sim1, trajectories=mu_traj_sim1_np,
                              speed_color=v_mu_full_sim1,
                              dims_3d=(0,1,2), title="simulated (none)",
                              figsize=(5, 5), quiver_alpha=0.5,
                              speed_norm=shared_norm)
ax.set_xlim(xlim); ax.set_ylim(ylim); ax.set_zlim(zlim)
ax.view_init(**view)
plt.savefig(plotFolder + "/2_1_sim_v_3d_none.pdf")
plt.savefig(plotFolder + "/2_1_sim_v_3d_none.png")

# %% [code]
fig, ax = plot_mu_velocity_3d(mu_pts_sim2, v_mu_sim2,
                    speed_color=v_mu_full_sim2,
                    trajectories=mu_traj_sim2_np, dims_3d=(0,1,2),
                    title="simulated (ridge)", figsize=(5, 5), quiver_alpha=0.5,
                              speed_norm=shared_norm)
ax.set_xlim(xlim); ax.set_ylim(ylim); ax.set_zlim(zlim)
ax.view_init(**view)
plt.savefig(plotFolder + "/2_2_sim_v_3d_ridge.pdf")
plt.savefig(plotFolder + "/2_2_sim_v_3d_ridge.png")

# %% [code]
fig, ax = plot_mu_velocity_3d(mu_pts_sim3, v_mu_sim3, trajectories=mu_traj_sim3_np, dims_3d=(0,1,2),
                    speed_color=v_mu_full_sim3,
                    title="simulated (LASSO)", figsize=(5, 5), quiver_alpha=0.5,
                              speed_norm=shared_norm)
ax.set_xlim(xlim); ax.set_ylim(ylim); ax.set_zlim(zlim)
ax.view_init(**view)
plt.savefig(plotFolder + "/2_3_sim_v_3d_lasso.pdf")
plt.savefig(plotFolder + "/2_3_sim_v_3d_lasso.png")

# %% [code]
fig, ax = plot_mu_velocity_3d(mu_pts_sim4, v_mu_sim4, trajectories=mu_traj_sim4_np, dims_3d=(0,1,2),
                    speed_color=v_mu_full_sim4,
                    title="simulated (horseshoe)", figsize=(5, 5), quiver_alpha=0.5,
                              speed_norm=shared_norm)
ax.set_xlim(xlim); ax.set_ylim(ylim); ax.set_zlim(zlim)
ax.view_init(**view)
plt.savefig(plotFolder + "/2_4_sim_v_3d_hs.pdf")
plt.savefig(plotFolder + "/2_4_sim_v_3d_hs.png")

# %% [markdown]
# Add flowed version

# %% [code]
fig, ax = plot_mu_velocity_3d(mu_pts_flow1, v_mu_flow1, trajectories=mu_traj_flow1_np, dims_3d=(0,1,2),
                    speed_color=v_mu_full_flow1,
                    title="flowed: (none)", figsize=(5, 5), quiver_alpha=0.5,
                              speed_norm=shared_norm)
ax.set_xlim(xlim); ax.set_ylim(ylim); ax.set_zlim(zlim)
ax.view_init(**view)
plt.savefig(plotFolder + "/3_1_flow_v_3d_none.pdf")
plt.savefig(plotFolder + "/3_1_flow_v_3d_none.png")

# %% [code]
fig, ax = plot_mu_velocity_3d(mu_pts_flow2, v_mu_flow2, trajectories=mu_traj_flow2_np, dims_3d=(0,1,2),
                    speed_color=v_mu_full_flow2,
                    title="flowed: (ridge)", figsize=(5, 5), quiver_alpha=0.5,
                              speed_norm=shared_norm)
ax.set_xlim(xlim); ax.set_ylim(ylim); ax.set_zlim(zlim)
ax.view_init(**view)
plt.savefig(plotFolder + "/3_2_flow_v_3d_ridge.pdf")
plt.savefig(plotFolder + "/3_2_flow_v_3d_ridge.png")

# %% [code]
fig, ax = plot_mu_velocity_3d(mu_pts_flow3, v_mu_flow3, trajectories=mu_traj_flow3_np, dims_3d=(0,1,2),
                    speed_color=v_mu_full_flow3,
                    title="flowed: (LASSO)", figsize=(5, 5), quiver_alpha=0.5,
                              speed_norm=shared_norm)
ax.set_xlim(xlim); ax.set_ylim(ylim); ax.set_zlim(zlim)
ax.view_init(**view)
plt.savefig(plotFolder + "/3_3_flow_v_3d_lasso.pdf")
plt.savefig(plotFolder + "/3_3_flow_v_3d_lasso.png")

# %% [code]
fig, ax = plot_mu_velocity_3d(mu_pts_flow4, v_mu_flow4, trajectories=mu_traj_flow4_np, dims_3d=(0,1,2),
                    speed_color=v_mu_full_flow4,
                    title="flowed: (horseshoe)", figsize=(5, 5), quiver_alpha=0.5,
                              speed_norm=shared_norm)
ax.set_xlim(xlim); ax.set_ylim(ylim); ax.set_zlim(zlim)
ax.view_init(**view)
plt.savefig(plotFolder + "/3_4_flow_v_3d_hs.pdf")
plt.savefig(plotFolder + "/3_4_flow_v_3d_hs.png")

# %% [code]


# %% [code]


# %% [code]
```
