## Supplementary material for "Dynamic Compression Flows for Neuroscience Data": Python code: paper_plot_bird.ipynb

```
# %% [code]
import numpy as np
import matplotlib.pyplot as plt
from matplotlib.collections import LineCollection
from matplotlib.colors import ListedColormap, BoundaryNorm, LinearSegmentedColormap, Normalize
from mpl_toolkits.mplot3d.art3d import Line3DCollection
import torch
import dnnlib.util_v5 as util
from tqdm import tqdm
import pickle
from matplotlib.ticker import FuncFormatter
import gc
from PIL import Image

import numpy as np
import pickle
import os
import re 
import json 
import click
import torch
from torch_utils import distributed as dist
from training import toy_training_loop_vfm_noDataGen
from tqdm import tqdm
import matplotlib.pyplot as plt
from matplotlib import colors
from plot_neural import *
from plot_sim_results_v7 import *
from plot_sim_results_img_v7 import * 
# from plot_musal_v5 import * 
import pickle, types
from torch_utils import persistence

import pandas as pd

import warnings
warnings.filterwarnings('ignore', 'Grad strides do not match bucket view strides') # False warning 
device = torch.device("cuda" if torch.cuda.is_available() else "cpu")
torch.set_default_device(device)
colorgrad = "PuOr"

# %% [code]
def plot_tau(T, out_imgs_tau, dset_samples, trial_idx, tau, title=None):
    # Plot True
    plot_frame(dset_samples, T, trial_idx)
    plt.suptitle(f'true, trial = {trial_idx}')
    plt.subplots_adjust(bottom=0.2)  # Make room for horizontal colorbar

    # Plot Simulated
    plot_frame(out_imgs_tau, T, trial_idx)
    if title is None:
        plt.suptitle(f'simulated- tau = {tau}')
    else:
        plt.suptitle(title)
    plt.subplots_adjust(bottom=0.2)

# %% [code]
def recon_check_x0(encoder, trial_check, T_check, dset_samples):
    encoder = encoder.to(device).eval()
    X1_ori = dset_samples[trial_check][T_check,0,:,:]
    data_flattent = [np.reshape(a, (a.shape[0], -1), order='C') for a in dset_samples]
    with torch.inference_mode():
        X1 = torch.from_numpy(data_flattent[trial_check][T_check,:]).to(device=device, dtype=torch.float32)
        X0 = encoder.rsample(X1)
    X0_back = X0.reshape(X0.shape[0], C, H, W)
    return X0_back[0,0,:,:].detach().cpu().numpy()

# %% [code]
def recon_check_x0_allT(encoder, trial_check, dset_samples):
    encoder = encoder.to(device).eval()
    X1_ori = dset_samples[trial_check][T_check,0,:,:]
    data_flattent = [np.reshape(a, (a.shape[0], -1), order='C') for a in dset_samples]
    with torch.inference_mode():
        X1 = torch.from_numpy(data_flattent[trial_check]).to(device=device, dtype=torch.float32)
        X0 = encoder.rsample(X1)
    X0_back = X0.reshape(X0.shape[0], C, H, W)
    return X0_back.detach().cpu().numpy()

# %% [markdown]
# 1. Read Data

# %% [code]
folder_data = 'out/bird_song_local_80/dimk_lag0_ori/t/none/data'
dataset_samples = np.load(folder_data + "/dataset_samples.npz", allow_pickle=True)
cov_static_samples = np.load(folder_data + "/cov_static_trunc_only.npz", allow_pickle=True)

dset_samples = to_list_of_arrays(dataset_samples['samples'])
cov_stat_samples = to_list_of_arrays(cov_static_samples['samples'])
# mean_img = np.load('data/musal_behavior/mean_image.npy')

# %% [code]
len(dset_samples)

# %% [code]
dset_samples[0].shape

# %% [code]
idx = 0
plt.imshow(dset_samples[0][idx,0], cmap = 'gray')
plt.title('trial 0, T = ' + str(idx));
plt.colorbar()
print(dset_samples[0].shape)

# %% [code]
eps = 1e-2
T_max = max(a.shape[0] for a in dset_samples)
padded = [np.pad(a, ((0, T_max-a.shape[0]), (0,0), (0,0), (0,0)), mode='constant', constant_values=np.nan)
          for a in dset_samples]                                              # -> (n_trial, T_max, C, H, W)
stacked = np.stack(padded, axis=0)
min_img = np.nanmin(stacked, axis=(0,1))                                      # shape (C,H,W)
max_img = np.nanmax(stacked, axis=(0,1))                                      # shape (C,H,W)
clamp_range = np.stack([min_img - eps, max_img + eps], axis=0).astype(np.float32)  # shape (2,C,H,W)

# %% [markdown]
# 2. Read Model

# %% [code]
folder = 'out/bird_song_local_80/dimk_lag0_ori/t/none/00001-bird_dimk_lag0_none-prr-uncond-regularFM-gpus1-batch256-fp32'
chk_pts = '/network-snapshot-032614.pkl'
useCov = False
dim_to_keep = 3

pkl_path = folder + chk_pts

# %% [code]
if not hasattr(persistence, "_orig_reconstruct"):
    persistence._orig_reconstruct = persistence._reconstruct_persistent_obj
def _tolerant_reconstruct(meta):
    try:
        return persistence._orig_reconstruct(meta)
    except Exception:
        class _Stub: pass
        return _Stub()

class PatchedUnpickler(pickle.Unpickler):
    def find_class(self, module, name):
        if module == "torch_utils.persistence" and name == "_reconstruct_persistent_obj":
            return _tolerant_reconstruct
        return super().find_class(module, name)

with open(pkl_path, "rb") as f:
    model_all = PatchedUnpickler(f).load()

# use the net; ignore any loss_fn in the pickle
net = model_all["ema"] if "ema" in model_all else model_all["net"]
flow_net = net.unet_model
dyn_net   = net.vnet_model
encoder   = net.encoder

# %% [code]
lag = infer_lag_k_from_dyn_net(dyn_net)

# %% [code]
lag

# %% [code]
len(dset_samples)

# %% [markdown]
# 3. Check Encoder

# %% [code]
is_image = (np.asarray(dset_samples[0]).ndim == 4)
if is_image:
    _, C, H, W = np.asarray(dset_samples[0]).shape
T_min = min(arr.shape[0] for arr in dset_samples)
T_max = max(arr.shape[0] for arr in dset_samples)
n_step = T_max
dset_samples_look = dset_samples
# del dset_samples

# %% [code]
print(T_max)

# %% [code]
print("k_max =", encoder.k_max, "use_pop_stats =", encoder.use_pop_stats)

# %% [code]
trial_check = 10
T_check = 0
X0_back = recon_check_x0(encoder, trial_check, T_check, dset_samples = dset_samples_look)

# %% [code]
fig, axs = plt.subplots(1, 2, figsize=(6, 5))

im1 = axs[0].imshow(X0_back)
fig.colorbar(im1, ax=axs[0], fraction=0.046, pad=0.04)  # smaller, aligned
axs[0].set_title('encoder recon')

im2 = axs[1].imshow(dset_samples_look[trial_check][T_check,0,:,:])
fig.colorbar(im2, ax=axs[1], fraction=0.046, pad=0.04)  # same settings
axs[1].set_title('training data')

plt.tight_layout()
plt.show()

# %% [code]
data_flattent = [np.reshape(a, (a.shape[0], -1), order='C') for a in dset_samples_look]
dim_to_keep = 3
mu_keep = get_mu_from_encoder(encoder, data_flattent, dim_to_keep, trial_check)
plot_latent_time_series([mu_keep], alpha = 0.4, dims=np.arange(dim_to_keep), 
                        dt=0.01, labels=None)

# %% [code]
plot_traj_3d(mu_keep, title = 'trial = ' + str(trial_check))

# %% [code]
mu_tmp, d, L, loading, bias = latent_mu(dset_samples_look, encoder.to(device), dim_to_keep, device, fullRes = True)
L_np = L.detach().cpu().numpy()
d_np = d.detach().cpu().numpy()
loading_np = loading.detach().cpu().numpy()
mu_all = get_mu_from_encoder(encoder, data_flattent, 64*64, 0)

# %% [code]
k_max = 200
k_target = 50
loading_norm_all = torch.norm(loading[:,0:k_max], dim= 0).detach().cpu().numpy()
energy = loading_norm_all ** 2 
cum_energy = np.cumsum(energy)
K_eff = 1 + np.searchsorted(cum_energy, 0.95 * cum_energy[-1])

# %% [code]
fig = plt.figure(figsize=(3, 3))
plt.plot(np.sqrt(d_np[0:k_max]))
plt.axvline(x=k_target-1, color='red', label = 'k_target')
plt.axvline(x=K_eff-1, color='orange', label = 'k_eff = ' + str(K_eff))
plt.title('sqrt{d}');
plt.legend()

# %% [code]
k_max = 200
dim_check_most = 200
k_target = 50
rows = loading.shape[0]
cols = dim_check_most
dim_check = np.arange(dim_check_most)
# rec = mu_group_recon(bias.detach().cpu().numpy(), loading_np, mu_all, dim_check, H=28, W=28)
# ball_idx = 0

fig, axs = plt.subplots(2, 2, figsize=(6, 5))
axs[0, 0].plot(np.sqrt(d_np[0:dim_check_most]))
axs[0, 0].axvline(x=k_target-1, color='red', label = 'k_target')
axs[0, 0].axvline(x=K_eff-1, color='orange', label = 'k_eff = ' + str(K_eff))
axs[0, 0].set_title('sqrt{d}');
axs[0, 0].legend()

im1 = axs[0, 1].imshow(np.abs(loading_np[:, 0:dim_check_most]), cmap='gray_r', aspect=cols / rows)
axs[0, 1].set_title('|loading|')
fig.colorbar(im1, ax=axs[0, 1])

axs[1, 0].plot(mu_all[:, :dim_check_most])
axs[1, 0].set_title('first ' + str(dim_check_most) + " dim of mu")

# # Plot 4: cumulative contribution
# im2 = axs[1, 1].imshow(rec[ball_idx], cmap='gray')
# axs[1, 1].set_title('cumulative contribution from ' + str(dim_check[0]) + ' to ' + str(dim_check[-1]))
# fig.colorbar(im2, ax=axs[1, 1])

fig.suptitle('Encoder Diagnostic', fontsize=16)
plt.tight_layout()
plt.show()

# %% [code]


# %% [code]


# %% [markdown]
# 3. Flowed Latent

# %% [code]
gc.collect()
torch.cuda.empty_cache()
dim_to_keep = 3
latent_traj_list = proj_to_latent(dset_samples_look, flow_net.to(device), 0.0, device)

# %% [code]
lag

# %% [code]
latent_traj_list_enc = []
for ii in range(len(dset_samples_look)):
    latent_traj_list_enc.append(recon_check_x0_allT(encoder, ii, dset_samples_look))

# %% [code]
mu_traj_sim, d, L, loading, bias = latent_mu(latent_traj_list, encoder.to(device), dim_to_keep, device,
                                             fullRes = True, resid_tol=1e-15)

# %% [code]
data_flow = [arr.reshape(arr.shape[0], C, H, W) for arr in latent_traj_list ]
data_enc = [arr.reshape(arr.shape[0], C, H, W) for arr in latent_traj_list_enc ]

T_check = 0
fig, axs = plt.subplots(1, 3, figsize=(12, 10))

im1 = axs[0].imshow(data_flow[trial_check][T_check,0])
fig.colorbar(im1, ax=axs[0], fraction=0.046, pad=0.04)  # smaller, aligned
axs[0].set_title('flow(X)')

im2 = axs[1].imshow(data_enc[trial_check][T_check,0])
fig.colorbar(im2, ax=axs[1], fraction=0.046, pad=0.04)  # smaller, aligned
axs[1].set_title('encoder(X)')

im3 = axs[2].imshow(data_flow[trial_check][T_check,0] - data_enc[trial_check][T_check,0])
fig.colorbar(im3, ax=axs[2], fraction=0.046, pad=0.04)  # smaller, aligned
axs[2].set_title('flow(X) - encoder(X)')

plt.tight_layout()
plt.show()

# %% [code]
mu_traj_flow_np = [mu_traj_sim[j].cpu().numpy() for j in range(len(mu_traj_sim))]
plot_latent_time_series(mu_traj_flow_np, alpha = 0.4, dims=np.arange(dim_to_keep), 
                        dt=0.01, labels=None)
plt.suptitle('flowed latent trajc for each trial')

# %% [code]
plot_traj_3d(mu_traj_flow_np[trial_check][:,0:3], title = 'trial = ' + str(trial_check))

# %% [code]
plot_mu_trajectories_gradient_steps_3d(
    mu_traj_flow_np,
    trial_indices=np.arange(len(mu_traj_flow_np)), 
    dims=(0,1,2),
    title="all flowed Latent trajectories",
    z_scale='auto',  # Automatically scales z to be visually comparable to x and y
    arrow_head_frac=0.08,
    arrow_length_ratio=0.9,
    elev_azim = None, # (20,45)
)

# %% [markdown]
# 4. Simulated Traj

# %% [code]
n_step

# %% [code]
tau = 1.0
traj_sim_tau1_flattent, lag_cov_list = generate_traj_image_onthefly(dyn_net.to(device),
                                                                    dset_samples_look, n_step,
                                                                    tau, device, lag,
                                                                    include_x0_tau=False,
                                                                    oracle=False,
                                                                    flow_net=None,
                                                                    clamp_range=clamp_range
                                                                    )

# %% [code]
out_imgs_tau1 = [arr.reshape(arr.shape[0], C, H, W) for arr in traj_sim_tau1_flattent]

# %% [code]
n_step_trial = dset_samples_look[trial_check].shape[0]
plot_tau(n_step_trial, out_imgs_tau1, dset_samples_look, trial_idx = trial_check, tau = 1.0)

# %% [code]
out_imgs_tau0_flow_flatten = proj_to_latent(out_imgs_tau1, flow_net.to(device), 0.0, device)
out_imgs_tau0_flow = [arr.reshape(arr.shape[0], C, H, W) for arr in out_imgs_tau0_flow_flatten ]

# %% [code]
tau = 0.0
traj_sim_tau0_flattent, _ = generate_traj_image_onthefly(dyn_net.to(device),
                                                         dset_samples_look, n_step,
                                                         tau, device, lag,
                                                         include_x0_tau=False,
                                                         oracle=False,
                                                         flow_net=flow_net.to(device),
                                                         clamp_range=clamp_range
                                                         )
out_imgs_tau0 = [arr.reshape(arr.shape[0], C, H, W) for arr in traj_sim_tau0_flattent ]

# %% [code]
n_step_trial = dset_samples_look[trial_check].shape[0]
plot_tau(n_step_trial, out_imgs_tau0_flow, dset_samples_look,
         trial_idx = trial_check, tau = 0.0, title = 'porjected, tau = 0.0; flow from tau = 1.0 directly')

# %% [code]
plot_tau(n_step_trial, out_imgs_tau0, dset_samples_look,
         trial_idx = trial_check, tau = 0.0, title = 'simulated, tau = 0.0')

# %% [code]
dim_to_keep = 3
mu_traj_sim, d, L, loading, bias = latent_mu(traj_sim_tau0_flattent, encoder.to(device), dim_to_keep, device, fullRes = True)

# %% [code]
len(traj_sim_tau0_flattent)

# %% [code]
mu_traj_sim_np = [mu_traj_sim[j].cpu().numpy() for j in range(len(mu_traj_sim))]
mu_traj_sim_np_plot = [mu_traj_sim_np[ii][:,:] for ii in range(len(mu_traj_sim_np))]
plot_latent_time_series(mu_traj_sim_np_plot, alpha = 0.4, dims=np.arange(dim_to_keep), 
                        dt=0.01, labels=None)
plt.suptitle('simulated latent trajc for each trial')

# %% [code]
plot_traj_3d(mu_traj_sim_np_plot[trial_check][:,0:3], title = 'trial = ' + str(trial_check))

# %% [code]
plot_mu_trajectories_gradient_steps_3d(
    mu_traj_sim_np_plot,
    trial_indices=np.arange(len(mu_traj_sim_np_plot)), 
    dims=(0,1,2),
    title="all simulated Latent trajectories",
    z_scale='auto',  # Automatically scales z to be visually comparable to x and y
    arrow_head_frac=0.08,
    arrow_length_ratio=0.9,
    elev_azim = None, # (20,45)
)

# %% [markdown]
# 5. Flow Field

# %% [markdown]
Just lot 10% of traj

# %% [code]
# dset_samples_look_v2 = []
# for ii in range(len(dset_samples_look)//10):
#     dset_samples_look_v2.append(dset_samples_look[ii])

N = len(dset_samples_look)
k = N // 10

rng = np.random.default_rng(0)
idx = rng.choice(N, size=k, replace=False)
dset_samples_look_v2 = [dset_samples_look[i] for i in idx]

# %% [code]
latent_traj_list_enc = []
for ii in range(len(dset_samples_look_v2)):
    latent_traj_list_enc.append(recon_check_x0_allT(encoder, ii, dset_samples_look))
mu_traj_sim_enc, d, L, loading, bias = latent_mu(latent_traj_list_enc, encoder.to(device), dim_to_keep, device,
                                             fullRes = True, resid_tol=1e-15)

# %% [code]
len(latent_traj_list_enc)

# %% [code]
dim_to_keep = 3  # μ-dims you kept elsewhere
mu_pts_encoder, v_mu_encoder, v_mu_full_encoder = compute_mu_velocity_field(
    latent_traj_list=latent_traj_list_enc,
    dyn_net=dyn_net,
    encoder=encoder,
    device=device,
    dim_preserve=dim_to_keep,
    lag=lag,
    t_plot=0.0,
    return_full=True
)

# %% [code]
plot_mu_velocity_panel(mu_pts_encoder, v_mu_encoder,
                       trajectories=mu_traj_sim_enc,
                       speed_color=v_mu_full_encoder,
                       title_2d = "encoder: mu1–mu2 field (tau =0.0)",
                       title_3d = "encoder: mu1–mu2-mu3 field (tau =0.0)",
                       arrow_frac_2d=0.05,
                       arrow_size_2d=1.2,
                       linewidth_2d=1.5,
                       
                       arrow_frac_3d=0.1,
                       arrow_size_3d=1.5,
                       linewidth_3d=1.5,
                       base_arrow_length_ratio_3d=0.3,
                       )

# %% [code]
latent_traj_list = proj_to_latent(dset_samples_look_v2, flow_net.to(device), 0.0, device)
mu_traj_flow, d, L, loading, bias = latent_mu(latent_traj_list, encoder.to(device), dim_to_keep, device,
                                             fullRes = True, resid_tol=1e-15)

dim_to_keep = 3  # μ-dims you kept elsewhere
mu_pts_flow, v_mu_flow, v_mu_full_flow = compute_mu_velocity_field(
    latent_traj_list=latent_traj_list,
    dyn_net=dyn_net,
    encoder=encoder,
    device=device,
    dim_preserve=dim_to_keep,
    lag=lag,
    t_plot=0.0,
    return_full=True
)

plot_mu_velocity_panel(mu_pts_flow, v_mu_flow,
                       trajectories=mu_traj_flow, # mu_traj_sim
                       speed_color=v_mu_full_flow,
                       title_2d = "flowed: mu1–mu2 field (tau =0.0)",
                       title_3d = "flowed: mu1–mu2-mu3 field (tau =0.0)",
                       arrow_frac_2d=0.05,
                       arrow_size_2d=1.2,
                       linewidth_2d=1.5,
                       
                       arrow_frac_3d=0.1,
                       arrow_size_3d=1.5,
                       linewidth_3d=1.5,
                       base_arrow_length_ratio_3d=0.3,
                       )

# %% [code]
def calculate_mean_trajectory(traj_list):
    if not traj_list:
        return None
    min_len = min(arr.shape[0] for arr in traj_list)
    print(f"Minimum length found across trials: {min_len}")
    truncated_trajs = [arr[:min_len, :] for arr in traj_list]
    stacked_trajs = np.array(truncated_trajs)
    mean_traj = np.mean(stacked_trajs, axis=0)

    return mean_traj
mean_traj = calculate_mean_trajectory(mu_traj_flow_np)

# %% [code]
plot_traj_3d(mean_traj, title = 'mean traj')

# %% [code]
rng = np.random.default_rng(0)

N = len(traj_sim_tau0_flattent)
k = N // 10  # 10%

idx = rng.choice(N, size=k, replace=False)
traj_sim_tau0_flattent_2 = [traj_sim_tau0_flattent[i] for i in idx]


# %% [code]
len(traj_sim_tau0_flattent_2)

# %% [code]
mu_pts_sim, v_mu_sim, v_mu_full_sim = compute_mu_velocity_field(
    latent_traj_list=traj_sim_tau0_flattent_2,
    dyn_net=dyn_net,
    encoder=encoder,
    device=device,
    dim_preserve=dim_to_keep,
    lag=lag,
    t_plot=0.0,
    return_full=True
)

# %% [code]
plot_mu_velocity_panel(mu_pts_sim, v_mu_sim,
                       trajectories=mu_traj_sim, # mu_traj_sim
                       speed_color=v_mu_full_sim,
                       title_2d = "simulated: mu1–mu2 field (tau =0.0)",
                       title_3d = "simulated: mu1–mu2-mu3 field (tau =0.0)",
                       arrow_frac_2d=0.05,
                       arrow_size_2d=1.2,
                       linewidth_2d=1.5,
                       
                       arrow_frac_3d=0.1,
                       arrow_size_3d=1.5,
                       linewidth_3d=1.5,
                       base_arrow_length_ratio_3d=0.3,
                       )

# %% [markdown]
# Plot

# %% [markdown]
## Functions

# %% [code]
def plot_mu_velocity_2d(
    mu,
    v_mu,
    trajectories=None,
    dims_2d=(0, 1),
    title=None,
    arrow_frac=0.05,
    arrow_size=1.2,
    linewidth=1.5,
    traj_alpha=0.4,
    traj_width=0.8,
    traj_color="gray",
    max_trajs=300,
    quiver_alpha=0.6,
    max_points=3000,
    figsize=(7, 6),
    speed_color=None,                      # NEW
    speed_norm=None,                       # NEW
    colorbar_label=r"$\|\dot{\mu}\|$",     # NEW
):
    fig, ax = plt.subplots(figsize=figsize, constrained_layout=True)
    ax.grid(False)

    traj_list = []
    if trajectories is not None:
        traj_source = trajectories
        for i, traj in enumerate(traj_source):
            if i >= max_trajs:
                break
            t_np = traj.detach().cpu().numpy() if hasattr(traj, "detach") else np.array(traj)
            traj_list.append(t_np)

    d1, d2 = dims_2d
    for t_np in traj_list:
        ax.plot(
            t_np[:, d1], t_np[:, d2],
            color=traj_color, alpha=traj_alpha, linewidth=traj_width, zorder=1
        )

    coords = mu[:, [d1, d2]]
    vecs   = v_mu[:, [d1, d2]]
    N = coords.shape[0]

    # --- preprocess speed_color (for coloring only) ---
    speed_color_np = None
    if speed_color is not None:
        speed_color_np = speed_color.detach().cpu().numpy() if hasattr(speed_color, "detach") else np.asarray(speed_color)
        speed_color_np = np.asarray(speed_color_np)
        if speed_color_np.ndim == 2:
            speed_color_np = np.linalg.norm(speed_color_np, axis=1)  # (N,)
        speed_color_np = speed_color_np.reshape(-1)
        assert speed_color_np.shape[0] == mu.shape[0]

    if max_points is not None and N > max_points:
        idx = np.random.choice(N, size=max_points, replace=False)
        coords_2d = coords[idx]
        vecs_2d   = vecs[idx]
        speed_c   = speed_color_np[idx] if speed_color_np is not None else None
    else:
        coords_2d = coords
        vecs_2d   = vecs
        speed_c   = speed_color_np if speed_color_np is not None else None

    # --- directions from projected velocity ---
    speed_proj = np.linalg.norm(vecs_2d, axis=1)
    eps = 1e-8
    dirs = vecs_2d / (speed_proj[:, None] + eps)

    x_min, x_max = coords_2d[:, 0].min(), coords_2d[:, 0].max()
    y_min, y_max = coords_2d[:, 1].min(), coords_2d[:, 1].max()
    span = max(x_max - x_min, y_max - y_min) + eps
    L = arrow_frac * span
    U = dirs[:, 0] * L
    V = dirs[:, 1] * L

    base_width = 0.004
    base_headwidth = 4.0
    base_headlength = 6.0
    base_headaxislength = 5.0
    width = base_width * arrow_size

    # --- coloring values ---
    if speed_c is None:
        speed_c = speed_proj

    norm = speed_norm if speed_norm is not None else Normalize(
        vmin=float(speed_c.min()),
        vmax=float(speed_c.max()),
    )

    q = ax.quiver(
        coords_2d[:, 0], coords_2d[:, 1], U, V, speed_c,
        cmap=cm.viridis, norm=norm, angles="xy", scale_units="xy", scale=1.0,
        width=width,
        headwidth=base_headwidth * arrow_size,
        headlength=base_headlength * arrow_size,
        headaxislength=base_headaxislength * arrow_size,
        linewidth=linewidth, alpha=quiver_alpha, zorder=2
    )
    cb = fig.colorbar(q, ax=ax, fraction=0.046, pad=0.04)
    cb.set_label(colorbar_label)

    ax.set_xlabel(fr"$\mu_{{{d1+1}}}$")
    ax.set_ylabel(fr"$\mu_{{{d2+1}}}$")
    if title:
        ax.set_title(title)
    ax.axis("equal")

    return fig, ax, q


def plot_mu_velocity_3d(
    mu,
    v_mu,
    trajectories=None,
    dims_3d=(0, 1, 2),
    title=None,
    arrow_frac=0.1,
    arrow_size=1.5,
    linewidth=1.5,
    base_arrow_length_ratio=0.3,
    traj_alpha=0.4,
    traj_width=0.8,
    quiver_alpha=0.6,
    traj_color="gray",
    max_trajs=300,
    max_points=3000,
    figsize=(10, 7),
    elev=None,
    azim=None,
    roll=None,
    speed_color=None,                      # NEW
    speed_norm=None,                       # NEW
    colorbar_label=r"$\|\dot{\mu}\|$",     # NEW
):
    fig = plt.figure(figsize=figsize, constrained_layout=True)
    ax = fig.add_subplot(1, 1, 1, projection="3d")

    # set camera/view rotation
    if elev is not None or azim is not None or roll is not None:
        ax.view_init(elev=elev, azim=azim, roll=roll)

    ax.set_box_aspect((1, 1, 1))
    ax.grid(False)
    for axis in (ax.xaxis, ax.yaxis, ax.zaxis):
        axis.pane.fill = False
        axis.pane.set_edgecolor((1, 1, 1, 0))
        axis._axinfo["grid"]["color"] = (1, 1, 1, 0)
        axis._axinfo["grid"]["linewidth"] = 0

    traj_list = []
    if trajectories is not None:
        traj_source = trajectories
        for i, traj in enumerate(traj_source):
            if i >= max_trajs:
                break
            t_np = traj.detach().cpu().numpy() if hasattr(traj, "detach") else np.array(traj)
            traj_list.append(t_np)

    d1, d2, d3 = dims_3d
    for t_np in traj_list:
        ax.plot(
            t_np[:, d1], t_np[:, d2], t_np[:, d3],
            color=traj_color, alpha=traj_alpha, linewidth=traj_width, zorder=1
        )

    coords = mu[:, [d1, d2, d3]]
    vecs   = v_mu[:, [d1, d2, d3]]
    N = coords.shape[0]

    # --- preprocess speed_color (for coloring only) ---
    speed_color_np = None
    if speed_color is not None:
        speed_color_np = speed_color.detach().cpu().numpy() if hasattr(speed_color, "detach") else np.asarray(speed_color)
        speed_color_np = np.asarray(speed_color_np)
        if speed_color_np.ndim == 2:
            speed_color_np = np.linalg.norm(speed_color_np, axis=1)  # (N,)
        speed_color_np = speed_color_np.reshape(-1)
        assert speed_color_np.shape[0] == mu.shape[0]

    if max_points is not None and N > max_points:
        idx = np.random.choice(N, size=max_points, replace=False)
        coords_3d = coords[idx]
        vecs_3d   = vecs[idx]
        speed_c   = speed_color_np[idx] if speed_color_np is not None else None
    else:
        coords_3d = coords
        vecs_3d   = vecs
        speed_c   = speed_color_np if speed_color_np is not None else None

    # --- directions from projected velocity ---
    speed_proj = np.linalg.norm(vecs_3d, axis=1)
    eps = 1e-8
    dirs = vecs_3d / (speed_proj[:, None] + eps)

    mins = coords_3d.min(axis=0)
    maxs = coords_3d.max(axis=0)
    span = float(np.max(maxs - mins) + eps)

    L = arrow_frac * span
    U, V, W = dirs[:, 0], dirs[:, 1], dirs[:, 2]

    # --- coloring values ---
    if speed_c is None:
        speed_c = speed_proj

    norm = speed_norm if speed_norm is not None else Normalize(
        vmin=float(speed_c.min()),
        vmax=float(speed_c.max()),
    )

    colors = cm.viridis(norm(speed_c))
    colors[:, 3] = quiver_alpha

    ax.quiver(
        coords_3d[:, 0], coords_3d[:, 1], coords_3d[:, 2],
        U, V, W,
        length=L, normalize=False, colors=colors,
        linewidth=linewidth * arrow_size,
        arrow_length_ratio=base_arrow_length_ratio * arrow_size,
        zorder=2
    )

    mappable = cm.ScalarMappable(norm=norm, cmap="viridis")
    mappable.set_array(speed_c)
    cb = fig.colorbar(mappable, ax=ax, fraction=0.035, pad=0.08)
    cb.set_label(colorbar_label)

    ax.set_xlabel(fr"$\mu_{{{d1+1}}}$")
    ax.set_ylabel(fr"$\mu_{{{d2+1}}}$")
    ax.set_zlabel(fr"$\mu_{{{d3+1}}}$")
    if title:
        ax.set_title(title)

    return fig, ax

# %% [code]
def plot_frame_compact(
    rows,                 # list like [dset_samples, out_imgs_tau0_flow, ...]
    trial_idx,
    idxs,                 # 1D int array, columns
    row_labels=None,      # list of strings, one per row
    mean_img=None,
    figsize=(6, 2.6),
    img_cmap="gray",
    vmin=None,
    vmax=None,
    border_from=None,
    border_lw=3.0,
    left=0.10, right=0.99, bottom=0.05, top=0.95,
    wspace=0.02, hspace=0.02,
):
    import numpy as np
    import matplotlib.pyplot as plt

    idxs = np.asarray(idxs).astype(int)
    n_rows = len(rows)
    n_cols = len(idxs)

    # --- global vmin/vmax (consistent across rows) ---
    if vmin is None or vmax is None:
        mn, mx = np.inf, -np.inf
        for data in rows:
            X = data[trial_idx]  # [T, 1, H, W] (torch or np)
            for i in idxs:
                img = X[i, 0]
                if hasattr(img, "detach"):
                    img = img.detach().cpu().numpy()
                else:
                    img = np.asarray(img)
                if mean_img is not None:
                    img = img + mean_img
                mn = min(mn, float(img.min()))
                mx = max(mx, float(img.max()))
        if vmin is None: vmin = mn
        if vmax is None: vmax = mx

    if border_from is not None:
        cmap = border_from.get_cmap()
        norm = border_from.norm
        col_colors = cmap(norm(idxs.astype(float)))
    else:
        col_colors = ["k"] * n_cols

    fig, axes = plt.subplots(n_rows, n_cols, figsize=figsize, squeeze=False)
    fig.subplots_adjust(left=left, right=right, bottom=bottom, top=top, wspace=wspace, hspace=hspace)

    for r, data in enumerate(rows):
        X = data[trial_idx]
        for c, i in enumerate(idxs):
            ax = axes[r, c]

            img = X[i, 0]
            if hasattr(img, "detach"):
                img = img.detach().cpu().numpy()
            else:
                img = np.asarray(img)
            if mean_img is not None:
                img = img + mean_img

            ax.imshow(img, cmap=img_cmap, vmin=vmin, vmax=vmax, origin='lower')

            # keep spines for colored border, so don't ax.axis("off")
            ax.set_xticks([]); ax.set_yticks([])

            # titles only on first row
            if r == 0:
                ax.set_title(f"t={i}")
            else:
                ax.set_title("")

            # colored border per column
            for sp in ax.spines.values():
                sp.set_visible(True)
                sp.set_linewidth(border_lw)
                sp.set_color(col_colors[c])

    # row labels on the left, centered per row (compact)
    if row_labels is not None:
        for r, lab in enumerate(row_labels):
            pos = axes[r, 0].get_position()
            y = 0.5 * (pos.y0 + pos.y1)
            x = pos.x0 - 0.02
            fig.text(x, y, lab, ha="right", va="center")

    return fig, axes, vmin

# %% [markdown]
## Plottings

# %% [code]
elev = 20 
azim = 60

# %% [code]
plotFolder = "paper_fig/bird"

# %% [code]
# 1. sqrt{d}
fig = plt.figure(figsize=(4, 4))
plt.plot(np.sqrt(d_np[0:k_max]))
plt.axvline(x=k_target-1, color='red', label = 'k_target')
plt.axvline(x=K_eff-1, color='orange', label = 'k_eff = ' + str(K_eff))
plt.title('sqrt{d}');
plt.legend()

plt.savefig(plotFolder + "/1_sqrt_d.pdf")
plt.savefig(plotFolder + "/1_sqrt_d.svg")
plt.savefig(plotFolder + "/1_sqrt_d.png")

# %% [code]
# 2. compressed flow field in mu (3D)
fig2, ax2 = plot_mu_velocity_3d(
    mu_pts_flow, v_mu_flow, trajectories=mu_traj_flow, dims_3d=(0,1,2),
    speed_color=v_mu_full_flow,
    title="flowed: mu1–mu2-mu3 field (tau =0.0)", figsize=(5, 5),
    arrow_frac=0.08, arrow_size=1.0, elev=elev, azim=azim
)

plt.savefig(plotFolder + "/2_flow_v_3d.pdf")
plt.savefig(plotFolder + "/2_flow_v_3d.svg")
plt.savefig(plotFolder + "/2_flow_v_3d.png")

# %% [code]
# 3. compressed flow field in mu (3D)
fig3, ax3 = plot_mu_velocity_3d(
    mu_pts_sim, v_mu_sim, trajectories=mu_traj_sim, dims_3d=(0,1,2),
    speed_color=v_mu_full_sim,
    traj_alpha=0.1,
    title="simulated: mu1–mu2-mu3 field (tau =0.0)", figsize=(5, 5),
    arrow_frac=0.06, arrow_size=0.8, elev=elev, azim=azim,
    base_arrow_length_ratio = 0.5,
    quiver_alpha=0.8
)

# xlim_ref, ylim_ref, zlim_ref = ax2.get_xlim3d(), ax2.get_ylim3d(), ax2.get_zlim3d()
# ax3.set_xlim3d(xlim_ref); ax3.set_ylim3d(ylim_ref); ax3.set_zlim3d(zlim_ref)

plt.savefig(plotFolder + "/3_sim_v_3d.pdf")
plt.savefig(plotFolder + "/3_sim_v_3d.svg")
plt.savefig(plotFolder + "/3_sim_v_3d.png")

# %% [code]
mu_traj_flow_all = np.concatenate(mu_traj_flow_np, axis=0)
mean_traj_flow = calculate_mean_trajectory(mu_traj_flow_np)

# %% [code]
idx_select = np.arange(0, dset_samples[0].shape[0], 5, dtype=int)

# %% [code]
print(len(mu_traj_sim_np_plot))
print(mu_traj_sim_np_plot[0].shape)

# %% [code]
from mpl_toolkits.mplot3d.art3d import Line3DCollection
from matplotlib.colors import Normalize

roll = None

trial_idx = 0
trial_sel = [trial_idx]   # edit
n_repr = 5            # edit

T_tr = mu_traj_sim_np_plot[0].shape[0]      # = 26
t_max = T_tr - 1                             # = 25
norm = Normalize(vmin=0, vmax=t_max)

mu_traj_sim_all = np.concatenate(mu_traj_sim_np_plot, axis=0)
mean_traj_sim = calculate_mean_trajectory(mu_traj_sim_np_plot)

fig = plt.figure(figsize=(5, 5), constrained_layout=True)
ax3d = fig.add_subplot(111, projection='3d')

if elev is not None or azim is not None or roll is not None:
    ax3d.view_init(elev=elev, azim=azim, roll=roll)

ax3d.set_box_aspect((1, 1, 1))
ax3d.grid(False)
for axis in (ax3d.xaxis, ax3d.yaxis, ax3d.zaxis):
    axis.pane.fill = False
    axis.pane.set_edgecolor((1, 1, 1, 0))
    axis._axinfo["grid"]["color"] = (1, 1, 1, 0)
    axis._axinfo["grid"]["linewidth"] = 0

# ax3d.scatter(mu_traj_sim_all[:, 0], mu_traj_sim_all[:, 1], mu_traj_sim_all[:, 2],
#              c="k", s=2, alpha=0.04)
for P_tr in mu_traj_sim_np_plot:   # each is (26,3)
    ax3d.plot(P_tr[:, 0], P_tr[:, 1], P_tr[:, 2],
              color="k", alpha=0.04, linewidth=0.8)


# representative points (colored by absolute time step 0..25)
pts_list, c_list = [], []
for tr in trial_sel:
    P_tr = mu_traj_sim_np_plot[tr]                  # [26, 3]
    t_idx = np.linspace(0, t_max, n_repr).round().astype(int)
    t_idx = np.unique(t_idx)

    pts_list.append(P_tr[t_idx])
    c_list.append(t_idx.astype(float))

    ax3d.plot(P_tr[:, 0], P_tr[:, 1], P_tr[:, 2], color="k", alpha=0.15, linewidth=1.0)

pts = np.concatenate(pts_list, axis=0)
cvals = np.concatenate(c_list, axis=0)

sc = ax3d.scatter(pts[:, 0], pts[:, 1], pts[:, 2],
                  c=cvals, cmap="plasma", norm=norm, s=60, alpha=1.0)

# colored trajectory = selected trial (same one used for representative points)
tr0 = trial_sel[0]
P = mu_traj_sim_np_plot[tr0]  # (26,3)

segs = np.stack([P[:-1], P[1:]], axis=1)
t_line = np.arange(P.shape[0] - 1).astype(float)

lc = Line3DCollection(segs, cmap="plasma", norm=norm)
lc.set_array(t_line)
lc.set_linewidth(3.0)
lc.set_alpha(1.0)
ax3d.add_collection3d(lc)


# P = mean_traj_sim
# segs = np.stack([P[:-1], P[1:]], axis=1)
# t_line = np.arange(P.shape[0] - 1).astype(float)

# lc = Line3DCollection(segs, cmap="plasma", norm=norm)
# lc.set_array(t_line)
# lc.set_linewidth(3.0)
# lc.set_alpha(1.0)
# ax3d.add_collection3d(lc)

cbar = fig.colorbar(sc, ax=ax3d, fraction=0.03, pad=0.08)
cbar.set_label("time step (t)")
fig.subplots_adjust(right=0.86)
ax3d.set_title("simulated mu")

xlim_train, ylim_train, zlim_train = ax3d.get_xlim3d(), ax3d.get_ylim3d(), ax3d.get_zlim3d()

plt.savefig(plotFolder + "/5_sel_sim_v_3d_train.pdf")
plt.savefig(plotFolder + "/5_sel_sim_v_3d_train.svg")
plt.savefig(plotFolder + "/5_sel_sim_v_3d_train.png")

# %% [code]
from mpl_toolkits.mplot3d.art3d import Line3DCollection
from matplotlib.colors import Normalize

roll = None

fig = plt.figure(figsize=(5, 5), constrained_layout=True)
ax3d = fig.add_subplot(111, projection='3d')

if elev is not None or azim is not None or roll is not None:
    ax3d.view_init(elev=elev, azim=azim, roll=roll)

ax3d.set_box_aspect((1, 1, 1))


for P_tr in mu_traj_flow_np:   # each is (26,3)
    ax3d.plot(P_tr[:, 0], P_tr[:, 1], P_tr[:, 2],
              color="k", alpha=0.04, linewidth=0.8)


# ax3d.scatter(mu_traj_flow_all[:, 0], mu_traj_flow_all[:, 1], mu_traj_flow_all[:, 2],
#              c="k", s=2, alpha=0.05)
ax3d.grid(False)
for axis in (ax3d.xaxis, ax3d.yaxis, ax3d.zaxis):
    axis.pane.fill = False
    axis.pane.set_edgecolor((1, 1, 1, 0))
    axis._axinfo["grid"]["color"] = (1, 1, 1, 0)
    axis._axinfo["grid"]["linewidth"] = 0

idx_sel = np.asarray(idx_select)
order = np.argsort(idx_sel)
idx_sel = idx_sel[order]

vals = idx_sel.astype(float)

# P = mean_traj_flow
tr0 = trial_sel[0]          # reuse from 5_sel_sim_v_3d_train
P = mu_traj_flow_np[tr0]    # selected trial trajectory (26,3)

segs = np.stack([P[:-1], P[1:]], axis=1)
t = np.linspace(vals.min(), vals.max(), P.shape[0] - 1)

lc = Line3DCollection(segs, cmap="plasma",
                      norm=Normalize(vmin=vals.min(), vmax=vals.max()))
lc.set_array(t)
lc.set_linewidth(3.0)
lc.set_alpha(1.0)
ax3d.add_collection3d(lc)

cbar = fig.colorbar(sc, ax=ax3d, fraction=0.03, pad=0.08)
cbar.set_label("selection order")
fig.subplots_adjust(right=0.86)
ax3d.set_title('flowed mu')

ax3d.set_xlim3d(xlim_train)
ax3d.set_ylim3d(ylim_train)
ax3d.set_zlim3d(zlim_train)

plt.savefig(plotFolder + "/4_flow_v_3d.pdf")
plt.savefig(plotFolder + "/4_flow_v_3d.svg")
plt.savefig(plotFolder + "/4_flow_v_3d.png")

# %% [code]
height1 = 1.3
_,_,vmin_ori = plot_frame_compact(
    rows=[dset_samples, out_imgs_tau1, out_imgs_tau0],
    row_labels=["data", "tau = 1.0", "tau = 0.0"],
    trial_idx=trial_idx,
    idxs=idx_select,
    border_from=sc,         
    figsize=(6.8, height1*3),
    mean_img=None,
);

plt.savefig(plotFolder + "/6_sel_sim_img_train.pdf")
plt.savefig(plotFolder + "/6_sel_sim_img_train.svg")
plt.savefig(plotFolder + "/6_sel_sim_img_train.png")

# %% [markdown]
## test part

# %% [code]
folder_data = 'data/bird_song_small_80_test'
dataset_samples_test = np.load(folder_data + "/dataset_samples.npz", allow_pickle=True)

dset_samples_test = to_list_of_arrays(dataset_samples_test['samples'])

# %% [code]
len(dset_samples_test)

# %% [code]
tau = 1.0
traj_sim_tau1_flattent_test, lag_cov_list_test = generate_traj_image_onthefly(dyn_net.to(device),
                                                                              dset_samples_test, n_step,
                                                                              tau, device, lag,
                                                                              include_x0_tau=False,
                                                                              oracle=False,
                                                                              flow_net=None,
                                                                              clamp_range=clamp_range
                                                                              )
out_imgs_tau1_test = [arr.reshape(arr.shape[0], C, H, W) for arr in traj_sim_tau1_flattent_test]

tau = 0.0
traj_sim_tau0_flattent_test, _ = generate_traj_image_onthefly(dyn_net.to(device),
                                                              dset_samples_test, n_step,
                                                              tau, device, lag,
                                                              include_x0_tau=False,
                                                              oracle=False,
                                                              flow_net=flow_net.to(device),
                                                              clamp_range=clamp_range
                                                              )
out_imgs_tau0_test = [arr.reshape(arr.shape[0], C, H, W) for arr in traj_sim_tau0_flattent_test ]

# %% [code]
dim_to_keep = 3
mu_traj_sim_test, d, L, loading, bias_test = latent_mu(traj_sim_tau0_flattent_test, encoder.to(device), dim_to_keep, device, fullRes = True)

# %% [code]
mu_traj_sim_np_test = [mu_traj_sim_test[j].cpu().numpy() for j in range(len(mu_traj_sim_test))]
mu_traj_sim_np_plot_test = [mu_traj_sim_np_test[ii][:,:] for ii in range(len(mu_traj_sim_np_test))]

# %% [code]
from mpl_toolkits.mplot3d.art3d import Line3DCollection
from matplotlib.colors import Normalize

roll = None

trial_idx = 0
trial_sel = [trial_idx]   # edit
n_repr = 5            # edit

T_tr = mu_traj_sim_np_plot_test[0].shape[0]      # = 26
t_max = T_tr - 1                             # = 25
norm = Normalize(vmin=0, vmax=t_max)

mu_traj_sim_all = np.concatenate(mu_traj_sim_np_plot_test, axis=0)
mean_traj_sim = calculate_mean_trajectory(mu_traj_sim_np_plot_test)

fig = plt.figure(figsize=(5, 5), constrained_layout=True)
ax3d = fig.add_subplot(111, projection='3d')

# NEW: set view rotation
if elev is not None or azim is not None or roll is not None:
    ax3d.view_init(elev=elev, azim=azim, roll=roll)

ax3d.set_box_aspect((1, 1, 1))

# ax3d.scatter(mu_traj_sim_all[:, 0], mu_traj_sim_all[:, 1], mu_traj_sim_all[:, 2],
#              c="k", s=2, alpha=0.05)


# background: ALL trials as faint trajectories
for P_tr in mu_traj_sim_np_plot_test:   # each (26,3)
    ax3d.plot(P_tr[:, 0], P_tr[:, 1], P_tr[:, 2],
              color="k", alpha=0.04, linewidth=0.8)


ax3d.grid(False)
for axis in (ax3d.xaxis, ax3d.yaxis, ax3d.zaxis):
    axis.pane.fill = False
    axis.pane.set_edgecolor((1, 1, 1, 0))
    axis._axinfo["grid"]["color"] = (1, 1, 1, 0)
    axis._axinfo["grid"]["linewidth"] = 0

# representative points (colored by absolute time step 0..25)
pts_list, c_list = [], []
for tr in trial_sel:
    P_tr = mu_traj_sim_np_plot_test[tr]                  # [26, 3]
    t_idx = np.linspace(0, t_max, n_repr).round().astype(int)
    t_idx = np.unique(t_idx)

    pts_list.append(P_tr[t_idx])
    c_list.append(t_idx.astype(float))

    ax3d.plot(P_tr[:, 0], P_tr[:, 1], P_tr[:, 2], color="k", alpha=0.15, linewidth=1.0)

pts = np.concatenate(pts_list, axis=0)
cvals = np.concatenate(c_list, axis=0)

sc = ax3d.scatter(pts[:, 0], pts[:, 1], pts[:, 2],
                  c=cvals, cmap="plasma", norm=norm, s=60, alpha=1.0)


tr0 = trial_sel[0]
P = mu_traj_sim_np_plot_test[tr0]  # (26,3)

segs = np.stack([P[:-1], P[1:]], axis=1)
t_line = np.arange(P.shape[0] - 1).astype(float)

lc = Line3DCollection(segs, cmap="plasma", norm=norm)
lc.set_array(t_line)
lc.set_linewidth(3.0)
lc.set_alpha(1.0)
ax3d.add_collection3d(lc)


# P = mean_traj_sim
# segs = np.stack([P[:-1], P[1:]], axis=1)
# t_line = np.arange(P.shape[0] - 1).astype(float)

# lc = Line3DCollection(segs, cmap="plasma", norm=norm)
# lc.set_array(t_line)
# lc.set_linewidth(3.0)
# lc.set_alpha(1.0)
# ax3d.add_collection3d(lc)

cbar = fig.colorbar(sc, ax=ax3d, fraction=0.03, pad=0.08)
cbar.set_label("time step (t)")
fig.subplots_adjust(right=0.86)
ax3d.set_title("simulated mu")

ax3d.set_xlim3d(xlim_train)
ax3d.set_ylim3d(ylim_train)
ax3d.set_zlim3d(zlim_train)

plt.savefig(plotFolder + "/7_sel_sim_v_3d_test.pdf")
plt.savefig(plotFolder + "/7_sel_sim_v_3d_test.svg")
plt.savefig(plotFolder + "/7_sel_sim_v_3d_test.png")

# %% [code]
height1 = 1.3
plot_frame_compact(
    rows=[dset_samples_test, out_imgs_tau1_test, out_imgs_tau0_test],
    row_labels=["data", "tau = 1.0", "tau = 0.0"],
    trial_idx=trial_idx,
    idxs=idx_select,
    border_from=sc,         
    figsize=(6.8, height1*3),
    mean_img=None,
);

plt.savefig(plotFolder + "/8_sel_sim_img_test.pdf")
plt.savefig(plotFolder + "/8_sel_sim_img_test.svg")
plt.savefig(plotFolder + "/8_sel_sim_img_test.png")

# %% [code]


# %% [markdown]
# Modified Version

# %% [code]
height1 = 1.3
plot_frame_compact(
    rows=[dset_samples, out_imgs_tau1, out_imgs_tau0],
    row_labels=["data", "tau = 1.0", "tau = 0.0"],
    trial_idx=trial_idx,
    idxs=idx_select,
    border_from=sc,
    img_cmap='gray',
    figsize=(6.8, height1*3),
    mean_img=None,
    vmin = vmin_ori+0.3
);

plt.savefig(plotFolder + "/9_sel_sim_img_train_v2.pdf")
plt.savefig(plotFolder + "/9_sel_sim_img_train_v2.svg")
plt.savefig(plotFolder + "/9_sel_sim_img_train_v2.png")

# %% [code]
height1 = 1.3
plot_frame_compact(
    rows=[dset_samples_test, out_imgs_tau1_test, out_imgs_tau0_test],
    row_labels=["data", "tau = 1.0", "tau = 0.0"],
    trial_idx=trial_idx,
    idxs=idx_select,
    border_from=sc,         
    figsize=(6.8, height1*3),
    mean_img=None,
    vmin = vmin_ori+0.3
);

plt.savefig(plotFolder + "/10_sel_sim_img_test_v2.pdf")
plt.savefig(plotFolder + "/10_sel_sim_img_test_v2.svg")
plt.savefig(plotFolder + "/10_sel_sim_img_test_v2.png")

# %% [code]


# %% [code]
```
