## Supplementary material for "Dynamic Compression Flows for Neuroscience Data": Python code: paper_plot_maze.ipynb

```
# %% [code]
import numpy as np
import pickle
import os
import re 
import json 
import click
import torch
import dnnlib.util_v4 as util
from torch_utils import distributed as dist
from training import toy_training_loop_vfm_noDataGen
from tqdm import tqdm
import matplotlib.pyplot as plt
from matplotlib import colors
from plot_neural import *
from plot_sim_results_v7 import *
from plot_sim_results_img_v7 import * 
import pandas as pd

import warnings
warnings.filterwarnings('ignore', 'Grad strides do not match bucket view strides') # False warning 
device = torch.device("cuda" if torch.cuda.is_available() else "cpu")
torch.set_default_device(device)

# %% [code]
# colorgrad = "PuOr"
colorgrad = 'viridis'

# %% [markdown]
# 1. Read Data

# %% [code]
folder_data = 'data/mc_maze3'
dataset_samples = np.load(folder_data + "/dataset_samples.npz", allow_pickle=True)
data_raw = dataset_samples['samples']
dset_samples_full = []
for ii in range(len(data_raw)):
    dset_samples_full.append(np.asarray(data_raw[ii], dtype=np.float64))

# %% [code]
# corresponding meta data
train_meta = np.load(folder_data + "/meta_train.npy", allow_pickle=True).tolist()
trial_idx_use = np.array(train_meta['trial_info']['trial_version']) == 0
# np.sum(trial_idx_use)

# %% [code]
dset_samples_full[0].shape

# %% [code]
len(dset_samples_full)

# %% [code]
import numpy as np

def _build_proj_history_and_target(dset_samples_full, k, d_proj=64, seed=0):
    """
    Build (H_proj, Y) where:
      H_proj[t] ~ random-projection of h_t^{(k)} = [x_t, x_{t-1}, ..., x_{t-k}]
      Y[t]      = x_{t+1}
    using only numpy, without ever materializing the full (k+1)*D history vector.
    """
    rng = np.random.default_rng(seed)

    B = len(dset_samples_full)
    X0 = np.asarray(dset_samples_full[0])
    assert X0.ndim == 2, "Expect each trial is (T, D)."
    T, D = X0.shape

    # random projection blocks: one (D,d_proj) per lag slice
    # scale by 1/sqrt(d_proj) so norms stay reasonable
    R = rng.standard_normal(size=(k + 1, D, d_proj)).astype(np.float32) / np.sqrt(d_proj)

    H_list, Y_list = [], []
    for j in range(B):
        X = np.asarray(dset_samples_full[j], dtype=np.float32)
        Tj, Dj = X.shape
        assert Dj == D, "All trials must share the same D."

        L = Tj - 1 - k
        if L <= 0:
            continue

        # targets: y_t = x_{t+1}, t = k..T-2  -> indices (k+1 .. T-1)
        Y = X[k + 1 : Tj]                         # (L, D)

        # projected history: sum_{s=0..k} x_{t-s} @ R[s]
        H = np.zeros((L, d_proj), dtype=np.float32)
        for s in range(k + 1):
            # x_{t-s}: indices (k-s .. T-2-s) length L
            H += X[k - s : (Tj - 1) - s] @ R[s]   # (L,d_proj)

        H_list.append(H)
        Y_list.append(Y)

    H_all = np.concatenate(H_list, axis=0)  # (N, d_proj)
    Y_all = np.concatenate(Y_list, axis=0)  # (N, D)
    return H_all, Y_all


def _knn_conditional_var_ratio(H, Y, m=20, n_ref=6000, n_query=800, seed=0):
    """
    Estimate  E[ Var(Y | H) ] / Var(Y)  via kNN in H-space.
    """
    rng = np.random.default_rng(seed)
    N = H.shape[0]
    n_ref = min(n_ref, N)
    n_query = min(n_query, n_ref)

    ref_idx = rng.choice(N, size=n_ref, replace=False)
    # choose queries from ref set so "self" exists in the reference pool
    q_pos = rng.choice(n_ref, size=n_query, replace=False)
    q_idx = ref_idx[q_pos]

    H_ref = H[ref_idx].astype(np.float32, copy=False)
    Y_ref = Y[ref_idx].astype(np.float32, copy=False)
    H_q   = H[q_idx].astype(np.float32, copy=False)

    # z-score H (stabilizes distances)
    mu = H_ref.mean(axis=0, keepdims=True)
    sd = H_ref.std(axis=0, keepdims=True) + 1e-6
    H_ref = (H_ref - mu) / sd
    H_q   = (H_q   - mu) / sd

    # precompute norms
    ref_norm = (H_ref * H_ref).sum(axis=1)  # (n_ref,)

    # brute-force kNN in chunks
    k_take = m + 1  # include self, then drop it
    nn_pos = np.empty((n_query, k_take), dtype=np.int64)

    bs = 64
    for a in range(0, n_query, bs):
        b = min(n_query, a + bs)
        Q = H_q[a:b]                              # (bs,d)
        q_norm = (Q * Q).sum(axis=1)              # (bs,)
        # dist^2 = ||q||^2 + ||r||^2 - 2 q·r
        dist2 = q_norm[:, None] + ref_norm[None, :] - 2.0 * (Q @ H_ref.T)
        # take k smallest positions
        part = np.argpartition(dist2, kth=k_take - 1, axis=1)[:, :k_take]
        # sort those k by actual dist
        row = np.arange(b - a)[:, None]
        part_sorted = part[row, np.argsort(dist2[row, part], axis=1)]
        nn_pos[a:b] = part_sorted

    # drop self if present, otherwise just drop the closest
    nn_pos_drop = np.empty((n_query, m), dtype=np.int64)
    for i in range(n_query):
        cand = nn_pos[i]  # (m+1,)
        # find self in candidates
        self_mask = (ref_idx[cand] == q_idx[i])
        if np.any(self_mask):
            cand = cand[~self_mask]
        else:
            cand = cand[1:]  # drop closest
        nn_pos_drop[i] = cand[:m]

    # neighbor targets
    Y_nb = Y_ref[nn_pos_drop]            # (n_query, m, D)
    local_var = np.var(Y_nb, axis=1)     # (n_query, D)
    local_var_mean = float(local_var.mean())

    global_var = np.var(Y_ref, axis=0)   # (D,)
    global_var_mean = float(global_var.mean()) + 1e-12

    return local_var_mean / global_var_mean


def decide_lag_knn(
    dset_samples_full,
    ks=(0, 1, 2, 3, 4, 6, 8, 12),
    d_proj=64,
    m=20,
    n_ref=6000,
    n_query=800,
    seed=0,
    plateau_rel=0.02,   # 2% improvement threshold
):
    """
    Prints a small table and returns a suggested lag k.
    """
    ratios = []
    print("k |  E[Var(x_{t+1}|h_t^k)] / Var(x_{t+1})   (lower is better)")
    print("--+--------------------------------------------------------")
    for k in ks:
        H, Y = _build_proj_history_and_target(dset_samples_full, k, d_proj=d_proj, seed=seed + 17*k)
        r = _knn_conditional_var_ratio(H, Y, m=m, n_ref=n_ref, n_query=n_query, seed=seed + 101*k)
        ratios.append(r)
        print(f"{k:2d} |  {r:.6f}")

    # pick smallest k where improvements plateau for 2 consecutive steps
    k_suggest = ks[-1]
    for i in range(2, len(ks)):
        r0, r1, r2 = ratios[i-2], ratios[i-1], ratios[i]
        imp1 = (r0 - r1) / max(r0, 1e-12)
        imp2 = (r1 - r2) / max(r1, 1e-12)
        if (imp1 < plateau_rel) and (imp2 < plateau_rel):
            k_suggest = ks[i-1]
            break

    print(f"\nSuggested lag k ≈ {k_suggest}  (first plateau with <{100*plateau_rel:.1f}% gains twice)")
    return k_suggest, dict(zip(ks, ratios))


# ---- example ----
k_star, ratio_table = decide_lag_knn(dset_samples_full)


# %% [markdown]
# 2. Behavior Data

# %% [code]
trial_type_use = np.array(train_meta['trial_info']['trial_type'])[trial_idx_use]
indices = np.where(trial_idx_use)[0]

target_pos_sel = [train_meta['trial_info']['target_pos'][idx] for idx in indices]
target_pos_sel = np.array(target_pos_sel).squeeze(1).astype(np.float32)

hand_pos_sel = [train_meta['behavior']['hand_pos'][idx] for idx in indices]
cursor_pos_sel = [train_meta['behavior']['cursor_pos'][idx] for idx in indices]
eye_pos_sel = [train_meta['behavior']['eye_pos'][idx] for idx in indices]

# %% [markdown]
# 3. Read Model and Data 

# %% [code]
# folder_data = 'out/maze/dimk_lag5/t/none/data'
# folder = 'out/maze/dimk_lag5/t/none/00002-maze_dimk_lag5_none-prp-uncond-regularFM-gpus1-batch256-fp32'
# chk_pts = '/network-snapshot-050176.pkl'

folder_data = 'out/maze/dimk_lag10/t/none/data'
folder = 'out/maze/dimk_lag10/t/none/00003-maze_dimk_lag10_none-prp-uncond-regularFM-gpus1-batch256-fp32'
# chk_pts = '/network-snapshot-262922.pkl'
chk_pts = '/network-snapshot-055695.pkl'

dataset_samples = np.load(folder_data + "/dataset_samples.npz", allow_pickle=True)
dset_samples = to_list_of_arrays(dataset_samples['samples'])

# %% [code]
dt = 0.001
eps_clamp = 1e-2
T_max = max(arr.shape[0] for arr in dset_samples)
padded_arrays = [np.pad(arr, ((0, T_max - arr.shape[0]), (0, 0)),
                        mode='constant', constant_values=np.nan) 
                 for arr in dset_samples]
stacked = np.stack(padded_arrays, axis=0)  # Shape [n_trial, T_max, dim]


min_vals = np.nanmin(stacked, axis=(0, 1))  # Shape [dim]
max_vals = np.nanmax(stacked, axis=(0, 1))  # Shape [dim]

clamp_range = np.stack([min_vals, max_vals], axis=1)  # Shape [dim, 2]
clamp_range[:,0] = clamp_range[:,0] - eps_clamp
clamp_range[:,1] = clamp_range[:,1] + eps_clamp

# %% [code]
useCov = False
dim_to_keep = 3

pkl_path = folder + chk_pts

# %% [code]
if not hasattr(persistence, "_orig_reconstruct"):
    persistence._orig_reconstruct = persistence._reconstruct_persistent_obj
def _tolerant_reconstruct(meta):
    try:
        return persistence._orig_reconstruct(meta)
    except Exception:
        class _Stub: pass
        return _Stub()

class PatchedUnpickler(pickle.Unpickler):
    def find_class(self, module, name):
        if module == "torch_utils.persistence" and name == "_reconstruct_persistent_obj":
            return _tolerant_reconstruct
        return super().find_class(module, name)

with open(pkl_path, "rb") as f:
    model_all = PatchedUnpickler(f).load()

# use the net; ignore any loss_fn in the pickle
net = model_all["ema"] if "ema" in model_all else model_all["net"]
flow_net = net.unet_model
dyn_net   = net.vnet_model
encoder   = net.encoder

# %% [code]
flow_net.training

# %% [code]
lag = dset_samples_full[0].shape[0] - dset_samples[0].shape[0]
print(lag)

# %% [markdown]
# 4. Check Encoder

# %% [code]
is_image = (np.asarray(dset_samples[0]).ndim == 4)
if is_image:
    _, C, H, W = np.asarray(dset_samples[0]).shape
T_min = min(arr.shape[0] for arr in dset_samples)
T_max = max(arr.shape[0] for arr in dset_samples)
n_step = T_max

# %% [code]
encoder = encoder.to(device).eval()

# %% [code]
# not good enough...
trial_check = 0
T_check = 0
X1_ori = dset_samples[trial_check][T_check,:][None,:]
with torch.inference_mode():
    X1 = torch.from_numpy(X1_ori).to(device=device, dtype=torch.float32)
    X0 = encoder.rsample(X1).detach().cpu().numpy()
plt.plot(X1_ori.T, alpha = 0.5);
plt.plot(X0.T, alpha = 0.5);

# %% [code]
dim_to_keep = 3
mu_tmp, d, L, loading, bias = latent_mu(dset_samples, encoder.to(device), dim_to_keep, device, fullRes = True)
L_np = L.detach().cpu().numpy()
d_np = d.detach().cpu().numpy()
loading_np = loading.detach().cpu().numpy()
mu_all = get_mu_from_encoder(encoder, dset_samples, 64*64, 0)

# %% [code]
# mu_keep = get_mu_from_encoder(encoder, dset_samples, dim_to_keep, trial_check)
# plot_latent_time_series([mu_keep], alpha = 0.4, dims=np.arange(dim_to_keep), 
#                         dt=0.01, labels=None)

# %% [code]
# plot_traj_3d(mu_keep, title = 'trial = ' + str(trial_check))

# %% [code]
k_max = 137
k_target = 50
d_np = d.detach().cpu().numpy()
cum_energy = np.cumsum(d_np); K_eff = 1 + np.searchsorted(cum_energy, 0.95 * cum_energy[-1])
fig = plt.figure(figsize=(3, 3))
plt.plot(np.sqrt(d_np[0:k_max]))
plt.axvline(x=k_target-1, color='red', label = 'k_target')
plt.axvline(x=K_eff-1, color='orange', label = 'k_eff = ' + str(K_eff))
plt.title('sqrt{d}');
plt.legend()

# %% [code]
k_max = 200
dim_check_most = 200
k_target = 50
rows = loading.shape[0]
cols = dim_check_most
dim_check = np.arange(dim_check_most)
# rec = mu_group_recon(bias.detach().cpu().numpy(), loading_np, mu_all, dim_check, H=28, W=28)
# ball_idx = 0

fig, axs = plt.subplots(2, 2, figsize=(6, 5))
axs[0, 0].plot(np.sqrt(d_np[0:dim_check_most]))
axs[0, 0].axvline(x=k_target-1, color='red', label = 'k_target')
axs[0, 0].axvline(x=K_eff-1, color='orange', label = 'k_eff = ' + str(K_eff))
axs[0, 0].set_title('sqrt{d}');
axs[0, 0].legend()

im1 = axs[0, 1].imshow(np.abs(loading_np[:, 0:dim_check_most]), cmap='gray_r', aspect=cols / rows)
axs[0, 1].set_title('|loading|')
fig.colorbar(im1, ax=axs[0, 1])

axs[1, 0].plot(mu_all[:, :dim_check_most])
axs[1, 0].set_title('first ' + str(dim_check_most) + " dim of mu")

fig.suptitle('Encoder Diagnostic', fontsize=16)
plt.tight_layout()
plt.show()

# %% [markdown]
# 5. Flowed Latent

# %% [code]
import gc
gc.collect()
torch.cuda.empty_cache()
latent_traj_list = proj_to_latent(dset_samples, flow_net.to(device), 0.0, device)

# %% [code]
mu_traj_sim, d, L, loading, bias = latent_mu(latent_traj_list, encoder.to(device), dim_to_keep, device,
                                             fullRes = True, resid_tol=1e-15)

# %% [code]
mu_traj_flow_np = [mu_traj_sim[j].cpu().numpy() for j in range(len(mu_traj_sim))]
plot_latent_time_series(mu_traj_flow_np, alpha = 0.4, dims=np.arange(dim_to_keep), 
                        dt=0.01, labels=None)
plt.suptitle('flowed latent trajc for each trial')

# %% [code]
plot_mu_trajectories_gradient_steps_3d(
    mu_traj_flow_np,
    trial_indices=np.arange(len(mu_traj_flow_np)), 
    dims=(0,1,2),
    title="all flowed Latent trajectories",
    z_scale='auto',
    arrow_head_frac=0.08,
    arrow_length_ratio=0.9,
    elev_azim = None,
)

# %% [markdown]
# 4. Simulated Traj

# %% [code]
n_step

# %% [code]
tau = 1.0
traj_sim_tau1, lag_cov_list = generate_traj(dyn_net.to(device),
                                            dset_samples, n_step,
                                            tau, device, lag,
                                            include_x0_tau=False,
                                            oracle=False,
                                            flow_net=None,
                                            clamp_range=clamp_range,
                                            dset_samples_full=dset_samples_full,
                                            )

# %% [code]
trial_check = 0
fig, axs = plt.subplots(1, 2, figsize=(10, 3), constrained_layout=True)
axs[0].plot(dset_samples[trial_check]);      axs[0].set_title("dset_samples")
axs[1].plot(traj_sim_tau1[trial_check]);    axs[1].set_title("traj_sim_tau1")

# %% [code]
out_tau0_flow = proj_to_latent(traj_sim_tau1, flow_net.to(device), 0.0, device)

# %% [code]
tau = 0.0
traj_sim_tau0, _ = generate_traj(dyn_net.to(device),
                                 dset_samples, n_step,
                                 tau, device, lag,
                                 include_x0_tau=False,
                                 oracle=False,
                                 flow_net=flow_net.to(device),
                                 clamp_range=clamp_range,
                                 dset_samples_full=dset_samples_full,
                                 )

# %% [code]
trial_check = 0
fig, axs = plt.subplots(1, 3, figsize=(10, 3), constrained_layout=True)
axs[0].plot(dset_samples[trial_check]);      axs[0].set_title("dset_samples")
axs[1].plot(out_tau0_flow[trial_check]);    axs[1].set_title("flowed")
axs[2].plot(traj_sim_tau0[trial_check]);    axs[1].set_title("traj_sim_tau0")

# %% [code]
conditions=['DownLeft','Left','UpLeft','Up','UpRight','Right','DownRight']
loc = target_pos_sel
train_lab = np.zeros(target_pos_sel.shape[0])

for ii in range(target_pos_sel.shape[0]):
    if loc[ii,1] >35: vertstring='Up'
    elif loc[ii,1] <-35: vertstring='Down'
    else: vertstring=''
    
    if loc[ii,0] < -50: horzstring='Left'
    elif loc[ii,0] > 50: horzstring='Right'
    else: horzstring=''

    cond_string=vertstring+horzstring
    cond_num = np.argwhere([c == cond_string for c in conditions]).squeeze()
    train_lab[ii] = cond_num

# %% [code]
mu_traj_sim = latent_mu(traj_sim_tau0, encoder, dim_to_keep, device)
mu_traj_sim_np = [mu_traj_sim[j].cpu().numpy() for j in range(len(mu_traj_sim))]

# %% [code]
len(mu_traj_sim_np)

# %% [code]
target_pos_sel.shape

# %% [code]
train_lab_unique, _, mean_traj_sim, _ = mean_traj_by_labels(np.array(mu_traj_sim_np), train_lab)
mean_traj_list_sim = list(mean_traj_sim)

# %% [code]
plot_latent_time_series(mu_traj_sim_np, alpha = 0.4, dims=np.arange(dim_to_keep), 
                        dt=dt, labels=train_lab, cont_cmap=colorgrad)
plt.suptitle('simulated latent trajc for each trial')

# %% [markdown]
# 5. Plot

# %% [code]
def plot_mu_velocity_2d(
    mu,
    v_mu,
    trajectories=None,
    dims_2d=(0, 1),
    title=None,
    arrow_frac=0.05,
    arrow_size=1.2,
    linewidth=1.5,
    traj_alpha=0.4,
    traj_width=0.8,
    traj_color="gray",
    max_trajs=300,
    quiver_alpha=0.6,
    max_points=3000,
    figsize=(7, 6),
):
    fig, ax = plt.subplots(figsize=figsize, constrained_layout=True)
    ax.grid(False)

    traj_list = []
    if trajectories is not None:
        traj_source = trajectories
        for i, traj in enumerate(traj_source):
            if i >= max_trajs:
                break
            t_np = traj.detach().cpu().numpy() if hasattr(traj, "detach") else np.array(traj)
            traj_list.append(t_np)

    d1, d2 = dims_2d
    for t_np in traj_list:
        ax.plot(
            t_np[:, d1], t_np[:, d2],
            color=traj_color, alpha=traj_alpha, linewidth=traj_width, zorder=1
        )

    coords = mu[:, [d1, d2]]
    vecs   = v_mu[:, [d1, d2]]
    N = coords.shape[0]

    if max_points is not None and N > max_points:
        idx = np.random.choice(N, size=max_points, replace=False)
        coords_2d = coords[idx]
        vecs_2d   = vecs[idx]
    else:
        coords_2d = coords
        vecs_2d   = vecs

    speed = np.linalg.norm(vecs_2d, axis=1)
    eps = 1e-8
    dirs = vecs_2d / (speed[:, None] + eps)

    x_min, x_max = coords_2d[:, 0].min(), coords_2d[:, 0].max()
    y_min, y_max = coords_2d[:, 1].min(), coords_2d[:, 1].max()
    span = max(x_max - x_min, y_max - y_min) + eps
    L = arrow_frac * span
    U = dirs[:, 0] * L
    V = dirs[:, 1] * L

    base_width = 0.004
    base_headwidth = 4.0
    base_headlength = 6.0
    base_headaxislength = 5.0
    width = base_width * arrow_size

    norm = Normalize(vmin=float(speed.min()), vmax=float(speed.max()))
    q = ax.quiver(
        coords_2d[:, 0], coords_2d[:, 1], U, V, speed,
        cmap=cm.viridis, norm=norm, angles="xy", scale_units="xy", scale=1.0,
        width=width,
        headwidth=base_headwidth * arrow_size,
        headlength=base_headlength * arrow_size,
        headaxislength=base_headaxislength * arrow_size,
        linewidth=linewidth, alpha=quiver_alpha, zorder=2
    )
    cb = fig.colorbar(q, ax=ax, fraction=0.046, pad=0.04)
    cb.set_label(r"$\|\dot{\mu}\|$")

    ax.set_xlabel(fr"$\mu_{{{d1+1}}}$")
    ax.set_ylabel(fr"$\mu_{{{d2+1}}}$")
    if title:
        ax.set_title(title)
    ax.axis("equal")

    # fig.tight_layout()
    return fig, ax, q


def plot_mu_velocity_3d(
    mu,
    v_mu,
    trajectories=None,
    dims_3d=(0, 1, 2),
    title=None,
    arrow_frac=0.1,
    arrow_size=1.5,
    linewidth=1.5,
    base_arrow_length_ratio=0.3,
    traj_alpha=0.4,
    traj_width=0.8,
    quiver_alpha=0.6,
    traj_color="gray",
    max_trajs=300,
    max_points=3000,
    figsize=(10, 7),
):
    fig = plt.figure(figsize=figsize, constrained_layout=True)
    ax = fig.add_subplot(1, 1, 1, projection="3d")
    ax.set_box_aspect((1, 1, 1))
    ax.grid(False)
    for axis in (ax.xaxis, ax.yaxis, ax.zaxis):
        axis.pane.fill = False
        axis.pane.set_edgecolor((1, 1, 1, 0))
        axis._axinfo["grid"]["color"] = (1, 1, 1, 0)
        axis._axinfo["grid"]["linewidth"] = 0

    traj_list = []
    if trajectories is not None:
        traj_source = trajectories
        for i, traj in enumerate(traj_source):
            if i >= max_trajs:
                break
            t_np = traj.detach().cpu().numpy() if hasattr(traj, "detach") else np.array(traj)
            traj_list.append(t_np)

    d1, d2, d3 = dims_3d
    for t_np in traj_list:
        ax.plot(
            t_np[:, d1], t_np[:, d2], t_np[:, d3],
            color=traj_color, alpha=traj_alpha, linewidth=traj_width, zorder=1
        )

    coords = mu[:, [d1, d2, d3]]
    vecs   = v_mu[:, [d1, d2, d3]]
    N = coords.shape[0]

    if max_points is not None and N > max_points:
        idx = np.random.choice(N, size=max_points, replace=False)
        coords_3d = coords[idx]
        vecs_3d   = vecs[idx]
    else:
        coords_3d = coords
        vecs_3d   = vecs

    speed = np.linalg.norm(vecs_3d, axis=1)
    eps = 1e-8
    dirs = vecs_3d / (speed[:, None] + eps)

    mins = coords_3d.min(axis=0)
    maxs = coords_3d.max(axis=0)
    span = float(np.max(maxs - mins) + eps)

    L = arrow_frac * span
    U, V, W = dirs[:, 0], dirs[:, 1], dirs[:, 2]

    norm = Normalize(vmin=float(speed.min()), vmax=float(speed.max()))
    colors = cm.viridis(norm(speed))
    colors[:, 3] = quiver_alpha

    ax.quiver(
        coords_3d[:, 0], coords_3d[:, 1], coords_3d[:, 2],
        U, V, W,
        length=L, normalize=False, colors=colors,
        linewidth=linewidth * arrow_size,
        arrow_length_ratio=base_arrow_length_ratio * arrow_size,
        zorder=2
    )

    mappable = cm.ScalarMappable(norm=norm, cmap="viridis")
    mappable.set_array(speed)
    cb = fig.colorbar(mappable, ax=ax, fraction=0.035, pad=0.08)
    cb.set_label(r"$\|\dot{\mu}\|$")

    ax.set_xlabel(fr"$\mu_{{{d1+1}}}$")
    ax.set_ylabel(fr"$\mu_{{{d2+1}}}$")
    ax.set_zlabel(fr"$\mu_{{{d3+1}}}$")
    if title:
        ax.set_title(title)

    # fig.tight_layout()
    return fig, ax

# %% [code]
def plot_frame_compact(
    rows,                 # list like [dset_samples, out_imgs_tau0_flow, ...]
    trial_idx,
    idxs,                 # 1D int array, columns
    row_labels=None,      # list of strings, one per row
    mean_img=None,
    figsize=(6, 2.6),
    img_cmap="gray",
    vmin=None,
    vmax=None,
    border_from=None,
    border_lw=3.0,
    left=0.10, right=0.99, bottom=0.05, top=0.95,
    wspace=0.02, hspace=0.02,
):
    import numpy as np
    import matplotlib.pyplot as plt

    idxs = np.asarray(idxs).astype(int)
    n_rows = len(rows)
    n_cols = len(idxs)

    # --- global vmin/vmax (consistent across rows) ---
    if vmin is None or vmax is None:
        mn, mx = np.inf, -np.inf
        for data in rows:
            X = data[trial_idx]  # [T, 1, H, W] (torch or np)
            for i in idxs:
                img = X[i, 0]
                if hasattr(img, "detach"):
                    img = img.detach().cpu().numpy()
                else:
                    img = np.asarray(img)
                if mean_img is not None:
                    img = img + mean_img
                mn = min(mn, float(img.min()))
                mx = max(mx, float(img.max()))
        if vmin is None: vmin = mn
        if vmax is None: vmax = mx

    if border_from is not None:
        cmap = border_from.get_cmap()
        norm = border_from.norm
        col_colors = cmap(norm(idxs.astype(float)))
    else:
        col_colors = ["k"] * n_cols

    fig, axes = plt.subplots(n_rows, n_cols, figsize=figsize, squeeze=False)
    fig.subplots_adjust(left=left, right=right, bottom=bottom, top=top, wspace=wspace, hspace=hspace)

    for r, data in enumerate(rows):
        X = data[trial_idx]
        for c, i in enumerate(idxs):
            ax = axes[r, c]

            img = X[i, 0]
            if hasattr(img, "detach"):
                img = img.detach().cpu().numpy()
            else:
                img = np.asarray(img)
            if mean_img is not None:
                img = img + mean_img

            ax.imshow(img, cmap=img_cmap, vmin=vmin, vmax=vmax)

            # keep spines for colored border, so don't ax.axis("off")
            ax.set_xticks([]); ax.set_yticks([])

            # titles only on first row
            if r == 0:
                ax.set_title(f"t={i}")
            else:
                ax.set_title("")

            # colored border per column
            for sp in ax.spines.values():
                sp.set_visible(True)
                sp.set_linewidth(border_lw)
                sp.set_color(col_colors[c])

    # row labels on the left, centered per row (compact)
    if row_labels is not None:
        for r, lab in enumerate(row_labels):
            pos = axes[r, 0].get_position()
            y = 0.5 * (pos.y0 + pos.y1)
            x = pos.x0 - 0.02
            fig.text(x, y, lab, ha="right", va="center")

    return fig, axes

# %% [code]
def plot_mu_trajectories_gradient_steps_3d_v2(
    mu_traj_np,
    trial_indices=None,
    dims=(0, 1, 2),
    # line & markers
    linewidth=2.0, alpha=0.5,
    start_marker_size=36, start_marker_alpha=0.95,
    # colors / labels
    labels=None, label_mode='auto', cmap=None, vrange=None,
    start_color="#fdae61", end_color="#313695", circmap='twilight',
    # layout
    fig_w=7, fig_h=6, decimate=1,
    z_scale='auto', aspect_ratio=(1, 1, 1), elev_azim=None,
    title="Latent trajectories",
    # arrows
    n_arrows=3,
    arrow_size_frac=0.03,      # shaft length as fraction of mean axis span (data units)
    arrow_head_pts=12,         # head size in screen points (constant on screen)
    end_arrow_alpha=0.95
):
    import numpy as np
    import matplotlib.pyplot as plt
    from matplotlib.colors import Normalize, ListedColormap, BoundaryNorm, LinearSegmentedColormap
    from matplotlib.ticker import MaxNLocator, FuncFormatter
    from mpl_toolkits.mplot3d.art3d import Line3DCollection
    from matplotlib.patches import FancyArrowPatch
    from mpl_toolkits.mplot3d import proj3d

    class Arrow3D(FancyArrowPatch):
        """3D arrow with a screen-space head (constant size in points)."""
        def __init__(self, xs, ys, zs, *args, **kwargs):
            super().__init__((0, 0), (0, 0), *args, **kwargs)
            self._verts3d = (np.asarray(xs, float),
                             np.asarray(ys, float),
                             np.asarray(zs, float))
    
        def draw(self, renderer):
            # project 3D -> 2D every draw
            xs3d, ys3d, zs3d = self._verts3d
            xs, ys, zs = proj3d.proj_transform(xs3d, ys3d, zs3d, self.axes.get_proj())
            self.set_positions((xs[0], ys[0]), (xs[1], ys[1]))
            super().draw(renderer)
    
        def do_3d_projection(self, renderer=None):
            # used by mplot3d for depth-sorting
            xs3d, ys3d, zs3d = self._verts3d
            xs, ys, zs = proj3d.proj_transform(xs3d, ys3d, zs3d, self.axes.get_proj())
            self.set_positions((xs[0], ys[0]), (xs[1], ys[1]))
            return float(np.mean(zs))  # z-depth for sorting

    if trial_indices is None:
        trial_indices = range(len(mu_traj_np))

    # collect
    trajs, max_len = [], 0
    for idx in trial_indices:
        S = mu_traj_np[idx]
        if S is None or S.size == 0: continue
        trajs.append((idx, S)); max_len = max(max_len, S.shape[0])
    if not trajs: raise ValueError("No non-empty trials to plot.")

    # global ranges -> z scale
    xs_all, ys_all, zs_all = [], [], []
    for _, S in trajs:
        xs_all.extend(S[:, dims[0]]); ys_all.extend(S[:, dims[1]]); zs_all.extend(S[:, dims[2]])
    x_rng = (np.max(xs_all) - np.min(xs_all)) if xs_all else 1.0
    y_rng = (np.max(ys_all) - np.min(ys_all)) if ys_all else 1.0
    z_rng = (np.max(zs_all) - np.min(zs_all)) if zs_all else 1.0
    if z_scale == 'auto':
        xy_avg = (x_rng + y_rng) / 2.0
        scale_factor = (xy_avg / z_rng) if z_rng > 0 else 1.0
    elif z_scale == 'none':
        scale_factor = 1.0
    else:
        scale_factor = float(z_scale)

    # colors
    use_labels = labels is not None
    if use_labels:
        lab_for = {}
        if isinstance(labels, dict):
            for i, _ in trajs:
                if i in labels: lab_for[i] = labels[i]
        else:
            arr = np.asarray(labels)
            for i, _ in trajs:
                if i < arr.shape[0]: lab_for[i] = arr[i]
        vals_exist = [lab_for[i] for i, _ in trajs if i in lab_for]
        if not vals_exist: raise ValueError("Provided labels are empty or misaligned.")
        if label_mode == 'continuous':
            use_cont = True
        elif label_mode == 'categorical':
            use_cont = False
        else:
            all_int = all(isinstance(v, (int, np.integer)) for v in vals_exist)
            all_num = all(isinstance(v, (int, float, np.integer, np.floating)) for v in vals_exist)
            use_cont = (not all_int) and all_num
        if use_cont:
            vals = np.array(vals_exist, float)
            vmin, vmax = (float(np.nanmin(vals)), float(np.nanmax(vals))) if vrange is None else vrange
            if np.isclose(vmin, -180.0) and np.isclose(vmax, 180.0) and cmap is None:
                cmap_obj = plt.get_cmap(circmap)
            else:
                default_name = plt.rcParams.get('image.cmap', 'viridis')
                cmap_obj = plt.get_cmap(default_name if cmap is None else cmap)
            norm = Normalize(vmin=vmin, vmax=vmax); cat_to_i = None
        else:
            cats = []
            for i, _ in trajs:
                if i in lab_for and lab_for[i] not in cats: cats.append(lab_for[i])
            base = plt.get_cmap('tab10') if cmap is None else plt.get_cmap(cmap)
            cmap_obj = ListedColormap([base(i % base.N) for i in range(len(cats))])
            norm = BoundaryNorm(np.arange(len(cats)+1)-0.5, len(cats))
            cat_to_i = {c:i for i,c in enumerate(cats)}
    else:
        cmap_obj = LinearSegmentedColormap.from_list("o2b", [start_color, end_color])
        norm = Normalize(vmin=0, vmax=(1 if max_len <= 1 else max_len - 1)); cat_to_i = None

    # figure
    fig = plt.figure(figsize=(fig_w, fig_h))
    try:
        fig.set_constrained_layout(True)  # more reliable than tight_layout for 3D + colorbar
    except Exception:
        pass
    ax = fig.add_subplot(111, projection="3d")
    ax.grid(False)
    for a in (ax.xaxis, ax.yaxis, ax.zaxis):
        try: a.pane.fill = False; a.pane.set_edgecolor('none')
        except Exception: pass

    # plot lines & cache arrow starts
    arrows = []  # (x0,y0,z0, ux,uy,uz, color)
    for idx, S in trajs:
        X = np.asarray(S[:, dims[0]], float)
        Y = np.asarray(S[:, dims[1]], float)
        Z = np.asarray(S[:, dims[2]], float) * scale_factor
        if decimate > 1: X, Y, Z = X[::decimate], Y[::decimate], Z[::decimate]
        T = len(X)
        if T == 0: continue

        if not use_labels:
            start_col = cmap_obj(norm(0))
        else:
            if isinstance(labels, dict):
                has_lab = (idx in labels); lab_val = labels.get(idx, None)
            else:
                has_lab = (idx < np.asarray(labels).shape[0]); lab_val = labels[idx] if has_lab else None
            start_col = (cmap_obj(norm(cat_to_i[lab_val])) if (cat_to_i is not None and has_lab)
                         else (cmap_obj(norm(float(lab_val))) if has_lab else (0.6,0.6,0.6,1.0)))

        ax.scatter(X[0], Y[0], Z[0], s=start_marker_size, marker="o",
                   facecolor=start_col, edgecolor="black", linewidths=0.5,
                   alpha=start_marker_alpha, zorder=3)
        if T == 1: continue

        if not use_labels:
            pts = np.column_stack([X, Y, Z])
            segs = np.stack([pts[:-1], pts[1:]], axis=1)
            seg_steps = np.arange(segs.shape[0])
            cols = cmap_obj(norm(seg_steps))
            lc = Line3DCollection(segs, linewidths=linewidth, alpha=alpha)
            lc.set_colors(cols); ax.add_collection3d(lc)
            traj_col = cmap_obj(norm(T-1))
        else:
            traj_col = start_col
            ax.plot(X, Y, Z, lw=linewidth, alpha=alpha, color=traj_col)

        if n_arrows and T >= 2:
            lo, hi = 1, T-2
            if hi < lo: lo = hi
            idxs = np.unique(np.linspace(lo, hi, int(n_arrows), dtype=int))
            for k in idxs:
                x0, y0, z0 = X[k], Y[k], Z[k]
                vx, vy, vz = X[k+1]-x0, Y[k+1]-y0, Z[k+1]-z0
                nrm = float(np.linalg.norm([vx,vy,vz]))
                if nrm < 1e-12: continue
                ux, uy, uz = vx/nrm, vy/nrm, vz/nrm
                col = (cmap_obj(norm(int(k))) if not use_labels else traj_col)
                arrows.append((x0, y0, z0, ux, uy, uz, col))

    # axes/view
    ax.set_xlabel(f"μ{dims[0]+1}"); ax.set_ylabel(f"μ{dims[1]+1}"); ax.set_zlabel(f"μ{dims[2]+1}")
    ax.set_title(title)

    # autoscale to plotted points
    xcat = []; ycat = []; zcat = []
    for _, S in trajs:
        x = np.asarray(S[:, dims[0]], float)
        y = np.asarray(S[:, dims[1]], float)
        z = np.asarray(S[:, dims[2]], float) * scale_factor
        if decimate > 1: x, y, z = x[::decimate], y[::decimate], z[::decimate]
        if x.size: xcat.append(x); ycat.append(y); zcat.append(z)
    if xcat:
        xcat = np.concatenate(xcat); ycat = np.concatenate(ycat); zcat = np.concatenate(zcat)
        ax.auto_scale_xyz(xcat, ycat, zcat, had_data=False)
        dx = max(1e-12, xcat.max()-xcat.min())
        dy = max(1e-12, ycat.max()-ycat.min())
        dz = max(1e-12, zcat.max()-zcat.min())
        pad = 0.03
        ax.set_xlim(xcat.min()-pad*dx, xcat.max()+pad*dx)
        ax.set_ylim(ycat.min()-pad*dy, ycat.max()+pad*dy)
        ax.set_zlim(zcat.min()-pad*dz, zcat.max()+pad*dz)

    ax.xaxis.set_major_locator(MaxNLocator(nbins=5, prune='both'))
    ax.yaxis.set_major_locator(MaxNLocator(nbins=5, prune='both'))
    ax.zaxis.set_major_locator(MaxNLocator(nbins=5, prune='both'))
    try: ax.set_box_aspect('auto' if aspect_ratio == (1,1,1) else aspect_ratio)
    except Exception: pass
    if elev_azim is not None: ax.view_init(elev=elev_azim[0], azim=elev_azim[1])
    if scale_factor != 1.0:
        ax.zaxis.set_major_formatter(FuncFormatter(lambda v, p: f"{v/scale_factor:.2f}"))

    # ---- draw arrows with screen-space heads (no fat/skinny effect) ----
    if arrows:
        xlim, ylim, zlim = ax.get_xlim(), ax.get_ylim(), ax.get_zlim()
        Rx = float(xlim[1]-xlim[0]); Ry = float(ylim[1]-ylim[0]); Rz = float(zlim[1]-zlim[0])
        fixed_len = max(1e-12, arrow_size_frac * (Rx + Ry + Rz) / 3.0)
        for x0, y0, z0, ux, uy, uz, col in arrows:
            x1, y1, z1 = x0 + ux*fixed_len, y0 + uy*fixed_len, z0 + uz*fixed_len
            a = Arrow3D([x0, x1], [y0, y1], [z0, z1],
                        arrowstyle="-|>", mutation_scale=arrow_head_pts,
                        lw=0.8*linewidth, color=col, alpha=end_arrow_alpha)
            ax.add_artist(a)

    # colorbar
    sm = plt.cm.ScalarMappable(cmap=cmap_obj, norm=norm); sm.set_array([])
    cbar = fig.colorbar(sm, ax=ax, fraction=0.035, pad=0.08, shrink=0.85)
    if not use_labels:
        ticks = [0, 1] if max_len <= 2 else [0, (max_len-1)//2, max_len-1]
        cbar.set_ticks(ticks); cbar.set_ticklabels([str(t) for t in ticks])
        cbar.set_label("Step index")
    else:
        if cat_to_i is not None:
            cbar.set_ticks(range(len(cat_to_i)))
            inv = [k for k,_ in sorted(cat_to_i.items(), key=lambda kv: kv[1])]
            cbar.set_ticklabels([str(k) for k in inv]); cbar.set_label("Category")
        else:
            cbar.set_label("Label" if vrange is None else (vrange,))

    # fig.tight_layout()

# %% [code]
plotFolder = "paper_fig/maze"

# %% [code]
# 1. sqrt{d}
fig = plt.figure(figsize=(4, 4))
plt.plot(np.sqrt(d_np[0:k_max]))
plt.axvline(x=k_target-1, color='red', label = 'k_target')
plt.axvline(x=K_eff-1, color='orange', label = 'k_eff = ' + str(K_eff))
plt.title('sqrt{d}');
plt.legend()

plt.savefig(plotFolder + "/1_sqrt_d.pdf")
plt.savefig(plotFolder + "/1_sqrt_d.png")

# %% [code]
latent_traj_list[0].shape

# %% [code]
dim_to_keep = 3  # μ-dims you kept elsewhere
mu_pts_flow, v_mu_flow = compute_mu_velocity_field(
    latent_traj_list=latent_traj_list,
    dyn_net=dyn_net,
    encoder=encoder,
    device=device,
    dim_preserve=dim_to_keep,
    lag=lag,
    t_plot=0.0,
)

# %% [code]
# 2. compressed flow field in mu (3D)
plot_mu_velocity_3d(mu_pts_flow, v_mu_flow, trajectories=[mu_pts_flow], dims_3d=(0,1,2),
                    title="flowed: mu1–mu2-mu3 field (tau =0.0)", figsize=(5, 5), quiver_alpha=0.5)

plt.savefig(plotFolder + "/2_flow_v_3d.pdf")
plt.savefig(plotFolder + "/2_flow_v_3d.png")

# %% [code]
mu_pts_sim, v_mu_sim = compute_mu_velocity_field(
    latent_traj_list=traj_sim_tau0,
    dyn_net=dyn_net,
    encoder=encoder,
    device=device,
    dim_preserve=dim_to_keep,
    lag=lag,
    t_plot=0.0,
)

# %% [code]
# 3. compressed flow field in mu (3D)
plot_mu_velocity_3d(mu_pts_sim, v_mu_sim, trajectories=[mu_pts_sim], dims_3d=(0,1,2),
                    title="simulated: mu1–mu2-mu3 field (tau =0.0)", figsize=(5, 5), quiver_alpha=0.5)

plt.savefig(plotFolder + "/3_sim_v_3d.pdf")
plt.savefig(plotFolder + "/3_sim_v_3d.png")

# %% [code]
train_lab_unique, _, mean_traj_flow, _ = mean_traj_by_labels(np.array(mu_traj_flow_np), train_lab)
mean_traj_list_flow = list(mean_traj_flow)

# %% [code]
plot_mu_trajectories_gradient_steps_3d_v2(
    mean_traj_list_flow,
    labels=train_lab_unique,
    trial_indices=np.arange(len(mean_traj_list_flow)), 
    dims=(0,1,2),
    title="Flowed Mean Latent trajectories over target location",
    z_scale='auto',  # Automatically scales z to be visually comparable to x and y
    aspect_ratio=(1, 1, 1),
    fig_w=5, fig_h=5,
    circmap = colorgrad,
    # elev_azim = (10,45), # (20,45)
    n_arrows = 5,
    arrow_size_frac=0.02
)

plt.savefig(plotFolder + "/4_LatentFlow_target_3d.pdf")
plt.savefig(plotFolder + "/4_LatentFlow_target_3d.png")

# %% [code]
plot_mu_trajectories_gradient_steps_3d_v2(
    mean_traj_list_sim,
    labels=train_lab_unique,
    trial_indices=np.arange(len(mean_traj_list_sim)), 
    dims=(0,1,2),
    title="Simulated Mean Latent trajectories over target location",
    z_scale='auto',  # Automatically scales z to be visually comparable to x and y
    aspect_ratio=(1, 1, 1),
    fig_w=5, fig_h=5,
    circmap = colorgrad,
    # elev_azim = (10,45), # (20,45)
    n_arrows = 5,
    arrow_size_frac=0.02,
)

plt.savefig(plotFolder + "/5_LatentSim_target_3d.pdf")
plt.savefig(plotFolder + "/5_LatentSim_target_3d.png")

# %% [code]
fig = plt.figure(figsize=(5, 5), constrained_layout=True)
ax3d = fig.add_subplot(111, projection='3d')
ax3d.set_box_aspect((1, 1, 1))
ax3d.grid(False)
for axis in (ax3d.xaxis, ax3d.yaxis, ax3d.zaxis):
    axis.pane.fill = False
    axis.pane.set_edgecolor((1, 1, 1, 0))
    axis._axinfo["grid"]["color"] = (1, 1, 1, 0)
    axis._axinfo["grid"]["linewidth"] = 0

ax3d.scatter(mu_pts_sim[:, 0], mu_pts_sim[:, 1], mu_pts_sim[:, 2],
             c="k", s=2, alpha=0.005)

labs_show = [0.0, 3.0, 6.0]

cmap_obj = plt.get_cmap(plt.rcParams.get("image.cmap", "viridis"))
norm = Normalize(vmin=float(np.min(train_lab)), vmax=float(np.max(train_lab)))

rep_idx = {}
for lab in labs_show:
    idxs = np.flatnonzero(train_lab == lab)   # indices in 0..591
    if idxs.size == 0:
        continue

    arr = np.stack([mu_traj_sim_np[i] for i in idxs], axis=0)  # [n, T, 3]
    m = arr.mean(axis=0)                                       # [T, 3]
    d = ((arr - m) ** 2).mean(axis=(1, 2))                     # [n]
    i_rep = int(idxs[np.argmin(d)])                            # traj index in original list
    rep_idx[lab] = i_rep

    tr = mu_traj_sim_np[i_rep]
    col = cmap_obj(norm(float(lab)))

    ax3d.plot(tr[:, 0], tr[:, 1], tr[:, 2], color=col, lw=2.5, alpha=0.95)
    ax3d.scatter(tr[0, 0], tr[0, 1], tr[0, 2],
                 s=40, facecolor=col, edgecolor="k", linewidths=0.6, alpha=1.0)

fig.subplots_adjust(right=0.86)
ax3d.set_title('simulated mu')

plt.savefig(plotFolder + "/6_3traj_3d_train.pdf")
plt.savefig(plotFolder + "/6_3traj_3d_train.png")

# %% [code]
row_labels = ["data", r"$\tau = 1.0$", r"$\tau = 0.0$"]

labs_valid = [lab for lab in labs_show if lab in rep_idx]
ncol = len(labs_valid)

# compact, wide cells
cell_w = 2.5
cell_h = 1.15
fig, axes = plt.subplots(
    3, ncol,
    figsize=(cell_w * ncol, cell_h * 3),
    constrained_layout=True
)
fig.subplots_adjust(left=0.10)  # small room for row labels

for j, lab in enumerate(labs_valid):
    idx = rep_idx[lab]
    col = cmap_obj(norm(float(lab)))
    imgs = [dset_samples[idx], traj_sim_tau1[idx], traj_sim_tau0[idx]]

    axes[0, j].set_title(f"lab {int(lab)}", pad=2)

    for i in range(3):
        ax = axes[i, j]
        ax.imshow(imgs[i].T, cmap="gray_r", aspect="auto")  # key line
        ax.set_xticks([]); ax.set_yticks([])

        for sp in ax.spines.values():
            sp.set_linewidth(2.5)
            sp.set_edgecolor(col)

# row labels on the left (only first column)
for i, rlab in enumerate(row_labels):
    ax0 = axes[i, 0]
    ax0.set_ylabel(rlab, rotation=0, labelpad=26)
    ax0.yaxis.set_label_coords(-0.22, 0.5)

plt.savefig(plotFolder + "/7_3traj_img_train.pdf")
plt.savefig(plotFolder + "/7_3traj_img_train.png")

# %% [markdown]
### test

# %% [code]
folder_data_test = 'data/mc_maze3_test'
dataset_samples = np.load(folder_data_test + "/dataset_samples.npz", allow_pickle=True)
data_raw = dataset_samples['samples']
dset_samples_test_full = []
for ii in range(len(data_raw)):
    dset_samples_test_full.append(np.asarray(data_raw[ii], dtype=np.float64))

# %% [code]
len(dset_samples_test_full)

# %% [code]
dset_samples_test = []
for ii in range(len(dset_samples_test_full)):
    dset_samples_test.append(dset_samples_test_full[ii][lag:,:])

# %% [code]
dset_samples_test[0].shape

# %% [code]
tau = 1.0
traj_sim_tau1_test, lag_cov_list = generate_traj(dyn_net.to(device),
                                            dset_samples_test, n_step,
                                            tau, device, lag,
                                            include_x0_tau=False,
                                            oracle=False,
                                            flow_net=None,
                                            clamp_range=clamp_range,
                                            dset_samples_full=dset_samples_test_full,
                                            )

tau = 0.0
traj_sim_tau0_test, _ = generate_traj(dyn_net.to(device),
                                 dset_samples_test, n_step,
                                 tau, device, lag,
                                 include_x0_tau=False,
                                 oracle=False,
                                 flow_net=flow_net.to(device),
                                 clamp_range=clamp_range,
                                 dset_samples_full=dset_samples_test_full,
                                 )

# %% [code]
mu_traj_sim_test = latent_mu(traj_sim_tau0_test, encoder, dim_to_keep, device)
mu_traj_sim_np_test = [mu_traj_sim_test[j].cpu().numpy() for j in range(len(mu_traj_sim_test))]

# %% [code]
mu_pts_sim_test, v_mu_sim_test = compute_mu_velocity_field(
    latent_traj_list=traj_sim_tau0_test,
    dyn_net=dyn_net,
    encoder=encoder,
    device=device,
    dim_preserve=dim_to_keep,
    lag=lag,
    t_plot=0.0,
)

# %% [code]
conditions=['DownLeft','Left','UpLeft','Up','UpRight','Right','DownRight']
test_meta = np.load(folder_data_test + "/meta_test.npy", allow_pickle=True).tolist()
trial_idx_use = np.array(test_meta['trial_info']['trial_version']) == 0

trial_type_use = np.array(test_meta['trial_info']['trial_type'])[trial_idx_use]
indices = np.where(trial_idx_use)[0]

target_pos_sel_test = [test_meta['trial_info']['target_pos'][idx] for idx in indices]
target_pos_sel_test = np.array(target_pos_sel_test).squeeze(1).astype(np.float32)

loc = target_pos_sel_test
test_lab = np.zeros(target_pos_sel_test.shape[0])

for ii in range(target_pos_sel_test.shape[0]):
    if loc[ii,1] >35: vertstring='Up'
    elif loc[ii,1] <-35: vertstring='Down'
    else: vertstring=''
    
    if loc[ii,0] < -50: horzstring='Left'
    elif loc[ii,0] > 50: horzstring='Right'
    else: horzstring=''

    cond_string=vertstring+horzstring
    cond_num = np.argwhere([c == cond_string for c in conditions]).squeeze()
    test_lab[ii] = cond_num

# %% [code]
labs_show = [0.0, 3.0, 6.0]

# same color code as your train plot (continuous label -> cmap(norm(label)))
cmap_obj = plt.get_cmap(plt.rcParams.get("image.cmap", "viridis"))
norm = Normalize(vmin=float(np.min(train_lab)), vmax=float(np.max(train_lab)))

# ---------- 1) TRAIN means per lab (in mu-space) ----------
train_mean_mu = {}
for lab in labs_show:
    idxs = np.flatnonzero(train_lab == lab)
    if idxs.size == 0:
        continue
    arr = np.stack([mu_traj_sim_np[i] for i in idxs], axis=0)  # [n, T, 3]
    train_mean_mu[lab] = arr.mean(axis=0)                      # [T, 3]

# ---------- 2) pick TEST representative by distance to TRAIN mean ----------
rep_idx_test = {}
for lab in labs_show:
    if lab not in train_mean_mu:
        continue
    idxs = np.flatnonzero(test_lab == lab)   # indices in test list
    if idxs.size == 0:
        continue

    m = train_mean_mu[lab]  # [T, 3]

    # assume fixed length; if not, safely truncate to common T
    arr = np.stack([mu_traj_sim_np_test[i] for i in idxs], axis=0)  # [n, Tt, 3]
    T = min(arr.shape[1], m.shape[0])
    d = ((arr[:, :T, :] - m[:T, :]) ** 2).mean(axis=(1, 2))         # [n]
    i_rep = int(idxs[np.argmin(d)])
    rep_idx_test[lab] = i_rep

# ===================== Plot A: 3D scatter + reps (TEST) =====================
fig = plt.figure(figsize=(5, 5), constrained_layout=True)
ax3d = fig.add_subplot(111, projection='3d')
ax3d.set_box_aspect((1, 1, 1))
ax3d.grid(False)
for axis in (ax3d.xaxis, ax3d.yaxis, ax3d.zaxis):
    axis.pane.fill = False
    axis.pane.set_edgecolor((1, 1, 1, 0))
    axis._axinfo["grid"]["color"] = (1, 1, 1, 0)
    axis._axinfo["grid"]["linewidth"] = 0

ax3d.scatter(mu_pts_sim_test[:, 0], mu_pts_sim_test[:, 1], mu_pts_sim_test[:, 2],
             c="k", s=2, alpha=0.005)

for lab in labs_show:
    if lab not in rep_idx_test:
        continue
    i_rep = rep_idx_test[lab]
    tr = mu_traj_sim_np_test[i_rep]
    col = cmap_obj(norm(float(lab)))

    ax3d.plot(tr[:, 0], tr[:, 1], tr[:, 2], color=col, lw=2.5, alpha=0.95)
    ax3d.scatter(tr[0, 0], tr[0, 1], tr[0, 2],
                 s=40, facecolor=col, edgecolor="k", linewidths=0.6, alpha=1.0)

fig.subplots_adjust(right=0.86)
ax3d.set_title('simulated mu (test)')

plt.savefig(plotFolder + "/8_3traj_3d_test.pdf")
plt.savefig(plotFolder + "/8_3traj_3d_test.png")

# %% [code]
row_labels = ["data", r"$\tau = 1.0$", r"$\tau = 0.0$"]
labs_valid = [lab for lab in labs_show if lab in rep_idx_test]
ncol = len(labs_valid)

cell_w = 2.5
cell_h = 1.15
fig, axes = plt.subplots(
    3, ncol,
    figsize=(cell_w * ncol, cell_h * 3),
    constrained_layout=True
)
fig.subplots_adjust(left=0.10)

for j, lab in enumerate(labs_valid):
    idx = rep_idx_test[lab]
    col = cmap_obj(norm(float(lab)))

    imgs = [dset_samples_test[idx], traj_sim_tau1_test[idx], traj_sim_tau0_test[idx]]

    axes[0, j].set_title(f"lab {int(lab)}", pad=2)

    for i in range(3):
        ax = axes[i, j]
        ax.imshow(imgs[i].T, cmap="gray_r", aspect="auto")
        ax.set_xticks([]); ax.set_yticks([])

        for sp in ax.spines.values():
            sp.set_linewidth(2.5)
            sp.set_edgecolor(col)

for i, rlab in enumerate(row_labels):
    ax0 = axes[i, 0]
    ax0.set_ylabel(rlab, rotation=0, labelpad=26)
    ax0.yaxis.set_label_coords(-0.22, 0.5)

plt.savefig(plotFolder + "/9_3traj_img_test.pdf")
plt.savefig(plotFolder + "/9_3traj_img_test.png")

print("rep_idx_test:", rep_idx_test)

# %% [markdown]
### R2 for simulated and flowed

# %% [code]
from sklearn.linear_model import RidgeCV
def train_asses_linear_predictions(train_embeddings,test_embeddings,train_cursor_pos,test_cursor_pos,lag_ms = 100,spike_bin_size_ms=5):

    """
    train and assess linear predictions of cursor position and velocity from model embeddings. accepts as input:
        :train_embeddings: np.array, T1 (number of trials) x L1 (length of trials) x d (embedding dimension)
        :test_embeddings: np.array,  T2 x L2 x d
        :train_cursor_pos: np.array, T1 x L1 x 2 (x,y position of cursor)
        :test_cursor_pos: np.array, T2 x L2 x 2
        :lag_ms: int, time lag in ms of cursor from embeddings. Will predict cursor from embeddings `lag_ms` ms prior
        :spike_bin_size_ms: int, size of spike bin in ms. used to compute lag (in samples) of cursor from embeddings

    returns:
        :(train_r2_pos,test_r2_pos): (np.array,np.array), r^2 for cursor position predictions for all (train trials,test trials)
        :(train_r2_vel,test_r2_vel): (np.array,np.array), r^2 for cursor velocity predictions for all (train trials,test trials)
        :(train_preds_pos,test_preds_pos): (np.array,np.array), cursor position predictions for all (train_trials,test trials)
        :(train_preds_vel,test_preds_vel): (np.array,np.array), cursor velocity predictions for all (train_trials,test trials)
    """

    vel_train = np.diff(train_cursor_pos,axis=1)
    vel_test = np.diff(test_cursor_pos,axis=1)
    
    lag_samples = lag_ms//spike_bin_size_ms

    ### fit position regression
    train_r2_pos,test_r2_pos = [],[]
    train_preds_pos,test_preds_pos = [],[]
    
    lr_pos = RidgeCV(alphas=np.logspace(-4,0,5))

    lr_pos.fit(np.reshape(train_embeddings[:,:-lag_samples,:],(-1,3)),np.reshape(train_cursor_pos[:,lag_samples:],(-1,2)))

    for x,y in zip(train_embeddings[:,:-lag_samples,:],train_cursor_pos[:,lag_samples:,:]):

        train_r2_pos.append(lr_pos.score(x,y))
        train_preds_pos.append(lr_pos.predict(x))


    for x,y in zip(test_embeddings[:,:-lag_samples,:],test_cursor_pos[:,lag_samples:,:]):
    
        test_r2_pos.append(lr_pos.score(x,y))
        test_preds_pos.append(lr_pos.predict(x))

    train_r2_pos,test_r2_pos = np.hstack(train_r2_pos),np.hstack(test_r2_pos)
    train_preds_pos,test_preds_pos = np.stack(train_preds_pos,axis=0),np.stack(test_preds_pos,axis=0)

    
    ### fit velocity regression
    
    train_r2_vel,test_r2_vel = [],[]
    train_preds_vel,test_preds_vel = [],[]
    
    lr_vel = RidgeCV(alphas=np.logspace(-4,0,5))

    lr_vel.fit(np.reshape(train_embeddings[:,:-lag_samples-1,:],(-1,3)),np.reshape(vel_train[:,lag_samples:],(-1,2)))

    for x,y in zip(train_embeddings[:,:-lag_samples-1,:],vel_train[:,lag_samples:,:]):

        pred = lr_vel.predict(x)
        sse = ((pred - y)**2).sum(axis=0)
        
        ssq = (y**2).sum(axis=0)
        #print(sse.shape,ssq.shape)
        train_r2_vel.append(np.nanmean(1 - sse/ssq))
        train_preds_vel.append(pred)


    for x,y in zip(test_embeddings[:,:-lag_samples-1,:],vel_test[:,lag_samples:,:]):
    
        pred = lr_vel.predict(x)
        sse = ((pred - y)**2).sum(axis=0)
        ssq = (y**2).sum(axis=0)

        test_r2_vel.append(np.nanmean(1 - sse/ssq))
        test_preds_vel.append(pred)

    train_r2_vel,test_r2_vel = np.hstack(train_r2_vel),np.hstack(test_r2_vel)
    train_preds_vel,test_preds_vel = np.stack(train_preds_vel,axis=0),np.stack(test_preds_vel,axis=0)

    return (train_r2_pos,test_r2_pos),(train_r2_vel,test_r2_vel),(train_preds_pos,test_preds_pos),(train_preds_vel,test_preds_vel)

# %% [code]
latent_traj_list_test = proj_to_latent(dset_samples_test,
                                       flow_net.to(device), 0.0, device)

# %% [code]
trial_idx_use_train = np.array(train_meta['trial_info']['trial_version']) == 0
trial_type_use = np.array(train_meta['trial_info']['trial_type'])[trial_idx_use_train]
indices_train = np.where(trial_idx_use_train)[0]
cursor_pos_train = [train_meta['behavior']['cursor_pos'][idx] for idx in indices_train]
cursor_pos_train_3d = np.stack([np.asarray(a)[:100, :] for a in cursor_pos_train],
                               axis=0)

# %% [code]
trial_idx_use_test = np.array(test_meta['trial_info']['trial_version']) == 0
trial_type_use = np.array(test_meta['trial_info']['trial_type'])[trial_idx_use_test]
indices_test = np.where(trial_idx_use_test)[0]
cursor_pos_test = [test_meta['behavior']['cursor_pos'][idx] for idx in indices_test]
cursor_pos_test_3d = np.stack([np.asarray(a)[:100, :] for a in cursor_pos_test],
                              axis=0)

# %% [markdown]
#### Flowed

# %% [code]
# :train_embeddings: np.array, T1 (number of trials) x L1 (length of trials) x d (embedding dimension)
# :test_embeddings: np.array,  T2 x L2 x d
# :train_cursor_pos: np.array, T1 x L1 x 2 (x,y position of cursor)
# :test_cursor_pos: np.array, T2 x L2 x 2
# :lag_ms: int, time lag in ms of cursor from embeddings. Will predict cursor from embeddings `lag_ms` ms prior
# :spike_bin_size_ms: int, size of spike bin in ms. used to compute lag (in samples) of cursor from embeddings

# %% [code]
latent_traj_list_full = proj_to_latent(dset_samples_full, flow_net.to(device),
                                       0.0, device)
latent_traj_list_test_full = proj_to_latent(dset_samples_test_full, flow_net.to(device),
                                            0.0, device)

# %% [code]
print(len(latent_traj_list_full))
print(latent_traj_list_full[0].shape)

# %% [code]
print(len(latent_traj_list_test_full))
print(latent_traj_list_test_full[0].shape)

# %% [code]
mu_traj_flow_train, _, _, _, _ = latent_mu(latent_traj_list_full,
                                           encoder.to(device),
                                           dim_to_keep, device,
                                           fullRes = True, resid_tol=1e-15)

# %% [code]
mu_traj_flow_train_3d = np.stack([np.asarray(a) for a in mu_traj_flow_train], axis=0)

# %% [code]
mu_traj_flow_train_3d.shape

# %% [code]
mu_traj_flow_test, _, _, _, _ = latent_mu(latent_traj_list_test_full,
                                          encoder.to(device),
                                          dim_to_keep, device,
                                          fullRes = True, resid_tol=1e-15)

# %% [code]
mu_traj_flow_test_3d = np.stack([np.asarray(a) for a in mu_traj_flow_test], axis=0)

# %% [code]
mu_traj_flow_test_3d.shape

# %% [code]
cursor_pos_train_3d.shape

# %% [code]
cursor_pos_test_3d.shape

# %% [code]
from sklearn.linear_model import RidgeCV

(train_r2_pos,test_r2_pos),(train_r2_vel,test_r2_vel),(train_preds_pos,test_preds_pos),(train_preds_vel,test_preds_vel)=\
train_asses_linear_predictions(mu_traj_flow_train_3d,
                               mu_traj_flow_test_3d,
                               cursor_pos_train_3d,
                               cursor_pos_test_3d)

# %% [code]
np.mean(test_r2_vel)

# %% [code]
np.std(test_r2_vel)

# %% [code]
test_r2_vel.shape

# %% [code]
np.percentile(test_r2_vel, [25, 50, 75])

# %% [code]
np.save('test_r2_vel_flowed.npy', test_r2_vel)

# %% [code]


# %% [markdown]
### Simulated

# %% [code]
dset_samples_full[0].shape

# %% [code]
dset_samples_test_full[0].shape

# %% [code]
print(len(traj_sim_tau0))
print(traj_sim_tau0[0].shape)

# %% [code]
print(len(traj_sim_tau0_test))
print(traj_sim_tau0_test[0].shape)

# %% [code]
traj_sim_tau0[0].shape

# %% [code]
traj_sim_tau0_list = [
    np.concatenate([dset_samples_full[i][:10, :], traj_sim_tau0[i]], axis=0)
    for i in range(len(dset_samples_full))
]

# %% [code]
traj_sim_tau0_test_list = [
    np.concatenate([dset_samples_test_full[i][:10, :], traj_sim_tau0_test[i]], axis=0)
    for i in range(len(dset_samples_test_full))
]

# %% [code]
print(len(traj_sim_tau0_list))
print(traj_sim_tau0_list[0].shape)

# %% [code]
print(len(traj_sim_tau0_test_list))
print(traj_sim_tau0_test_list[0].shape)

# %% [code]
mu_traj_sim_train, _, _, _, _ = latent_mu(traj_sim_tau0_list,
                                          encoder.to(device),
                                          dim_to_keep, device,
                                          fullRes = True, resid_tol=1e-15)

# %% [code]
mu_traj_sim_test, _, _, _, _ = latent_mu(traj_sim_tau0_test_list,
                                         encoder.to(device),
                                         dim_to_keep, device,
                                         fullRes = True, resid_tol=1e-15)

# %% [code]
mu_traj_sim_train_3d = np.stack([np.asarray(a) for a in mu_traj_sim_train], axis=0)
mu_traj_sim_test_3d = np.stack([np.asarray(a) for a in mu_traj_sim_test], axis=0)

# %% [code]
(train_r2_pos,test_r2_pos),(train_r2_vel,test_r2_vel),(train_preds_pos,test_preds_pos),(train_preds_vel,test_preds_vel)=\
train_asses_linear_predictions(mu_traj_sim_train_3d,
                               mu_traj_sim_test_3d,
                               cursor_pos_train_3d,
                               cursor_pos_test_3d)

# %% [code]
np.mean(test_r2_vel)

# %% [code]
np.std(test_r2_vel)

# %% [code]
np.percentile(test_r2_vel, [25, 50, 75])

# %% [code]
np.save('test_r2_vel_simulated.npy', test_r2_vel)

# %% [code]
```
