## Supplementary material for "Dynamic Compression Flows for Neuroscience Data": Python code: paper_plot_musal.ipynb

```
# %% [code]
import numpy as np
import matplotlib.pyplot as plt
from matplotlib.collections import LineCollection
from matplotlib.colors import ListedColormap, BoundaryNorm, LinearSegmentedColormap, Normalize
from mpl_toolkits.mplot3d.art3d import Line3DCollection
import torch
import dnnlib.util_v5 as util
from tqdm import tqdm
import pickle
from matplotlib.ticker import FuncFormatter
import gc
from PIL import Image
from sklearn.cluster import DBSCAN, KMeans
import matplotlib.cm as cm

import numpy as np
import pickle
import os
import re 
import json 
import click
import torch
from torch_utils import distributed as dist
from training import toy_training_loop_vfm_noDataGen
from tqdm import tqdm
import matplotlib.pyplot as plt
from matplotlib import colors
from plot_neural import *
from plot_sim_results_v7 import *
from plot_sim_results_img_v7 import * 
# from plot_musal_v5 import * 
import pickle, types
from torch_utils import persistence

import pandas as pd

import warnings
warnings.filterwarnings('ignore', 'Grad strides do not match bucket view strides') # False warning 
device = torch.device("cuda" if torch.cuda.is_available() else "cpu")
torch.set_default_device(device)
colorgrad = "PuOr"

# %% [code]
def plot_tau(T, out_imgs_tau, dset_samples, trial_idx, tau, title=None):
    # Plot True
    plot_frame(dset_samples, T, trial_idx)
    plt.suptitle(f'true, trial = {trial_idx}')
    plt.subplots_adjust(bottom=0.2)  # Make room for horizontal colorbar

    # Plot Simulated
    plot_frame(out_imgs_tau, T, trial_idx)
    if title is None:
        plt.suptitle(f'simulated- tau = {tau}')
    else:
        plt.suptitle(title)
    plt.subplots_adjust(bottom=0.2)

# %% [code]
def recon_check_x0(encoder, trial_check, T_check, dset_samples):
    encoder = encoder.to(device).eval()
    X1_ori = dset_samples[trial_check][T_check,0,:,:]
    data_flattent = [np.reshape(a, (a.shape[0], -1), order='C') for a in dset_samples]
    with torch.inference_mode():
        X1 = torch.from_numpy(data_flattent[trial_check][T_check,:]).to(device=device, dtype=torch.float32)
        X0 = encoder.rsample(X1)
    X0_back = X0.reshape(X0.shape[0], C, H, W)
    return X0_back[0,0,:,:].detach().cpu().numpy()

# %% [code]
def recon_check_x0_allT(encoder, trial_check, dset_samples):
    encoder = encoder.to(device).eval()
    X1_ori = dset_samples[trial_check][T_check,0,:,:]
    data_flattent = [np.reshape(a, (a.shape[0], -1), order='C') for a in dset_samples]
    with torch.inference_mode():
        X1 = torch.from_numpy(data_flattent[trial_check]).to(device=device, dtype=torch.float32)
        X0 = encoder.rsample(X1)
    X0_back = X0.reshape(X0.shape[0], C, H, W)
    return X0_back.detach().cpu().numpy()

# %% [code]
def plot_mu_velocity_2d(
    mu,
    v_mu,
    trajectories=None,
    dims_2d=(0, 1),
    title=None,
    arrow_frac=0.05,
    arrow_size=1.2,
    linewidth=1.5,
    traj_alpha=0.4,
    traj_width=0.8,
    traj_color="gray",
    max_trajs=300,
    quiver_alpha=0.6,
    max_points=3000,
    figsize=(7, 6),
    speed_color=None,                      # NEW
    speed_norm=None,                       # NEW
    colorbar_label=r"$\|\dot{\mu}\|$",     # NEW
):
    fig, ax = plt.subplots(figsize=figsize, constrained_layout=True)
    ax.grid(False)

    traj_list = []
    if trajectories is not None:
        traj_source = trajectories
        for i, traj in enumerate(traj_source):
            if i >= max_trajs:
                break
            t_np = traj.detach().cpu().numpy() if hasattr(traj, "detach") else np.array(traj)
            traj_list.append(t_np)

    d1, d2 = dims_2d
    for t_np in traj_list:
        ax.plot(
            t_np[:, d1], t_np[:, d2],
            color=traj_color, alpha=traj_alpha, linewidth=traj_width, zorder=1
        )

    coords = mu[:, [d1, d2]]
    vecs   = v_mu[:, [d1, d2]]
    N = coords.shape[0]

    # --- preprocess speed_color (for coloring only) ---
    speed_color_np = None
    if speed_color is not None:
        speed_color_np = speed_color.detach().cpu().numpy() if hasattr(speed_color, "detach") else np.asarray(speed_color)
        speed_color_np = np.asarray(speed_color_np)
        if speed_color_np.ndim == 2:
            speed_color_np = np.linalg.norm(speed_color_np, axis=1)  # (N,)
        speed_color_np = speed_color_np.reshape(-1)
        assert speed_color_np.shape[0] == mu.shape[0]

    if max_points is not None and N > max_points:
        idx = np.random.choice(N, size=max_points, replace=False)
        coords_2d = coords[idx]
        vecs_2d   = vecs[idx]
        speed_c   = speed_color_np[idx] if speed_color_np is not None else None
    else:
        coords_2d = coords
        vecs_2d   = vecs
        speed_c   = speed_color_np if speed_color_np is not None else None

    # --- directions from projected velocity ---
    speed_proj = np.linalg.norm(vecs_2d, axis=1)
    eps = 1e-8
    dirs = vecs_2d / (speed_proj[:, None] + eps)

    x_min, x_max = coords_2d[:, 0].min(), coords_2d[:, 0].max()
    y_min, y_max = coords_2d[:, 1].min(), coords_2d[:, 1].max()
    span = max(x_max - x_min, y_max - y_min) + eps
    L = arrow_frac * span
    U = dirs[:, 0] * L
    V = dirs[:, 1] * L

    base_width = 0.004
    base_headwidth = 4.0
    base_headlength = 6.0
    base_headaxislength = 5.0
    width = base_width * arrow_size

    # --- coloring values ---
    if speed_c is None:
        speed_c = speed_proj

    norm = speed_norm if speed_norm is not None else Normalize(
        vmin=float(speed_c.min()),
        vmax=float(speed_c.max()),
    )

    q = ax.quiver(
        coords_2d[:, 0], coords_2d[:, 1], U, V, speed_c,
        cmap=cm.viridis, norm=norm, angles="xy", scale_units="xy", scale=1.0,
        width=width,
        headwidth=base_headwidth * arrow_size,
        headlength=base_headlength * arrow_size,
        headaxislength=base_headaxislength * arrow_size,
        linewidth=linewidth, alpha=quiver_alpha, zorder=2
    )
    cb = fig.colorbar(q, ax=ax, fraction=0.046, pad=0.04)
    cb.set_label(colorbar_label)

    ax.set_xlabel(fr"$\mu_{{{d1+1}}}$")
    ax.set_ylabel(fr"$\mu_{{{d2+1}}}$")
    if title:
        ax.set_title(title)
    ax.axis("equal")

    return fig, ax, q


def plot_mu_velocity_3d(
    mu,
    v_mu,
    trajectories=None,
    dims_3d=(0, 1, 2),
    title=None,
    arrow_frac=0.1,
    arrow_size=1.5,
    linewidth=1.5,
    base_arrow_length_ratio=0.3,
    traj_alpha=0.4,
    traj_width=0.8,
    quiver_alpha=0.6,
    traj_color="gray",
    max_trajs=300,
    max_points=3000,
    figsize=(10, 7),
    elev=None,
    azim=None,
    roll=None,
    speed_color=None,                      # NEW
    speed_norm=None,                       # NEW
    colorbar_label=r"$\|\dot{\mu}\|$",     # NEW
):
    fig = plt.figure(figsize=figsize, constrained_layout=True)
    ax = fig.add_subplot(1, 1, 1, projection="3d")

    # set camera/view rotation
    if elev is not None or azim is not None or roll is not None:
        ax.view_init(elev=elev, azim=azim, roll=roll)

    ax.set_box_aspect((1, 1, 1))
    ax.grid(False)
    for axis in (ax.xaxis, ax.yaxis, ax.zaxis):
        axis.pane.fill = False
        axis.pane.set_edgecolor((1, 1, 1, 0))
        axis._axinfo["grid"]["color"] = (1, 1, 1, 0)
        axis._axinfo["grid"]["linewidth"] = 0

    traj_list = []
    if trajectories is not None:
        traj_source = trajectories
        for i, traj in enumerate(traj_source):
            if i >= max_trajs:
                break
            t_np = traj.detach().cpu().numpy() if hasattr(traj, "detach") else np.array(traj)
            traj_list.append(t_np)

    d1, d2, d3 = dims_3d
    for t_np in traj_list:
        ax.plot(
            t_np[:, d1], t_np[:, d2], t_np[:, d3],
            color=traj_color, alpha=traj_alpha, linewidth=traj_width, zorder=1
        )

    coords = mu[:, [d1, d2, d3]]
    vecs   = v_mu[:, [d1, d2, d3]]
    N = coords.shape[0]

    # --- preprocess speed_color (for coloring only) ---
    speed_color_np = None
    if speed_color is not None:
        speed_color_np = speed_color.detach().cpu().numpy() if hasattr(speed_color, "detach") else np.asarray(speed_color)
        speed_color_np = np.asarray(speed_color_np)
        if speed_color_np.ndim == 2:
            speed_color_np = np.linalg.norm(speed_color_np, axis=1)  # (N,)
        speed_color_np = speed_color_np.reshape(-1)
        assert speed_color_np.shape[0] == mu.shape[0]

    if max_points is not None and N > max_points:
        idx = np.random.choice(N, size=max_points, replace=False)
        coords_3d = coords[idx]
        vecs_3d   = vecs[idx]
        speed_c   = speed_color_np[idx] if speed_color_np is not None else None
    else:
        coords_3d = coords
        vecs_3d   = vecs
        speed_c   = speed_color_np if speed_color_np is not None else None

    # --- directions from projected velocity ---
    speed_proj = np.linalg.norm(vecs_3d, axis=1)
    eps = 1e-8
    dirs = vecs_3d / (speed_proj[:, None] + eps)

    mins = coords_3d.min(axis=0)
    maxs = coords_3d.max(axis=0)
    span = float(np.max(maxs - mins) + eps)

    L = arrow_frac * span
    U, V, W = dirs[:, 0], dirs[:, 1], dirs[:, 2]

    # --- coloring values ---
    if speed_c is None:
        speed_c = speed_proj

    norm = speed_norm if speed_norm is not None else Normalize(
        vmin=float(speed_c.min()),
        vmax=float(speed_c.max()),
    )

    colors = cm.viridis(norm(speed_c))
    colors[:, 3] = quiver_alpha

    ax.quiver(
        coords_3d[:, 0], coords_3d[:, 1], coords_3d[:, 2],
        U, V, W,
        length=L, normalize=False, colors=colors,
        linewidth=linewidth * arrow_size,
        arrow_length_ratio=base_arrow_length_ratio * arrow_size,
        zorder=2
    )

    mappable = cm.ScalarMappable(norm=norm, cmap="viridis")
    mappable.set_array(speed_c)
    cb = fig.colorbar(mappable, ax=ax, fraction=0.035, pad=0.08)
    cb.set_label(colorbar_label)

    ax.set_xlabel(fr"$\mu_{{{d1+1}}}$")
    ax.set_ylabel(fr"$\mu_{{{d2+1}}}$")
    ax.set_zlabel(fr"$\mu_{{{d3+1}}}$")
    if title:
        ax.set_title(title)

    return fig, ax


    
# def plot_mu_velocity_3d(
#     mu,
#     v_mu,
#     trajectories=None,
#     dims_3d=(0, 1, 2),
#     title=None,
#     arrow_frac=0.1,
#     arrow_size=1.5,
#     linewidth=1.5,
#     base_arrow_length_ratio=0.3,
#     traj_alpha=0.4,
#     traj_width=0.8,
#     quiver_alpha=0.6,
#     traj_color="gray",
#     max_trajs=300,
#     max_points=3000,
#     figsize=(10, 7),
# ):
#     fig = plt.figure(figsize=figsize, constrained_layout=True)
#     ax = fig.add_subplot(1, 1, 1, projection="3d")
#     ax.set_box_aspect((1, 1, 1))
#     ax.grid(False)
#     for axis in (ax.xaxis, ax.yaxis, ax.zaxis):
#         axis.pane.fill = False
#         axis.pane.set_edgecolor((1, 1, 1, 0))
#         axis._axinfo["grid"]["color"] = (1, 1, 1, 0)
#         axis._axinfo["grid"]["linewidth"] = 0

#     traj_list = []
#     if trajectories is not None:
#         traj_source = trajectories
#         for i, traj in enumerate(traj_source):
#             if i >= max_trajs:
#                 break
#             t_np = traj.detach().cpu().numpy() if hasattr(traj, "detach") else np.array(traj)
#             traj_list.append(t_np)

#     d1, d2, d3 = dims_3d
#     for t_np in traj_list:
#         ax.plot(
#             t_np[:, d1], t_np[:, d2], t_np[:, d3],
#             color=traj_color, alpha=traj_alpha, linewidth=traj_width, zorder=1
#         )

#     coords = mu[:, [d1, d2, d3]]
#     vecs   = v_mu[:, [d1, d2, d3]]
#     N = coords.shape[0]

#     if max_points is not None and N > max_points:
#         idx = np.random.choice(N, size=max_points, replace=False)
#         coords_3d = coords[idx]
#         vecs_3d   = vecs[idx]
#     else:
#         coords_3d = coords
#         vecs_3d   = vecs

#     speed = np.linalg.norm(vecs_3d, axis=1)
#     eps = 1e-8
#     dirs = vecs_3d / (speed[:, None] + eps)

#     mins = coords_3d.min(axis=0)
#     maxs = coords_3d.max(axis=0)
#     span = float(np.max(maxs - mins) + eps)

#     L = arrow_frac * span
#     U, V, W = dirs[:, 0], dirs[:, 1], dirs[:, 2]

#     norm = Normalize(vmin=float(speed.min()), vmax=float(speed.max()))
#     colors = cm.viridis(norm(speed))
#     colors[:, 3] = quiver_alpha

#     ax.quiver(
#         coords_3d[:, 0], coords_3d[:, 1], coords_3d[:, 2],
#         U, V, W,
#         length=L, normalize=False, colors=colors,
#         linewidth=linewidth * arrow_size,
#         arrow_length_ratio=base_arrow_length_ratio * arrow_size,
#         zorder=2
#     )

#     mappable = cm.ScalarMappable(norm=norm, cmap="viridis")
#     mappable.set_array(speed)
#     cb = fig.colorbar(mappable, ax=ax, fraction=0.035, pad=0.08)
#     cb.set_label(r"$\|\dot{\mu}\|$")

#     ax.set_xlabel(fr"$\mu_{{{d1+1}}}$")
#     ax.set_ylabel(fr"$\mu_{{{d2+1}}}$")
#     ax.set_zlabel(fr"$\mu_{{{d3+1}}}$")
#     if title:
#         ax.set_title(title)

#     # fig.tight_layout()
#     return fig, ax

# %% [markdown]
# 1. Read Data

# %% [code]
folder_data = 'out/musal_behavior_local/dimk_lag0/t/none/data'
dataset_samples = np.load(folder_data + "/dataset_samples.npz", allow_pickle=True)
cov_static_samples = np.load(folder_data + "/cov_static_trunc_only.npz", allow_pickle=True)

dset_samples = to_list_of_arrays(dataset_samples['samples'])
cov_stat_samples = to_list_of_arrays(cov_static_samples['samples'])
mean_img = np.load('data/musal_behavior/mean_image.npy')

# %% [code]
len(dset_samples)

# %% [code]
dset_samples[0].shape

# %% [code]
idx = 0
plt.imshow(dset_samples[0][idx,0] + mean_img, cmap = 'gray')
plt.title('trial 0, T = ' + str(idx))

# %% [code]
eps = 1e-2
T_max = max(a.shape[0] for a in dset_samples)
padded = [np.pad(a, ((0, T_max-a.shape[0]), (0,0), (0,0), (0,0)), mode='constant', constant_values=np.nan)
          for a in dset_samples]                                              # -> (n_trial, T_max, C, H, W)
stacked = np.stack(padded, axis=0)
min_img = np.nanmin(stacked, axis=(0,1))                                      # shape (C,H,W)
max_img = np.nanmax(stacked, axis=(0,1))                                      # shape (C,H,W)
clamp_range = np.stack([min_img - eps, max_img + eps], axis=0).astype(np.float32)  # shape (2,C,H,W)

# %% [code]
is_image = (np.asarray(dset_samples[0]).ndim == 4)
if is_image:
    _, C, H, W = np.asarray(dset_samples[0]).shape
T_min = min(arr.shape[0] for arr in dset_samples)
T_max = max(arr.shape[0] for arr in dset_samples)
n_step = T_max//50
dset_samples_look = [dset_samples[0][:n_step]]
del dset_samples

# %% [markdown]
# 2. Read Model

# %% [code]
folder = 'out/musal_behavior_local/dimk_lag0/t/none/00000-musal_dim50_lag0_none-prr-uncond-regularFM-gpus1-batch256-fp32'
chk_pts = '/network-snapshot-020823.pkl'
useCov = False
dim_to_keep = 3

pkl_path = folder + chk_pts

# %% [code]
if not hasattr(persistence, "_orig_reconstruct"):
    persistence._orig_reconstruct = persistence._reconstruct_persistent_obj
def _tolerant_reconstruct(meta):
    try:
        return persistence._orig_reconstruct(meta)
    except Exception:
        class _Stub: pass
        return _Stub()

class PatchedUnpickler(pickle.Unpickler):
    def find_class(self, module, name):
        if module == "torch_utils.persistence" and name == "_reconstruct_persistent_obj":
            return _tolerant_reconstruct
        return super().find_class(module, name)

with open(pkl_path, "rb") as f:
    model_all = PatchedUnpickler(f).load()

# use the net; ignore any loss_fn in the pickle
net = model_all["ema"] if "ema" in model_all else model_all["net"]
flow_net = net.unet_model
dyn_net   = net.vnet_model
encoder   = net.encoder

# %% [code]
lag = infer_lag_k_from_dyn_net(dyn_net)
print(lag)

# %% [markdown]
# 3. Check Encoder

# %% [code]
trial_check = 0
T_check = 0
X0_back = recon_check_x0(encoder, trial_check, T_check, dset_samples = dset_samples_look)

# %% [code]
fig, axs = plt.subplots(1, 2, figsize=(6, 5))

im1 = axs[0].imshow(X0_back)
fig.colorbar(im1, ax=axs[0], fraction=0.046, pad=0.04)  # smaller, aligned
axs[0].set_title('encoder recon')

im2 = axs[1].imshow(dset_samples_look[trial_check][T_check,0,:,:])
fig.colorbar(im2, ax=axs[1], fraction=0.046, pad=0.04)  # same settings
axs[1].set_title('training data')

plt.tight_layout()
plt.show()

# %% [code]
data_flattent = [np.reshape(a, (a.shape[0], -1), order='C') for a in dset_samples_look]
dim_to_keep = 3
mu_keep = get_mu_from_encoder(encoder, data_flattent, dim_to_keep, 0)
plot_latent_time_series([mu_keep], alpha = 0.4, dims=np.arange(dim_to_keep), 
                        dt=0.03333333333333333, labels=None)

# %% [code]
plot_traj_3d(mu_keep)

# %% [code]
mu_encoder, d, L, loading, bias = latent_mu(dset_samples_look, encoder.to(device), dim_to_keep, device, fullRes = True)
L_np = L.detach().cpu().numpy()
d_np = d.detach().cpu().numpy()
loading_np = loading.detach().cpu().numpy()
mu_all = get_mu_from_encoder(encoder, data_flattent, 64*64, 0)

# %% [code]
k_max = 200
k_target = 50
loading_norm_all = torch.norm(loading[:,0:k_max], dim= 0).detach().cpu().numpy()
energy = loading_norm_all ** 2 
cum_energy = np.cumsum(energy)
K_eff = 1 + np.searchsorted(cum_energy, 0.95 * cum_energy[-1])

# %% [code]
fig = plt.figure(figsize=(3, 3))
plt.plot(np.sqrt(d_np[0:k_max]))
plt.axvline(x=k_target-1, color='red', label = 'k_target')
plt.axvline(x=K_eff-1, color='orange', label = 'k_eff = ' + str(K_eff))
plt.title('sqrt{d}');
plt.legend()

# %% [code]
k_max = 100
dim_check_most = 100
k_target = 50
rows = loading.shape[0]
cols = dim_check_most
dim_check = np.arange(dim_check_most)
# rec = mu_group_recon(bias.detach().cpu().numpy(), loading_np, mu_all, dim_check, H=28, W=28)
# ball_idx = 0

fig, axs = plt.subplots(2, 2, figsize=(6, 5))
axs[0, 0].plot(np.sqrt(d_np[0:dim_check_most]))
axs[0, 0].axvline(x=k_target-1, color='red', label = 'k_target')
axs[0, 0].axvline(x=K_eff-1, color='orange', label = 'k_eff = ' + str(K_eff))
axs[0, 0].set_title('sqrt{d}');
axs[0, 0].legend()

im1 = axs[0, 1].imshow(np.abs(loading_np[:, 0:dim_check_most]), cmap='gray_r', aspect=cols / rows)
axs[0, 1].set_title('|loading|')
fig.colorbar(im1, ax=axs[0, 1])

axs[1, 0].plot(mu_all[:, :dim_check_most])
axs[1, 0].set_title('first ' + str(dim_check_most) + " dim of mu")

# # Plot 4: cumulative contribution
# im2 = axs[1, 1].imshow(rec[ball_idx], cmap='gray')
# axs[1, 1].set_title('cumulative contribution from ' + str(dim_check[0]) + ' to ' + str(dim_check[-1]))
# fig.colorbar(im2, ax=axs[1, 1])

fig.suptitle('Encoder Diagnostic', fontsize=16)
plt.tight_layout()
plt.show()

# %% [markdown]
# 3. Flowed Latent

# %% [code]
gc.collect()
torch.cuda.empty_cache()
dim_to_keep = 3
latent_traj_list = proj_to_latent(dset_samples_look, flow_net.to(device), 0.0, device)

# %% [code]
latent_traj_list_enc = []
for ii in range(len(dset_samples_look)):
    latent_traj_list_enc.append(recon_check_x0_allT(encoder, ii, dset_samples_look))

# %% [code]
mu_traj_sim, d, L, loading, bias = latent_mu(latent_traj_list, encoder.to(device), dim_to_keep, device,
                                             fullRes = True, resid_tol=1e-15)

# %% [code]
data_flow = [arr.reshape(arr.shape[0], C, H, W) for arr in latent_traj_list ]
data_enc = [arr.reshape(arr.shape[0], C, H, W) for arr in latent_traj_list_enc ]

trial_check = 0
T_check = 0
fig, axs = plt.subplots(1, 3, figsize=(12, 10))

im1 = axs[0].imshow(data_flow[trial_check][T_check,0])
fig.colorbar(im1, ax=axs[0], fraction=0.046, pad=0.04)  # smaller, aligned
axs[0].set_title('flow(X)')

im2 = axs[1].imshow(data_enc[trial_check][T_check,0])
fig.colorbar(im2, ax=axs[1], fraction=0.046, pad=0.04)  # smaller, aligned
axs[1].set_title('encoder(X)')

im3 = axs[2].imshow(data_flow[trial_check][T_check,0] - data_enc[trial_check][T_check,0])
fig.colorbar(im3, ax=axs[2], fraction=0.046, pad=0.04)  # smaller, aligned
axs[2].set_title('flow(X) - encoder(X)')

plt.tight_layout()
plt.show()

# %% [code]
mu_traj_flow_np = [mu_traj_sim[j].cpu().numpy() for j in range(len(mu_traj_sim))]
plot_latent_time_series(mu_traj_flow_np, alpha = 0.4, dims=np.arange(dim_to_keep), 
                        dt=0.033333333, labels=None)
plt.suptitle('flowed latent trajc for each trial')

# %% [code]
plot_traj_3d(mu_traj_flow_np[0][:,0:3])

# %% [markdown]
# 4. Simulated Traj

# %% [markdown]
The simulation doesn't make any sense. Just show for fun.

# %% [code]
tau = 1.0
traj_sim_tau1_flattent, lag_cov_list = generate_traj_image_onthefly(dyn_net.to(device),
                                                                    dset_samples_look, n_step,
                                                                    tau, device, lag,
                                                                    include_x0_tau=False,
                                                                    oracle=False,
                                                                    flow_net=None,
                                                                    clamp_range=clamp_range
                                                                    )

# %% [code]
out_imgs_tau1 = [arr.reshape(arr.shape[0], C, H, W) for arr in traj_sim_tau1_flattent]

# %% [code]
plot_tau(n_step, out_imgs_tau1, dset_samples_look, trial_idx = 0, tau = 1.0)

# %% [code]
out_imgs_tau0_flow_flatten = proj_to_latent(out_imgs_tau1, flow_net.to(device), 0.0, device)
out_imgs_tau0_flow = [arr.reshape(arr.shape[0], C, H, W) for arr in out_imgs_tau0_flow_flatten ]

# %% [code]
tau = 0.0
traj_sim_tau0_flattent, _ = generate_traj_image_onthefly(dyn_net.to(device),
                                                         dset_samples_look, n_step,
                                                         tau, device, lag,
                                                         include_x0_tau=False,
                                                         oracle=False,
                                                         flow_net=flow_net.to(device),
                                                         clamp_range=clamp_range
                                                         )
out_imgs_tau0 = [arr.reshape(arr.shape[0], C, H, W) for arr in traj_sim_tau0_flattent ]

# %% [code]
plot_tau(n_step, out_imgs_tau0_flow, dset_samples_look,
         trial_idx = 0, tau = 0.0, title = 'porjected, tau = 0.0; flow from tau = 1.0 directly')

# %% [code]
plot_tau(n_step, out_imgs_tau0, dset_samples_look,
         trial_idx = 0, tau = 0.0, title = 'simulated, tau = 0.0')

# %% [code]
dim_to_keep = 3
mu_traj_sim, d, L, loading, bias = latent_mu(traj_sim_tau0_flattent, encoder.to(device), dim_to_keep, device, fullRes = True)

# %% [code]
mu_traj_sim_np = [mu_traj_sim[j].cpu().numpy() for j in range(len(mu_traj_sim))]
mu_traj_sim_np_plot = [mu_traj_sim_np[ii][10:,:] for ii in range(len(mu_traj_sim_np))]
plot_latent_time_series(mu_traj_sim_np_plot, alpha = 0.4, dims=np.arange(dim_to_keep), 
                        dt=0.01, labels=None)
plt.suptitle('simulated latent trajc for each trial')

# %% [code]
plot_traj_3d(mu_traj_sim_np_plot[0][:,0:3])

# %% [markdown]
# 5. Flow Field

# %% [markdown]
## 5.1 from Encoder

# %% [markdown]
Workflow:

1. use encoder to project data $X$ ($\tau = 1.0$) to latent space $Z$ ($\tau = 0.0$) 


# %% [code]
latent_traj_list_enc = []
for ii in range(len(dset_samples_look)):
    latent_traj_list_enc.append(recon_check_x0_allT(encoder, ii, dset_samples_look))
mu_traj_sim_enc, d, L, loading, bias = latent_mu(latent_traj_list_enc, encoder.to(device), dim_to_keep, device,
                                             fullRes = True, resid_tol=1e-15)

# %% [code]
dim_to_keep = 3
mu_pts_encoder, v_mu_encoder, v_mu_full_encoder = compute_mu_velocity_field(
    latent_traj_list=latent_traj_list_enc,
    dyn_net=dyn_net,
    encoder=encoder,
    device=device,
    dim_preserve=dim_to_keep,
    lag=lag,
    t_plot=0.0,
    return_full=True
)

# %% [code]
plot_mu_velocity_panel(mu_pts_encoder, v_mu_encoder,
                       trajectories=mu_traj_sim_enc,
                       title_2d = "encoder: mu1–mu2 field (tau =0.0)",
                       title_3d = "encoder: mu1–mu2-mu3 field (tau =0.0)",
                       speed_color=v_mu_full_encoder,
                       arrow_frac_2d=0.05,
                       arrow_size_2d=1.2,
                       linewidth_2d=1.5,
                       
                       arrow_frac_3d=0.1,
                       arrow_size_3d=1.5,
                       linewidth_3d=1.5,
                       base_arrow_length_ratio_3d=0.3,
                       )

# %% [markdown]
## 5.2 from compression flow

# %% [code]
latent_traj_list = proj_to_latent(dset_samples_look, flow_net.to(device), 0.0, device)
mu_traj_sim, d, L, loading, bias = latent_mu(latent_traj_list, encoder.to(device), dim_to_keep, device,
                                             fullRes = True, resid_tol=1e-15)

# %% [code]
dim_to_keep = 3  # μ-dims you kept elsewhere
mu_pts_flow, v_mu_flow, v_mu_full_flow = compute_mu_velocity_field(
    latent_traj_list=latent_traj_list,
    dyn_net=dyn_net,
    encoder=encoder,
    device=device,
    dim_preserve=dim_to_keep,
    lag=lag,
    t_plot=0.0,
    return_full=True
)

plot_mu_velocity_panel(mu_pts_flow, v_mu_flow,
                       trajectories=[mu_pts_flow], # mu_traj_sim
                       speed_color=v_mu_full_flow,
                       title_2d = "flowed: mu1–mu2 field (tau =0.0)",
                       title_3d = "flowed: mu1–mu2-mu3 field (tau =0.0)",
                       arrow_frac_2d=0.05,
                       arrow_size_2d=1.2,
                       linewidth_2d=1.5,
                       
                       arrow_frac_3d=0.1,
                       arrow_size_3d=1.5,
                       linewidth_3d=1.5,
                       base_arrow_length_ratio_3d=0.3,
                       )

# %% [markdown]
# 6. Let's investigate more on flow field

# %% [markdown]
## 6.1 select points

# %% [code]
mu = mu_pts_flow
v = v_mu_flow
# speed = np.linalg.norm(v, axis=1)
speed = v_mu_full_flow
mu_2d = mu[:, :2]

# Step A: Outlier Detection (DBSCAN)
db = DBSCAN(eps=0.15, min_samples=10).fit(mu_2d)
is_outlier = (db.labels_ == -1)

non_outlier_indices = np.where(~is_outlier)[0]
outlier_indices = np.where(is_outlier)[0]

# Step B: Force 4 Bands on Dense Data (KMeans)
kmeans = KMeans(n_clusters=4, random_state=42, n_init=20)
band_labels_local = kmeans.fit_predict(mu_2d[non_outlier_indices])

final_labels = np.full(mu.shape[0], -1)
final_labels[non_outlier_indices] = band_labels_local

# %% [code]
# select representitive pts for each band
valid_z = mu[non_outlier_indices, 2]
z_top_target = np.percentile(valid_z, 98)
z_low_target = np.percentile(valid_z, 2)
z_mid_target = (z_top_target + z_low_target) / 2.0

bands = []
for label in range(4):
    indices = np.where(final_labels == label)[0]
    band_z = mu[indices, 2]
    
    idx_top = indices[np.argmin(np.abs(band_z - z_top_target))]
    idx_mid = indices[np.argmin(np.abs(band_z - z_mid_target))]
    idx_low = indices[np.argmin(np.abs(band_z - z_low_target))]
    
    bands.append({
        'label': label,
        'indices': indices,
        'reps': [idx_top, idx_mid, idx_low],
        'rep_names': ['Top', 'Mid', 'Low'],
        'avg_x': np.mean(mu[indices, 0])
    })

bands.sort(key=lambda b: b['avg_x'])

# %% [code]
# select representitive pts for outliers
# center2 = mu_2d.mean(axis=0)
# dists = np.linalg.norm(mu_2d[outlier_indices] - center2, axis=1)

center = mu.mean(axis=0)
dists = np.linalg.norm(mu[outlier_indices] - center, axis=1)


k_outliers = 3

# Row 1: Max Distance from Center
dist_sorted_args = np.argsort(dists)[::-1][:k_outliers]
outliers_dist = outlier_indices[dist_sorted_args]
dists_val = dists[dist_sorted_args] # Store distances for titles

# Row 2: Max Velocity
vel_sorted_args = np.argsort(speed[outlier_indices])[::-1][:k_outliers]
outliers_vel = outlier_indices[vel_sorted_args]

# %% [markdown]
## 6.2 Plots

# %% [code]
cmap_bands = plt.get_cmap("tab10")
band_colors = [cmap_bands(i) for i in range(len(bands))]
outlier_color = 'grey'
outlier_maxD_color = 'tab:brown'
outlier_maxV_color = 'tab:cyan'

# %% [markdown]
### 6.2.1 2D plots

# %% [code]
fig1, ax2d = plt.subplots(figsize=(6, 6))
ax2d.scatter(mu[outlier_indices, 0], mu[outlier_indices, 1], 
             c=outlier_color, marker='.', s=10, alpha=0.3, label='Outliers (All)')
ax2d.scatter(mu[outliers_dist, 0], mu[outliers_dist, 1], 
             c=outlier_maxD_color, marker='x', s=80, linewidth=2, label='Max Dist Reps', zorder=10)
ax2d.scatter(mu[outliers_vel, 0], mu[outliers_vel, 1], 
             c=outlier_maxV_color, marker='x', s=80, linewidth=2, label='Max Vel Reps', zorder=10)

for i, band in enumerate(bands):
    idx = band['indices']
    reps = band['reps']
    c = band_colors[i]
    ax2d.scatter(mu[idx, 0], mu[idx, 1], c=[c], s=15, alpha=0.4, label=f'Band {i+1}')
    ax2d.scatter(mu[reps, 0], mu[reps, 1], c=[c], s=100, edgecolors='k', linewidth=1.5, zorder=5)

ax2d.set_title("2D Projection (X vs Y)")
ax2d.set_xlabel("$\mu_1$")
ax2d.set_ylabel("$\mu_2$")
ax2d.legend()
plt.tight_layout()
plt.show()

# %% [markdown]
### 6.2.2 3D plot

# %% [code]
fig2 = plt.figure(figsize=(6, 6))
ax3d = fig2.add_subplot(111, projection='3d')

ax3d.scatter(mu[outlier_indices, 0], mu[outlier_indices, 1], mu[outlier_indices, 2], 
             c=outlier_color, s=10, alpha=0.2)

ax3d.scatter(mu[outliers_dist, 0], mu[outliers_dist, 1], mu[outliers_dist, 2], 
             c=outlier_maxD_color, marker='x', s=100, linewidth=2, zorder=10)
ax3d.scatter(mu[outliers_vel, 0], mu[outliers_vel, 1], mu[outliers_vel, 2], 
             c=outlier_maxV_color, marker='x', s=100, linewidth=2, zorder=10)

for i, band in enumerate(bands):
    idx = band['indices']
    reps = band['reps']
    c = band_colors[i]
    ax3d.scatter(mu[idx, 0], mu[idx, 1], mu[idx, 2], c=[c], s=15, alpha=0.2)
    ax3d.scatter(mu[reps, 0], mu[reps, 1], mu[reps, 2], c=[c], s=150, edgecolors='k', linewidth=2, zorder=10)
    # ax3d.text(mu[reps[0],0], mu[reps[0],1], mu[reps[0],2], f"B{i+1}", fontsize=10, weight='bold', color='k')

ax3d.set_title("3D Flow Structure")
ax3d.set_xlabel("$\mu_1$"); ax3d.set_ylabel("$\mu_2$"); ax3d.set_zlabel("$\mu_3$")
ax3d.view_init(elev=20, azim=-60)
plt.tight_layout()
plt.show()

# %% [markdown]
### 6.2.3 outlier images

# %% [code]
print(len(dset_samples_look))
print(dset_samples_look[0].shape)

# %% [code]
np.mean(dset_samples_look[0][:,0], axis =0).shape

# %% [code]
alpha_diff = 0.5

data_mean = np.mean(dset_samples_look[0][:,0], axis =0)
x_all = dset_samples_look[0][:, 0]   # (N,64,64), centered
if hasattr(x_all, "detach"): x_all = x_all.detach().cpu().numpy()

diff_max = (x_all - data_mean).max() + 1e-12
diff_min = (x_all - data_mean).min() - 1e-12

# %% [code]
fig3, axes3 = plt.subplots(2, k_outliers, figsize=(10, 5))

# Row 1: Max Distance
for i, idx in enumerate(outliers_dist):
    ax = axes3[0, i]
    img_data = dset_samples_look[0][idx, 0] + mean_img
    if hasattr(img_data, 'detach'): img_data = img_data.detach().cpu().numpy()
    
    ax.imshow(img_data, cmap='gray', interpolation="nearest", rasterized=True)
    # diff = np.abs(dset_samples_look[0][idx, 0])
    diff = dset_samples_look[0][idx, 0] - data_mean
    ax.imshow(diff, cmap="seismic", alpha=alpha_diff,
              vmin=diff_min, vmax=diff_max, interpolation="nearest", rasterized=True)

    
    # Title with Distance
    dist_val = dists_val[i]
    # ax.set_title(f"$|v|={speed[idx]:.2f}$\n$Dist={dist_val:.2f}$", fontsize=9)
    
    # Grey dashed border for distance outliers matches scatter marker color (black/grey)
    for spine in ax.spines.values():
        spine.set_edgecolor(outlier_maxD_color); spine.set_linewidth(3); 
        # spine.set_linestyle('--')
    ax.set_xticks([]); ax.set_yticks([])
    if i == 0: ax.set_ylabel("Max Distance", fontsize=12, fontweight='bold')

# Row 2: Max Velocity
for i, idx in enumerate(outliers_vel):
    ax = axes3[1, i]
    img_data = dset_samples_look[0][idx, 0] + mean_img
    if hasattr(img_data, 'detach'): img_data = img_data.detach().cpu().numpy()
    
    ax.imshow(img_data, cmap='gray', interpolation="nearest", rasterized=True)
    # ax.set_title(f"$|v|={speed[idx]:.2f}$", fontsize=9)
    # diff = np.abs(img_data - mean_img)
    # ax.imshow(diff, cmap="Reds", alpha=alpha_diff, vmin=0.0, vmax=diff_max)
    diff = dset_samples_look[0][idx, 0] - data_mean
    ax.imshow(diff, cmap="seismic", alpha=alpha_diff,
              vmin=diff_min, vmax=diff_max, interpolation="nearest", rasterized=True)
    
    # Red border for velocity outliers
    for spine in ax.spines.values():
        spine.set_edgecolor(outlier_maxV_color); spine.set_linewidth(3)
    ax.set_xticks([]); ax.set_yticks([])
    if i == 0: ax.set_ylabel("Max Velocity", fontsize=12, fontweight='bold')

fig3.suptitle("Outlier Analysis", fontsize=16)
plt.tight_layout()
plt.show()

# %% [markdown]
### 6.2.4 band image matrix (representitive pts)

# %% [code]
fig4, axes4 = plt.subplots(3, 4, figsize=(10, 6))
positions = ['Top', 'Mid', 'Low']

for col_idx, band in enumerate(bands):
    c = band_colors[col_idx]
    reps = band['reps']
    for row_idx, idx in enumerate(reps):
        ax = axes4[row_idx, col_idx]
        
        img_data = dset_samples_look[0][idx, 0] + mean_img
        if hasattr(img_data, 'detach'): img_data = img_data.detach().cpu().numpy()
        
        ax.imshow(img_data, cmap='gray')
        # diff = np.abs(dset_samples_look[0][idx, 0])
        # ax.imshow(diff, cmap="Reds", alpha=alpha_diff, vmin=0.0, vmax=diff_max)
        # ax.set_title(f"$|v|={speed[idx]:.2f}, z={mu[idx,2]:.1f}$", fontsize=10)
        diff = dset_samples_look[0][idx, 0] - data_mean
        ax.imshow(diff, cmap="seismic", alpha=alpha_diff, vmin=diff_min, vmax=diff_max)
        
        for spine in ax.spines.values():
            spine.set_edgecolor(c); spine.set_linewidth(4)
        ax.set_xticks([]); ax.set_yticks([])
        
        if row_idx == 0:
            ax.set_xlabel(f"Band {col_idx+1}", fontsize=14, fontweight='bold', color=c)
            ax.xaxis.set_label_position('top') 
        if col_idx == 0:
            ax.set_ylabel(positions[row_idx], fontsize=14, fontweight='bold')

fig4.suptitle("Band Evolution (Top to Low)", fontsize=16)
plt.tight_layout()
plt.show()

# %% [markdown]
### 6.2.5 band image (mean/ deviation from mean/ variability)

# %% [code]
fig5, axes5 = plt.subplots(3, 4, figsize=(12, 8))

imgs_all = []
for idx in non_outlier_indices:
    img = dset_samples_look[0][idx, 0] + mean_img
    if hasattr(img, 'detach'):
        img = img.detach().cpu().numpy()
    imgs_all.append(img)
imgs_all = np.asarray(imgs_all, dtype=np.float32)
global_mean_img = imgs_all.mean(axis=0)

band_mean_imgs = []
band_std_imgs = []
for band in bands:
    imgs = []
    for idx in band['indices']:
        img = dset_samples_look[0][idx, 0] + mean_img
        if hasattr(img, 'detach'):
            img = img.detach().cpu().numpy()
        imgs.append(img)
    imgs = np.asarray(imgs, dtype=np.float32)
    band_mean_imgs.append(imgs.mean(axis=0))
    band_std_imgs.append(imgs.std(axis=0))

diff_imgs = [m - global_mean_img for m in band_mean_imgs]
max_abs_diff = max(float(np.max(np.abs(d))) for d in diff_imgs)
max_std = max(float(np.max(s)) for s in band_std_imgs)

for col_idx, band in enumerate(bands):
    c = band_colors[col_idx]
    idxs = band['indices']
    n = len(idxs)
    mean_speed = float(np.mean(speed[idxs])) if n > 0 else 0.0

    ax = axes5[0, col_idx]
    ax.imshow(band_mean_imgs[col_idx], cmap='gray')
    
    ax.set_title(f"n={n}, mean|v|={mean_speed:.2f}", fontsize=10)
    ax.set_xticks([]); ax.set_yticks([])
    ax.set_xlabel(f"Band {col_idx+1}", fontsize=14, fontweight='bold', color=c)
    ax.xaxis.set_label_position('top')

    ax = axes5[1, col_idx]
    ax.imshow(diff_imgs[col_idx], cmap='seismic', vmin=-max_abs_diff, vmax=max_abs_diff)
    ax.set_xticks([]); ax.set_yticks([])

    ax = axes5[2, col_idx]
    ax.imshow(band_std_imgs[col_idx], cmap='gray', vmin=0.0, vmax=max_std)
    ax.set_xticks([]); ax.set_yticks([])

    # Colored borders (match Fig3 style)
    for r in range(3):
        for spine in axes5[r, col_idx].spines.values():
            spine.set_edgecolor(c)
            spine.set_linewidth(4)

# Row labels
axes5[0, 0].set_ylabel("Mean", fontsize=14, fontweight='bold')
axes5[1, 0].set_ylabel("Mean-Global", fontsize=14, fontweight='bold')
axes5[2, 0].set_ylabel("Std", fontsize=14, fontweight='bold')

fig5.suptitle("Band Fingerprints", fontsize=16)
plt.tight_layout(rect=[0, 0.20, 1, 0.95])

sm_diff = mpl.cm.ScalarMappable(
    norm=mpl.colors.Normalize(vmin=-max_abs_diff, vmax=max_abs_diff),
    cmap='seismic'
)
sm_diff.set_array([])

sm_std = mpl.cm.ScalarMappable(
    norm=mpl.colors.Normalize(vmin=0.0, vmax=max_std),
    cmap='gray'
)
sm_std.set_array([])

# side-by-side colorbars
cbax_diff = fig5.add_axes([0.12, 0.08, 0.36, 0.03])  # left bar
cbar_diff = fig5.colorbar(sm_diff, cax=cbax_diff, orientation='horizontal')
cbar_diff.set_label("Mean - Global", fontsize=11)

cbax_std = fig5.add_axes([0.56, 0.08, 0.36, 0.03])   # right bar
cbar_std = fig5.colorbar(sm_std, cax=cbax_std, orientation='horizontal')
cbar_std.set_label("Std", fontsize=11)

plt.show()

# %% [markdown]
# Plot

# %% [code]
plotFolder = "paper_fig/musal"

# %% [code]
# 1. sqrt{d}
k_max = 100
fig = plt.figure(figsize=(4, 4))
plt.plot(np.sqrt(d_np[0:k_max]))
plt.axvline(x=k_target-1, color='red', label = 'k_target')
plt.axvline(x=K_eff-1, color='orange', label = 'k_eff = ' + str(K_eff))
plt.title('sqrt{d}');
plt.legend()

plt.savefig(plotFolder + "/1_sqrt_d.pdf")
plt.savefig(plotFolder + "/1_sqrt_d.svg")
plt.savefig(plotFolder + "/1_sqrt_d.png")

# %% [code]
# 2.1 compressed flow field in mu (2D)
plot_mu_velocity_2d(mu_pts_flow, v_mu_flow, trajectories=[mu_pts_flow], dims_2d=(0,1),
                    speed_color= v_mu_full_flow,
                    title="flowed: mu1–mu2 field (tau =0.0)", figsize=(5, 4))

plt.savefig(plotFolder + "/2_1_flow_v_2d.pdf")
plt.savefig(plotFolder + "/2_1_flow_v_2d.svg")
plt.savefig(plotFolder + "/2_1_flow_v_2d.png")

# %% [code]
elev = 45
azim = -60

# %% [code]
# 2.2 compressed flow field in mu (3D)
fig_flow, ax_flow = plot_mu_velocity_3d(
    mu_pts_flow, v_mu_flow, trajectories=[mu_pts_flow], dims_3d=(0,1,2),
    speed_color=v_mu_full_flow,
    title="flowed: mu1–mu2-mu3 field (tau =0.0)", figsize=(5, 5), 
    arrow_frac=0.08, arrow_size=1.4,
    elev=elev, azim=azim)

xlim_ref, ylim_ref, zlim_ref = ax_flow.get_xlim3d(), ax_flow.get_ylim3d(), ax_flow.get_zlim3d()

plt.savefig(plotFolder + "/2_2_flow_v_3d.pdf")
plt.savefig(plotFolder + "/2_2_flow_v_3d.svg")
plt.savefig(plotFolder + "/2_2_flow_v_3d.png")

# %% [code]
# 3.1 scatter mu (2D)
fig1, ax2d = plt.subplots(figsize=(4.5, 4.5))
ax2d.scatter(mu[outlier_indices, 0], mu[outlier_indices, 1], 
             c=outlier_color, marker='.', s=10, alpha=0.3, label='Outliers (All)')
ax2d.scatter(mu[outliers_dist, 0], mu[outliers_dist, 1], 
             c=outlier_maxD_color, marker='x', s=80, linewidth=2, label='Max Dist Reps', zorder=10)
ax2d.scatter(mu[outliers_vel, 0], mu[outliers_vel, 1], 
             c=outlier_maxV_color, marker='x', s=80, linewidth=2, label='Max Vel Reps', zorder=10)

for i, band in enumerate(bands):
    idx = band['indices']
    reps = band['reps']
    c = band_colors[i]
    ax2d.scatter(mu[idx, 0], mu[idx, 1], c=[c], s=15, alpha=0.4, label=f'Band {i+1}')
    ax2d.scatter(mu[reps, 0], mu[reps, 1], c=[c], s=100, edgecolors='k', linewidth=1.5, zorder=5)

ax2d.set_title("2D Projection (X vs Y)")
ax2d.set_xlabel("$\mu_1$")
ax2d.set_ylabel("$\mu_2$")
ax2d.legend()
plt.tight_layout()

plt.savefig(plotFolder + "/3_1_mu_2d.pdf")
plt.savefig(plotFolder + "/3_1_mu_2d.svg")
plt.savefig(plotFolder + "/3_1_mu_2d.png")

# %% [code]
# 3.2 scatter mu (3D)
roll = None
fig2 = plt.figure(figsize=(4.5, 4.5))
ax3d = fig2.add_subplot(111, projection='3d')

ax3d.grid(False)
for axis in (ax3d.xaxis, ax3d.yaxis, ax3d.zaxis):
    axis.pane.fill = False
    axis.pane.set_edgecolor((1, 1, 1, 0))
    axis._axinfo["grid"]["color"] = (1, 1, 1, 0)
    axis._axinfo["grid"]["linewidth"] = 0

ax3d.scatter(mu[outlier_indices, 0], mu[outlier_indices, 1], mu[outlier_indices, 2],
             c=outlier_color, s=10, alpha=0.2)

ax3d.scatter(mu[outliers_dist, 0], mu[outliers_dist, 1], mu[outliers_dist, 2],
             c=outlier_maxD_color, marker='x', s=100, linewidth=2, zorder=10)
ax3d.scatter(mu[outliers_vel, 0], mu[outliers_vel, 1], mu[outliers_vel, 2],
             c=outlier_maxV_color, marker='x', s=100, linewidth=2, zorder=10)

for i, band in enumerate(bands):
    idx = band['indices']
    reps = band['reps']
    c = band_colors[i]
    ax3d.scatter(mu[idx, 0], mu[idx, 1], mu[idx, 2], c=[c], s=15, alpha=0.2)
    ax3d.scatter(mu[reps, 0], mu[reps, 1], mu[reps, 2], c=[c], s=150, edgecolors='k', linewidth=2, zorder=10)

ax3d.set_title("3D Flow Structure")
ax3d.set_xlabel("$\mu_1$"); ax3d.set_ylabel("$\mu_2$"); ax3d.set_zlabel("$\mu_3$")
ax3d.view_init(elev=elev, azim=azim)

ax3d.set_xlim3d(xlim_ref)
ax3d.set_ylim3d(ylim_ref)
ax3d.set_zlim3d(zlim_ref)

plt.tight_layout()
plt.savefig(plotFolder + "/3_2_mu_3d.pdf")
plt.savefig(plotFolder + "/3_2_mu_3d.svg")
plt.savefig(plotFolder + "/3_2_mu_3d.png")

# %% [code]
from matplotlib.colors import TwoSlopeNorm

diff_norm = TwoSlopeNorm(vmin=diff_min, vcenter=0.0, vmax=diff_max)

# %% [code]
alpha_diff = 0.4

# %% [code]
# # 4. outliers
# fig3, axes3 = plt.subplots(2, k_outliers, figsize=(6, 4))

# for i, idx in enumerate(outliers_dist):
#     ax = axes3[0, i]
#     img_data = dset_samples_look[0][idx, 0] + mean_img
#     if hasattr(img_data, 'detach'): img_data = img_data.detach().cpu().numpy()
    
#     ax.imshow(img_data, cmap='gray', interpolation="nearest", rasterized=True)
#     diff = dset_samples_look[0][idx, 0] - data_mean


#     base = plt.cm.gray(plt.Normalize(img_data.min(), img_data.max())(img_data))[..., :3]
#     heat = plt.cm.seismic(plt.Normalize(diff_min, diff_max)(diff))[..., :3]
#     ax.imshow((1 - alpha_diff) * base + alpha_diff * heat, interpolation="nearest")


    
#     # ax.imshow(diff, cmap="seismic", alpha=alpha_diff, vmin=diff_min, vmax=diff_max, interpolation="nearest", rasterized=True)
    
#     dist_val = dists_val[i]
#     # ax.set_title(f"$|v|={speed[idx]:.2f}$\n$Dist={dist_val:.2f}$", fontsize=9)
    
#     for spine in ax.spines.values():
#         spine.set_edgecolor(outlier_maxD_color); spine.set_linewidth(6); 
#         # spine.set_linestyle('--')
#     ax.set_xticks([]); ax.set_yticks([])
#     if i == 0: ax.set_ylabel("Max Distance", fontsize=12, fontweight='bold')

# for i, idx in enumerate(outliers_vel):
#     ax = axes3[1, i]
#     img_data = dset_samples_look[0][idx, 0] + mean_img
#     if hasattr(img_data, 'detach'): img_data = img_data.detach().cpu().numpy()
    
#     ax.imshow(img_data, cmap='gray', interpolation="nearest", rasterized=True)
#     # ax.set_title(f"$|v|={speed[idx]:.2f}$", fontsize=9)
#     diff = dset_samples_look[0][idx, 0] - data_mean

#     base = plt.cm.gray(plt.Normalize(img_data.min(), img_data.max())(img_data))[..., :3]
#     heat = plt.cm.seismic(plt.Normalize(diff_min, diff_max)(diff))[..., :3]
#     ax.imshow((1 - alpha_diff) * base + alpha_diff * heat, interpolation="nearest")

    
#     # ax.imshow(diff, cmap="seismic", alpha=alpha_diff, vmin=diff_min, vmax=diff_max, interpolation="nearest", rasterized=True)
    
#     for spine in ax.spines.values():
#         spine.set_edgecolor(outlier_maxV_color); spine.set_linewidth(6)
#     ax.set_xticks([]); ax.set_yticks([])
#     if i == 0: ax.set_ylabel("Max Velocity", fontsize=12, fontweight='bold')

# fig3.suptitle("Outlier Analysis", fontsize=16)
# plt.tight_layout()

# plt.savefig(plotFolder + "/4_outlier_img.pdf")
# plt.savefig(plotFolder + "/4_outlier_img.svg")
# plt.savefig(plotFolder + "/4_outlier_img.png")

# 4. outliers: k rows × 2 cols
fig3, axes3 = plt.subplots(k_outliers, 2, figsize=(4, 4))

for i, idx in enumerate(outliers_dist):
    ax = axes3[i, 0]   # column 0 = Max Distance
    img_data = dset_samples_look[0][idx, 0] + mean_img
    if hasattr(img_data, "detach"): img_data = img_data.detach().cpu().numpy()

    diff = dset_samples_look[0][idx, 0] - data_mean

    base = plt.cm.gray(plt.Normalize(img_data.min(), img_data.max())(img_data))[..., :3]
    # heat = plt.cm.seismic(plt.Normalize(diff_min, diff_max)(diff))[..., :3]
    heat = plt.cm.seismic(diff_norm(diff))[..., :3]
    ax.imshow((1 - alpha_diff) * base + alpha_diff * heat, interpolation="nearest")

    for spine in ax.spines.values():
        spine.set_edgecolor(outlier_maxD_color)
        spine.set_linewidth(6)

    ax.set_xticks([]); ax.set_yticks([])
    if i == 0:
        ax.set_title("Max Distance", fontsize=12, fontweight="bold")


for i, idx in enumerate(outliers_vel):
    ax = axes3[i, 1]   # column 1 = Max Velocity
    img_data = dset_samples_look[0][idx, 0] + mean_img
    if hasattr(img_data, "detach"): img_data = img_data.detach().cpu().numpy()

    diff = dset_samples_look[0][idx, 0] - data_mean

    base = plt.cm.gray(plt.Normalize(img_data.min(), img_data.max())(img_data))[..., :3]
    # heat = plt.cm.seismic(plt.Normalize(diff_min, diff_max)(diff))[..., :3]
    heat = plt.cm.seismic(diff_norm(diff))[..., :3]
    ax.imshow((1 - alpha_diff) * base + alpha_diff * heat, interpolation="nearest")

    for spine in ax.spines.values():
        spine.set_edgecolor(outlier_maxV_color)
        spine.set_linewidth(6)

    ax.set_xticks([]); ax.set_yticks([])
    if i == 0:
        ax.set_title("Max Velocity", fontsize=12, fontweight="bold")


fig3.suptitle("Outlier Analysis", fontsize=16)
plt.tight_layout()
plt.savefig(plotFolder + "/4_outlier_img.pdf")
plt.savefig(plotFolder + "/4_outlier_img.svg")
plt.savefig(plotFolder + "/4_outlier_img.png")


# %% [code]
# 5. band evolution
fig4, axes4 = plt.subplots(3, 4, figsize=(6.5, 6))
positions = ['Top', 'Mid', 'Low']

for col_idx, band in enumerate(bands):
    c = band_colors[col_idx]
    reps = band['reps']
    for row_idx, idx in enumerate(reps):
        ax = axes4[row_idx, col_idx]
        
        img_data = dset_samples_look[0][idx, 0] + mean_img
        if hasattr(img_data, 'detach'): img_data = img_data.detach().cpu().numpy()

        if row_idx == 0:
            ax.text(0.5, 1.22, f"Band {col_idx+1}",
                    transform=ax.transAxes, ha="center", va="bottom",
                    fontsize=14, fontweight="bold", color=c, clip_on=False)
        #     ax.text(0.5, 1.06, f"$|v|={speed[idx]:.2f}, z={mu[idx,2]:.1f}$",
        #             transform=ax.transAxes, ha="center", va="bottom",
        #             fontsize=10, clip_on=False)
        # else:
        #     ax.set_title(f"$|v|={speed[idx]:.2f}, z={mu[idx,2]:.1f}$", fontsize=10)
        
        ax.imshow(img_data, cmap='gray', interpolation="nearest", rasterized=True)
        diff = dset_samples_look[0][idx, 0] - data_mean

        base = plt.cm.gray(plt.Normalize(img_data.min(), img_data.max())(img_data))[..., :3]
        # heat = plt.cm.seismic(plt.Normalize(diff_min, diff_max)(diff))[..., :3]
        heat = plt.cm.seismic(diff_norm(diff))[..., :3]

        ax.imshow((1 - alpha_diff) * base + alpha_diff * heat, interpolation="nearest")
        # ax.imshow(diff, cmap="seismic", alpha=alpha_diff, vmin=diff_min, vmax=diff_max, interpolation="nearest", rasterized=True)
        
        for spine in ax.spines.values():
            spine.set_edgecolor(c); spine.set_linewidth(4)
        ax.set_xticks([]); ax.set_yticks([])
        

        if col_idx == 0:
            ax.set_ylabel(positions[row_idx], fontsize=14, fontweight='bold')

fig4.suptitle("Band Evolution (Top to Low)", fontsize=16)
plt.tight_layout()

plt.savefig(plotFolder + "/5_band_img.pdf")
plt.savefig(plotFolder + "/5_band_img.svg")
plt.savefig(plotFolder + "/5_band_img.png")

# %% [code]
fig5, axes5 = plt.subplots(3, 4, figsize=(7, 7))

imgs_all = []
for idx in non_outlier_indices:
    img = dset_samples_look[0][idx, 0] + mean_img
    if hasattr(img, 'detach'):
        img = img.detach().cpu().numpy()
    imgs_all.append(img)
imgs_all = np.asarray(imgs_all, dtype=np.float32)
global_mean_img = imgs_all.mean(axis=0)

band_mean_imgs = []
band_std_imgs = []
for band in bands:
    imgs = []
    for idx in band['indices']:
        img = dset_samples_look[0][idx, 0] + mean_img
        if hasattr(img, 'detach'):
            img = img.detach().cpu().numpy()
        imgs.append(img)
    imgs = np.asarray(imgs, dtype=np.float32)
    band_mean_imgs.append(imgs.mean(axis=0))
    band_std_imgs.append(imgs.std(axis=0))

diff_imgs = [m - global_mean_img for m in band_mean_imgs]
max_abs_diff = max(float(np.max(np.abs(d))) for d in diff_imgs)
max_std = max(float(np.max(s)) for s in band_std_imgs)

for col_idx, band in enumerate(bands):
    c = band_colors[col_idx]
    idxs = band['indices']
    n = len(idxs)
    mean_speed = float(np.mean(speed[idxs])) if n > 0 else 0.0

    ax = axes5[0, col_idx]
    ax.imshow(band_mean_imgs[col_idx], cmap='gray')
    ax.set_xticks([]); ax.set_yticks([])

    ax.text(0.5, 1.22, f"Band {col_idx+1}",
            transform=ax.transAxes, ha="center", va="bottom",
            fontsize=14, fontweight="bold", color=c, clip_on=False)
    ax.text(0.5, 1.06, f"n={n}, mean|v|={mean_speed:.2f}",
            transform=ax.transAxes, ha="center", va="bottom",
            fontsize=9, clip_on=False)

    ax = axes5[1, col_idx]
    ax.imshow(diff_imgs[col_idx], cmap='seismic', vmin=-max_abs_diff, vmax=max_abs_diff)
    ax.set_xticks([]); ax.set_yticks([])

    ax = axes5[2, col_idx]
    ax.imshow(band_std_imgs[col_idx], cmap='gray', vmin=0.0, vmax=max_std)
    ax.set_xticks([]); ax.set_yticks([])

    # Colored borders (match Fig3 style)
    for r in range(3):
        for spine in axes5[r, col_idx].spines.values():
            spine.set_edgecolor(c)
            spine.set_linewidth(4)

# Row labels
axes5[0, 0].set_ylabel("Mean", fontsize=14, fontweight='bold')
axes5[1, 0].set_ylabel("Mean-Global", fontsize=14, fontweight='bold')
axes5[2, 0].set_ylabel("Std", fontsize=14, fontweight='bold')

fig5.suptitle("Band Fingerprints", fontsize=16)
plt.tight_layout(rect=[0, 0.20, 1, 1.0])

sm_diff = mpl.cm.ScalarMappable(
    norm=mpl.colors.Normalize(vmin=-max_abs_diff, vmax=max_abs_diff),
    cmap='seismic'
)
sm_diff.set_array([])

sm_std = mpl.cm.ScalarMappable(
    norm=mpl.colors.Normalize(vmin=0.0, vmax=max_std),
    cmap='gray'
)
sm_std.set_array([])

# side-by-side colorbars
cbax_diff = fig5.add_axes([0.12, 0.2, 0.36, 0.03])  # left bar
cbar_diff = fig5.colorbar(sm_diff, cax=cbax_diff, orientation='horizontal')
cbar_diff.set_label("Mean - Global", fontsize=11)

cbax_std = fig5.add_axes([0.56, 0.2, 0.36, 0.03])   # right bar
cbar_std = fig5.colorbar(sm_std, cax=cbax_std, orientation='horizontal')
cbar_std.set_label("Std", fontsize=11)

plt.savefig(plotFolder + "/6_band_fingerprint.pdf")
plt.savefig(plotFolder + "/6_band_fingerprint.svg")
plt.savefig(plotFolder + "/6_band_fingerprint.png")

# %% [code]
# 4. outliers
# fig3, axes3 = plt.subplots(2, k_outliers, figsize=(6, 4))

# for i, idx in enumerate(outliers_dist):
#     ax = axes3[0, i]
#     img_data = dset_samples_look[0][idx, 0] + mean_img
#     if hasattr(img_data, 'detach'): img_data = img_data.detach().cpu().numpy()
    
#     ax.imshow(img_data, cmap='gray', interpolation="nearest", rasterized=True)
#     diff = dset_samples_look[0][idx, 0] - data_mean

#     base = plt.cm.gray(plt.Normalize(img_data.min(), img_data.max())(img_data))[..., :3]
#     heat = plt.cm.seismic(plt.Normalize(diff_min, diff_max)(diff))[..., :3]
#     ax.imshow((1 - alpha_diff) * base + alpha_diff * heat, interpolation="nearest")
    
#     # ax.imshow(diff, cmap="seismic", alpha=alpha_diff, vmin=diff_min, vmax=diff_max, interpolation="nearest", rasterized=True)
    
#     dist_val = dists_val[i]
#     # ax.set_title(f"$|v|={speed[idx]:.2f}$\n$Dist={dist_val:.2f}$", fontsize=9)
    
#     for spine in ax.spines.values():
#         spine.set_edgecolor(outlier_maxD_color); spine.set_linewidth(6); 
#         # spine.set_linestyle('--')
#     ax.set_xticks([]); ax.set_yticks([])
#     if i == 0: ax.set_ylabel("Max Distance", fontsize=12, fontweight='bold')

# for i, idx in enumerate(outliers_vel):
#     ax = axes3[1, i]
#     img_data = dset_samples_look[0][idx, 0] + mean_img
#     if hasattr(img_data, 'detach'): img_data = img_data.detach().cpu().numpy()
    
#     ax.imshow(img_data, cmap='gray', interpolation="nearest", rasterized=True)
#     # ax.set_title(f"$|v|={speed[idx]:.2f}$", fontsize=9)
#     diff = dset_samples_look[0][idx, 0] - data_mean

#     base = plt.cm.gray(plt.Normalize(img_data.min(), img_data.max())(img_data))[..., :3]
#     heat = plt.cm.seismic(plt.Normalize(diff_min, diff_max)(diff))[..., :3]
#     ax.imshow((1 - alpha_diff) * base + alpha_diff * heat, interpolation="nearest")

    
#     # ax.imshow(diff, cmap="seismic", alpha=alpha_diff, vmin=diff_min, vmax=diff_max, interpolation="nearest", rasterized=True)
    
#     for spine in ax.spines.values():
#         spine.set_edgecolor(outlier_maxV_color); spine.set_linewidth(6)
#     ax.set_xticks([]); ax.set_yticks([])
#     if i == 0: ax.set_ylabel("Max Velocity", fontsize=12, fontweight='bold')
fig3, axes3 = plt.subplots(k_outliers, 2, figsize=(4, 4))

for i, idx in enumerate(outliers_dist):
    ax = axes3[i, 0]   # column 0 = Max Distance
    img_data = dset_samples_look[0][idx, 0] + mean_img
    if hasattr(img_data, "detach"): img_data = img_data.detach().cpu().numpy()

    diff = dset_samples_look[0][idx, 0] - data_mean

    base = plt.cm.gray(plt.Normalize(img_data.min(), img_data.max())(img_data))[..., :3]
    # heat = plt.cm.seismic(plt.Normalize(diff_min, diff_max)(diff))[..., :3]
    heat = plt.cm.seismic(diff_norm(diff))[..., :3]

    ax.imshow((1 - alpha_diff) * base + alpha_diff * heat, interpolation="nearest")

    for spine in ax.spines.values():
        spine.set_edgecolor(outlier_maxD_color)
        spine.set_linewidth(6)

    ax.set_xticks([]); ax.set_yticks([])
    if i == 0:
        ax.set_title("Max Distance", fontsize=12, fontweight="bold")


for i, idx in enumerate(outliers_vel):
    ax = axes3[i, 1]   # column 1 = Max Velocity
    img_data = dset_samples_look[0][idx, 0] + mean_img
    if hasattr(img_data, "detach"): img_data = img_data.detach().cpu().numpy()

    diff = dset_samples_look[0][idx, 0] - data_mean

    base = plt.cm.gray(plt.Normalize(img_data.min(), img_data.max())(img_data))[..., :3]
    # heat = plt.cm.seismic(plt.Normalize(diff_min, diff_max)(diff))[..., :3]
    heat = plt.cm.seismic(diff_norm(diff))[..., :3]

    ax.imshow((1 - alpha_diff) * base + alpha_diff * heat, interpolation="nearest")

    for spine in ax.spines.values():
        spine.set_edgecolor(outlier_maxV_color)
        spine.set_linewidth(6)

    ax.set_xticks([]); ax.set_yticks([])
    if i == 0:
        ax.set_title("Max Velocity", fontsize=12, fontweight="bold")


fig3.suptitle("Outlier Analysis", fontsize=16)
plt.tight_layout()

# fig3.suptitle("Outlier Analysis", fontsize=16)

# fig3.tight_layout(rect=[0.0, 0.0, 0.90, 0.95])

pos = [ax.get_position() for ax in axes3.ravel()]
y0 = min(p.y0 for p in pos)
y1 = max(p.y1 for p in pos)
x1 = max(p.x1 for p in pos)

pad = 0.015
cbar_w = 0.045
cax = fig3.add_axes([x1 + pad, y0, cbar_w, y1 - y0])


# sm = cm.ScalarMappable(norm=Normalize(vmin=diff_min, vmax=diff_max), cmap="seismic")
sm = cm.ScalarMappable(norm=diff_norm, cmap="seismic")

sm.set_array([])

cbar = fig3.colorbar(sm, cax=cax)
cbar.set_label("img - mean")


plt.savefig(plotFolder + "/7_outlier_img_cbar.pdf",
             bbox_inches="tight", facecolor="white", dpi=300)
plt.savefig(plotFolder + "/7_outlier_img_cbar.svg")
plt.savefig(plotFolder + "/7_outlier_img_cbar.png")

# %% [code]


# %% [code]


# %% [code]
```
