## Supplementary material for "Dynamic Compression Flows for Neuroscience Data": Python code: plot_musal_v5.py

```
import numpy as np
import matplotlib.pyplot as plt
from matplotlib.collections import LineCollection
from matplotlib.colors import ListedColormap, BoundaryNorm, LinearSegmentedColormap, Normalize
from mpl_toolkits.mplot3d.art3d import Line3DCollection
import torch
import dnnlib.util_v5 as util
from tqdm import tqdm
import pickle
from matplotlib.ticker import FuncFormatter
import gc
from PIL import Image

import numpy as np
import pickle
import os
import re 
import json 
import click
import torch
from torch_utils import distributed as dist
from training import toy_training_loop_vfm_noDataGen
from tqdm import tqdm
import matplotlib.pyplot as plt
from matplotlib import colors
from plot_neural import *
from plot_sim_results_v5 import *
from plot_sim_results_img_v5 import * 

import pandas as pd

import warnings
warnings.filterwarnings('ignore', 'Grad strides do not match bucket view strides') # False warning 
device = torch.device("cuda" if torch.cuda.is_available() else "cpu")
torch.set_default_device(device)
colorgrad = "PuOr"


def scale_factor_like_plot(mu_traj_np, trial_indices, dims=(0,1,2), z_scale='auto'):
    # gather ranges exactly like your function
    xs, ys, zs = [], [], []
    for idx in trial_indices:
        S = mu_traj_np[idx]
        if S is None or (hasattr(S, "size") and S.size == 0):  # numpy
            continue
        A = np.asarray(S)
        xs.extend(A[:, dims[0]].tolist())
        ys.extend(A[:, dims[1]].tolist())
        zs.extend(A[:, dims[2]].tolist())
    xR = (max(xs) - min(xs)) if xs else 1.0
    yR = (max(ys) - min(ys)) if ys else 1.0
    zR = (max(zs) - min(zs)) if zs else 1.0
    if z_scale == 'auto':
        xy_avg = (xR + yR) / 2.0
        return (xy_avg / zR) if zR > 0 else 1.0
    if z_scale == 'none':
        return 1.0
    return float(z_scale)


def plot_selected_image(topk_idx, dset_samples_look, mean_img, h=3,
                        coords=None, row_color=None, fmt=".3f"):
    W = 2*h + 1
    idxs = torch.as_tensor(topk_idx).cpu().tolist()
    T = np.asarray(dset_samples_look[0]).shape[0]
    imgs = np.asarray(dset_samples_look[0])
    mean = np.asarray(mean_img)

    fig, axes = plt.subplots(len(idxs), W, figsize=(W*2.0, max(1, len(idxs))*2.0))
    if len(idxs) == 1:
        axes = np.expand_dims(axes, 0)

    for r, i in enumerate(idxs):
        cols_ts = list(range(i - h, i + h + 1))
        for c, t in enumerate(cols_ts):
            ax = axes[r, c]
            if 0 <= t < T:
                ax.imshow(imgs[t, 0] + mean, cmap='gray')
                rel = t - i
                if rel == 0:
                    # center frame title
                    if coords is not None and i in coords:
                        u1, u2, u3 = coords[i]
                        ttl = f"t={t}\n[μ1={u1:{fmt}}, μ2={u2:{fmt}}, μ3={u3:{fmt}}]"
                        ax.set_title(ttl, fontsize=12,
                                     color=(row_color if row_color is not None else 'k'),
                                     fontweight='bold')
                    else:
                        ax.set_title(f"t={t}", fontsize=12)
                else:
                    ax.set_title(f"t+{rel}" if rel > 0 else f"t{rel}", fontsize=10)
            ax.axis("off")
    plt.tight_layout()


def plot_frame(data, T, trial_idx, mean_img = None):
    idxs = np.linspace(0, T-1, 6, dtype=int)
    fig, axes = plt.subplots(1, len(idxs), figsize=(12, 2))
    for ax, i in zip(axes, idxs):
        if mean_img is None:
            ax.imshow(data[trial_idx][i,0,:,:], cmap = 'gray')
        else:
            ax.imshow(data[trial_idx][i,0,:,:] + mean_img, cmap = 'gray')
        ax.set_title(f"t={i}")
        ax.axis("off")
    plt.tight_layout()
    # plt.show()

def plot_grad_frame(data, trial_idx, n=6, idxs=None, signed=False, it_mode='forward', idxs_cbar=None):
    """
    it_mode: 'forward' | 'backward' | 'central'
    idxs_cbar: indices used to set the colorbar range (defaults to idxs)
    """
    x = data[trial_idx]                  # (T, 1, H, W)
    T = x.shape[0]
    X = x[:, 0]

    if it_mode == 'forward':
        g = X[1:] - X[:-1];         i_min, i_max = 0, T-2
        title_fn = lambda i: f"t={i}→{i+1}"
        map_g_idx = lambda i: i
    elif it_mode == 'backward':
        g = X[1:] - X[:-1];         i_min, i_max = 1, T-1
        title_fn = lambda i: f"t={i-1}→{i}"
        map_g_idx = lambda i: i-1
    elif it_mode == 'central':
        g = 0.5 * (X[2:] - X[:-2]); i_min, i_max = 1, T-2
        title_fn = lambda i: f"centered t={i}"
        map_g_idx = lambda i: i-1
    else:
        raise ValueError("it_mode must be 'forward', 'backward', or 'central'.")

    if idxs is None:
        if i_max < i_min: raise ValueError(f"Not enough frames for it_mode='{it_mode}'.")
        idxs = np.linspace(i_min, i_max, n, dtype=int)
    else:
        idxs = [i for i in idxs if i_min <= i <= i_max]
        if not idxs: raise ValueError(f"All idxs out of range for it_mode='{it_mode}' (valid {i_min}..{i_max}).")

    # --- NEW: colorbar indices (defaults to idxs), clipped to valid range
    if idxs_cbar is None:
        idxs_cbar = idxs
    else:
        idxs_cbar = [i for i in idxs_cbar if i_min <= i <= i_max]
        if not idxs_cbar: raise ValueError(f"All idxs_cbar out of range for it_mode='{it_mode}' (valid {i_min}..{i_max}).")
    g_idx_cbar = [map_g_idx(i) for i in idxs_cbar]
    g_for_cbar = g[g_idx_cbar]

    # vmin/vmax from idxs_cbar
    if signed:
        vmax = np.max(np.abs(g_for_cbar)); vmin, vmax = -vmax, vmax
        cmap = 'seismic'; transform = (lambda arr: arr)
    else:
        vmax = np.max(np.abs(g_for_cbar)); vmin = 0.0
        cmap = 'viridis'; transform = np.abs

    fig, axes = plt.subplots(1, len(idxs), figsize=(12, 2))
    ims = []
    for ax, i in zip(axes, idxs):
        gi = map_g_idx(i)
        im = ax.imshow(transform(g[gi]), cmap=cmap, vmin=vmin, vmax=vmax)
        ax.set_title(title_fn(i)); ax.axis("off"); ims.append(im)

    cbar = fig.colorbar(ims[0], ax=axes, orientation='horizontal', fraction=0.06, pad=0.02)
    cbar.set_label('temporal gradient' if signed else '|temporal gradient|')
    plt.tight_layout(rect=[0, 0.12, 1, 1])


def _box_sum(A, r):
    P = np.pad(A, ((r+1, r), (r+1, r)), mode='edge')
    I = P.cumsum(0).cumsum(1)
    return (I[2*r+1:, 2*r+1:] - I[:-2*r-1, 2*r+1:]
            - I[2*r+1:, :-2*r-1] + I[:-2*r-1, :-2*r-1])

# Lucas–Kanade optical flow
def lk_flow(img0, img1, r=2, lam=1e-3, imgm1=None, it_mode='central'):
    """
    it_mode controls how I_t is computed at time t:
      'forward' : I_t = img1 - img0                (aligned at t in [0..T-2])
      'backward': I_t = img0 - imgm1               (requires imgm1; aligned at t in [1..T-1])
      'central' : I_t = 0.5*(img1 - imgm1)         (requires imgm1; aligned at t in [1..T-2])
    """
    Ibar = 0.5*(img0 + img1)
    Iy, Ix = np.gradient(Ibar)

    if it_mode == 'forward' or imgm1 is None:
        It = (img1 - img0)
    elif it_mode == 'backward':
        It = (img0 - imgm1)
    elif it_mode == 'central':
        It = 0.5 * (img1 - imgm1)
    else:
        raise ValueError("it_mode must be 'forward', 'backward', or 'central'.")

    Sxx = _box_sum(Ix*Ix, r);  Syy = _box_sum(Iy*Iy, r);  Sxy = _box_sum(Ix*Iy, r)
    Sxt = _box_sum(Ix*It, r);  Syt = _box_sum(Iy*It, r)
    det = Sxx*Syy - Sxy*Sxy + lam
    u = (-Syy*Sxt + Sxy*Syt) / det
    v = ( Sxy*Sxt - Sxx*Syt) / det
    return u, v, np.abs(It)


# custom "white -> LIGHT red" cmap with transparency increasing with value
def _light_red_cmap(alpha_max=0.95, alpha_min=0.10,
                    red_rgb=(1.0, 0.45, 0.45), deepen=0.15):
    """
    alpha_max: opacity for largest gradients (0..1)
    alpha_min: baseline opacity so faint gradients are still visible
    red_rgb:   base red tint (R,G,B)
    deepen:    how much to further reduce G,B at the top stop (more = stronger red)
    """
    r, g, b = red_rgb
    top_rgba = (1.0, max(0.0, g - deepen), max(0.0, b - deepen), alpha_max)  # deeper red + more opaque
    low_rgba = (1.0, 1.0, 1.0, alpha_min)                                    # white with some opacity
    return LinearSegmentedColormap.from_list("white_to_stronger_red", [low_rgba, top_rgba])

def plot_optical_flow_frames(
        data, trial_idx, n=6, idxs=None,
        r=2, lam=1e-3, stride=2,
        img_alpha=0.30,
        arrow_scale=8.0, arrow_width=0.012,
        grad_quantile=99,
        show_image_bg=True,
        mean_img=None,
        arrow_scale_mode='global',
        draw_arrows=True,
        signed_grad=False,
        grad_it_mode='forward',
        flow_it_mode='forward',
        idxs_cbar=None,                   # NEW: indices to set colorbar range
):
    x = data[trial_idx][:, 0]  # (T,H,W)
    T = x.shape[0]; H, W = x.shape[1:]

    def valid_range(T, mode):
        if mode == 'forward':  return 0, T-2
        if mode == 'backward': return 1, T-1
        if mode == 'central':  return 1, T-2
        raise ValueError

    lo_g, hi_g = valid_range(T, grad_it_mode)
    lo_f, hi_f = valid_range(T, flow_it_mode)
    lo, hi = max(lo_g, lo_f), min(hi_g, hi_f)
    if lo > hi:
        raise ValueError(f"Not enough frames for grad_it_mode='{grad_it_mode}' and flow_it_mode='{flow_it_mode}'.")

    if idxs is None:
        idxs = np.linspace(lo, hi, n, dtype=int)
    else:
        idxs = [i for i in idxs if lo <= i <= hi]
        if not idxs: raise ValueError(f"All idxs out of range (valid {lo}..{hi}).")

    # --- NEW: colorbar indices (defaults to idxs), but must be valid for grad_it_mode
    if idxs_cbar is None:
        idxs_cbar = idxs
    else:
        lo_cb, hi_cb = valid_range(T, grad_it_mode)
        idxs_cbar = [i for i in idxs_cbar if lo_cb <= i <= hi_cb]
        if not idxs_cbar: raise ValueError(f"All idxs_cbar out of range for grad_it_mode='{grad_it_mode}' (valid {lo_cb}..{hi_cb}).")

    def grad_at(i):
        if grad_it_mode == 'forward':   g = x[i+1] - x[i]
        elif grad_it_mode == 'backward': g = x[i] - x[i-1]
        else:                            g = 0.5*(x[i+1] - x[i-1])
        return g if signed_grad else np.abs(g)

    if grad_it_mode == 'forward':
        title_fn = lambda i: f"t={i}→{i+1}"
    elif grad_it_mode == 'backward':
        title_fn = lambda i: f"t={i-1}→{i}"
    else:
        title_fn = lambda i: f"centered t={i}"

    yy, xx = np.meshgrid(np.arange(H), np.arange(W), indexing='ij')

    # Gradients for display and for colorbar range
    grads = [grad_at(i) for i in idxs]
    grads_cbar = [grad_at(i) for i in idxs_cbar]
    all_gm = np.stack(grads_cbar, 0)

    # Color scaling from idxs_cbar
    if signed_grad:
        gmax = np.percentile(np.abs(all_gm), grad_quantile) + 1e-9
        cmap = 'seismic'; use_norm = False
        vmin, vmax = -gmax, gmax; norm = None
        grad_label = 'temporal gradient (signed)'
    else:
        gm_vmax = np.percentile(all_gm, grad_quantile) + 1e-9
        cmap = _light_red_cmap(alpha_max=0.9, alpha_min=0.12, red_rgb=(1.0, 0.50, 0.50), deepen=0.10)
        use_norm = True; vmin, vmax = 0.0, gm_vmax
        norm = Normalize(vmin=0.0, vmax=gm_vmax)
        grad_label = '|temporal gradient|'

    # Flows (optional)
    flows = []
    if draw_arrows:
        for i in idxs:
            prev = x[i-1] if i-1 >= 0 else None
            u, v, _ = lk_flow(x[i], x[i+1], r=r, lam=lam, imgm1=prev, it_mode=flow_it_mode)
            flows.append((u, v))
        if arrow_scale_mode == 'global':
            all_mag = [np.hypot(u, v) for (u, v) in flows]
            global_denom = np.percentile(np.stack(all_mag), 98) + 1e-9

    fig, axes = plt.subplots(1, len(idxs), figsize=(12, 2.8), constrained_layout=True)
    axes = np.atleast_1d(axes)
    mappable_for_cb = None

    for ax, i, gm in zip(axes, idxs, grads):
        if show_image_bg:
            base = x[i] if mean_img is None else (x[i] + mean_img)
            ax.imshow(base, origin='upper', cmap='gray', alpha=img_alpha)

        im = ax.imshow(gm, origin='upper', cmap=cmap,
                       norm=norm if use_norm else None,
                       vmin=None if use_norm else vmin,
                       vmax=None if use_norm else vmax)
        if mappable_for_cb is None: mappable_for_cb = im

        if draw_arrows:
            u, v = flows.pop(0)
            if arrow_scale_mode == 'global':
                s = arrow_scale / global_denom
            else:
                mag = np.hypot(u, v)
                denom = np.percentile(mag, 98) + 1e-9
                s = arrow_scale / denom
            ax.quiver(xx[::stride, ::stride], yy[::stride, ::stride],
                      (u*s)[::stride, ::stride], (v*s)[::stride, ::stride],
                      color='black', angles='xy', scale_units='xy', scale=1.0,
                      pivot='middle', width=arrow_width)

        ax.set_title(title_fn(i)); ax.axis('off')

    cb = fig.colorbar(mappable_for_cb, ax=axes, orientation='horizontal', fraction=0.06, pad=0.08)
    cb.set_label(grad_label)


def mu2_bands_from_hist(y, m=3, bins=80, smooth=5):
    y_np = y.detach().cpu().numpy()
    cnt, edges = np.histogram(y_np, bins=bins)
    # gentle smoothing
    if smooth > 0:
        ker = np.array([1,2,3,2,1], float)
        ker /= ker.sum()
        for _ in range(smooth): cnt = np.convolve(cnt, ker, mode='same')
    centers = (edges[:-1] + edges[1:]) / 2

    # peaks
    pk = np.where((cnt[1:-1] > cnt[:-2]) & (cnt[1:-1] >= cnt[2:]))[0] + 1
    if pk.size == 0: pk = np.array([cnt.argmax()])
    pk = sorted(pk, key=lambda i: cnt[i], reverse=True)[:m]
    pk.sort()  # low → high

    # boundaries: minima between consecutive peaks
    bounds = [-np.inf]
    for a,b in zip(pk[:-1], pk[1:]):
        valley = a + np.argmin(cnt[a:b+1])
        boundary = centers[valley]
        bounds.append(boundary)
    bounds.append(np.inf)

    # label each point
    labels = np.zeros_like(y_np, dtype=int)
    for bi in range(m):
        mask = (y_np > bounds[bi]) & (y_np <= bounds[bi+1])
        labels[mask] = bi
    return torch.from_numpy(labels).to(y.device), torch.from_numpy(centers[pk]).to(y.device)

def reps_for_band(band_id, k, z_keep_q_seq=(0.20, 0.30, 0.50, 0.75, 1.00)):
    idx_all = torch.nonzero(band == band_id, as_tuple=False).squeeze(1)
    if idx_all.numel() == 0:
        return []

    # --- 1) ON-BAND in μ3 (use plotted z = mu3) ---
    z = mu3[idx_all]
    z_med = z.median()
    z_dev = (z - z_med).abs()

    cand = None
    for q in z_keep_q_seq:                          # progressively widen if too tight
        thr = torch.quantile(z_dev, q)
        cand = idx_all[z_dev <= thr]
        if cand.numel() >= min(k, idx_all.numel()):
            break
    if cand is None or cand.numel() == 0:
        order = torch.argsort(z_dev)
        return idx_all[order[:min(k, idx_all.numel())]].cpu().tolist()

    # --- 2) Spread along μ1 inside this band (greedy farthest-point) ---
    x = mu1[cand]
    if cand.numel() <= k:
        return cand.cpu().tolist()

    # seed: closest to μ1 median (stable)
    seed_rel = (x - x.median()).abs().argmin()
    chosen_abs = [int(cand[seed_rel].item())]

    # remaining pool
    mask = torch.ones(cand.numel(), dtype=torch.bool, device=cand.device)
    mask[seed_rel] = False
    remaining = cand[mask]
    if remaining.numel() == 0:
        return chosen_abs

    # track min |Δμ1| to chosen set
    min_dist = (mu1[remaining] - mu1[chosen_abs[0]]).abs()
    while len(chosen_abs) < k and remaining.numel() > 0:
        j_rel = torch.argmax(min_dist)
        j_abs = int(remaining[j_rel].item())
        chosen_abs.append(j_abs)

        if remaining.numel() == 1:
            break
        keep = torch.arange(remaining.numel(), device=remaining.device) != j_rel
        remaining = remaining[keep]
        if remaining.numel() == 0:
            break
        new_d = (mu1[remaining] - mu1[j_abs]).abs()
        min_dist = torch.minimum(min_dist[keep], new_d)

    return chosen_abs


def band_mu1_range(b, qlo=0.10, qhi=0.90):
    idx = torch.nonzero(band == b, as_tuple=False).squeeze(1)
    if idx.numel() == 0:
        return None
    lo = torch.quantile(mu1[idx], qlo)
    hi = torch.quantile(mu1[idx], qhi)
    return lo, hi


def _kmeans1d(s, K=3, iters=25):
    s = s.float().view(-1)
    qs = torch.linspace(0.1, 0.9, K, device=s.device)
    c  = torch.quantile(s, qs)                   # quantile init
    for _ in range(iters):
        a = ((s[:, None] - c[None, :])**2).argmin(dim=1)
        new_c = torch.stack([s[a==j].mean() if (a==j).any() else c[j] for j in range(K)])
        if torch.allclose(new_c, c, atol=1e-7): break
        c = new_c
    order = torch.argsort(c); c = c[order]
    remap = torch.empty_like(order); remap[order] = torch.arange(K, device=s.device)
    a = remap[((s[:, None] - c[None, :])**2).argmin(dim=1)]
    return c, a

def _sep_score(s, K=3):
    c, a = _kmeans1d(s, K)
    N = s.numel()
    # pooled stdev
    sig2 = 0.0
    for j in range(K):
        sj = s[a==j]
        if sj.numel() >= 2:
            sig2 += sj.var(unbiased=False) * sj.numel()
    sig = torch.sqrt(sig2 / max(N, 1))
    gaps = c[1:] - c[:-1]
    min_gap = gaps.min() if gaps.numel() else torch.tensor(0.0, device=s.device)
    # small size penalty to avoid degenerate tiny bands
    size_pen = torch.stack([(a==j).float().mean() for j in range(K)]).min()
    return (min_gap / (sig + 1e-12)) * (0.5 + 0.5*size_pen), c, a

def find_bands_by_projection(x0, K=3, scale_z=1.0, num_dirs=48, seed=0):
    """
    x0: torch.Tensor [T,3] in your plot's coordinates (use raw x0; we scale z by scale_z)
    Returns:
      labels  : LongTensor [T] with values 0..K-1 (ordered low→high along best projection)
      idx_sets: list of index lists per band
      n       : best projection direction (unit vector, in scaled space)
      centers : 1D cluster centers along that projection
    """
    X = x0.detach().float().clone()
    X[:, 2] *= float(scale_z)                 # match your plotted z

    # center (translation doesn’t affect bands; helps direction search)
    Xm = X - X.mean(dim=0, keepdim=True)

    # PCA directions
    _, _, Vh = torch.linalg.svd(Xm, full_matrices=False)
    PCs = Vh.T  # 3x3 (columns)

    cand_dirs = []
    for i in range(3):
        v = PCs[:, i]; cand_dirs += [v, -v]
    combos = [PCs[:,0]+PCs[:,1], PCs[:,0]-PCs[:,1],
              PCs[:,0]+PCs[:,2], PCs[:,0]-PCs[:,2],
              PCs[:,1]+PCs[:,2], PCs[:,1]-PCs[:,2]]
    cand_dirs += combos

    # a few random directions
    g = torch.Generator(device=X.device); g.manual_seed(seed)
    R = torch.randn((3, num_dirs), generator=g, device=X.device)
    R = R / (torch.norm(R, dim=0, keepdim=True) + 1e-12)
    cand_dirs += [R[:,i] for i in range(R.shape[1])]

    best_score, best_v, best_centers, best_labels = None, None, None, None
    for v in cand_dirs:
        v = v / (torch.norm(v) + 1e-12)
        s = Xm @ v
        score, c, a = _sep_score(s, K)
        if (best_score is None) or (score > best_score):
            best_score, best_v, best_centers, best_labels = score, v, c, a

    labels = best_labels            # 0..K-1 ordered by center
    centers = best_centers
    idx_sets = [torch.nonzero(labels==b, as_tuple=False).squeeze(1).cpu().tolist()
                for b in range(K)]
    return labels, idx_sets, best_v, centers
def band_reps_onband(x0, labels, n, centers, band_id, m=2,
                     scale_z=1.0, keep_q=0.30, trim=0.05):
    """
    Pick m representative indices for one band:
      1) ON-BAND gate: keep the inner 'keep_q' quantile by distance along 'n'
         to that band's center (robust slab inliers).
      2) ALONG-BAND spread: PCA on inliers -> take trimmed quantiles (ends/middle).

    x0      : [T,3] raw coords (same you plot)
    labels  : LongTensor from find_bands_by_projection
    n       : unit normal returned by find_bands_by_projection (in scaled space)
    centers : 1D centers along the projection (same order as labels 0..K-1)
    band_id : which band
    m       : reps per band (2 => ends, 3 => start/middle/end)
    scale_z : same z-scale you used in the 3D plot
    keep_q  : keep this central fraction on the slab (e.g., 0.30 = tight)
    trim    : ignore extreme ends along the band when placing quantile targets
    """
    # scale & center exactly as in the finder
    X = x0.detach().float().clone()
    X[:, 2] *= float(scale_z)
    Xc = X - X.mean(0, keepdim=True)

    idx_band = torch.nonzero(labels == band_id, as_tuple=False).squeeze(1)
    if idx_band.numel() == 0:
        return []

    # --- ON-BAND gate along normal n ---
    s = Xc @ n                                   # projection used for band assignment
    d = (s[idx_band] - centers[band_id]).abs()   # distance to this slab’s center
    if idx_band.numel() <= m:
        inliers = idx_band
    else:
        tau = torch.quantile(d, keep_q)          # keep central keep_q fraction
        inliers = idx_band[d <= tau]
        if inliers.numel() < m:                  # gently widen once if too strict
            tau = torch.quantile(d, min(0.50, max(keep_q, 0.40)))
            inliers = idx_band[d <= tau]
        if inliers.numel() == 0:
            # last resort: the closest m to the slab center
            order = torch.argsort(d)
            return idx_band[order[:m]].cpu().tolist()

    # --- ALONG-BAND: ends / middle via PCA on inliers ---
    Xb = X[inliers]
    Xbc = Xb - Xb.mean(0, keepdim=True)
    _, _, Vh = torch.linalg.svd(Xbc, full_matrices=False)
    t = Vh.T[:, 0]                               # along-band axis (unit)

    u = Xbc @ t                                  # 1-D coordinate along the band
    m_eff = min(m, u.numel())
    qs = torch.tensor([0.5], device=u.device) if m_eff == 1 \
         else torch.linspace(trim, 1.0 - trim, steps=m_eff, device=u.device)
    targets = torch.quantile(u, qs)

    chosen = []
    avail = torch.arange(u.numel(), device=u.device)
    for tgt in targets:
        if avail.numel() == 0: break
        j_rel = torch.argmin((u[avail] - tgt).abs())
        j_band = int(avail[j_rel].item())        # index inside inliers
        chosen.append(int(inliers[j_band].item()))
        avail = avail[avail != j_band]           # no duplicates
    return chosen

def reps_for_all_bands_onband(x0, labels, n, centers, K, m=2, scale_z=1.0, keep_q=0.30, trim=0.05):
    return [band_reps_onband(x0, labels, n, centers, b, m=m, scale_z=scale_z,
                             keep_q=keep_q, trim=trim)
            for b in range(K)]
```
