## Supplementary material for "Dynamic Compression Flows for Neuroscience Data": Python code: plot_neural.py

```
import numpy as np
import matplotlib.pyplot as plt
from matplotlib.collections import LineCollection
from matplotlib.colors import LinearSegmentedColormap, Normalize, ListedColormap, BoundaryNorm

from mpl_toolkits.mplot3d.art3d import Line3DCollection
from mpl_toolkits.axes_grid1 import make_axes_locatable


#### 1. PCA related
## 1.1 PCA
def pca_stream(trials, n_components=None, dtype=np.float64):
    """
    trials: list of arrays with shapes [T_k, d] (d=137)
    n_components: keep top-k PCs (default: all)
    returns:
        mean (d,),
        components (d, k)  # columns are PC directions
        explained_variance (k,),
        explained_variance_ratio (k,),
        transform_fn  # function to stream-project new trials
    """
    # --- accumulate mean and M2 (sum of centered outer products) in a single pass
    N = 0
    d = trials[0].shape[1]
    mean = np.zeros(d, dtype=dtype)
    M2 = np.zeros((d, d), dtype=dtype)

    for X in trials:
        if X.size == 0: 
            continue
        X = np.asarray(X, dtype=dtype)
        m = X.shape[0]
        # block stats
        block_mean = X.mean(axis=0)
        Xc = X - block_mean
        block_M2 = Xc.T @ Xc  # d x d

        # combine A (N, mean, M2) with B (m, block_mean, block_M2)
        delta = block_mean - mean
        newN = N + m
        if newN > 0:
            mean = mean + delta * (m / newN)
        M2 = M2 + block_M2 + np.outer(delta, delta) * (N * m / max(newN, 1))
        N = newN

    if N < 2:
        raise ValueError("Not enough total samples for PCA.")

    # sample covariance
    cov = M2 / (N - 1)

    # eigendecomposition (cov is symmetric)
    evals, evecs = np.linalg.eigh(cov)
    order = np.argsort(evals)[::-1]
    evals = evals[order]
    evecs = evecs[:, order]

    # keep top-k
    if n_components is None or n_components > d:
        n_components = d
    evals_k = np.clip(evals[:n_components], 0, None)
    comps_k = evecs[:, :n_components]
    evr_k = evals_k / np.maximum(evals.sum(), 1e-12)

    # transformer that streams projections trial-by-trial
    def transform(trial_iterable):
        """
        Yields projected scores for each trial in trial_iterable
        without stacking, each as [T_k, n_components].
        """
        for X in trial_iterable:
            X = np.asarray(X, dtype=dtype)
            if X.size == 0:
                yield np.empty((0, n_components), dtype=dtype)
            else:
                yield (X - mean) @ comps_k

    return mean, comps_k, evals_k, evr_k, transform


## 1.2 scree plot
def plot_scree(explained_variance, explained_variance_ratio, fig_w = 10, fig_h = 4):
    k = len(explained_variance)
    xs = np.arange(1, k+1)
    fig, ax = plt.subplots(1, 2, figsize=(fig_w, fig_h))
    ax[0].plot(xs, explained_variance_ratio, marker='o')
    ax[0].set_xlabel("PC")
    ax[0].set_ylabel("Explained variance ratio")
    ax[0].set_title("Scree")
    ax[1].plot(xs, np.cumsum(explained_variance_ratio), marker='o')
    ax[1].set_xlabel("PC")
    ax[1].set_ylabel("Cumulative EVR")
    ax[1].set_title("Cumulative variance")
    ax[1].set_ylim(0,1.01)
    # plt.show()


def _labels_array_for_indices(labels, trial_indices):
    """
    Returns labels_for_plot: np.ndarray of shape [K, M] (M can be 1+),
    aligned to trial_indices order.
    Supports:
      - labels as np.ndarray [N, M]
      - labels as dict {trial_idx: scalar or 1D array-like}
    """
    if isinstance(labels, dict):
        rows = []
        for i in trial_indices:
            v = labels.get(i)
            if v is None:
                raise ValueError(f"labels dict missing trial {i}")
            a = np.atleast_1d(np.asarray(v))
            rows.append(a)
        # pad to same width if needed
        maxm = max(r.size for r in rows)
        rows2 = []
        for r in rows:
            if r.size < maxm:
                rr = np.zeros(maxm, dtype=float)
                rr[:r.size] = r
                rows2.append(rr)
            else:
                rows2.append(r.astype(float))
        return np.stack(rows2, axis=0)
    else:
        arr = np.asarray(labels)
        if arr.ndim == 1:
            arr = arr[:, None]
        # select only requested indices
        return arr[np.asarray(trial_indices)]

def _group_means_by_labels(trials, trial_indices, labels_for_plot):
    """
    Group selected trials by identical label rows. Within each group,
    truncate all member trials to the group's min length and average.

    Returns:
      mean_trials: list of [Tg, d]
      unique_labels: np.ndarray [G, M]
      group_sizes: np.ndarray [G]
    """
    # build groups
    key_list = [tuple(labels_for_plot[k].tolist()) for k in range(len(trial_indices))]
    groups = {}
    for pos, (trial_idx, key) in enumerate(zip(trial_indices, key_list)):
        X = trials[trial_idx]
        if X is None or np.asarray(X).size == 0:
            continue
        groups.setdefault(key, []).append(np.asarray(X))

    unique_keys = list(groups.keys())
    unique_labels = np.array(unique_keys, dtype=float) if len(unique_keys) and \
                    np.issubdtype(np.asarray(unique_keys[0], dtype=float).dtype, np.number) else \
                    np.array(unique_keys, dtype=object)

    mean_trials, sizes = [], []
    for key in unique_keys:
        seqs = [np.asarray(X) for X in groups[key] if np.asarray(X).size > 0]
        if not seqs:
            continue
        min_len = min(s.shape[0] for s in seqs)
        if min_len <= 0:
            continue
        stacked = np.stack([s[:min_len] for s in seqs], axis=0)  # [n_g, T, d]
        mean_trials.append(stacked.mean(axis=0))
        sizes.append(stacked.shape[0])
    return mean_trials, np.array(unique_keys), np.asarray(sizes, int)


## 1.3 2D projection
def plot_trial_trajectories_gradient_steps(
    transform,
    trials,
    trial_indices,
    comps=(0, 1),
    evr=None,
    start_color="#fdae61",
    end_color="#313695",
    linewidth=2.0,
    start_marker_size=36,
    alpha=0.5,
    start_marker_alpha=0.8,
    end_arrow_alpha=0.8,
    title=None,
    fig_w=6,
    fig_h=6,
    labels=None,
    circmap="twilight",
    vrange=None,
    cbar_label=None,
    # NEW:
    plot_mean=False
):
    # --- NEW: optionally aggregate by labels over the selected trials
    if plot_mean:
        if labels is None:
            raise ValueError("plot_mean=True requires labels.")
        lab_arr = _labels_array_for_indices(labels, trial_indices)  # [K, M]
        mean_trials, uniq_labels, _sizes = _group_means_by_labels(trials, trial_indices, lab_arr)
        # Replace trials/indexing with group means
        trials_plot = mean_trials
        trial_indices_plot = list(range(len(mean_trials)))
        # For coloring:
        labels_for_plot = uniq_labels
    else:
        trials_plot = trials
        trial_indices_plot = trial_indices
        labels_for_plot = None  # will be set from `labels` below if provided

    # 1) Project selected trials
    proj, max_len = [], 0
    for idx in trial_indices_plot:
        S = next(transform([trials_plot[idx]]))
        if S.size == 0: 
            continue
        proj.append((idx, S))
        max_len = max(max_len, S.shape[0])
    if not proj: raise ValueError("No non-empty trials to plot.")

    # 2) Colormap / labeling
    use_labels = (labels is not None) or (labels_for_plot is not None)
    # decide label source per plotted item
    if use_labels:
        if labels_for_plot is None:
            arr_all = _labels_array_for_indices(labels, trial_indices)  # original selection
            # align to plotted order (not mean)
            labels_for_plot = arr_all
        # reduce to one row per plotted index
        if plot_mean:
            lab_rows = np.asarray(labels_for_plot)                # [G, M]
        else:
            lab_rows = np.asarray(labels_for_plot)                # [K, M]
        # numeric-scalar vs categorical
        numeric_scalar = (lab_rows.ndim == 2) and (lab_rows.shape[1] == 1) and np.issubdtype(lab_rows.dtype, np.number)
        if numeric_scalar:
            vals = lab_rows[:, 0].astype(float)
            if vrange is None:
                vmin, vmax = float(np.nanmin(vals)), float(np.nanmax(vals))
                if (vmin >= -180-1e-6) and (vmax <= 180+1e-6):
                    vmin, vmax = -180.0, 180.0
            else:
                vmin, vmax = vrange
            cmap = plt.get_cmap(circmap)
            norm = Normalize(vmin=vmin, vmax=vmax)
            def color_for(pos):
                return cmap(norm(float(vals[pos])))
        else:
            # categorical by tuple of label values
            tuples = [tuple(r.tolist()) for r in lab_rows]
            cats = list(dict.fromkeys(tuples))  # stable unique
            idx_of = {c:i for i,c in enumerate(cats)}
            K = max(1, len(cats))
            base = plt.get_cmap("tab20" if K <= 20 else "hsv", K)
            from matplotlib.colors import ListedColormap, BoundaryNorm
            cmap = ListedColormap([base(i) for i in range(K)])
            norm = BoundaryNorm(np.arange(K+1)-0.5, K)
            def color_for(pos):
                return cmap(norm(idx_of[tuples[pos]]))
    else:
        cmap = LinearSegmentedColormap.from_list("o2b", [start_color, end_color])
        norm = Normalize(vmin=0, vmax=max(0, max_len-1))

    # 3) Figure
    fig, ax = plt.subplots(figsize=(fig_w, fig_h))

    for plot_pos, (idx, S) in enumerate(proj):
        x, y = S[:, comps[0]], S[:, comps[1]]

        if not use_labels:
            if S.shape[0] == 1:
                ax.scatter(x[0], y[0], s=start_marker_size, facecolor=cmap(norm(0)),
                           edgecolor="black", linewidths=0.5, zorder=3)
                continue
            pts = np.column_stack([x, y])
            segs = np.stack([pts[:-1], pts[1:]], axis=1)
            seg_steps = np.arange(segs.shape[0])
            lc = LineCollection(segs, cmap=cmap, norm=norm, linewidths=linewidth, alpha=alpha)
            lc.set_array(seg_steps)
            ax.add_collection(lc)
            ax.scatter(x[0], y[0], s=start_marker_size, facecolor=cmap(norm(0)),
                       edgecolor="black", linewidths=0.5, alpha=start_marker_alpha, zorder=3)
            end_color_i = cmap(norm(S.shape[0]-1))
        else:
            col = color_for(plot_pos)
            ax.plot(x, y, lw=linewidth, alpha=alpha, color=col)
            ax.scatter(x[0], y[0], s=start_marker_size, facecolor=col,
                       edgecolor="black", linewidths=0.5, alpha=start_marker_alpha, zorder=3)
            end_color_i = col

        if S.shape[0] > 1:
            ax.annotate("", xy=(x[-1], y[-1]), xytext=(x[-2], y[-2]),
                        arrowprops=dict(arrowstyle="->", color=end_color_i,
                                        lw=linewidth*1.2, shrinkA=0, shrinkB=0,
                                        alpha=end_arrow_alpha),
                        zorder=4)

    # 4) Labels
    xlab = f"PC{comps[0]+1}"; ylab = f"PC{comps[1]+1}"
    if evr is not None:
        try:
            xlab += f" ({evr[comps[0]]*100:.1f}% EV)"
            ylab += f" ({evr[comps[1]]*100:.1f}% EV)"
        except Exception:
            pass
    ax.set_xlabel(xlab); ax.set_ylabel(ylab)
    ttl = "PC trajectories"
    if plot_mean: ttl += " (group means)"
    ax.set_title(title or ttl + " (color = " + ("label" if use_labels else "step index") + ")")
    ax.axhline(0, lw=0.5, color="k", alpha=0.3); ax.axvline(0, lw=0.5, color="k", alpha=0.3)
    ax.set_aspect("equal", adjustable="datalim")

    # 5) Colorbar
    sm = plt.cm.ScalarMappable(cmap=cmap, norm=norm); sm.set_array([])
    cbar = fig.colorbar(sm, ax=ax, pad=0.015)
    if not use_labels:
        cbar.set_label("Step index")
        ticks = [0, (max_len - 1)//2, max_len - 1] if max_len > 2 else [0, max_len - 1]
        cbar.set_ticks(ticks); cbar.set_ticklabels([str(t) for t in ticks])
    else:
        # numeric-scalar vs categorical
        if 'numeric_scalar' in locals() and numeric_scalar:
            lbl = cbar_label or ("Angle (deg)" if np.isclose(norm.vmin,-180) and np.isclose(norm.vmax,180) else "Label")
            cbar.set_label(lbl)
            if lbl == "Angle (deg)": cbar.set_ticks([-180, -90, 0, 90, 180])
        else:
            cbar.set_label("Group")
    plt.tight_layout(); # plt.show()

# def plot_trial_trajectories_gradient_steps(
#     transform,
#     trials,
#     trial_indices,
#     comps=(0, 1),
#     evr=None,
#     start_color="#fdae61",
#     end_color="#313695",
#     linewidth=2.0,
#     start_marker_size=36,
#     alpha=0.5,
#     start_marker_alpha=0.8,
#     end_arrow_alpha=0.8,
#     title=None,
#     fig_w=6,
#     fig_h=6,
#     # NEW:
#     labels=None,               # per-trial numeric labels (e.g., theta_deg)
#     circmap="twilight",        # cyclic cmap when labels are provided
#     vrange=None,               # (vmin, vmax) for labels; default auto
#     cbar_label=None            # override colorbar label
# ):
#     # 1) Project selected trials
#     proj, max_len = [], 0
#     for idx in trial_indices:
#         S = next(transform([trials[idx]]))
#         if S.size == 0: 
#             continue
#         proj.append((idx, S))
#         max_len = max(max_len, S.shape[0])
#     if not proj: raise ValueError("No non-empty trials to plot.")

#     # 2) Colormap
#     use_labels = labels is not None
#     if not use_labels:
#         cmap = LinearSegmentedColormap.from_list("o2b", [start_color, end_color])
#         norm = Normalize(vmin=0, vmax=max_len - 1)
#     else:
#         # per-trial label lookup
#         if isinstance(labels, dict):
#             lab_for = {i: labels.get(i) for i, _ in proj}
#         else:
#             arr = np.asarray(labels)
#             lab_for = {i: float(arr[i]) for i, _ in proj}
#         vals = np.array([lab_for[i] for i, _ in proj], dtype=float)
#         if vrange is None:
#             vmin, vmax = float(np.nanmin(vals)), float(np.nanmax(vals))
#             if (vmin >= -180-1e-6) and (vmax <= 180+1e-6):
#                 vmin, vmax = -180.0, 180.0
#         else:
#             vmin, vmax = vrange
#         cmap = plt.get_cmap(circmap)
#         norm = Normalize(vmin=vmin, vmax=vmax)

#     # 3) Figure
#     fig, ax = plt.subplots(figsize=(fig_w, fig_h))

#     for idx, S in proj:
#         x, y = S[:, comps[0]], S[:, comps[1]]

#         if not use_labels:
#             if S.shape[0] == 1:
#                 ax.scatter(x[0], y[0], s=start_marker_size, facecolor=cmap(norm(0)),
#                            edgecolor="black", linewidths=0.5, zorder=3)
#                 continue
#             pts = np.column_stack([x, y])
#             segs = np.stack([pts[:-1], pts[1:]], axis=1)
#             seg_steps = np.arange(segs.shape[0])
#             lc = LineCollection(segs, cmap=cmap, norm=norm, linewidths=linewidth, alpha=alpha)
#             lc.set_array(seg_steps)
#             ax.add_collection(lc)
#             # start + end (arrow)
#             ax.scatter(x[0], y[0], s=start_marker_size, facecolor=cmap(norm(0)),
#                        edgecolor="black", linewidths=0.5, alpha=start_marker_alpha, zorder=3)
#             end_color_i = cmap(norm(S.shape[0]-1))
#         else:
#             # solid color per trajectory based on label
#             color = cmap(norm(float(lab_for[idx])))
#             ax.plot(x, y, lw=linewidth, alpha=alpha, color=color)
#             ax.scatter(x[0], y[0], s=start_marker_size, facecolor=color,
#                        edgecolor="black", linewidths=0.5, alpha=start_marker_alpha, zorder=3)
#             end_color_i = color

#         if S.shape[0] > 1:
#             ax.annotate("", xy=(x[-1], y[-1]), xytext=(x[-2], y[-2]),
#                         arrowprops=dict(arrowstyle="->", color=end_color_i,
#                                         lw=linewidth*1.2, shrinkA=0, shrinkB=0,
#                                         alpha=end_arrow_alpha),
#                         zorder=4)

#     # 4) Labels
#     xlab = f"PC{comps[0]+1}"; ylab = f"PC{comps[1]+1}"
#     if evr is not None:
#         try:
#             xlab += f" ({evr[comps[0]]*100:.1f}% EV)"
#             ylab += f" ({evr[comps[1]]*100:.1f}% EV)"
#         except Exception:
#             pass
#     ax.set_xlabel(xlab); ax.set_ylabel(ylab)
#     ax.set_title(title or ("PC trajectories (color = " + ("label" if use_labels else "step index") + ")"))
#     ax.axhline(0, lw=0.5, color="k", alpha=0.3); ax.axvline(0, lw=0.5, color="k", alpha=0.3)
#     ax.set_aspect("equal", adjustable="datalim")

#     # 5) Colorbar
#     sm = plt.cm.ScalarMappable(cmap=cmap, norm=norm); sm.set_array([])
#     cbar = fig.colorbar(sm, ax=ax, pad=0.015)
#     if not use_labels:
#         cbar.set_label("Step index")
#         ticks = [0, (max_len - 1)//2, max_len - 1] if max_len > 2 else [0, max_len - 1]
#         cbar.set_ticks(ticks); cbar.set_ticklabels([str(t) for t in ticks])
#     else:
#         lbl = cbar_label or "Angle (deg)" if np.isclose(norm.vmin,-180) and np.isclose(norm.vmax,180) else (cbar_label or "Label")
#         cbar.set_label(lbl)
#         if lbl == "Angle (deg)": cbar.set_ticks([-180, -90, 0, 90, 180])

#     plt.tight_layout(); plt.show()


## 1.4 3D projection
def plot_trial_trajectories_gradient_steps_3d(
    transform,
    trials,
    trial_indices,
    comps=(0, 1, 2),
    evr=None,
    start_color="#fdae61",
    end_color="#313695",
    linewidth=2.0,
    alpha=0.5,
    start_marker_size=36,
    start_marker_alpha=0.95,
    end_arrow_alpha=0.95,
    arrow_head_frac=0.03,
    arrow_length_ratio=0.6,
    title=None,
    zlabel_on_left=True,
    fig_w=6,
    fig_h=6,
    labels=None,
    circmap="twilight",
    vrange=None,
    cbar_label=None,
    # NEW:
    plot_mean=False
):
    # NEW aggregation
    if plot_mean:
        if labels is None:
            raise ValueError("plot_mean=True requires labels.")
        lab_arr = _labels_array_for_indices(labels, trial_indices)
        mean_trials, uniq_labels, _sizes = _group_means_by_labels(trials, trial_indices, lab_arr)
        trials_plot = mean_trials
        trial_indices_plot = list(range(len(mean_trials)))
        labels_for_plot = uniq_labels
    else:
        trials_plot = trials
        trial_indices_plot = trial_indices
        labels_for_plot = None

    proj, max_len = [], 0
    for idx in trial_indices_plot:
        S = next(transform([trials_plot[idx]]))
        if S.size == 0: continue
        proj.append((idx, S)); max_len = max(max_len, S.shape[0])
    if not proj: raise ValueError("No non-empty trials to plot.")

    use_labels = (labels is not None) or (labels_for_plot is not None)
    if use_labels:
        if labels_for_plot is None:
            labels_for_plot = _labels_array_for_indices(labels, trial_indices)
        lab_rows = np.asarray(labels_for_plot)
        if lab_rows.ndim == 1:
            lab_rows = lab_rows[:, None]
        numeric_scalar = (lab_rows.shape[1] == 1) and np.issubdtype(lab_rows.dtype, np.number)
        if numeric_scalar:
            vals = lab_rows[:, 0].astype(float)
            if vrange is None:
                vmin, vmax = float(np.nanmin(vals)), float(np.nanmax(vals))
                if (vmin >= -180-1e-6) and (vmax <= 180+1e-6): vmin, vmax = -180.0, 180.0
            else:
                vmin, vmax = vrange
            cmap = plt.get_cmap(circmap)
            norm = Normalize(vmin=vmin, vmax=vmax)
            def color_for(pos): return cmap(norm(float(vals[pos])))
        else:
            tuples = [tuple(r.tolist()) for r in lab_rows]
            cats = list(dict.fromkeys(tuples))
            idx_of = {c:i for i,c in enumerate(cats)}
            K = max(1, len(cats))
            base = plt.get_cmap("tab20" if K <= 20 else "hsv", K)
            from matplotlib.colors import ListedColormap, BoundaryNorm
            cmap = ListedColormap([base(i) for i in range(K)])
            norm = BoundaryNorm(np.arange(K+1)-0.5, K)
            def color_for(pos): return cmap(norm(idx_of[tuples[pos]]))
    else:
        cmap = LinearSegmentedColormap.from_list("o2b", [start_color, end_color])
        norm = Normalize(vmin=0, vmax=(1 if max_len <= 1 else max_len - 1))

    fig = plt.figure(figsize=(fig_w, fig_h))
    fig.subplots_adjust(left=0.12, right=0.86, bottom=0.08, top=0.92)
    ax = fig.add_subplot(111, projection="3d")

    for plot_pos, (idx, S) in enumerate(proj):
        x, y, z = S[:, comps[0]], S[:, comps[1]], S[:, comps[2]]
        if not use_labels:
            pts = np.column_stack([x, y, z])
            segs = np.stack([pts[:-1], pts[1:]], axis=1)
            seg_steps = np.arange(segs.shape[0])
            colors = cmap(norm(seg_steps))
            lc = Line3DCollection(segs, linewidths=linewidth, alpha=alpha); lc.set_colors(colors)
            ax.add_collection3d(lc)
            start_col = cmap(norm(0)); end_col = cmap(norm(S.shape[0]-1))
        else:
            col = color_for(plot_pos)
            ax.plot(x, y, z, lw=linewidth, alpha=alpha, color=col)
            start_col = end_col = col

        ax.scatter(x[0], y[0], z[0], s=start_marker_size, marker="o",
                   facecolor=start_col, edgecolor="black",
                   linewidths=0.5, alpha=start_marker_alpha, zorder=3)

        if S.shape[0] > 1:
            dx, dy, dz = x[-1]-x[-2], y[-1]-y[-2], z[-1]-z[-2]
            rng = np.array([x.max()-x.min(), y.max()-y.min(), z.max()-z.min()])
            diag = float(np.linalg.norm(rng))
            L = max(1e-12, arrow_head_frac * (diag if diag > 0 else 1.0))
            u = np.array([dx, dy, dz], float); n = float(np.linalg.norm(u))
            if n < 1e-12:
                ax.scatter(x[-1], y[-1], z[-1], s=start_marker_size*0.9, marker="^",
                           facecolor=end_col, edgecolor="black", linewidths=0.5,
                           alpha=end_arrow_alpha, zorder=10)
            else:
                u_hat = u / max(n, 1e-12)
                tail = np.array([x[-1], y[-1], z[-1]]) - u_hat * L
                ax.quiver(tail[0], tail[1], tail[2],
                          u_hat[0], u_hat[1], u_hat[2],
                          length=L, normalize=False, color=end_col,
                          linewidth=linewidth*1.2, arrow_length_ratio=arrow_length_ratio,
                          alpha=end_arrow_alpha, zorder=10, clip_on=False)

    # labels/title
    def lab(i):
        base = f"PC{comps[i]+1}"
        if evr is not None and len(evr) > comps[i]: base += f" ({evr[comps[i]]*100:.1f}% EV)"
        return base
    ax.set_xlabel(lab(0)); ax.set_ylabel(lab(1))
    z_text = lab(2)
    if zlabel_on_left: ax.set_zlabel(""); fig.text(0.035, 0.52, z_text, rotation=90, va="center", ha="center")
    else: ax.set_zlabel(z_text, labelpad=12)
    ttl = "3D PC trajectories"
    if plot_mean: ttl += " (group means)"
    ax.set_title(title or (ttl + " (color = " + ("label" if use_labels else "step index") + ")"))
    ax.set_box_aspect((1,1,1))

    # colorbar
    cax = fig.add_axes([0.88, 0.18, 0.03, 0.64])
    sm = plt.cm.ScalarMappable(cmap=cmap, norm=norm); sm.set_array([])
    cbar = fig.colorbar(sm, cax=cax)
    if not use_labels:
        cbar.set_label("Step index")
        ticks = [0, (max_len - 1)//2, max_len - 1] if max_len > 2 else [0, max_len - 1]
        cbar.set_ticks(ticks); cbar.set_ticklabels([str(t) for t in ticks])
    else:
        if 'numeric_scalar' in locals() and numeric_scalar:
            lbl = cbar_label or ("Angle (deg)" if np.isclose(norm.vmin,-180) and np.isclose(norm.vmax,180) else "Label")
            cbar.set_label(lbl)
            if lbl == "Angle (deg)": cbar.set_ticks([-180, -90, 0, 90, 180])
        else:
            cbar.set_label("Group")
    ax.autoscale_view(); # plt.show()


# def plot_trial_trajectories_gradient_steps_3d(
#     transform,
#     trials,
#     trial_indices,
#     comps=(0, 1, 2),
#     evr=None,
#     start_color="#fdae61",
#     end_color="#313695",
#     linewidth=2.0,
#     alpha=0.5,
#     start_marker_size=36,
#     start_marker_alpha=0.95,
#     end_arrow_alpha=0.95,
#     arrow_head_frac=0.03,
#     arrow_length_ratio=0.6,
#     title=None,
#     zlabel_on_left=True,
#     fig_w=6,
#     fig_h=6,
#     # NEW:
#     labels=None,
#     circmap="twilight",
#     vrange=None,
#     cbar_label=None
# ):
#     proj, max_len = [], 0
#     for idx in trial_indices:
#         S = next(transform([trials[idx]]))
#         if S.size == 0: continue
#         proj.append((idx, S)); max_len = max(max_len, S.shape[0])
#     if not proj: raise ValueError("No non-empty trials to plot.")

#     use_labels = labels is not None
#     if not use_labels:
#         cmap = LinearSegmentedColormap.from_list("o2b", [start_color, end_color])
#         norm = Normalize(vmin=0, vmax=(1 if max_len <= 1 else max_len - 1))
#     else:
#         if isinstance(labels, dict):
#             lab_for = {i: labels.get(i) for i, _ in proj}
#         else:
#             arr = np.asarray(labels); lab_for = {i: float(arr[i]) for i,_ in proj}
#         vals = np.array([lab_for[i] for i,_ in proj], float)
#         if vrange is None:
#             vmin, vmax = float(np.nanmin(vals)), float(np.nanmax(vals))
#             if (vmin >= -180-1e-6) and (vmax <= 180+1e-6): vmin, vmax = -180.0, 180.0
#         else:
#             vmin, vmax = vrange
#         cmap = plt.get_cmap(circmap)
#         norm = Normalize(vmin=vmin, vmax=vmax)

#     fig = plt.figure(figsize=(fig_w, fig_h))
#     fig.subplots_adjust(left=0.12, right=0.86, bottom=0.08, top=0.92)
#     ax = fig.add_subplot(111, projection="3d")

#     for idx, S in proj:
#         x, y, z = S[:, comps[0]], S[:, comps[1]], S[:, comps[2]]
#         if not use_labels:
#             pts = np.column_stack([x, y, z])
#             segs = np.stack([pts[:-1], pts[1:]], axis=1)
#             seg_steps = np.arange(segs.shape[0])
#             colors = cmap(norm(seg_steps))
#             lc = Line3DCollection(segs, linewidths=linewidth, alpha=alpha); lc.set_colors(colors)
#             ax.add_collection3d(lc)
#             start_col = cmap(norm(0)); end_col = cmap(norm(S.shape[0]-1))
#         else:
#             col = cmap(norm(float(lab_for[idx])))
#             ax.plot(x, y, z, lw=linewidth, alpha=alpha, color=col)
#             start_col = end_col = col

#         ax.scatter(x[0], y[0], z[0], s=start_marker_size, marker="o",
#                    facecolor=start_col, edgecolor="black",
#                    linewidths=0.5, alpha=start_marker_alpha, zorder=3)

#         if S.shape[0] > 1:
#             dx, dy, dz = x[-1]-x[-2], y[-1]-y[-2], z[-1]-z[-2]
#             rng = np.array([x.max()-x.min(), y.max()-y.min(), z.max()-z.min()])
#             diag = float(np.linalg.norm(rng))
#             L = max(1e-12, arrow_head_frac * (diag if diag > 0 else 1.0))
#             u = np.array([dx, dy, dz], float); n = float(np.linalg.norm(u))
#             if n < 1e-12:
#                 ax.scatter(x[-1], y[-1], z[-1], s=start_marker_size*0.9, marker="^",
#                            facecolor=end_col, edgecolor="black", linewidths=0.5,
#                            alpha=end_arrow_alpha, zorder=10)
#             else:
#                 u_hat = u / max(n, 1e-12)
#                 tail = np.array([x[-1], y[-1], z[-1]]) - u_hat * L
#                 ax.quiver(tail[0], tail[1], tail[2],
#                           u_hat[0], u_hat[1], u_hat[2],
#                           length=L, normalize=False, color=end_col,
#                           linewidth=linewidth*1.2, arrow_length_ratio=arrow_length_ratio,
#                           alpha=end_arrow_alpha, zorder=10, clip_on=False)

#     # axis labels + title (unchanged) ...
#     def lab(i):
#         base = f"PC{comps[i]+1}"
#         if evr is not None and len(evr) > comps[i]: base += f" ({evr[comps[i]]*100:.1f}% EV)"
#         return base
#     ax.set_xlabel(lab(0)); ax.set_ylabel(lab(1))
#     z_text = lab(2)
#     if zlabel_on_left: ax.set_zlabel(""); fig.text(0.035, 0.52, z_text, rotation=90, va="center", ha="center")
#     else: ax.set_zlabel(z_text, labelpad=12)
#     ax.set_title(title or ("3D PC trajectories (color = " + ("label" if use_labels else "step index") + ")"))
#     ax.set_box_aspect((1,1,1))

#     # colorbar
#     cax = fig.add_axes([0.88, 0.18, 0.03, 0.64])
#     sm = plt.cm.ScalarMappable(cmap=cmap, norm=norm); sm.set_array([])
#     cbar = fig.colorbar(sm, cax=cax)
#     if not use_labels:
#         cbar.set_label("Step index")
#         ticks = [0, (max_len - 1)//2, max_len - 1] if max_len > 2 else [0, max_len - 1]
#         cbar.set_ticks(ticks); cbar.set_ticklabels([str(t) for t in ticks])
#     else:
#         lbl = cbar_label or "Angle (deg)" if np.isclose(norm.vmin,-180) and np.isclose(norm.vmax,180) else (cbar_label or "Label")
#         cbar.set_label(lbl)
#         if lbl == "Angle (deg)": cbar.set_ticks([-180, -90, 0, 90, 180])

#     ax.autoscale_view(); plt.show()


## 1.4 trajectory for each PC
def plot_pc_time_series(
    transform,
    trials,
    pcs=(0,),
    trial_indices=None,
    labels=None,
    evr=None,
    dt=None,
    linewidth=1.2,
    alpha=0.9,
    fig_w=8,
    fig_h_per=2,
    circmap="twilight",
    vrange=None,
    cbar_label=None,
    # NEW:
    plot_mean=False
):
    if trial_indices is None:
        trial_indices = list(range(len(trials)))

    # NEW aggregation
    if plot_mean:
        if labels is None:
            raise ValueError("plot_mean=True requires labels.")
        lab_arr = _labels_array_for_indices(labels, trial_indices)
        mean_trials, uniq_labels, _sizes = _group_means_by_labels(trials, trial_indices, lab_arr)
        trials_plot = mean_trials
        trial_indices_plot = list(range(len(mean_trials)))
        labels_for_plot = uniq_labels
    else:
        trials_plot = trials
        trial_indices_plot = trial_indices
        labels_for_plot = None

    pcs = (pcs,) if isinstance(pcs, (int, np.integer)) else list(pcs)

    proj = {}
    for idx in trial_indices_plot:
        S = next(transform([trials_plot[idx]]))
        if S.size > 0: proj[idx] = S
    if not proj: raise ValueError("No non-empty trials to plot.")

    # coloring (continuous scalar vs categorical)
    use_labels = (labels is not None) or (labels_for_plot is not None)
    use_cont = False
    if use_labels:
        if labels_for_plot is None:
            labels_for_plot = _labels_array_for_indices(labels, trial_indices)
        lab_rows = np.asarray(labels_for_plot)
        if lab_rows.ndim == 1:
            lab_rows = lab_rows[:, None]
        if (lab_rows.shape[1] == 1) and np.issubdtype(lab_rows.dtype, np.number):
            use_cont = True
            vals = lab_rows[:, 0].astype(float)
            if vrange is None:
                vmin, vmax = float(np.nanmin(vals)), float(np.nanmax(vals))
                if (vmin >= -180-1e-6) and (vmax <= 180+1e-6): vmin, vmax = -180.0, 180.0
            else:
                vmin, vmax = vrange
            cmap = plt.get_cmap(circmap); norm = Normalize(vmin=vmin, vmax=vmax)
            def get_color_by_pos(pos): return cmap(norm(float(vals[pos])))
        else:
            # categorical
            tuples = [tuple(r.tolist()) for r in lab_rows]
            cats = list(dict.fromkeys(tuples))
            cat_to_i = {c:i for i,c in enumerate(cats)}
            K = max(1, len(cats))
            hsv = plt.get_cmap("hsv", K)
            from matplotlib.colors import ListedColormap, BoundaryNorm
            cmap = ListedColormap([hsv(i) for i in range(K)])
            norm = BoundaryNorm(np.arange(K+1)-0.5, K)
            def get_color_by_pos(pos): return cmap(norm(cat_to_i[tuples[pos]]))
    else:
        def get_color_by_pos(pos): return "k"

    fig_h = fig_h_per * len(pcs)
    fig, axes = plt.subplots(len(pcs), 1, figsize=(fig_w, fig_h), squeeze=False)
    axes = axes.ravel()

    for ax, pc in zip(axes, pcs):
        for plot_pos, idx in enumerate(trial_indices_plot):
            if idx not in proj: continue
            y = proj[idx][:, pc]
            x = np.arange(len(y)) if dt is None else np.arange(len(y)) * dt
            ax.plot(x, y, lw=linewidth, alpha=alpha, color=get_color_by_pos(plot_pos))

        ylab = f"PC{pc+1}"
        if evr is not None and pc < len(evr): ylab += f" ({evr[pc]*100:.1f}% EV)"
        ax.set_ylabel(ylab); ax.grid(True, alpha=0.25)
    axes[-1].set_xlabel("Time (step)" if dt is None else "Time")

    # colorbar
    if use_labels:
        fig.subplots_adjust(bottom=0.18)
        cax = fig.add_axes([0.12, 0.08, 0.76, 0.04])
        sm = plt.cm.ScalarMappable(norm=norm, cmap=cmap); sm.set_array([])
        cbar = fig.colorbar(sm, cax=cax, orientation="horizontal")
        if use_cont:
            lbl = cbar_label or ("Angle (deg)" if np.isclose(norm.vmin,-180) and np.isclose(norm.vmax,180) else "Label")
            cbar.set_label(lbl)
            if lbl == "Angle (deg)": cbar.set_ticks([-180, -90, 0, 90, 180])
        else:
            cbar.set_label("Group")

    plt.tight_layout(rect=[0.02, 0.18 if use_labels else 0.02, 0.98, 0.98])
    # plt.show()


# def plot_pc_time_series(
#     transform,
#     trials,
#     pcs=(0,),
#     trial_indices=None,
#     labels=None,           # None | numeric array/dict (continuous) | non-numeric (categorical)
#     evr=None,
#     dt=None,
#     linewidth=1.2,
#     alpha=0.9,
#     fig_w=8,
#     fig_h_per=2,
#     # NEW:
#     circmap="twilight",
#     vrange=None,
#     cbar_label=None
# ):
#     if trial_indices is None:
#         trial_indices = list(range(len(trials)))
#     pcs = (pcs,) if isinstance(pcs, (int, np.integer)) else list(pcs)

#     proj = {}
#     for idx in trial_indices:
#         S = next(transform([trials[idx]]))
#         if S.size > 0: proj[idx] = S
#     if not proj: raise ValueError("No non-empty trials to plot.")

#     use_labels = labels is not None
#     use_cont = False
#     if use_labels:
#         if isinstance(labels, dict):
#             lab_for_all = {i: labels.get(i) for i in trial_indices}
#         else:
#             arr = np.asarray(labels, dtype=object)
#             lab_for_all = {i: arr[i] for i in trial_indices}
#         # decide continuous vs categorical
#         vals = [lab_for_all[i] for i in trial_indices if i in proj]
#         use_cont = all(isinstance(v, (int, float, np.integer, np.floating)) for v in vals)

#         if use_cont:
#             vals = np.array(vals, float)
#             if vrange is None:
#                 vmin, vmax = float(np.nanmin(vals)), float(np.nanmax(vals))
#                 if (vmin >= -180-1e-6) and (vmax <= 180+1e-6): vmin, vmax = -180.0, 180.0
#             else:
#                 vmin, vmax = vrange
#             cmap = plt.get_cmap(circmap); norm = Normalize(vmin=vmin, vmax=vmax)
#             get_color = lambda i: cmap(norm(float(lab_for_all[i])))
#         else:
#             # categorical path (existing style)
#             cats, order = [], []
#             for i in trial_indices:
#                 if i in proj:
#                     v = lab_for_all[i]
#                     if v not in cats: cats.append(v)
#             K = max(1, len(cats))
#             hsv = plt.get_cmap("hsv", K)
#             cmap = ListedColormap([hsv(i) for i in range(K)])
#             norm = BoundaryNorm(np.arange(K+1)-0.5, K)
#             cat_to_i = {c:i for i,c in enumerate(cats)}
#             get_color = lambda i: cmap(norm(cat_to_i[lab_for_all[i]]))
#     else:
#         get_color = lambda i: "k"

#     fig_h = fig_h_per * len(pcs)
#     fig, axes = plt.subplots(len(pcs), 1, figsize=(fig_w, fig_h), squeeze=False)
#     axes = axes.ravel()

#     for ax, pc in zip(axes, pcs):
#         for idx in trial_indices:
#             if idx not in proj: continue
#             y = proj[idx][:, pc]
#             x = np.arange(len(y)) if dt is None else np.arange(len(y)) * dt
#             ax.plot(x, y, lw=linewidth, alpha=alpha, color=get_color(idx))

#         ylab = f"PC{pc+1}"
#         if evr is not None and pc < len(evr): ylab += f" ({evr[pc]*100:.1f}% EV)"
#         ax.set_ylabel(ylab); ax.grid(True, alpha=0.25)
#     axes[-1].set_xlabel("Time (step)" if dt is None else "Time")

#     # colorbar
#     if use_labels:
#         if use_cont:
#             fig.subplots_adjust(bottom=0.18)
#             cax = fig.add_axes([0.12, 0.08, 0.76, 0.04])
#             sm = plt.cm.ScalarMappable(norm=norm, cmap=cmap); sm.set_array([])
#             cbar = fig.colorbar(sm, cax=cax, orientation="horizontal")
#             lbl = cbar_label or ("Angle (deg)" if np.isclose(norm.vmin,-180) and np.isclose(norm.vmax,180) else "Label")
#             cbar.set_label(lbl)
#             if lbl == "Angle (deg)": cbar.set_ticks([-180, -90, 0, 90, 180])
#         else:
#             fig.subplots_adjust(bottom=0.18)
#             cax = fig.add_axes([0.12, 0.08, 0.76, 0.04])
#             sm = plt.cm.ScalarMappable(norm=norm, cmap=cmap); sm.set_array([])
#             cbar = fig.colorbar(sm, cax=cax, orientation="horizontal")
#             cbar.set_ticks(range(len(cat_to_i)))
#             cbar.set_ticklabels(list(cat_to_i.keys()))
#             cbar.set_label("Trial category")

#     plt.tight_layout(rect=[0.02, 0.18 if use_labels else 0.02, 0.98, 0.98])
#     plt.show()


def plot_2d_hand_trajectories(
    hand_pos_sel,                 # list of arrays [T_k, 2]
    trial_indices=None,           # None => plot all trials
    trial_type_use=None,          # labels for *plotted* trials (same length as trial_indices or all)
    linewidth=1.6,
    alpha=0.9,
    fig_w=6,
    fig_h=6,
    title="2D hand trajectories",
):
    # ---- decide which trials to plot ----
    if trial_indices is None:
        idx_list = list(range(len(hand_pos_sel)))
    else:
        idx_list = list(trial_indices)

    # ---- prepare categorical coloring (optional) ----
    use_cats = trial_type_use is not None
    if use_cats:
        trial_type_use = list(trial_type_use)
        if len(trial_type_use) != len(idx_list):
            raise ValueError("trial_type_use must have the same length as the number of plotted trials.")
        # preserve first-seen category order
        cats = []
        for lab in trial_type_use:
            if lab not in cats:
                cats.append(lab)
        K = max(1, len(cats))
        base = plt.get_cmap("tab20", K) if K <= 20 else plt.get_cmap("hsv", K)
        cmap = ListedColormap([base(i) for i in range(K)])
        norm = BoundaryNorm(np.arange(K+1) - 0.5, K)
        cat_to_i = {c: i for i, c in enumerate(cats)}

    # ---- plot ----
    fig, ax = plt.subplots(figsize=(fig_w, fig_h))
    for j, idx in enumerate(idx_list):
        arr = hand_pos_sel[idx]
        if arr is None or len(arr) == 0:
            continue
        if use_cats:
            ci = cat_to_i[trial_type_use[j]]
            color = cmap(norm(ci))
        else:
            color = "k"
        ax.plot(arr[:, 0], arr[:, 1], lw=linewidth, alpha=alpha, color=color)

    ax.set_aspect("equal", adjustable="datalim")
    ax.set_xlabel("X"); ax.set_ylabel("Y"); ax.grid(True, alpha=0.25)
    ax.set_title(title)

    # ---- bottom categorical colorbar (only if labels provided) ----
    if use_cats:
        fig.subplots_adjust(bottom=0.2)                 # reserve space
        cax = fig.add_axes([0.12, 0.08, 0.76, 0.05])    # [left, bottom, width, height]
        sm = plt.cm.ScalarMappable(norm=norm, cmap=cmap); sm.set_array([])
        cbar = fig.colorbar(sm, cax=cax, orientation="horizontal")
        cbar.set_ticks(range(len(cats)))
        cbar.set_ticklabels([str(c) for c in cats])
        cbar.set_label("Trial type")

    # plt.show()
```
