## Supplementary material for "Dynamic Compression Flows for Neuroscience Data": Python code: plot_sim_results_img_v7.py

```
import numpy as np
import matplotlib.pyplot as plt
from matplotlib.colors import ListedColormap, BoundaryNorm, Normalize, LinearSegmentedColormap
import torch
import dnnlib.util_v7 as util
from tqdm import tqdm
import torch.nn.functional as F
from torch import nn
from PIL import Image

import warnings
warnings.filterwarnings('ignore', 'Grad strides do not match bucket view strides') # False warning
device = torch.device("cuda" if torch.cuda.is_available() else "cpu")
torch.set_default_device(device)
colorgrad = "PuOr"

def _crop_or_resize4d(x, target_h, target_w):
    Hx, Wx = x.shape[-2], x.shape[-1]
    if Hx == target_h and Wx == target_w:
        return x
    if Hx >= target_h and Wx >= target_w:
        top  = (Hx - target_h) // 2
        left = (Wx - target_w) // 2
        return x[:, :, top:top+target_h, left:left+target_w]
    return F.interpolate(x, size=(target_h, target_w), mode='bilinear', align_corners=False)

class _FlowFlatAdapter(torch.nn.Module):
    def __init__(self, base, C, H, W):
        super().__init__()
        self.base, self.C, self.H, self.W = base, C, H, W
    def forward(self, x, t):
        # x: (B, D) -> (B, C, H, W)
        B = x.shape[0]
        x_img = x.view(B, self.C, self.H, self.W)
        y = self.base(x_img, t)                 # possibly (B, C, H', W')
        if y.dim() == 4:
            y = _crop_or_resize4d(y, self.H, self.W).reshape(B, -1)  # -> (B, D)
        else:
            y = y.reshape(B, -1)
        return y

def to_list_of_arrays(x):
    if isinstance(x, list): return [np.asarray(a) for a in x]
    x = np.asarray(x, dtype=object); return [np.asarray(a) for a in x.tolist()]

# def plot_frame(data, T, trial_idx, mean_img = None, idxs=None, n = 6):
#     if idxs is None:
#         idxs = np.linspace(0, T-2, n, dtype=int)
#     fig, axes = plt.subplots(1, len(idxs), figsize=(12, 2))
#     for ax, i in zip(axes, idxs):
#         if mean_img is None:
#             ax.imshow(data[trial_idx][i,0,:,:], cmap = 'gray')
#         else:
#             ax.imshow(data[trial_idx][i,0,:,:] + mean_img, cmap = 'gray')
#         ax.set_title(f"t={i}")
#         ax.axis("off")
#     plt.tight_layout()
    # plt.show()

def plot_frame(data, T, trial_idx, mean_img=None, idxs=None, n=6, colorbar=True, figsize=(12,2)):
    if idxs is None:
        idxs = np.linspace(0, T-2, n, dtype=int)

    X = data[trial_idx][:, 0]                       # [T, H, W]
    base = (X + mean_img) if mean_img is not None else X
    vmin, vmax = float(base.min()), float(base.max())

    fig, axes = plt.subplots(1, len(idxs), figsize=figsize)
    last_im = None
    for ax, i in zip(axes, idxs):
        img = (data[trial_idx][i, 0] + mean_img) if mean_img is not None else data[trial_idx][i, 0]
        last_im = ax.imshow(img, cmap='gray', vmin=vmin, vmax=vmax)
        ax.set_title(f"t={i}")
        ax.axis("off")
    if colorbar:
        fig.colorbar(last_im, ax=axes.ravel().tolist(), orientation='horizontal', fraction=0.05, pad=0.10)
    plt.tight_layout()


def plot_grad_frame(data, T, trial_idx, n=6, idxs=None, signed=False):
    x = data[trial_idx]                  # (T, 1, 28, 28)
    g = np.diff(x[:, 0], axis=0)         # (T-1, 28, 28), temporal gradient

    if idxs is None:
        idxs = np.linspace(0, T-2, n, dtype=int)

    if signed:
        vmax = np.max(np.abs(g))
        vmin, vmax = -vmax, vmax
        cmap = 'seismic'
        gshow = g
    else:
        vmin, vmax = 0, np.max(np.abs(g))
        cmap = 'viridis'
        gshow = np.abs(g)

    fig, axes = plt.subplots(1, len(idxs), figsize=(12, 2))
    ims = []
    for ax, i in zip(axes, idxs):
        im = ax.imshow(gshow[i], cmap=cmap, vmin=vmin, vmax=vmax)
        ax.set_title(f"t={i}→{i+1}")
        ax.axis("off")
        ims.append(im)
    
    cbar = fig.colorbar(ims[0], ax=axes.ravel().tolist(), orientation='horizontal', fraction=0.06, pad=0.08)
    cbar.set_label('temporal gradient' if signed else '|temporal gradient|')
    plt.tight_layout()

# --- helper: box-sum via integral image (no scipy)
def _box_sum(A, r):
    P = np.pad(A, ((r+1, r), (r+1, r)), mode='edge')
    I = P.cumsum(0).cumsum(1)
    return (I[2*r+1:, 2*r+1:] - I[:-2*r-1, 2*r+1:]
            - I[2*r+1:, :-2*r-1] + I[:-2*r-1, :-2*r-1])

# Lucas–Kanade optical flow (between two frames)
def lk_flow(img0, img1, r=2, lam=1e-3):
    Ibar = 0.5*(img0 + img1)
    Iy, Ix = np.gradient(Ibar)
    It = img1 - img0
    Sxx = _box_sum(Ix*Ix, r);  Syy = _box_sum(Iy*Iy, r);  Sxy = _box_sum(Ix*Iy, r)
    Sxt = _box_sum(Ix*It, r);  Syt = _box_sum(Iy*It, r)
    det = Sxx*Syy - Sxy*Sxy + lam
    u = (-Syy*Sxt + Sxy*Syt) / det
    v = ( Sxy*Sxt - Sxx*Syt) / det
    return u, v, np.abs(It)

# custom "white -> LIGHT red" cmap with transparency increasing with value
def _light_red_cmap():
    # RGBA: low -> white transparent, high -> light red semi-opaque
    return LinearSegmentedColormap.from_list(
        "white_to_lightred",
        [(1,1,1,0.0), (1.0, 0.75, 0.75, 0.85)]
    )

def plot_optical_flow_frames(
        data, T, trial_idx, n=6, idxs=None,
        r=2, lam=1e-3, stride=2,
        img_alpha=0.30,                # make base image light
        arrow_scale=8.0,               # bigger arrows; tune 4–12
        arrow_width=0.012,             # thicker arrows
        grad_quantile=99,              # robust grad vmax for color/scale
        show_image_bg=True,
):
    """
    Overlay order per panel: gray image (light) + red/white grad magnitude + big arrows.
    Colorbar shows gradient magnitude (|It|). Orientation matches your plots (origin='upper').
    """
    x = data[trial_idx][:, 0]  # (T,H,W)
    H, W = x.shape[1:]

    if idxs is None:
        idxs = np.linspace(0, T-2, n, dtype=int)

    yy, xx = np.meshgrid(np.arange(H), np.arange(W), indexing='ij')

    # Precompute flows and gradient magnitudes
    flows = []
    all_gm = []
    for i in idxs:
        u, v, gm = lk_flow(x[i], x[i+1], r=r, lam=lam)
        flows.append((i, u, v, gm))
        all_gm.append(gm)
    all_gm = np.stack(all_gm, 0)
    gm_vmax = np.percentile(all_gm, grad_quantile) + 1e-9
    norm = Normalize(vmin=0.0, vmax=gm_vmax)
    cmap = _light_red_cmap()

    fig, axes = plt.subplots(1, len(idxs), figsize=(12, 2.8), constrained_layout=True)
    axes = np.atleast_1d(axes)
    mappable_for_cb = None

    for ax, (i, u, v, gm) in zip(axes, flows):
        # 1) light gray base image
        if show_image_bg:
            ax.imshow(x[i], origin='upper', cmap='gray', alpha=img_alpha)

        # 2) red-white gradient overlay (large => light red)
        im = ax.imshow(gm, origin='upper', cmap=cmap, norm=norm)
        if mappable_for_cb is None: mappable_for_cb = im

        # 3) big arrows; scale by robust max so they are visible
        mag = np.hypot(u, v)
        denom = np.percentile(mag, 98) + 1e-9
        s = arrow_scale / denom

        ax.quiver(xx[::stride, ::stride], yy[::stride, ::stride],
                  (u*s)[::stride, ::stride], (v*s)[::stride, ::stride],
                  color='black', angles='xy', scale_units='xy', scale=1.0,
                  pivot='middle', width=arrow_width)
        ax.set_title(f"t={i}→{i+1}")
        ax.axis('off')

    # single horizontal colorbar for |It|
    cb = fig.colorbar(mappable_for_cb, ax=axes, orientation='horizontal', fraction=0.06, pad=0.08)
    cb.set_label('|It| (gradient magnitude)')
    # plt.show()

def _to_uint8(x):
    x = x.astype(np.float32)
    x = (x - x.min()) / (x.max() - x.min() + 1e-8)
    return (x * 255).astype(np.uint8)

def _save_gif(frames_THW, out_path, fps=30):
    # frames_THW: [T, H, W], float/uint8
    arr = frames_THW
    if arr.dtype != np.uint8:
        arr = _to_uint8(arr)
    pil_frames = [Image.fromarray(arr[t]) for t in range(arr.shape[0])]
    pil_frames[0].save(out_path, save_all=True, append_images=pil_frames[1:],
                       duration=int(1000 / fps), loop=0)

def plot_latent_time_series(
    mu_traj_np,             # list of [T_i, dim_preserve]
    dims=(0,),              # which latent dims to plot (0-based)
    trial_indices=None,
    labels=None,            # None -> black; numeric -> depends on label_mode
    label_mode='auto',      # 'auto' | 'categorical' | 'continuous'
    dt=None,
    linewidth=1.2,
    alpha=0.9,
    fig_w=8,
    fig_h_per=2,
    cont_cmap="twilight", # "twilight_shifted"
    cont_vrange=None        # (vmin, vmax)
):
    if trial_indices is None:
        trial_indices = list(range(len(mu_traj_np)))
    dims = (dims,) if isinstance(dims, (int, np.integer)) else list(dims)

    # collect non-empty trials
    data = {i: mu_traj_np[i] for i in trial_indices if len(mu_traj_np[i]) > 0}
    if not data:
        raise ValueError("No non-empty trials to plot.")

    # build label lookup
    lab_for = None
    use_labels = labels is not None
    if use_labels:
        if isinstance(labels, dict):
            lab_for = {i: labels.get(i) for i in trial_indices if i in data}
        else:
            arr = np.asarray(labels)
            lab_for = {i: (arr[i] if i < arr.shape[0] else None) for i in trial_indices if i in data}

        # determine mode
        if label_mode == 'auto':
            vals = [lab_for[i] for i in lab_for if lab_for[i] is not None]
            # if all ints -> categorical; else continuous if all numeric
            all_int = all(isinstance(v, (int, np.integer)) for v in vals)
            all_num = all(isinstance(v, (int, float, np.integer, np.floating)) for v in vals)
            use_cont = (not all_int) and all_num
        elif label_mode == 'continuous':
            use_cont = True
        else:
            use_cont = False
    else:
        use_cont = False

    # set up color mapping
    cmap = None
    norm = None
    cat_to_i = None
    cats = None

    if not use_labels:
        pass
    elif use_cont:
        vals = np.array([float(lab_for[i]) for i in lab_for], dtype=float)
        if cont_vrange is None:
            vmin, vmax = np.nanmin(vals), np.nanmax(vals)
        else:
            vmin, vmax = cont_vrange

        default_name = plt.rcParams.get('image.cmap', 'viridis')
        cmap = plt.get_cmap(default_name if cont_cmap is None else cont_cmap)
        norm = Normalize(vmin=vmin, vmax=vmax)
    else:
        # categorical palette (keep your original hsv style)
        cats = []
        for i in trial_indices:
            if i in data:
                v = lab_for.get(i, None)
                if v not in cats:
                    cats.append(v)
        K = max(1, len(cats))
        hsv = plt.get_cmap("hsv", K)
        cmap = ListedColormap([hsv(i) for i in range(K)])
        norm = BoundaryNorm(np.arange(K+1)-0.5, K)
        cat_to_i = {c: i for i, c in enumerate(cats)}

    # figure
    fig_h = fig_h_per * len(dims)
    fig, axes = plt.subplots(len(dims), 1, figsize=(fig_w, fig_h), squeeze=False)
    axes = axes.ravel()

    for ax, d in zip(axes, dims):
        for i in trial_indices:
            if i not in data:
                continue
            y = data[i][:, d]
            x = np.arange(len(y)) if dt is None else np.arange(len(y)) * dt

            if not use_labels:
                color = "k"
            elif use_cont:
                v = lab_for.get(i, np.nan)
                color = cmap(norm(v)) if np.isfinite(v) else (0.5, 0.5, 0.5, 1.0)  # gray if missing
            else:
                color = cmap(norm(cat_to_i[lab_for[i]]))

            ax.plot(x, y, lw=linewidth, alpha=alpha, color=color)

        ax.set_ylabel(f"μ{d+1}")
        ax.grid(True, alpha=0.25)

    axes[-1].set_xlabel("Time (step)" if dt is None else "Time")

    # colorbar
    if use_labels:
        fig.subplots_adjust(bottom=0.18)
        cax = fig.add_axes([0.12, 0.08, 0.76, 0.04])
        sm = plt.cm.ScalarMappable(norm=norm, cmap=cmap)
        sm.set_array([])
        cbar = plt.colorbar(sm, cax=cax, orientation="horizontal")
        if use_cont:
            cbar.set_label("Angle (deg)")
            if np.isclose(norm.vmin, -180) and np.isclose(norm.vmax, 180):
                cbar.set_ticks([-180, -90, 0, 90, 180])
        else:
            cbar.set_ticks(range(len(cats)))
            cbar.set_ticklabels([str(c) for c in cats])
            cbar.set_label("Trial category")

    plt.tight_layout(rect=[0.02, 0.18 if use_labels else 0.02, 0.98, 0.98])


def proj_to_latent(dset_samples, flow_net, tau_latent, device, show_progress=True, chunk_size=None):

    flow_net = flow_net.to(device).eval()
    is_image = (np.asarray(dset_samples[0]).ndim == 4)
    if is_image:
        _, C, H, W = np.asarray(dset_samples[0]).shape
        # Probe whether flow_net already returns flat; if not, wrap it
        with torch.inference_mode():
            test_x = torch.zeros((1, C*H*W), device=device, dtype=torch.float32)
            test_t = torch.tensor([1.0], device=device, dtype=torch.float32)
            test_y = flow_net(test_x, test_t)
        if test_y.dim() != 2:   # returns image-shaped → adapt to (B,D)->(B,D)
            flow_for_ode = _FlowFlatAdapter(flow_net, C, H, W).to(device).eval()
        else:
            flow_for_ode = flow_net
    else:
        flow_for_ode = flow_net
    
    latent_traj_list = []
    it = tqdm(range(len(dset_samples)), desc="Project → latent τ", leave=False) if show_progress \
         else range(len(dset_samples))

    with torch.inference_mode():
        for i in it:
            X = np.asarray(dset_samples[i])
            # flatten to (T, D)
            if X.ndim == 4:  # (T, C, H, W)
                T_i = X.shape[0]
                X_flat = X.reshape(T_i, -1)
            elif X.ndim == 2:  # (T, D)
                X_flat = X
            else:
                raise ValueError(f"Unexpected shape for dset_samples[{i}]: {X.shape}")

            X_flat_t = torch.from_numpy(X_flat).to(device=device, dtype=torch.float32)

            if chunk_size is None:
                # one shot: batch = T_i
                traj = util.calc_flow_trajectories(flow_for_ode, X_flat_t, 1.0, tau_latent, nt=2)
                X_tau = traj[-1]  # (T_i, D_flat)
                latent_traj_list.append(X_tau.detach().cpu().numpy())
            else:
                # optional chunking along T to reduce memory
                outs = []
                for s in range(0, X_flat_t.shape[0], int(chunk_size)):
                    e = min(s + int(chunk_size), X_flat_t.shape[0])
                    traj = util.calc_flow_trajectories(flow_for_ode, X_flat_t[s:e], 1.0, tau_latent, nt=2)
                    outs.append(traj[-1].detach().cpu())
                latent_traj_list.append(torch.cat(outs, dim=0).numpy())

    return latent_traj_list

def proj_to_latent(dset_samples, flow_net, tau_latent, device):
    latent_traj_list = []
    for ii in tqdm(range(len(dset_samples))):
        latent_traj_tmp = util.calc_flow_trajectories(flow_net,
                                                      torch.from_numpy(dset_samples[ii]).type(torch.float32).to(device),
                                                      1.0, tau_latent, nt=2)
        latent_traj_list.append(latent_traj_tmp[-1,:,:].cpu().numpy())
    return latent_traj_list

def latent_mu(latent_traj_list, encoder, dim_preserve, device, fullRes=False,
              min_var=None, rcond=None, resid_tol=1e-10, use_fp64=True):
    import torch
    enc = encoder.to(device).eval()
    mu_traj, loading = [], None

    def _flatten_Tx(z: torch.Tensor) -> torch.Tensor:
        return z.view(z.size(0), -1)

    with torch.inference_mode():
        arr0 = np.asarray(latent_traj_list[0], dtype=np.float32)
        Z0 = torch.from_numpy(arr0[:1]).to(device=device, dtype=torch.float32)
        _, D, L, bias = enc.encode(Z0, return_data_mean=True, update_stats=False)

        bias = bias.view(-1)

        # D is full diag; only first K are used
        d_vec = torch.diagonal(D) if D.ndim == 2 else D
        d_len = d_vec.numel()
        K = int(min(getattr(enc, 'k_max', d_len), d_len))

        mv = (max(1e-12, 1e-6 * torch.median(d_vec).item())
              if min_var is None else float(min_var))
        d_safe = torch.clamp(d_vec, min=mv)
        d_sqrt = torch.sqrt(d_safe[:K])               # [K]
        inv_sqrt_d = d_sqrt.reciprocal()              # [K]

        # Orthonormal columns, use only first K
        L_K = L[:, :K]                                 # [d, K]

        def solve_mu_batch(Z):
            # Z: [T,...] -> [T,d]
            Z_flat = _flatten_Tx(Z).to(device=device, dtype=torch.float32)
            Zb = Z_flat - bias.unsqueeze(0).to(Z_flat.dtype)   # [T,d]

            ZbT = Zb.T  # [d,T]
            if use_fp64:
                Ld = L_K.double()
                ZbT = ZbT.double()
                invsd = inv_sqrt_d.double()
                Y = Ld.T @ ZbT                # [K,T]
                MUt = invsd.unsqueeze(1) * Y  # [K,T]
                mu_full = MUt.T.float()       # [T,K]
            else:
                Y = L_K.T @ ZbT               # [K,T]
                MUt = inv_sqrt_d.unsqueeze(1) * Y
                mu_full = MUt.T               # [T,K]

            return mu_full  # [T,K]

        bad = False
        for arr in latent_traj_list:
            Z = torch.from_numpy(arr).to(device=device, dtype=torch.float32)
            mu_full = solve_mu_batch(Z)                     # [T,K]
            Z_flat  = _flatten_Tx(Z).to(mu_full.dtype)      # [T,d]

            # Reconstruct for residual check
            loading_now = (L_K * d_sqrt.unsqueeze(0)).to(mu_full.dtype)  # [d,K]
            Z_hat = (loading_now @ mu_full.T).T + bias.to(mu_full.dtype)  # [T,d]

            r = (Z_hat - Z_flat).norm() / (Z_flat.norm() + 1e-12)
            if torch.isnan(mu_full).any() or torch.isinf(mu_full).any() or r > resid_tol:
                bad = True
                break

            mu_traj.append(mu_full[:, :dim_preserve].cpu())

        if bad:
            # Exact left inverse on span(L_K): W = D_K^{-1/2} L_K^T
            loading = L_K * d_sqrt.unsqueeze(0)             # [d,K]
            W = (L_K.T * inv_sqrt_d.unsqueeze(1)).float()   # [K,d]
            mu_traj = []
            for arr in latent_traj_list:
                Z = torch.from_numpy(arr).to(device=device, dtype=torch.float32)
                Zf = _flatten_Tx(Z)
                Zb = Zf - bias
                mu_full = (W @ Zb.T).T                      # [T,K]
                mu_traj.append(mu_full[:, :dim_preserve].cpu())

    if fullRes:
        # return effective loading L_K sqrt(D_K)
        if loading is None:
            loading = L_K * d_sqrt.unsqueeze(0)
        return mu_traj, d_vec, L_K, loading, bias
    return mu_traj


# def generate_traj_image_onthefly(
#     dyn_net, dset_samples, n_step, tau, device, lag_k,
#     include_x0_tau=False, oracle=False,
#     flow_net=None, clamp_range=None, cov_static_list=None,
#     init_lag_mode="repeat_first"
# ):
    
#     # --- Infer geometry ---
#     assert len(dset_samples) > 0 and np.asarray(dset_samples[0]).ndim == 4, \
#         "This function is for image data only."
#     T0, C, H, W = np.asarray(dset_samples[0]).shape
#     D = C * H * W
#     B = len(dset_samples)

#     # --- clamp prep ---
#     gmin = gmax = None
#     cmin = cmax = None
#     if clamp_range is not None:
#         cr = np.asarray(clamp_range)
#         if cr.ndim == 1 and cr.shape[0] == 2:
#             gmin = torch.tensor(float(cr[0]), dtype=torch.float32, device=device)
#             gmax = torch.tensor(float(cr[1]), dtype=torch.float32, device=device)
#         elif cr.ndim == 4 and cr.shape[0] == 2 and cr.shape[1:] == (C, H, W):
#             cmin = torch.as_tensor(cr[0], dtype=torch.float32, device=device)
#             cmax = torch.as_tensor(cr[1], dtype=torch.float32, device=device)

#     # --- x(τ) at first frame ---
#     x0 = np.stack([np.asarray(dset_samples[i][0]).reshape(-1) for i in range(B)], 0)
#     curr_tau_pts = torch.as_tensor(x0, dtype=torch.float32, device=device).unsqueeze(1)  # (B,1,D)
#     flow_for_ode = flow_net
#     if tau < 1.0:
#         assert flow_net is not None, "flow_net required when tau<1.0."
#         with torch.no_grad():
#             # ensure (B,D)->(B,D) interface for flow trajectories
#             test_out = flow_net(torch.zeros((1, C*H*W), device=device), torch.tensor([1.0], device=device))
#             if test_out.dim() != 2:
#                 flow_for_ode = _FlowFlatAdapter(flow_net, C, H, W).to(device).eval()
#             xt = util.calc_flow_trajectories(flow_for_ode, curr_tau_pts.squeeze(1), 1.0, tau, nt=2)[-1]
#         curr_tau_pts = xt.unsqueeze(1)

#     # --- helpers ---
#     def static_t(ii):
#         if cov_static_list is None:
#             return None
#         rows = []
#         for j in range(B):
#             S = np.asarray(cov_static_list[j])  # (Tj, S)
#             idx = min(ii, S.shape[0] - 1)
#             rows.append(torch.as_tensor(S[idx], dtype=torch.float32, device=device))
#         return torch.stack(rows, 0) if rows else None  # (B,S) or None


#     def build_lag_from_truncated(ii):
#         """Oracle: lag from truncated frames [t-k..t], pad on left; map to τ if τ<1."""
#         lag_tensors = []
#         for j in range(B):
#             X = np.asarray(dset_samples[j])  # (Tj, C, H, W)
#             if lag_k == 0:
#                 frames = [X[min(ii, X.shape[0]-1)]]
#             else:
#                 start = max(0, ii - lag_k)
#                 frames = [X[start + s if (start + s) < X.shape[0] else X.shape[0]-1]
#                           for s in range(min(lag_k, ii) + 1)]
#                 if len(frames) < (lag_k + 1):
#                     frames = [X[0]] * ((lag_k + 1) - len(frames)) + frames
#             # ((k+1), C, H, W) -> ((k+1), D)
#             fr = torch.as_tensor(np.stack(frames, 0), dtype=torch.float32, device=device)
#             fr_flat = fr.view(fr.shape[0], -1)
#             if tau < 1.0:
#                 assert flow_for_ode is not None, "flow_net required for oracle τ<1.0."
#                 with torch.no_grad():
#                     fr_flat = util.calc_flow_trajectories(flow_for_ode, fr_flat, 1.0, tau, nt=2)[-1]
#             lag_tensors.append(fr_flat.view(-1))  # ((k+1)*D)
#         return torch.stack(lag_tensors, 0)  # (B,(k+1)*D)

#     # --- init τ-history lag from current τ state ---
#     x0_tau_img = curr_tau_pts.squeeze(1).view(B, C, H, W)       # (B,C,H,W)
#     lag_vec = x0_tau_img.repeat(1, lag_k + 1, 1, 1).view(B, -1) if lag_k > 0 else curr_tau_pts.squeeze(1)

#     # --- rollout ---
#     traj_out = [curr_tau_pts.cpu().numpy()]
#     lag_cov_list = []

#     for ii in tqdm(range(n_step - 1)):
#         # state & lag for this step
#         if oracle:
#             # x0 from data -> map to τ if needed
#             x0_tmp = torch.stack([
#                 torch.as_tensor(dset_samples[j][min(ii, len(dset_samples[j]) - 1)],
#                                 dtype=torch.float32, device=device).reshape(-1)
#                 for j in range(B)
#             ], dim=0)  # (B,D)
#             if tau < 1.0:
#                 assert flow_for_ode is not None, "flow_net required for oracle τ<1.0."
#                 with torch.no_grad():
#                     x0_tmp = util.calc_flow_trajectories(flow_for_ode, x0_tmp, 1.0, tau, nt=2)[-1]
#             lag_now = build_lag_from_truncated(ii)  # already τ-mapped when τ<1
#         else:
#             x0_tmp = curr_tau_pts.squeeze(1)  # (B,D)
#             lag_now = lag_vec  

#         # clamp lag
#         if clamp_range is not None:
#             if cmin is not None:
#                 lag_now = torch.clamp(lag_now.view(B, lag_k + 1, C, H, W), min=cmin, max=cmax).view(B, -1)
#             elif gmin is not None:
#                 lag_now = lag_now.clamp(min=gmin, max=gmax)

#         # stitch static cov
#         S_now = static_t(ii)
#         cov_stat_total = torch.cat([S_now, lag_now], dim=1) if S_now is not None else lag_now

#         # one step in τ
#         with torch.no_grad():
#             x_next = util.calc_dyn_trajectories(
#                 dyn_net, x0_tmp, tau, x0_tau=None,
#                 covariates=None, next_covariates=None,
#                 covariates_static=cov_stat_total,
#                 include_x0_tau=include_x0_tau, nt=2
#             )[-1]  # (B,D)

#         # clamp state
#         if clamp_range is not None:
#             if gmin is not None:
#                 x_next = x_next.clamp(min=gmin, max=gmax)
#             elif cmin is not None:
#                 xi = x_next.view(B, C, H, W)
#                 xi = torch.clamp(xi, min=cmin, max=cmax)
#                 x_next = xi.view(B, -1)

#         # push time
#         curr_tau_pts = x_next.unsqueeze(1)
#         traj_out.append(curr_tau_pts.cpu().numpy())

#         # roll tau-history
#         if (not oracle):
#             ximg_tau = x_next.view(B, C, H, W)
#             if lag_k > 0:
#                 if ii == 0 and 'lag_img_stack' not in locals():
#                     lag_img_stack = ximg_tau.repeat(1, lag_k + 1, 1, 1)
#                 else:
#                     lag_img_stack = torch.cat([lag_img_stack[:, C:], ximg_tau], dim=1)
#                 lag_vec = lag_img_stack.view(B, -1)
#             else:
#                 lag_vec = x_next
#         # record lag used this step
#         lag_cov_list.append(lag_now)
        

#     out = np.concatenate(traj_out, axis=1)  # (B,Tgen,D)
#     out_trunc = [out[i, :np.asarray(dset_samples[i]).shape[0], :] for i in range(B)]
#     return out_trunc, lag_cov_list

def generate_traj_image_onthefly(
    dyn_net, dset_samples, n_step, tau, device, lag_k,
    include_x0_tau=False, oracle=False,
    flow_net=None, clamp_range=None, cov_static_list=None,
    dset_samples_full=None,
):
    # --- Infer geometry ---
    assert len(dset_samples) > 0 and np.asarray(dset_samples[0]).ndim == 4, \
        "This function is for image data only."
    T0, C, H, W = np.asarray(dset_samples[0]).shape
    D = C * H * W
    B = len(dset_samples)

    # --- offsets if full dataset is provided ---
    use_full = dset_samples_full is not None
    if use_full:
        assert len(dset_samples_full) == B
        offsets = []
        for j in range(B):
            Tf = int(np.asarray(dset_samples_full[j]).shape[0])
            Tt = int(np.asarray(dset_samples[j]).shape[0])
            off = Tf - Tt
            # Expect off == lag_k in your setup, but do not hard fail
            offsets.append(off)
    else:
        offsets = [0] * B

    # --- clamp prep ---
    gmin = gmax = None
    cmin = cmax = None
    if clamp_range is not None:
        cr = np.asarray(clamp_range)
        if cr.ndim == 1 and cr.shape[0] == 2:
            gmin = torch.tensor(float(cr[0]), dtype=torch.float32, device=device)
            gmax = torch.tensor(float(cr[1]), dtype=torch.float32, device=device)
        elif cr.ndim == 4 and cr.shape[0] == 2 and cr.shape[1:] == (C, H, W):
            cmin = torch.as_tensor(cr[0], dtype=torch.float32, device=device)
            cmax = torch.as_tensor(cr[1], dtype=torch.float32, device=device)

    # --- x(τ) at first frame (from truncated dataset) ---
    x0 = np.stack([np.asarray(dset_samples[i][0]).reshape(-1) for i in range(B)], 0)
    curr_tau_pts = torch.as_tensor(x0, dtype=torch.float32, device=device).unsqueeze(1)  # (B,1,D)

    flow_for_ode = flow_net
    if tau < 1.0:
        assert flow_net is not None, "flow_net required when tau<1.0."
        with torch.no_grad():
            test_out = flow_net(torch.zeros((1, C*H*W), device=device), torch.tensor([1.0], device=device))
            if test_out.dim() != 2:
                flow_for_ode = _FlowFlatAdapter(flow_net, C, H, W).to(device).eval()
            xt = util.calc_flow_trajectories(flow_for_ode, curr_tau_pts.squeeze(1), 1.0, tau, nt=2)[-1]
        curr_tau_pts = xt.unsqueeze(1)

    # --- helpers ---
    def static_t(ii):
        if cov_static_list is None:
            return None
        rows = []
        for j in range(B):
            S = np.asarray(cov_static_list[j])  # (Tj, S)
            idx = min(ii, S.shape[0] - 1)
            rows.append(torch.as_tensor(S[idx], dtype=torch.float32, device=device))
        return torch.stack(rows, 0) if rows else None  # (B,S) or None

    def _tau_map_frames(fr_flat):
        # fr_flat: (N, D)
        if tau < 1.0:
            assert flow_for_ode is not None, "flow_net required when tau<1.0."
            with torch.no_grad():
                fr_flat = util.calc_flow_trajectories(flow_for_ode, fr_flat, 1.0, tau, nt=2)[-1]
        return fr_flat

    def build_lag_from_truncated(ii):
        """Pad on left using first truncated frame."""
        lag_tensors = []
        for j in range(B):
            X = np.asarray(dset_samples[j])  # (Tj, C, H, W)
            Tj = X.shape[0]
            t = min(ii, Tj - 1)

            if lag_k == 0:
                frames = [X[t]]
            else:
                start = max(0, t - lag_k)
                frames = [X[tt] for tt in range(start, t + 1)]
                if len(frames) < (lag_k + 1):
                    frames = [X[0]] * ((lag_k + 1) - len(frames)) + frames

            fr = torch.as_tensor(np.stack(frames, 0), dtype=torch.float32, device=device)  # (L,C,H,W)
            fr_flat = fr.view(fr.shape[0], -1)  # (L,D)
            fr_flat = _tau_map_frames(fr_flat)
            lag_tensors.append(fr_flat.view(-1))  # ((k+1)*D)
        return torch.stack(lag_tensors, 0)  # (B,(k+1)*D)

    def build_lag_from_full(ii):
        """True history from full dataset aligned by offset = T_full - T_trunc."""
        lag_tensors = []
        for j in range(B):
            Xf = np.asarray(dset_samples_full[j])  # (Tf, C, H, W)
            Tf = Xf.shape[0]
            t_abs = min(ii + offsets[j], Tf - 1)  # absolute time in full

            if lag_k == 0:
                frames = [Xf[t_abs]]
            else:
                frames = []
                for tt in range(t_abs - lag_k, t_abs + 1):
                    if tt < 0:
                        frames.append(Xf[0])
                    else:
                        frames.append(Xf[min(tt, Tf - 1)])

            fr = torch.as_tensor(np.stack(frames, 0), dtype=torch.float32, device=device)  # (L,C,H,W)
            fr_flat = fr.view(fr.shape[0], -1)  # (L,D)
            fr_flat = _tau_map_frames(fr_flat)
            lag_tensors.append(fr_flat.view(-1))  # ((k+1)*D)
        return torch.stack(lag_tensors, 0)  # (B,(k+1)*D)

    def init_lag_from_full_at_start():
        # history for the initial truncated frame corresponds to ii=0 in truncated
        # which is t_abs = offsets[j] in full
        lag0 = build_lag_from_full(ii=0)  # (B, (k+1)*D)
        if lag_k > 0:
            # also keep image-stacked form for efficient rolling
            hist = lag0.view(B, lag_k + 1, D).view(B, lag_k + 1, C, H, W)
            lag_img_stack = hist.view(B, (lag_k + 1) * C, H, W)  # (B,(k+1)*C,H,W)
            lag_vec = lag_img_stack.view(B, -1)
            return lag_img_stack, lag_vec
        else:
            return None, lag0  # lag_vec is (B,D)

    # --- init τ-history lag ---
    if lag_k > 0:
        if use_full:
            lag_img_stack, lag_vec = init_lag_from_full_at_start()
        else:
            # fallback: pad by repeating current τ state
            x0_tau_img = curr_tau_pts.squeeze(1).view(B, C, H, W)
            lag_img_stack = x0_tau_img.repeat(1, lag_k + 1, 1, 1)  # (B,(k+1)*C,H,W)
            lag_vec = lag_img_stack.view(B, -1)
    else:
        lag_img_stack = None
        lag_vec = curr_tau_pts.squeeze(1)  # (B,D)

    # --- rollout ---
    traj_out = [curr_tau_pts.cpu().numpy()]
    lag_cov_list = []

    for ii in tqdm(range(n_step - 1)):
        # state & lag for this step
        if oracle:
            # x0 from truncated data -> map to τ if needed
            x0_tmp = torch.stack([
                torch.as_tensor(dset_samples[j][min(ii, len(dset_samples[j]) - 1)],
                                dtype=torch.float32, device=device).reshape(-1)
                for j in range(B)
            ], dim=0)  # (B,D)
            if tau < 1.0:
                assert flow_for_ode is not None, "flow_net required for oracle τ<1.0."
                with torch.no_grad():
                    x0_tmp = util.calc_flow_trajectories(flow_for_ode, x0_tmp, 1.0, tau, nt=2)[-1]

            if lag_k > 0:
                lag_now = build_lag_from_full(ii) if use_full else build_lag_from_truncated(ii)
            else:
                lag_now = x0_tmp  # (B,D)
        else:
            x0_tmp = curr_tau_pts.squeeze(1)  # (B,D)
            lag_now = lag_vec                 # (B,(k+1)*D) or (B,D)

        # clamp lag
        if clamp_range is not None:
            if cmin is not None and lag_k > 0:
                lag_now = torch.clamp(lag_now.view(B, lag_k + 1, C, H, W), min=cmin, max=cmax).view(B, -1)
            elif gmin is not None:
                lag_now = lag_now.clamp(min=gmin, max=gmax)

        # stitch static cov
        S_now = static_t(ii)
        cov_stat_total = torch.cat([S_now, lag_now], dim=1) if S_now is not None else lag_now

        # one step in τ
        with torch.no_grad():
            x_next = util.calc_dyn_trajectories(
                dyn_net, x0_tmp, tau, x0_tau=None,
                covariates=None, next_covariates=None,
                covariates_static=cov_stat_total,
                include_x0_tau=include_x0_tau, nt=2
            )[-1]  # (B,D)

        # clamp state
        if clamp_range is not None:
            if gmin is not None:
                x_next = x_next.clamp(min=gmin, max=gmax)
            elif cmin is not None:
                xi = x_next.view(B, C, H, W)
                xi = torch.clamp(xi, min=cmin, max=cmax)
                x_next = xi.view(B, -1)

        # push time
        curr_tau_pts = x_next.unsqueeze(1)
        traj_out.append(curr_tau_pts.cpu().numpy())

        # roll τ-history (only in non-oracle mode)
        if not oracle:
            if lag_k > 0:
                ximg_tau = x_next.view(B, C, H, W)
                lag_img_stack = torch.cat([lag_img_stack[:, C:], ximg_tau], dim=1)  # shift, append
                lag_vec = lag_img_stack.view(B, -1)
            else:
                lag_vec = x_next

        # record lag actually used at this step (do not record the rolled version)
        lag_cov_list.append(lag_now)

    out = np.concatenate(traj_out, axis=1)  # (B,Tgen,D)
    out_trunc = [out[i, :np.asarray(dset_samples[i]).shape[0], :] for i in range(B)]
    return out_trunc, lag_cov_list
```
