## Supplementary material for "Dynamic Compression Flows for Neuroscience Data": Python code: plot_sim_results_v7.py

```
import numpy as np
import matplotlib.pyplot as plt
from matplotlib.collections import LineCollection
from matplotlib.colors import ListedColormap, BoundaryNorm, LinearSegmentedColormap, Normalize
from mpl_toolkits.mplot3d.art3d import Line3DCollection
import torch
import dnnlib.util_v7 as util
from tqdm import tqdm
import pickle
from matplotlib.ticker import FuncFormatter
from torch_utils import persistence
from torch import nn
import types
import matplotlib as mpl
from matplotlib import cm
import matplotlib.gridspec as gridspec

device = torch.device("cuda" if torch.cuda.is_available() else "cpu")
torch.set_default_device(device)

def infer_lag_k_from_dyn_net(dyn_net):
    # Works for ConvVNetWrapper only
    lag_k = None
    if hasattr(dyn_net, 'lag_ch') and hasattr(dyn_net, 'img_ch'):
        # lag_ch = (k+1) * C  ->  k = lag_ch / C - 1
        C = int(dyn_net.img_ch)
        lag_k = int(dyn_net.lag_ch // C) - 1 if C > 0 else None
    return lag_k

def read_model(folder, chk_pts):
    pkl_path = folder + chk_pts
    if not hasattr(persistence, "_orig_reconstruct"):
        persistence._orig_reconstruct = persistence._reconstruct_persistent_obj
    def _tolerant_reconstruct(meta):
        try:
            return persistence._orig_reconstruct(meta)
        except Exception:
            class _Stub: pass
            return _Stub()

    class PatchedUnpickler(pickle.Unpickler):
        def find_class(self, module, name):
            if module == "torch_utils.persistence" and name == "_reconstruct_persistent_obj":
                return _tolerant_reconstruct
            return super().find_class(module, name)

    with open(pkl_path, "rb") as f:
        model_all = PatchedUnpickler(f).load()

    net = model_all["ema"] if "ema" in model_all else model_all["net"]
    flow_net = net.unet_model  # Compressive flow (u_theta)
    dyn_net = net.vnet_model   # Dynamics flow (v_theta)
    encoder = net.encoder
    lag = infer_lag_k_from_dyn_net(dyn_net)
    return flow_net, dyn_net, encoder, lag

def apply_posthoc_alignment(target_encoder, ref_encoder):
    with torch.no_grad():
        tgt = getattr(target_encoder, 'module', target_encoder)
        ref = getattr(ref_encoder, 'module', ref_encoder)

        # Determine shared dimensions
        K = min(tgt.k_max, ref.k_max)
        D = min(tgt.input_dim, ref.input_dim)

        # Get L matrices
        tgt_L = tgt.L_head[:D, :K]
        ref_L = ref.L_head[:D, :K].to(tgt_L.device)

        # Calculate alignment
        dots = (tgt_L * ref_L).sum(dim=0) # [K]
        signs = torch.sign(dots)
        signs[signs == 0] = 1.0
        
        # Save signs as a buffer so it moves with the model (cpu/gpu)
        if not hasattr(tgt, 'posthoc_signs'):
            tgt.register_buffer('posthoc_signs', signs)
        else:
            tgt.posthoc_signs.copy_(signs)

        print(f"Aligning signs for {(signs < 0).sum().item()} columns.")

    # Flip L_head
    with torch.no_grad():
        tgt.L_head.data[:D, :K] *= tgt.posthoc_signs.unsqueeze(0)

    if not hasattr(tgt, '_original_encode'):
        tgt._original_encode = tgt.encode

    # Define the wrapper function
    def aligned_encode(self, x, return_data_mean=True, update_stats=False):
        
        original_out = self._original_encode(x, return_data_mean=return_data_mean, update_stats=update_stats)
        
        # Unpack, align mu, and repack
        mu, D, L = original_out[:3]
        K_signs = self.posthoc_signs.shape[0]
        
        mu_aligned = mu.clone()
        mu_aligned[:, :K_signs] *= self.posthoc_signs.unsqueeze(0)
        
        if return_data_mean:
            return (mu_aligned, D, L, original_out[3])
        else:
            return (mu_aligned, D, L)

            
    tgt.encode = types.MethodType(aligned_encode, tgt)

    print("Target encoder successfully patched.")


def plotTraj(data, data_sim = None, title = None):
    data_min = np.min(data)
    data_max = np.max(data)
    data_range = data_max - data_min
    offset = data_range * 1.1

    time = np.arange(data.shape[0])
    if data_sim is not None:
        time_sim = np.arange(data_sim.shape[0])
    
    # Plot the trajectories
    plt.figure(figsize=(6, 4))  # Adjust figure size as needed
    for i in range(data.shape[1]):
        adjusted_y = data[:, i] + i * offset
        plt.plot(time, adjusted_y, label=f'Trajectory {i+1}', linewidth=0.5, c='red') 
        if data_sim is not None:
            adjusted_y_sim = data_sim[:, i] + i * offset
            plt.plot(time_sim, adjusted_y_sim, label=f'Trajectory {i+1}', linewidth=0.5, c='black')
        
    # Customize the plot
    plt.xlabel('Time')
    plt.tight_layout()
    if title is not None:
        plt.title(title)
    # plt.show()

def plot_traj_3d(X, title = '3D Trajectory (color = time)'):
    X = np.asarray(X)                    # (T, 3)
    T = len(X)
    t = np.arange(T)

    # line segments between consecutive points
    segs = np.stack([X[:-1], X[1:]], axis=1)   # (T-1, 2, 3)

    fig = plt.figure(figsize=(6,5))
    ax = fig.add_subplot(projection='3d')

    norm = mpl.colors.Normalize(vmin=0, vmax=T-1)
    cmap = plt.get_cmap()               # use current default colormap

    lc = Line3DCollection(segs, cmap=cmap, norm=norm, linewidth=2)
    lc.set_array(t[1:])                 # color by time index of segment end
    ax.add_collection3d(lc)
    
    ax.scatter(X[:,0], X[:,1], X[:,2], c=t, s=12, cmap=cmap, norm=norm)

    # nice bounds/aspect
    lo, hi = X.min(0), X.max(0)
    ax.set_xlim(lo[0], hi[0]); ax.set_ylim(lo[1], hi[1]); ax.set_zlim(lo[2], hi[2])
    ax.set_box_aspect(hi - lo)

    ax.set_xlabel('x'); ax.set_ylabel('y'); ax.set_zlabel('z'); ax.set_title(title)
    fig.colorbar(lc, ax=ax, pad=0.02, fraction=0.05, label='time step')
    plt.tight_layout()
    plt.show()


def get_mu_from_encoder(encoder, data_flattent, dim_to_keep, trial_idx=None):
    import numpy as np
    encoder = encoder.to(device).eval()
    with torch.inference_mode():
        # normalize indices
        if trial_idx is None or trial_idx == 'all':
            indices = range(len(data_flattent))
        elif isinstance(trial_idx, slice):
            indices = range(len(data_flattent))[trial_idx]
        elif np.isscalar(trial_idx):
            indices = [int(trial_idx)]
        else:  # list/tuple/array
            indices = list(trial_idx)

        outs = []
        for idx in indices:
            X0 = torch.from_numpy(data_flattent[idx]).to(device=device, dtype=torch.float32)
            mu_data, D, L, _ = encoder.encode(X0)
            outs.append(mu_data[:, :dim_to_keep].detach().cpu().numpy())

    return np.concatenate(outs, axis=0)  # shape: (sum T_i, dim_to_keep)

# def generate_traj(dyn_net, dset_samples, n_step, tau, lag_samples, device, lag,
#                   include_x0_tau=False,
#                   oracle=False, flow_net=None, clamp_range=None):
#     # clamp prep
#     full_min = full_max = None
#     if clamp_range is not None and len(clamp_range.shape) == 2:
#         clamp_range = torch.from_numpy(clamp_range).to(device).float()
#         full_min = clamp_range[:, 0].tile(lag + 1).unsqueeze(0)
#         full_max = clamp_range[:, 1].tile(lag + 1).unsqueeze(0)

#     # x(τ) at first point
#     starting_pts = np.stack([dset_samples[ii][0, :] for ii in range(len(dset_samples))], 0)
#     starting_pts = torch.from_numpy(starting_pts).unsqueeze(1).float().to(device)  # (B,1,D)
#     if tau < 1.0:
#         assert flow_net is not None, "flow_net required when tau<1.0."
#         curr_tau_pts = util.calc_flow_trajectories(flow_net, starting_pts.squeeze(1), 1.0, tau, nt=2)[-1].unsqueeze(1)
#     else:
#         curr_tau_pts = starting_pts

#     B, _, D = curr_tau_pts.shape
#     x0_tau = curr_tau_pts.squeeze(1)                    # (B,D)
#     lag_cov = x0_tau.repeat(1, lag + 1) if lag > 0 else x0_tau.clone()
#     curr_tau_trajs, lag_cov_list = [curr_tau_pts.cpu().numpy()], []

#     for ii in tqdm(range(n_step - 1)):
#         # source state and lag
#         if oracle:
#             # lag at t -> τ if needed
#             lag_cov = torch.stack([
#                 torch.as_tensor(lag_samples[j][ii], dtype=torch.float32, device=device)
#                 if ii < lag_samples[j].shape[0] else
#                 torch.as_tensor(lag_samples[j][-1], dtype=torch.float32, device=device)
#                 for j in range(len(lag_samples))
#             ], dim=0)  # (B,(k+1)·D)
#             if tau < 1.0:
#                 assert flow_net is not None, "flow_net required for oracle τ<1.0."
#                 L = lag + 1
#                 lag_cov = lag_cov.view(B * L, D)
#                 with torch.no_grad():
#                     lag_cov = util.calc_flow_trajectories(flow_net, lag_cov, 1.0, tau, nt=2)[-1]
#                 lag_cov = lag_cov.view(B, L * D)
        
#             # x0 at t -> τ if needed
#             x0_tmp = torch.stack([
#                 torch.as_tensor(dset_samples[j][ii], dtype=torch.float32, device=device)
#                 if ii < dset_samples[j].shape[0] else
#                 torch.as_tensor(dset_samples[j][-1], dtype=torch.float32, device=device)
#                 for j in range(len(dset_samples))
#             ], dim=0)  # (B,D)
#             if tau < 1.0:
#                 with torch.no_grad():
#                     x0_tmp = util.calc_flow_trajectories(flow_net, x0_tmp, 1.0, tau, nt=2)[-1]
#         else:
#             x0_tmp = curr_tau_pts.squeeze(1)

#         # clamp lag
#         if clamp_range is not None:
#             if len(clamp_range.shape) == 1:
#                 lag_cov = lag_cov.clamp(min=clamp_range[0], max=clamp_range[1])
#             else:
#                 lag_cov = torch.clamp(lag_cov, min=full_min, max=full_max)

#         # one step in τ
#         traj = util.calc_dyn_trajectories(
#             dyn_net, x0_tmp, tau, x0_tau=None,
#             covariates=None, next_covariates=None,
#             covariates_static=lag_cov,
#             include_x0_tau=include_x0_tau, nt=2
#         )
#         x_next = traj[-1]  # (B,D)

#         # clamp state
#         if clamp_range is not None:
#             if len(clamp_range.shape) == 1:
#                 x_next = x_next.clamp(min=clamp_range[0], max=clamp_range[1])
#             else:
#                 x_next = torch.clamp(x_next, min=clamp_range[:, 0].unsqueeze(0), max=clamp_range[:, 1].unsqueeze(0))

#         # push time
#         curr_tau_pts = x_next.unsqueeze(1)
#         curr_tau_trajs.append(curr_tau_pts.cpu().numpy())

#         # roll tau
#         if not oracle:
#             if lag > 0:
#                 lag_cov = torch.cat([lag_cov[:, D:], x_next], dim=-1)
#             else:
#                 lag_cov = x_next
#         # record lag used this step
#         lag_cov_list.append(lag_cov)

#     curr_tau_trajs = np.concatenate(curr_tau_trajs, axis=1)
#     curr_tau_trajs_trunc = [curr_tau_trajs[i, :dset_samples[i].shape[0], :] for i in range(curr_tau_trajs.shape[0])]
#     return curr_tau_trajs_trunc, lag_cov_list

def generate_traj(
    dyn_net, dset_samples, n_step, tau,
    device, lag,
    include_x0_tau=False,
    oracle=False, flow_net=None, clamp_range=None,
    dset_samples_full=None,
):
    # ---- clamp prep (same spirit as your original) ----
    full_min = full_max = None
    clamp_t = None
    if clamp_range is not None:
        cr = np.asarray(clamp_range)
        if cr.ndim == 2:
            # per-dim clamp: (D,2)
            clamp_t = torch.as_tensor(cr, device=device, dtype=torch.float32)
            if lag > 0:
                full_min = clamp_t[:, 0].repeat(lag + 1).unsqueeze(0)  # (1,(lag+1)*D)
                full_max = clamp_t[:, 1].repeat(lag + 1).unsqueeze(0)
        else:
            # global clamp: (2,)
            clamp_t = torch.as_tensor(cr, device=device, dtype=torch.float32)

    # ---- helper: map to τ if tau<1 ----
    def _tau_map(x):  # x: (N,D)
        if tau < 1.0:
            assert flow_net is not None, "flow_net required when tau<1.0."
            with torch.no_grad():
                x = util.calc_flow_trajectories(flow_net, x, 1.0, tau, nt=2)[-1]
        return x

    # ---- offsets if full dataset is provided ----
    use_full = dset_samples_full is not None
    B = len(dset_samples)
    if use_full:
        assert len(dset_samples_full) == B
        offsets = []
        for j in range(B):
            Tf = int(np.asarray(dset_samples_full[j]).shape[0])
            Tt = int(np.asarray(dset_samples[j]).shape[0])
            offsets.append(Tf - Tt)  # in your pipeline this should equal lag
    else:
        offsets = [0] * B

    # ---- x(τ) at first point (from truncated dataset) ----
    starting_pts = np.stack([np.asarray(dset_samples[ii][0, :]) for ii in range(B)], 0)  # (B,D)
    starting_pts = torch.as_tensor(starting_pts, device=device, dtype=torch.float32)
    x0_tau = _tau_map(starting_pts)          # (B,D)
    curr_tau_pts = x0_tau.unsqueeze(1)       # (B,1,D)
    B, D = x0_tau.shape

    # ---- build lag (k+1)*D at a given truncated time ii ----
    # IMPORTANT: match training order in vfm_train_v7.py:
    #   [x_t, x_{t-1}, ..., x_{t-k}]  (newest -> oldest)
    def build_lag_from_trunc(ii):
        rows = []
        for j in range(B):
            X = np.asarray(dset_samples[j])  # (Tj,D)
            Tj = X.shape[0]
            t = min(ii, Tj - 1)
            if lag == 0:
                frames = [X[t]]
            else:
                frames = []
                for back in range(lag + 1):  # back=0 => x_t first
                    tt = t - back
                    frames.append(X[tt] if tt >= 0 else X[0])  # left pad with first available
            fr = torch.as_tensor(np.stack(frames, 0), device=device, dtype=torch.float32)  # (L,D)
            fr = _tau_map(fr)  # (L,D)
            rows.append(fr.reshape(-1))      # ((lag+1)*D)
        return torch.stack(rows, 0)          # (B,(lag+1)*D)

    def build_lag_from_full(ii):
        rows = []
        for j in range(B):
            Xf = np.asarray(dset_samples_full[j])  # (Tf,D)
            Tf = Xf.shape[0]
            t_abs = min(ii + offsets[j], Tf - 1)   # absolute time aligned to truncated index
            if lag == 0:
                frames = [Xf[t_abs]]
            else:
                frames = []
                for back in range(lag + 1):        # back=0 => x_t first
                    tt = t_abs - back
                    if tt < 0:
                        frames.append(Xf[0])
                    else:
                        frames.append(Xf[min(tt, Tf - 1)])
            fr = torch.as_tensor(np.stack(frames, 0), device=device, dtype=torch.float32)  # (L,D)
            fr = _tau_map(fr)
            rows.append(fr.reshape(-1))
        return torch.stack(rows, 0)

    # ---- init lag cov at start ----
    if lag > 0:
        lag_cov = build_lag_from_full(ii=0) if use_full else x0_tau.repeat(1, lag + 1)
    else:
        lag_cov = x0_tau.clone()  # (B,D)

    curr_tau_trajs = [curr_tau_pts.detach().cpu().numpy()]
    lag_cov_list = []

    for ii in tqdm(range(n_step - 1)):
        # ---- choose x0 and lag for this step ----
        if oracle:
            if use_full:
                x0_np = np.stack([
                    np.asarray(dset_samples_full[j])[min(ii + offsets[j], np.asarray(dset_samples_full[j]).shape[0] - 1)]
                    for j in range(B)
                ], 0)
            else:
                x0_np = np.stack([
                    np.asarray(dset_samples[j])[min(ii, np.asarray(dset_samples[j]).shape[0] - 1)]
                    for j in range(B)
                ], 0)

            x0_tmp = _tau_map(torch.as_tensor(x0_np, device=device, dtype=torch.float32))  # (B,D)
            if lag > 0:
                lag_now = build_lag_from_full(ii) if use_full else build_lag_from_trunc(ii)
            else:
                lag_now = x0_tmp
        else:
            x0_tmp = curr_tau_pts.squeeze(1)  # (B,D)
            lag_now = lag_cov                 # (B,(lag+1)*D) or (B,D)

        # ---- clamp lag ----
        if clamp_t is not None:
            if clamp_t.ndim == 1:
                lag_now = lag_now.clamp(min=clamp_t[0], max=clamp_t[1])
            else:
                if lag > 0:
                    lag_now = torch.clamp(lag_now, min=full_min, max=full_max)
                else:
                    lag_now = torch.clamp(lag_now,
                                          min=clamp_t[:, 0].unsqueeze(0),
                                          max=clamp_t[:, 1].unsqueeze(0))

        # ---- one step in τ ----
        with torch.no_grad():
            traj = util.calc_dyn_trajectories(
                dyn_net, x0_tmp, tau, x0_tau=None,
                covariates=None, next_covariates=None,
                covariates_static=lag_now,
                include_x0_tau=include_x0_tau, nt=2
            )
            x_next = traj[-1]  # (B,D)

        # ---- clamp state ----
        if clamp_t is not None:
            if clamp_t.ndim == 1:
                x_next = x_next.clamp(min=clamp_t[0], max=clamp_t[1])
            else:
                x_next = torch.clamp(x_next,
                                     min=clamp_t[:, 0].unsqueeze(0),
                                     max=clamp_t[:, 1].unsqueeze(0))

        # ---- push time ----
        curr_tau_pts = x_next.unsqueeze(1)
        curr_tau_trajs.append(curr_tau_pts.detach().cpu().numpy())

        # record the lag actually used this step (pre-roll)
        lag_cov_list.append(lag_now)

        # ---- roll lag (non-oracle only), keep newest-first order ----
        if not oracle:
            if lag > 0:
                # lag_cov = [x_t, x_{t-1}, ..., x_{t-lag}]
                # -> [x_{t+1}, x_t, ..., x_{t-lag+1}]
                lag_cov = torch.cat([x_next, lag_cov[:, :lag * D]], dim=-1)
            else:
                lag_cov = x_next

    curr_tau_trajs = np.concatenate(curr_tau_trajs, axis=1)  # (B,Tgen,D)
    curr_tau_trajs_trunc = [
        curr_tau_trajs[i, :np.asarray(dset_samples[i]).shape[0], :]
        for i in range(curr_tau_trajs.shape[0])
    ]
    return curr_tau_trajs_trunc, lag_cov_list


def plot_latent_time_series(
    mu_traj_np,             # list of [T_i, dim_preserve]
    dims=(0,),              # which latent dims to plot (0-based)
    trial_indices=None,
    labels=None,            # None -> black; numeric -> depends on label_mode
    label_mode='auto',      # 'auto' | 'categorical' | 'continuous'
    dt=None,
    linewidth=1.2,
    alpha=0.9,
    fig_w=8,
    fig_h_per=2,
    cont_cmap="twilight", # "twilight_shifted"
    cont_vrange=None        # (vmin, vmax)
):
    if trial_indices is None:
        trial_indices = list(range(len(mu_traj_np)))
    dims = (dims,) if isinstance(dims, (int, np.integer)) else list(dims)

    # collect non-empty trials
    data = {i: mu_traj_np[i] for i in trial_indices if len(mu_traj_np[i]) > 0}
    if not data:
        raise ValueError("No non-empty trials to plot.")

    # build label lookup
    lab_for = None
    use_labels = labels is not None
    if use_labels:
        if isinstance(labels, dict):
            lab_for = {i: labels.get(i) for i in trial_indices if i in data}
        else:
            arr = np.asarray(labels)
            lab_for = {i: (arr[i] if i < arr.shape[0] else None) for i in trial_indices if i in data}

        # determine mode
        if label_mode == 'auto':
            vals = [lab_for[i] for i in lab_for if lab_for[i] is not None]
            # if all ints -> categorical; else continuous if all numeric
            all_int = all(isinstance(v, (int, np.integer)) for v in vals)
            all_num = all(isinstance(v, (int, float, np.integer, np.floating)) for v in vals)
            use_cont = (not all_int) and all_num
        elif label_mode == 'continuous':
            use_cont = True
        else:
            use_cont = False
    else:
        use_cont = False

    # set up color mapping
    cmap = None
    norm = None
    cat_to_i = None
    cats = None

    if not use_labels:
        pass
    elif use_cont:
        vals = np.array([float(lab_for[i]) for i in lab_for], dtype=float)
        if cont_vrange is None:
            vmin, vmax = np.nanmin(vals), np.nanmax(vals)
        else:
            vmin, vmax = cont_vrange

        default_name = plt.rcParams.get('image.cmap', 'viridis')
        cmap = plt.get_cmap(default_name if cont_cmap is None else cont_cmap)
        norm = Normalize(vmin=vmin, vmax=vmax)
    else:
        # categorical palette (keep your original hsv style)
        cats = []
        for i in trial_indices:
            if i in data:
                v = lab_for.get(i, None)
                if v not in cats:
                    cats.append(v)
        K = max(1, len(cats))
        hsv = plt.get_cmap("hsv", K)
        cmap = ListedColormap([hsv(i) for i in range(K)])
        norm = BoundaryNorm(np.arange(K+1)-0.5, K)
        cat_to_i = {c: i for i, c in enumerate(cats)}

    # figure
    fig_h = fig_h_per * len(dims)
    fig, axes = plt.subplots(len(dims), 1, figsize=(fig_w, fig_h), squeeze=False)
    axes = axes.ravel()

    for ax, d in zip(axes, dims):
        for i in trial_indices:
            if i not in data:
                continue
            y = data[i][:, d]
            x = np.arange(len(y)) if dt is None else np.arange(len(y)) * dt

            if not use_labels:
                color = "k"
            elif use_cont:
                v = lab_for.get(i, np.nan)
                color = cmap(norm(v)) if np.isfinite(v) else (0.5, 0.5, 0.5, 1.0)  # gray if missing
            else:
                color = cmap(norm(cat_to_i[lab_for[i]]))

            ax.plot(x, y, lw=linewidth, alpha=alpha, color=color)

        ax.set_ylabel(f"μ{d+1}")
        ax.grid(True, alpha=0.25)

    axes[-1].set_xlabel("Time (step)" if dt is None else "Time")

    # colorbar
    if use_labels:
        fig.subplots_adjust(bottom=0.18)
        cax = fig.add_axes([0.12, 0.08, 0.76, 0.04])
        sm = plt.cm.ScalarMappable(norm=norm, cmap=cmap)
        sm.set_array([])
        cbar = plt.colorbar(sm, cax=cax, orientation="horizontal")
        if use_cont:
            cbar.set_label("Angle (deg)")
            if np.isclose(norm.vmin, -180) and np.isclose(norm.vmax, 180):
                cbar.set_ticks([-180, -90, 0, 90, 180])
        else:
            cbar.set_ticks(range(len(cats)))
            cbar.set_ticklabels([str(c) for c in cats])
            cbar.set_label("Trial category")

    plt.tight_layout(rect=[0.02, 0.18 if use_labels else 0.02, 0.98, 0.98])
    # plt.show()

def plot_mu_trajectories_gradient_steps_3d(
    mu_traj_np,                   # list of arrays [T_i, D]
    trial_indices=None,           # iterable of trial ids to plot; None => all
    dims=(0, 1, 2),               # which μ dims (0-based)
    start_color="#fdae61",
    end_color="#313695",
    linewidth=2.0,
    alpha=0.5,                    # line alpha
    start_marker_size=36,
    start_marker_alpha=0.95,
    end_arrow_alpha=0.95,
    arrow_head_frac=0.03,         # head length as fraction of per-trial diagonal
    arrow_length_ratio=0.6,
    title="Latent trajectories",
    fig_w=7, fig_h=6,
    decimate=1,                   # plot every k-th step along each trial (speed-up)
    z_scale='auto',               # 'auto', 'none', or a number to multiply z values by
    aspect_ratio=(1, 1, 1),       # visual aspect ratio for the box
    elev_azim=None,               # tuple (elev, azim)
    # NEW:
    labels=None,                  # per-trial values -> color each whole trajectory
    label_mode='auto',            # 'auto' | 'continuous' | 'categorical'
    cmap=None,                    # None -> Matplotlib default (e.g., 'viridis') for continuous; tab10 for categorical
    vrange=None,                  # (vmin, vmax) for continuous labels
    cbar_label=None,               # override colorbar label (continuous labels)
    circmap = 'twilight',
):
    # ----- choose trials -----
    if trial_indices is None:
        trial_indices = range(len(mu_traj_np))

    # ----- gather trajectories -----
    trajs, max_len = [], 0
    for idx in trial_indices:
        S = mu_traj_np[idx]
        if S is None or S.size == 0:
            continue
        trajs.append((idx, S))
        max_len = max(max_len, S.shape[0])
    if not trajs:
        raise ValueError("No non-empty trials to plot.")

    # ----- compute data ranges for auto-scaling -----
    all_x, all_y, all_z = [], [], []
    for _, S in trajs:
        all_x.extend(S[:, dims[0]].tolist())
        all_y.extend(S[:, dims[1]].tolist())
        all_z.extend(S[:, dims[2]].tolist())
    x_range = (max(all_x) - min(all_x)) if all_x else 1
    y_range = (max(all_y) - min(all_y)) if all_y else 1
    z_range = (max(all_z) - min(all_z)) if all_z else 1

    # ----- determine z scaling factor -----
    if z_scale == 'auto':
        xy_avg_range = (x_range + y_range) / 2
        scale_factor = (xy_avg_range / z_range) if z_range > 0 else 1.0
    elif z_scale == 'none':
        scale_factor = 1.0
    else:
        scale_factor = float(z_scale)

    # ===== label mode setup (if labels given) =====
    use_labels = labels is not None
    lab_for = {}
    use_cont = False
    cats = None
    cmap_obj = None
    norm = None

    if use_labels:
        # build per-trial label lookup
        if isinstance(labels, dict):
            for idx, _S in trajs:
                if idx in labels:
                    lab_for[idx] = labels[idx]
        else:
            arr = np.asarray(labels)
            for idx, _S in trajs:
                if idx < arr.shape[0]:
                    lab_for[idx] = arr[idx]

        vals_exist = [lab_for[i] for i, _ in trajs if i in lab_for]
        if len(vals_exist) == 0:
            raise ValueError("Provided labels are empty or misaligned with trials.")

        # decide continuous vs categorical
        if label_mode == 'continuous':
            use_cont = True
        elif label_mode == 'categorical':
            use_cont = False
        else:  # 'auto'
            all_int = all(isinstance(v, (int, np.integer)) for v in vals_exist)
            all_num = all(isinstance(v, (int, float, np.integer, np.floating)) for v in vals_exist)
            use_cont = (not all_int) and all_num

        if use_cont:
            vals = np.array(vals_exist, dtype=float)
            if vrange is None:
                vmin, vmax = float(np.nanmin(vals)), float(np.nanmax(vals))
                # if looks like degrees in [-180, 180], lock it
                if (vmin >= -180-1e-6) and (vmax <= 180+1e-6):
                    vmin, vmax = -180.0, 180.0
            else:
                vmin, vmax = vrange

            if np.isclose(vmin, -180.0) and np.isclose(vmax, 180.0) and cmap is None:
                cmap_obj = plt.get_cmap(circmap)
            else:
                default_name = plt.rcParams.get('image.cmap', 'viridis')
                cmap_obj = plt.get_cmap(default_name if cmap is None else cmap)
            
            norm = Normalize(vmin=vmin, vmax=vmax)
        else:
            # categorical palette
            cats = []
            for idx, _ in trajs:
                if idx in lab_for:
                    v = lab_for[idx]
                    if v not in cats:
                        cats.append(v)
            if cmap is None:
                base = plt.get_cmap('tab10')
                cmap_obj = ListedColormap([base(i % base.N) for i in range(len(cats))])
            else:
                cmap_obj = plt.get_cmap(cmap)
            norm = BoundaryNorm(np.arange(len(cats)+1)-0.5, len(cats))
            cat_to_i = {c: i for i, c in enumerate(cats)}
    else:
        # no labels -> we'll use a time-gradient colormap per your original code
        cmap_obj = LinearSegmentedColormap.from_list("o2b", [start_color, end_color])
        norm = Normalize(vmin=0, vmax=(1 if max_len <= 1 else max_len - 1))

    # ----- figure & axes -----
    fig = plt.figure(figsize=(fig_w, fig_h))
    ax = fig.add_subplot(111, projection="3d")

    # ----- track global data limits (with scaled z) -----
    xmin = ymin = zmin_scaled = np.inf
    xmax = ymax = zmax_scaled = -np.inf

    # ===== plot =====
    for idx, S in trajs:
        # dims + scale z
        x = np.asarray(S[:, dims[0]], dtype=float)
        y = np.asarray(S[:, dims[1]], dtype=float)
        z_raw = np.asarray(S[:, dims[2]], dtype=float)
        z = z_raw * scale_factor

        # decimate
        if decimate > 1:
            x = x[::decimate]; y = y[::decimate]; z = z[::decimate]; z_raw = z_raw[::decimate]
        T = x.shape[0]
        if T == 0:
            continue

        # update global bounds (scaled z)
        xmin = min(xmin, x.min()); xmax = max(xmax, x.max())
        ymin = min(ymin, y.min()); ymax = max(ymax, y.max())
        zmin_scaled = min(zmin_scaled, z.min()); zmax_scaled = max(zmax_scaled, z.max())

        # start marker
        if not use_labels:
            start_col = cmap_obj(norm(0))
        else:
            if idx in lab_for:
                if use_cont:
                    start_col = cmap_obj(norm(float(lab_for[idx])))
                else:
                    start_col = cmap_obj(norm(cat_to_i[lab_for[idx]]))
            else:
                start_col = (0.6, 0.6, 0.6, 1.0)  # gray when missing
        ax.scatter(x[0], y[0], z[0], s=start_marker_size, marker="o",
                   facecolor=start_col, edgecolor="black",
                   linewidths=0.5, alpha=start_marker_alpha, zorder=3)

        if T == 1:
            continue

        if not use_labels:
            # time-gradient segments
            pts = np.column_stack([x, y, z])             # (T, 3)
            segs = np.stack([pts[:-1], pts[1:]], axis=1) # (T-1, 2, 3)
            seg_steps = np.arange(segs.shape[0])         # 0..T-2
            colors = cmap_obj(norm(seg_steps))
            lc = Line3DCollection(segs, linewidths=linewidth, alpha=alpha)
            lc.set_colors(colors)
            ax.add_collection3d(lc)
            end_col = cmap_obj(norm(T-1))
        else:
            # solid color per trajectory from labels
            if idx in lab_for:
                if use_cont:
                    color = cmap_obj(norm(float(lab_for[idx])))
                else:
                    color = cmap_obj(norm(cat_to_i[lab_for[idx]]))
            else:
                color = (0.6, 0.6, 0.6, 1.0)
            ax.plot(x, y, z, lw=linewidth, alpha=alpha, color=color)
            end_col = color

        # end arrow
        dx, dy, dz = x[-1]-x[-2], y[-1]-y[-2], z[-1]-z[-2]
        rng = np.array([x.max()-x.min(), y.max()-y.min(), z.max()-z.min()])
        diag = float(np.linalg.norm(rng))
        head_len = max(1e-12, arrow_head_frac * (diag if diag > 0 else 1.0))

        uv = np.array([dx, dy, dz], float)
        nrm = float(np.linalg.norm(uv))

        # Use a slightly larger minimum so the head is visible
        min_head = 0.02 * max(rng.max(), 1e-12)   # 2% of local span
        head_len = max(head_len, min_head)
        
        # If the last step is (near) zero, just place an end marker instead of an arrow
        if nrm < 1e-12:
            ax.scatter(x[-1], y[-1], z[-1],
                       s=start_marker_size*0.9,
                       marker="^", facecolor=end_col, edgecolor="black",
                       linewidths=0.5, alpha=end_arrow_alpha, zorder=3)
        else:
            u_hat = uv / nrm
            tail = np.array([x[-1], y[-1], z[-1]]) - u_hat * head_len
            ax.quiver(
                tail[0], tail[1], tail[2],
                u_hat[0], u_hat[1], u_hat[2],
                length=head_len, normalize=False,
                color=end_col, linewidth=linewidth*1.2,
                arrow_length_ratio=arrow_length_ratio,
                alpha=end_arrow_alpha
            )

    # ----- labels & view -----
    ax.set_xlabel(f"μ{dims[0]+1}")
    ax.set_ylabel(f"μ{dims[1]+1}")
    ax.set_zlabel(f"μ{dims[2]+1}")
    ax.set_title(title)

    # limits + aspect
    # --- Let Matplotlib autoscale using the exact points we plotted (with z scaling) ---
    xs, ys, zs = [], [], []
    for idx, S in trajs:
        x = np.asarray(S[:, dims[0]], float)
        y = np.asarray(S[:, dims[1]], float)
        z = np.asarray(S[:, dims[2]], float) * scale_factor
        if decimate > 1:
            x = x[::decimate]; y = y[::decimate]; z = z[::decimate]
        if x.size:
            xs.append(x); ys.append(y); zs.append(z)
    
    if xs:  # concatenate and hand to the built-in autoscaler
        xs = np.concatenate(xs); ys = np.concatenate(ys); zs = np.concatenate(zs)
        ax.auto_scale_xyz(xs, ys, zs, had_data=False)
    
        # Optional: tiny uniform pad so points aren't glued to the box
        dx = max(1e-12, xs.max() - xs.min())
        dy = max(1e-12, ys.max() - ys.min())
        dz = max(1e-12, zs.max() - zs.min())
        pad = 0.03  # 3% visual padding
        ax.set_xlim(xs.min() - pad*dx, xs.max() + pad*dx)
        ax.set_ylim(ys.min() - pad*dy, ys.max() + pad*dy)
        ax.set_zlim(zs.min() - pad*dz, zs.max() + pad*dz)
    
    # Aspect: keep data aspect unless you explicitly pass something else
    try:
        if aspect_ratio == (1, 1, 1):
            ax.set_box_aspect('auto')   # natural data aspect (no forced cube)
        else:
            ax.set_box_aspect(aspect_ratio)
    except Exception:
        pass

    if elev_azim is not None:
        ax.view_init(elev=elev_azim[0], azim=elev_azim[1])

    # rescale tick labels on z back (if we visually scaled z)
    if scale_factor != 1.0:
        ax.zaxis.set_major_formatter(
            FuncFormatter(lambda val, pos: f"{val/scale_factor:.2f}")
        )

    # ----- colorbar -----
    sm = None
    if not use_labels:
        sm = plt.cm.ScalarMappable(cmap=cmap_obj, norm=norm); sm.set_array([])
        cbar = plt.colorbar(sm, ax=ax, fraction=0.05, pad=0.08)
        cbar.set_label("Step index")
        ticks = [0, 1] if max_len <= 2 else [0, (max_len - 1)//2, max_len - 1]
        cbar.set_ticks(ticks); cbar.set_ticklabels([str(t) for t in ticks])
    else:
        sm = plt.cm.ScalarMappable(cmap=cmap_obj, norm=norm); sm.set_array([])
        cbar = plt.colorbar(sm, ax=ax, fraction=0.05, pad=0.08)
        if use_cont:
            lbl = cbar_label if cbar_label is not None else "Label"
            # nice default for angles
            if isinstance(vrange, tuple):
                vmin, vmax = vrange
            else:
                vmin, vmax = norm.vmin, norm.vmax
            if np.isclose(vmin, -180) and np.isclose(vmax, 180) and cbar_label is None:
                lbl = "Angle (deg)"
                cbar.set_ticks([-180, -90, 0, 90, 180])
            cbar.set_label(lbl)
        else:
            cbar.set_ticks(range(len(cats)))
            cbar.set_ticklabels([str(c) for c in cats])
            cbar.set_label("Category")


def generate_traj_cov(dyn_net, dset_samples, n_step, tau, lag_samples, device, lag,
                      include_x0_tau=False, covStat_samples=None,
                      oracle=False, flow_net=None,
                      clamp_range=None):
    
    # clamp prep
    full_min = full_max = None
    if clamp_range is not None and len(clamp_range.shape) == 2:
        clamp_range = torch.from_numpy(clamp_range).to(device).float()
        full_min = clamp_range[:, 0].tile(lag + 1).unsqueeze(0)
        full_max = clamp_range[:, 1].tile(lag + 1).unsqueeze(0)

    
    # x(τ) at first point
    starting_pts = np.stack([dset_samples[ii][0, :] for ii in range(len(dset_samples))], 0)
    starting_pts = torch.from_numpy(starting_pts).unsqueeze(1).float().to(device)  # (B,1,D)
    if tau < 1.0:
        assert flow_net is not None, "flow_net required when tau<1.0."
        curr_tau_pts = util.calc_flow_trajectories(flow_net, starting_pts.squeeze(1), 1.0, tau, nt=2)[-1].unsqueeze(1)
    else:
        curr_tau_pts = starting_pts

    B, _, D = curr_tau_pts.shape
    # init τ-history lag from current τ state
    x0_tau = curr_tau_pts.squeeze(1)                    # (B,D)
    lag_cov = x0_tau.repeat(1, lag + 1) if lag > 0 else x0_tau.clone()
    curr_tau_trajs, lag_cov_list = [curr_tau_pts.cpu().numpy()], []

    for ii in tqdm(range(n_step - 1)):
        # state & lag
        if oracle:
            # lag at t -> τ if needed
            lag_cov = torch.stack([
                torch.as_tensor(lag_samples[j][ii], dtype=torch.float32, device=device)
                if ii < lag_samples[j].shape[0] else
                torch.as_tensor(lag_samples[j][-1], dtype=torch.float32, device=device)
                for j in range(len(lag_samples))
            ], dim=0)  # (B,(k+1)·D)
            if tau < 1.0:
                assert flow_net is not None, "flow_net required for oracle τ<1.0."
                L = lag + 1
                lag_cov = lag_cov.view(B * L, D)
                with torch.no_grad():
                    lag_cov = util.calc_flow_trajectories(flow_net, lag_cov, 1.0, tau, nt=2)[-1]
                lag_cov = lag_cov.view(B, L * D)
        
            # x0 at t -> τ if needed
            x0_tmp = torch.stack([
                torch.as_tensor(dset_samples[j][ii], dtype=torch.float32, device=device)
                if ii < dset_samples[j].shape[0] else
                torch.as_tensor(dset_samples[j][-1], dtype=torch.float32, device=device)
                for j in range(len(dset_samples))
            ], dim=0)  # (B,D)
            if tau < 1.0:
                with torch.no_grad():
                    x0_tmp = util.calc_flow_trajectories(flow_net, x0_tmp, 1.0, tau, nt=2)[-1]
        else:
            x0_tmp = curr_tau_pts.squeeze(1)

        # clamp lag
        if clamp_range is not None:
            if len(clamp_range.shape) == 1:
                lag_cov = lag_cov.clamp(min=clamp_range[0], max=clamp_range[1])
            else:
                lag_cov = torch.clamp(lag_cov, min=full_min, max=full_max)

        # prepend covStat if provided
        if covStat_samples is not None:
            covStat_tmp = torch.stack([
                torch.as_tensor(covStat_samples[j][ii], dtype=torch.float32, device=device)
                if ii < covStat_samples[j].shape[0] else
                torch.as_tensor(covStat_samples[j][-1], dtype=torch.float32, device=device)
                for j in range(len(covStat_samples))
            ], dim=0)
            covStat_all = torch.cat((covStat_tmp, lag_cov), dim=1)
        else:
            covStat_all = lag_cov

        # one step in τ
        traj = util.calc_dyn_trajectories(
            dyn_net, x0_tmp, tau, x0_tau=None,
            covariates=None, next_covariates=None,
            covariates_static=covStat_all,
            include_x0_tau=include_x0_tau, nt=2
        )
        x_next = traj[-1]  # (B,D)

        # clamp state
        if clamp_range is not None:
            if len(clamp_range.shape) == 1:
                x_next = x_next.clamp(min=clamp_range[0], max=clamp_range[1])
            else:
                x_next = torch.clamp(x_next, min=clamp_range[:, 0].unsqueeze(0), max=clamp_range[:, 1].unsqueeze(0))

        # push time
        curr_tau_pts = x_next.unsqueeze(1)
        curr_tau_trajs.append(curr_tau_pts.cpu().numpy())

        # roll tau
        if not oracle:
            if lag > 0:
                lag_cov = torch.cat([lag_cov[:, D:], x_next], dim=-1)
            else:
                lag_cov = x_next
        # record lag used this step
        lag_cov_list.append(lag_cov)

    curr_tau_trajs = np.concatenate(curr_tau_trajs, axis=1)
    curr_tau_trajs_trunc = [curr_tau_trajs[i, :dset_samples[i].shape[0], :] for i in range(curr_tau_trajs.shape[0])]
    return curr_tau_trajs_trunc, lag_cov_list

def proj_to_latent(dset_samples, flow_net, tau_latent, device):
    latent_traj_list = []
    for ii in tqdm(range(len(dset_samples))):
        latent_traj_tmp = util.calc_flow_trajectories(flow_net,
                                                      torch.from_numpy(dset_samples[ii]).type(torch.float32).to(device),
                                                      1.0, tau_latent, nt=2)
        latent_traj_list.append(latent_traj_tmp[-1,:,:].cpu().numpy())
    return latent_traj_list

def latent_mu(latent_traj_list, encoder, dim_preserve, device, fullRes=False,
              min_var=None, rcond=None, resid_tol=1e-10, use_fp64=True):
    enc = encoder.to(device).eval()
    mu_traj, loading = [], None

    def _flatten_Tx(z: torch.Tensor) -> torch.Tensor:
        return z.view(z.size(0), -1)

    with torch.inference_mode():

        arr0 = np.asarray(latent_traj_list[0], dtype=np.float32)
        Z0 = torch.from_numpy(arr0[:1]).to(device=device, dtype=torch.float32)
        _, D, L, bias = enc.encode(Z0, return_data_mean=True, update_stats=False)

        bias = bias.view(-1)

        # D is full diag; only first K are used
        d_vec = torch.diagonal(D) if D.ndim == 2 else D
        d_len = d_vec.numel()
        K = int(min(getattr(enc, 'k_max', d_len), d_len))

        mv = (max(1e-12, 1e-6 * torch.median(d_vec).item())
              if min_var is None else float(min_var))
        d_safe = torch.clamp(d_vec, min=mv)
        d_sqrt = torch.sqrt(d_safe[:K])               # [K]
        inv_sqrt_d = d_sqrt.reciprocal()              # [K]

        # Orthonormal columns, use only first K
        L_K = L[:, :K]                                 # [d, K]

        def solve_mu_batch(Z):
            # Z: [T,...] -> [T,d]
            Z_flat = _flatten_Tx(Z).to(device=device, dtype=torch.float32)
            Zb = Z_flat - bias.unsqueeze(0).to(Z_flat.dtype)   # [T,d]

            ZbT = Zb.T  # [d,T]
            if use_fp64:
                Ld = L_K.double()
                ZbT = ZbT.double()
                invsd = inv_sqrt_d.double()
                Y = Ld.T @ ZbT                # [K,T]
                MUt = invsd.unsqueeze(1) * Y  # [K,T]
                mu_full = MUt.T.float()       # [T,K]
            else:
                Y = L_K.T @ ZbT               # [K,T]
                MUt = inv_sqrt_d.unsqueeze(1) * Y
                mu_full = MUt.T               # [T,K]

            return mu_full  # [T,K]

        bad = False
        for arr in latent_traj_list:
            Z = torch.from_numpy(arr).to(device=device, dtype=torch.float32)
            mu_full = solve_mu_batch(Z)                     # [T,K]
            Z_flat  = _flatten_Tx(Z).to(mu_full.dtype)      # [T,d]

            # Reconstruct for residual check
            loading_now = (L_K * d_sqrt.unsqueeze(0)).to(mu_full.dtype)  # [d,K]
            Z_hat = (loading_now @ mu_full.T).T + bias.to(mu_full.dtype)  # [T,d]

            r = (Z_hat - Z_flat).norm() / (Z_flat.norm() + 1e-12)
            if torch.isnan(mu_full).any() or torch.isinf(mu_full).any() or r > resid_tol:
                bad = True
                break

            mu_traj.append(mu_full[:, :dim_preserve].cpu())

        if bad:
            # Exact left inverse on span(L_K): W = D_K^{-1/2} L_K^T
            loading = L_K * d_sqrt.unsqueeze(0)             # [d,K]
            W = (L_K.T * inv_sqrt_d.unsqueeze(1)).float()   # [K,d]
            mu_traj = []
            for arr in latent_traj_list:
                Z = torch.from_numpy(arr).to(device=device, dtype=torch.float32)
                Zf = _flatten_Tx(Z)
                Zb = Zf - bias
                mu_full = (W @ Zb.T).T                      # [T,K]
                mu_traj.append(mu_full[:, :dim_preserve].cpu())

    if fullRes:
        # return effective loading L_K sqrt(D_K)
        if loading is None:
            loading = L_K * d_sqrt.unsqueeze(0)
        return mu_traj, d_vec, L_K, loading, bias
    return mu_traj


def list_mse_equal_weight(data_list, sim_list):
    """
    Returns:
      overall_mse: float                      # mean of per-trial scalar MSEs
      per_trial:   (n_trial,) array           # scalar MSE per trial
      per_trial_dim: (n_trial, D) array       # MSE per trial per dimension
    """
    assert len(data_list) == len(sim_list), "Trial count mismatch"
    per_trial = []
    per_trial_dim = []

    D_ref = None
    for X, Y in zip(data_list, sim_list):
        X = np.asarray(X, dtype=float)
        Y = np.asarray(Y, dtype=float)
        if X.shape != Y.shape:
            raise ValueError(f"Shape mismatch in a trial: {X.shape} vs {Y.shape}")
        if D_ref is None:
            D_ref = X.shape[1]
        elif X.shape[1] != D_ref:
            raise ValueError(f"Inconsistent dims across trials: expected {D_ref}, got {X.shape[1]}")

        diff2 = (X - Y) ** 2            # [T, D]
        per_trial.append(diff2.mean())  # mean over time and dims
        per_trial_dim.append(diff2.mean(axis=0))  # mean over time -> [D]

    per_trial = np.asarray(per_trial)                  # [n_trial]
    per_trial_dim = np.vstack(per_trial_dim)          # [n_trial, D]
    overall_mse = float(per_trial.mean())             # scalar
    overall_mse_dim = np.mean(per_trial_dim, axis=0)
    
    return overall_mse, per_trial, overall_mse_dim, per_trial_dim


def mean_traj_by_labels(traj_array, label_array, round_labels=None):
    N, T, D = traj_array.shape
    labels = label_array
    if round_labels is not None:
        labels = np.round(labels.astype(float), round_labels)

    unique_labels, inv, counts = np.unique(labels, axis=0, return_inverse=True, return_counts=True)
    G = unique_labels.shape[0]

    sums = np.zeros((G, T, D), dtype=np.float64)
    np.add.at(sums, inv, traj_array)            # accumulate trial-wise
    means = sums / counts[:, None, None]        # broadcast divide

    return unique_labels, counts, means, inv


def get_theta_deg(XY):
    x = XY[:, 0]
    y = XY[:, 1]
    r = np.hypot(x, y)
    theta = np.arctan2(x, y)              # <-- note: x first, y second (swapped)
    theta = (theta + np.pi) % (2*np.pi) - np.pi
    theta[np.isclose(theta, np.pi)] = 0.0
    theta_deg = np.degrees(theta)
    return theta_deg

def model_gen(folder_data, folder, chk_pts, dim_preserve, clamp_range, device, useCov = False):
    pkl_path = folder + chk_pts
    dataset_samples = np.load(folder_data + "/dataset_samples.npz", allow_pickle=True)
    lagRead_samples = np.load(folder_data + "/lag_cov_list.npz", allow_pickle=True)
    if useCov:
        cov_static_all = np.load(folder_data + "/cov_static_new.npz", allow_pickle=True)
    
    data_raw = dataset_samples['samples']
    lag_raw = lagRead_samples['samples']
    dset_samples = []
    lag_samples = []

    if useCov:
        cov_static_raw = cov_static_all['samples']
        covStat_samples = []
    
    for ii in range(len(data_raw)):
        dset_samples.append(np.asarray(data_raw[ii], dtype=np.float64))
        lag_samples.append(np.asarray(lag_raw[ii], dtype=np.float64))
        if useCov:
            covStat_samples.append(np.asarray(cov_static_raw[ii][:, 0:(cov_static_raw[ii].shape[1] - lag_raw[ii].shape[1])], dtype=np.float64))
    
    
    lag = lag_samples[0].shape[1] // dset_samples[0].shape[1] - 1
    T_min = min(arr.shape[0] for arr in dset_samples)
    T_max = max(arr.shape[0] for arr in dset_samples)
    
    # with open(pkl_path, 'rb') as f:
    #     model_all = pickle.load(f)
    # if 'ema' in model_all: net = model_all['ema']  # EMA-stabilized model (recommended for inference/simulation)
    # else: net = model_all['net']  # Raw trained model
    # flow_net = net.unet_model  # Compressive flow (u_theta)
    # dyn_net = net.vnet_model   # Dynamics flow (v_theta)
    # encoder = net.encoder      # For latent proposals


    if not hasattr(persistence, "_orig_reconstruct"):
        persistence._orig_reconstruct = persistence._reconstruct_persistent_obj
    def _tolerant_reconstruct(meta):
        try:
            return persistence._orig_reconstruct(meta)
        except Exception:
            class _Stub: pass
            return _Stub()
    class PatchedUnpickler(pickle.Unpickler):
        def find_class(self, module, name):
            if module == "torch_utils.persistence" and name == "_reconstruct_persistent_obj":
                return _tolerant_reconstruct
            return super().find_class(module, name)
    with open(pkl_path, "rb") as f:
        model_all = PatchedUnpickler(f).load()

    net = model_all["ema"] if "ema" in model_all else model_all["net"]
    flow_net = net.unet_model  # Compressive flow (u_theta)
    dyn_net = net.vnet_model   # Dynamics flow (v_theta)
    encoder = net.encoder
    
    _, d, L, loading, bias = latent_mu(dset_samples, encoder, dim_preserve, device, fullRes = True)
    
    if useCov:
        traj_sim_tau1, lag_cov_list = generate_traj_cov(dyn_net.to(device), dset_samples,
                                                        n_step = T_max, tau = 1.0,
                                                        lag_samples = lag_samples,
                                                        device = device, lag = lag,
                                                        include_x0_tau = False,
                                                        covStat_samples = covStat_samples,
                                                        oracle = False,
                                                        clamp_range = clamp_range)
        
        traj_sim_tau0, _ = generate_traj_cov(dyn_net.to(device), dset_samples,
                                             n_step = T_max, tau = 0.0,
                                             lag_samples = lag_samples,
                                             device = device, lag = lag,
                                             include_x0_tau = False,
                                             covStat_samples = covStat_samples,
                                             oracle = False,
                                             flow_net = flow_net.to(device),
                                             clamp_range = clamp_range)
    else:
        traj_sim_tau1, lag_cov_list = generate_traj(dyn_net.to(device), dset_samples,
                                                    n_step = T_max, tau = 1.0,
                                                    lag_samples = lag_samples,
                                                    device = device, lag = lag,
                                                    include_x0_tau = False,
                                                    oracle = False,
                                                    clamp_range = clamp_range)
        
        traj_sim_tau0, _ = generate_traj(dyn_net.to(device), dset_samples,
                                         n_step = T_max, tau = 0.0,
                                         lag_samples = lag_samples, device = device, lag = lag,
                                         include_x0_tau = False,
                                         oracle = False,
                                         flow_net = flow_net.to(device),
                                         clamp_range = clamp_range)

    if useCov:
        return dset_samples, lag_samples, covStat_samples, flow_net, dyn_net, encoder, d, L, loading, bias, traj_sim_tau1, traj_sim_tau0
    else:
        return dset_samples, lag_samples, flow_net, dyn_net, encoder, d, L, loading, bias, traj_sim_tau1, traj_sim_tau0


def plot_mu_trajectories_gradient_steps_2d(
    mu_traj_np,                   # list of arrays [T_i, D]
    trial_indices=None,           # iterable of trial ids to plot; None => all
    dims=(0, 1),                  # which μ dims (0-based)
    start_color="#fdae61",
    end_color="#313695",
    linewidth=2.0,
    alpha=0.5,                    # line alpha
    start_marker_size=36,
    start_marker_alpha=0.95,
    end_arrow_alpha=0.95,
    arrow_head_frac=0.03,         # head length as fraction of per-trial diagonal (data units)
    arrow_length_ratio=0.6,       # kept for API symmetry (not used by quiver directly)
    title="Latent trajectories (2D)",
    fig_w=6, fig_h=6,
    decimate=1,                   # plot every k-th step along each trial (speed-up)
    aspect_equal=False,            # equal axes by default
    # Labels (same semantics as your 3D version)
    labels=None,                  # per-trial values -> color each whole trajectory
    label_mode='auto',            # 'auto' | 'continuous' | 'categorical'
    cmap=None,                    # None -> default (e.g., 'viridis') for continuous; tab10 for categorical
    vrange=None,                  # (vmin, vmax) for continuous labels
    cbar_label=None,              # override colorbar label (continuous labels)
    circmap='twilight'            # used for angles if [-180, 180]
):
    # ----- choose trials -----
    if trial_indices is None:
        trial_indices = range(len(mu_traj_np))

    # ----- gather trajectories -----
    trajs, max_len = [], 0
    for idx in trial_indices:
        S = mu_traj_np[idx]
        if S is None or getattr(S, "size", 0) == 0:
            continue
        trajs.append((idx, S))
        max_len = max(max_len, S.shape[0])
    if not trajs:
        raise ValueError("No non-empty trials to plot.")

    # ===== label mode setup (if labels given) =====
    use_labels = labels is not None
    lab_for = {}
    use_cont = False
    cats = None
    cmap_obj = None
    norm = None

    if use_labels:
        # map trial index -> label
        if isinstance(labels, dict):
            for idx, _S in trajs:
                if idx in labels:
                    lab_for[idx] = labels[idx]
        else:
            arr = np.asarray(labels)
            for idx, _S in trajs:
                if idx < arr.shape[0]:
                    lab_for[idx] = arr[idx]

        vals_exist = [lab_for[i] for i, _ in trajs if i in lab_for]
        if len(vals_exist) == 0:
            raise ValueError("Provided labels are empty or misaligned with trials.")

        # decide continuous vs categorical
        if label_mode == 'continuous':
            use_cont = True
        elif label_mode == 'categorical':
            use_cont = False
        else:  # 'auto'
            all_int = all(isinstance(v, (int, np.integer)) for v in vals_exist)
            all_num = all(isinstance(v, (int, float, np.integer, np.floating)) for v in vals_exist)
            use_cont = (not all_int) and all_num

        if use_cont:
            vals = np.array(vals_exist, dtype=float)
            if vrange is None:
                vmin, vmax = float(np.nanmin(vals)), float(np.nanmax(vals))
                # lock to degrees if it fits [-180, 180]
                if (vmin >= -180-1e-6) and (vmax <= 180+1e-6):
                    vmin, vmax = -180.0, 180.0
            else:
                vmin, vmax = vrange

            if np.isclose(vmin, -180.0) and np.isclose(vmax, 180.0) and cmap is None:
                cmap_obj = plt.get_cmap(circmap)
            else:
                default_name = plt.rcParams.get('image.cmap', 'viridis')
                cmap_obj = plt.get_cmap(default_name if cmap is None else cmap)

            norm = Normalize(vmin=vmin, vmax=vmax)
        else:
            # categorical palette
            cats = []
            for idx, _ in trajs:
                if idx in lab_for:
                    v = lab_for[idx]
                    if v not in cats:
                        cats.append(v)
            if cmap is None:
                base = plt.get_cmap('tab10')
                cmap_obj = ListedColormap([base(i % base.N) for i in range(len(cats))])
            else:
                cmap_obj = plt.get_cmap(cmap)
            norm = BoundaryNorm(np.arange(len(cats)+1)-0.5, len(cats))
            cat_to_i = {c: i for i, c in enumerate(cats)}
    else:
        # no labels -> time-gradient colormap per trajectory
        cmap_obj = LinearSegmentedColormap.from_list("o2b", [start_color, end_color])
        norm = Normalize(vmin=0, vmax=(1 if max_len <= 1 else max_len - 1))

    # ----- figure & axes -----
    fig, ax = plt.subplots(figsize=(fig_w, fig_h))

    # ----- global limits -----
    xmin = ymin = np.inf
    xmax = ymax = -np.inf

    # ===== plot =====
    for idx, S in trajs:
        x = np.asarray(S[:, dims[0]], dtype=float)
        y = np.asarray(S[:, dims[1]], dtype=float)

        # decimate
        if decimate > 1:
            x = x[::decimate]
            y = y[::decimate]
        T = x.shape[0]
        if T == 0:
            continue

        # update bounds
        xmin = min(xmin, np.min(x))
        xmax = max(xmax, np.max(x))
        ymin = min(ymin, np.min(y))
        ymax = max(ymax, np.max(y))

        # start marker color
        if not use_labels:
            start_col = cmap_obj(norm(0))
        else:
            if idx in lab_for:
                if use_cont:
                    start_col = cmap_obj(norm(float(lab_for[idx])))
                else:
                    start_col = cmap_obj(norm(cat_to_i[lab_for[idx]]))
            else:
                start_col = (0.6, 0.6, 0.6, 1.0)
        ax.scatter(x[0], y[0], s=start_marker_size, marker="o",
                   facecolor=start_col, edgecolor="black",
                   linewidths=0.5, alpha=start_marker_alpha, zorder=3)

        if T == 1:
            continue

        if not use_labels:
            # time-gradient colored segments
            pts = np.column_stack([x, y])               # (T, 2)
            segs = np.stack([pts[:-1], pts[1:]], axis=1)# (T-1, 2, 2)
            seg_steps = np.arange(segs.shape[0])        # 0..T-2
            colors = cmap_obj(norm(seg_steps))
            lc = LineCollection(segs, linewidths=linewidth, alpha=alpha)
            lc.set_colors(colors)
            ax.add_collection(lc)
            end_col = cmap_obj(norm(T-1))
        else:
            # solid color per trajectory
            if idx in lab_for:
                if use_cont:
                    color = cmap_obj(norm(float(lab_for[idx])))
                else:
                    color = cmap_obj(norm(cat_to_i[lab_for[idx]]))
            else:
                color = (0.6, 0.6, 0.6, 1.0)
            ax.plot(x, y, lw=linewidth, alpha=alpha, color=color)
            end_col = color

        # end arrow
        dx, dy = x[-1] - x[-2], y[-1] - y[-2]
        uv = np.array([dx, dy], float)
        nrm = float(np.linalg.norm(uv))

        # compute diagonal span for head length in data units
        rng = np.array([x.max()-x.min(), y.max()-y.min()], dtype=float)
        diag = float(np.linalg.norm(rng))
        head_len = max(1e-12, arrow_head_frac * (diag if diag > 0 else 1.0))

        # ensure visibility (2% of local span)
        min_head = 0.02 * max(rng.max(), 1e-12)
        head_len = max(head_len, min_head)

        if nrm < 1e-12:
            ax.scatter(x[-1], y[-1],
                       s=start_marker_size*0.9, marker="^",
                       facecolor=end_col, edgecolor="black",
                       linewidths=0.5, alpha=end_arrow_alpha, zorder=3)
        else:
            u_hat = uv / nrm
            # draw arrow of length=head_len in data units
            ax.quiver(
                x[-1] - u_hat[0]*head_len, y[-1] - u_hat[1]*head_len,
                u_hat[0]*head_len, u_hat[1]*head_len,
                angles='xy', scale_units='xy', scale=1.0,
                color=end_col, alpha=end_arrow_alpha, width=0.003
            )

    # ----- axes cosmetics -----
    ax.set_xlabel(f"μ{dims[0]+1}")
    ax.set_ylabel(f"μ{dims[1]+1}")
    ax.set_title(title)
    # limits + aspect
    if not np.isfinite([xmin, xmax, ymin, ymax]).all():
        xmin, xmax, ymin, ymax = -1, 1, -1, 1
    pad_x = 0.02 * max(1e-12, xmax - xmin)
    pad_y = 0.02 * max(1e-12, ymax - ymin)
    ax.set_xlim(xmin - pad_x, xmax + pad_x)
    ax.set_ylim(ymin - pad_y, ymax + pad_y)
    if aspect_equal:
        ax.set_aspect("equal", adjustable="datalim")
    ax.grid(True, alpha=0.25)

    # ----- colorbar -----
    sm = None
    if not use_labels:
        sm = plt.cm.ScalarMappable(cmap=cmap_obj, norm=norm); sm.set_array([])
        cbar = plt.colorbar(sm, ax=ax, fraction=0.05, pad=0.04)
        cbar.set_label("Step index")
        ticks = [0, 1] if max_len <= 2 else [0, (max_len - 1)//2, max_len - 1]
        cbar.set_ticks(ticks); cbar.set_ticklabels([str(t) for t in ticks])
    else:
        sm = plt.cm.ScalarMappable(cmap=cmap_obj, norm=norm); sm.set_array([])
        if use_cont:
            cbar = plt.colorbar(sm, ax=ax, fraction=0.05, pad=0.04)
            lbl = cbar_label if cbar_label is not None else "Label"
            if isinstance(vrange, tuple):
                vmin, vmax = vrange
            else:
                vmin, vmax = norm.vmin, norm.vmax
            if np.isclose(vmin, -180) and np.isclose(vmax, 180) and cbar_label is None:
                lbl = "Angle (deg)"
                cbar.set_ticks([-180, -90, 0, 90, 180])
            cbar.set_label(lbl)
        else:
            # bottom categorical legend-style colorbar (like your PCA example)
            fig.subplots_adjust(bottom=0.2)
            cax = fig.add_axes([0.12, 0.08, 0.76, 0.05])  # [left, bottom, width, height]
            cbar = fig.colorbar(sm, cax=cax, orientation="horizontal")
            cbar.set_ticks(range(len(cats)))
            cbar.set_ticklabels([str(c) for c in cats])
            cbar.set_label("Category")

    return fig, ax


def compute_mu_velocity_field(
    latent_traj_list,
    dyn_net,
    encoder,
    device,
    dim_preserve,
    lag=0,
    t_plot=0.0,
    rcond=1e-6,
    return_full=False,
):

    # ---------- 1) Z -> μ and loading from encoder ----------
    # mu_traj: list of [T_j, dim_preserve] tensors (on CPU)
    # loading: L√D flattened-compatible matrix
    mu_traj, d_vec, L, loading, bias = latent_mu(
        latent_traj_list,
        encoder,
        dim_preserve,
        device,
        fullRes=True,
    )

    # ---------- 2) Build eval set (Z_t, history h_t, μ_t) ----------
    Z_eval_list = []
    cov_static_list = []
    mu_eval_list = []
    
    for j, Z_traj in enumerate(latent_traj_list):
        # Z_traj can be [T, D], [T, H, W], or [T, C, H, W]
        Z_arr = np.asarray(Z_traj, dtype=np.float32)
        T_j = Z_arr.shape[0]
        Z_flat = Z_arr.reshape(T_j, -1)          # [T_j, D_flat]
        mu_traj_j = mu_traj[j].cpu().numpy()     # [T_j, dim_preserve]
        assert mu_traj_j.shape[0] == T_j
    
        for t in range(T_j):
            # need full history when lag > 0
            if lag > 0 and t < lag:
                continue
    
            Z_t = Z_flat[t]  # [D_flat]
    
            if lag == 0:
                # lag-0: history = current Z_t
                hist = Z_t.copy()                # [D_flat]
            else:
                # lag-k: history = [Z_{t-k}, ..., Z_t] flattened
                frames = Z_flat[t - lag : t + 1] # [lag+1, D_flat]
                hist = frames.reshape(-1)        # [(lag+1)*D_flat]
    
            Z_eval_list.append(Z_t)
            cov_static_list.append(hist)
            mu_eval_list.append(mu_traj_j[t, :dim_preserve])

    Z_eval_np      = np.stack(Z_eval_list, axis=0)        # [N_eval, D]
    cov_static_np  = np.stack(cov_static_list, axis=0)    # [N_eval, cov_dim]
    mu_eval_np     = np.stack(mu_eval_list, axis=0)       # [N_eval, dim_preserve]

    # ---------- 3) Evaluate v_Z via dyn_torch_wrapper at τ = 0 ----------
    Z_eval = torch.from_numpy(Z_eval_np).to(device=device, dtype=torch.float32)
    cov_static = torch.from_numpy(cov_static_np).to(device=device, dtype=torch.float32)

    dyn_net = dyn_net.to(device).eval()
    with torch.no_grad():
        dyn_wrapper = util.dyn_torch_wrapper(
            dyn_net,
            tau=0.0,                 # Z is at τ = 0
            x0_tau=None,
            cov_start=None,
            cov_delta=None,
            include_x0_tau=False,
            cov_static=cov_static,   # history (lag-0 or lag-k)
        )
        t = torch.tensor(float(t_plot), device=device, dtype=torch.float32)
        v_Z = dyn_wrapper(t, Z_eval)
        vZ_np = v_Z.detach().cpu().numpy()
        speed_full_np = np.linalg.norm(vZ_np, axis=1)
        
    # ---------- 5) Map v_Z -> v_μ using pseudo-inverse of loading ----------
    loading_t = loading.to(device=device, dtype=torch.float32)
    if loading_t.ndim > 2:
        # flatten spatial dims for images, etc.
        loading_t = loading_t.view(-1, loading_t.shape[-1])  # [D_flat, K_full]

    W = torch.linalg.pinv(loading_t, rcond=float(rcond))     # [K_full, D_flat]
    v_mu_full = (W @ v_Z.T).T                                # [N_eval, K_full]
    v_mu_np = v_mu_full[:, :dim_preserve].detach().cpu().numpy()

    if return_full:
        return mu_eval_np, v_mu_np, speed_full_np
    else:
        return mu_eval_np, v_mu_np


def plot_mu_velocity_2d(
    mu,
    v_mu,
    dims=(0, 1),
    trajectories=None,
    title=None,
    arrow_frac=0.05,
    arrow_size=1.0,
    max_points=5000,
    linewidth=1.0,
    traj_alpha=0.4,
    traj_width=0.8,
    traj_color='gray',
    max_trajs=500000,
):
    d1, d2 = dims
    coords = mu[:, [d1, d2]]
    vecs   = v_mu[:, [d1, d2]]

    N = coords.shape[0]
    if max_points is not None and N > max_points:
        idx = np.random.choice(N, size=max_points, replace=False)
        coords = coords[idx]
        vecs   = vecs[idx]

    speed = np.linalg.norm(vecs, axis=1)
    eps   = 1e-8
    dirs  = vecs / (speed[:, None] + eps)

    # Calculate arrow lengths
    x_min, x_max = coords[:, 0].min(), coords[:, 0].max()
    y_min, y_max = coords[:, 1].min(), coords[:, 1].max()
    span = max(x_max - x_min, y_max - y_min) + 1e-8
    L = arrow_frac * span

    U = dirs[:, 0] * L
    V = dirs[:, 1] * L

    # Visual size configuration
    base_width          = 0.004
    base_headwidth      = 4.0
    base_headlength     = 6.0
    base_headaxislength = 5.0

    width          = base_width * arrow_size
    headwidth      = base_headwidth * arrow_size
    headlength     = base_headlength * arrow_size
    headaxislength = base_headaxislength * arrow_size

    fig, ax = plt.subplots(figsize=(5, 5))

    if trajectories is not None:
        if hasattr(trajectories, 'shape') and len(trajectories.shape) == 3:
            traj_list = trajectories
        else:
            traj_list = trajectories

        count = 0
        for traj in traj_list:
            if count >= max_trajs: break
            t_np = traj.detach().cpu().numpy() if hasattr(traj, 'detach') else np.array(traj)
            
            ax.plot(
                t_np[:, d1], t_np[:, d2],
                color=traj_color,
                alpha=traj_alpha,
                linewidth=traj_width,
                zorder=1
            )
            count += 1

    norm = Normalize(vmin=float(speed.min()), vmax=float(speed.max()))
    cmap = cm.viridis

    q = ax.quiver(
        coords[:, 0], coords[:, 1],
        U, V,
        speed,
        cmap=cmap,
        norm=norm,
        angles="xy",
        scale_units="xy",
        scale=1.0,
        width=width,
        headwidth=headwidth,
        headlength=headlength,
        headaxislength=headaxislength,
        linewidth=linewidth,
        alpha=0.9,
        zorder=2
    )

    cb = fig.colorbar(q, ax=ax)
    cb.set_label(r"$\|\dot{\mu}\|$ (projection)")

    ax.set_xlabel(fr"$\mu_{{{d1+1}}}$")
    ax.set_ylabel(fr"$\mu_{{{d2+1}}}$")
    if title is not None:
        ax.set_title(title)
    ax.axis("equal")
    plt.tight_layout()
    plt.show()

def plot_mu_velocity_3d(
    mu,
    v_mu,
    dims=(0, 1, 2),
    trajectories=None,
    title=None,
    arrow_frac=0.1,
    arrow_size=1.0,
    max_points=3000,
    linewidth=1.5,
    base_arrow_length_ratio=0.3,
    traj_alpha=0.4,
    traj_width=0.8,
    traj_color='gray',
    max_trajs=300,
):
    d1, d2, d3 = dims
    coords = mu[:, [d1, d2, d3]]
    vecs   = v_mu[:, [d1, d2, d3]]

    N = coords.shape[0]
    if max_points is not None and N > max_points:
        idx = np.random.choice(N, size=max_points, replace=False)
        coords = coords[idx]
        vecs   = vecs[idx]

    speed = np.linalg.norm(vecs, axis=1)
    eps   = 1e-8
    dirs  = vecs / (speed[:, None] + eps)

    mins = coords.min(axis=0)
    maxs = coords.max(axis=0)
    span = float(np.max(maxs - mins) + 1e-8)
    L = arrow_frac * span

    U = dirs[:, 0]
    V = dirs[:, 1]
    W = dirs[:, 2]

    arrow_length_ratio = base_arrow_length_ratio * arrow_size
    linew_eff = linewidth * arrow_size

    fig = plt.figure(figsize=(6, 5))
    ax  = fig.add_subplot(111, projection="3d")

    if trajectories is not None:
        if hasattr(trajectories, 'shape') and len(trajectories.shape) == 3:
            traj_list = trajectories
        else:
            traj_list = trajectories
            
        count = 0
        for traj in traj_list:
            if count >= max_trajs: break
            t_np = traj.detach().cpu().numpy() if hasattr(traj, 'detach') else np.array(traj)
            
            ax.plot(
                t_np[:, d1], t_np[:, d2], t_np[:, d3],
                color=traj_color,
                alpha=traj_alpha,
                linewidth=traj_width,
                zorder=1
            )
            count += 1

    norm   = Normalize(vmin=float(speed.min()), vmax=float(speed.max()))
    colors = cm.viridis(norm(speed))

    ax.quiver(
        coords[:, 0], coords[:, 1], coords[:, 2],
        U, V, W,
        length=L,
        normalize=False,
        colors=colors,
        linewidth=linew_eff,
        arrow_length_ratio=arrow_length_ratio,
        zorder=2
    )

    mappable = cm.ScalarMappable(norm=norm, cmap="viridis")
    mappable.set_array(speed)
    cb = fig.colorbar(mappable, ax=ax)
    cb.set_label(r"$\|\dot{\mu}\|$ (projection)")

    ax.set_xlabel(fr"$\mu_{{{d1+1}}}$")
    ax.set_ylabel(fr"$\mu_{{{d2+1}}}$")
    ax.set_zlabel(fr"$\mu_{{{d3+1}}}$")
    if title is not None:
        ax.set_title(title)

    plt.tight_layout()
    plt.show()

def plot_mu_velocity_panel(
    mu,
    v_mu,
    trajectories=None,
    dims_2d=(0, 1),
    dims_3d=(0, 1, 2),
    title_2d=None,
    title_3d=None,
    speed_color=None,
    colorbar_label=r"$\|\dot{\mu}\|$",
    # 2D params
    arrow_frac_2d=0.05,
    arrow_size_2d=1.2,
    linewidth_2d=1.5,
    # 3D params
    arrow_frac_3d=0.1,
    arrow_size_3d=1.5,
    linewidth_3d=1.5,
    base_arrow_length_ratio_3d=0.3,
    # Trajectory params
    traj_alpha=0.4,
    traj_width=0.8,
    traj_color='gray',
    max_trajs=300,
    # Shared
    max_points=3000,
    figsize=(14, 6)
):
    fig = plt.figure(figsize=figsize)
    gs = gridspec.GridSpec(1, 2, width_ratios=[1, 1.5], figure=fig)
    
    traj_list = []
    if trajectories is not None:
        if hasattr(trajectories, 'shape') and len(trajectories.shape) == 3:
            traj_source = trajectories
        else:
            traj_source = trajectories
        
        for i, traj in enumerate(traj_source):
            if i >= max_trajs: break
            t_np = traj.detach().cpu().numpy() if hasattr(traj, 'detach') else np.array(traj)
            traj_list.append(t_np)

    if speed_color is not None:
        speed_color = speed_color.detach().cpu().numpy() if hasattr(speed_color, "detach") else np.asarray(speed_color)
        assert speed_color.shape[0] == mu.shape[0]
        vmin_c = float(speed_color.min())
        vmax_c = float(speed_color.max())

    
    ax1 = fig.add_subplot(gs[0, 0])
    d1, d2 = dims_2d
    
    for t_np in traj_list:
        ax1.plot(
            t_np[:, d1], t_np[:, d2],
            color=traj_color, alpha=traj_alpha, linewidth=traj_width, zorder=1
        )

    coords = mu[:, [d1, d2]]
    vecs   = v_mu[:, [d1, d2]]
    N = coords.shape[0]
    
    if max_points is not None and N > max_points:
        idx = np.random.choice(N, size=max_points, replace=False)
        coords_2d = coords[idx]
        vecs_2d   = vecs[idx]
    else:
        coords_2d = coords
        vecs_2d   = vecs

    # speed = np.linalg.norm(vecs_2d, axis=1)
    speed_proj = np.linalg.norm(vecs_2d, axis=1)  # for directions only
    speed_c = speed_color[idx] if (speed_color is not None and max_points is not None and N > max_points) else speed_color
    if speed_color is None:
        speed_c = speed_proj
    eps   = 1e-8
    dirs  = vecs_2d / (speed_proj[:, None] + eps)
    
    x_min, x_max = coords_2d[:, 0].min(), coords_2d[:, 0].max()
    y_min, y_max = coords_2d[:, 1].min(), coords_2d[:, 1].max()
    span = max(x_max - x_min, y_max - y_min) + 1e-8
    L = arrow_frac_2d * span
    U = dirs[:, 0] * L
    V = dirs[:, 1] * L

    base_width = 0.004; base_headwidth = 4.0; base_headlength = 6.0; base_headaxislength = 5.0
    width = base_width * arrow_size_2d
    
    norm = Normalize(vmin=vmin_c, vmax=vmax_c) if speed_color is not None else Normalize(vmin=float(speed_proj.min()), vmax=float(speed_proj.max()))

    q = ax1.quiver(
        coords_2d[:, 0], coords_2d[:, 1], U, V, speed_c,
        cmap=cm.viridis, norm=norm, angles="xy", scale_units="xy", scale=1.0,
        width=width, headwidth=base_headwidth*arrow_size_2d, 
        headlength=base_headlength*arrow_size_2d, 
        headaxislength=base_headaxislength*arrow_size_2d,
        linewidth=linewidth_2d, alpha=0.9, zorder=2
    )
    cb1 = fig.colorbar(q, ax=ax1, fraction=0.046, pad=0.04)
    cb1.set_label(colorbar_label)
    ax1.set_xlabel(fr"$\mu_{{{d1+1}}}$"); ax1.set_ylabel(fr"$\mu_{{{d2+1}}}$")
    if title_2d: ax1.set_title(title_2d)
    ax1.axis("equal")

    ax2 = fig.add_subplot(gs[0, 1], projection="3d")
    ax2.set_box_aspect((1, 1, 1))
    d1, d2, d3 = dims_3d

    for t_np in traj_list:
        ax2.plot(
            t_np[:, d1], t_np[:, d2], t_np[:, d3],
            color=traj_color, alpha=traj_alpha, linewidth=traj_width, zorder=1
        )

    coords = mu[:, [d1, d2, d3]]
    vecs   = v_mu[:, [d1, d2, d3]]
    
    if max_points is not None and N > max_points:
        idx = np.random.choice(N, size=max_points, replace=False)
        coords_3d = coords[idx]
        vecs_3d   = vecs[idx]
    else:
        coords_3d = coords
        vecs_3d   = vecs

    speed_proj = np.linalg.norm(vecs_3d, axis=1)
    dirs  = vecs_3d / (speed_proj[:, None] + eps)

    speed_c = speed_color[idx] if (speed_color is not None and max_points is not None and N > max_points) else speed_color
    if speed_color is None:
        speed_c = speed_proj
    mins = coords_3d.min(axis=0); maxs = coords_3d.max(axis=0)
    span = float(np.max(maxs - mins) + 1e-8)
    L = arrow_frac_3d * span
    U, V, W = dirs[:, 0], dirs[:, 1], dirs[:, 2]

    norm = Normalize(vmin=vmin_c, vmax=vmax_c) if speed_color is not None else Normalize(vmin=float(speed_proj.min()), vmax=float(speed_proj.max()))
    colors = cm.viridis(norm(speed_c))
    
    ax2.quiver(
        coords_3d[:, 0], coords_3d[:, 1], coords_3d[:, 2],
        U, V, W, length=L, normalize=False, colors=colors,
        linewidth=linewidth_3d * arrow_size_3d,
        arrow_length_ratio=base_arrow_length_ratio_3d * arrow_size_3d,
        zorder=2
    )
    
    mappable = cm.ScalarMappable(norm=norm, cmap="viridis")
    mappable.set_array(speed_c)
    cb2 = fig.colorbar(mappable, ax=ax2, fraction=0.035, pad=0.02)
    cb2.set_label(colorbar_label)
    ax2.set_xlabel(fr"$\mu_{{{d1+1}}}$"); ax2.set_ylabel(fr"$\mu_{{{d2+1}}}$"); ax2.set_zlabel(fr"$\mu_{{{d3+1}}}$")
    if title_3d: ax2.set_title(title_3d)

    plt.tight_layout()
    plt.show()
```
