## Supplementary material for "Dynamic Compression Flows for Neuroscience Data": Python code: __init__.py

```
# This work is licensed under a Creative Commons
# Attribution-NonCommercial-ShareAlike 4.0 International License.
# You should have received a copy of the license along with this
# work. If not, see http://creativecommons.org/licenses/by-nc-sa/4.0/

from .util import EasyDict, make_cache_dir_path
from .util_v2 import EasyDict, make_cache_dir_path
from .util_v4 import EasyDict, make_cache_dir_path
from .util_v5 import EasyDict, make_cache_dir_path
from .util_v6 import EasyDict, make_cache_dir_path
from .util_v7 import EasyDict, make_cache_dir_path

# from .util_v5 import EasyDict, make_cache_dir_path
```
