## Supplementary material for "Dynamic Compression Flows for Neuroscience Data": Python code: nlb_util_v2.py

```
import h5py
import remfile
import os
from dandi.download import download
from dandi.dandiapi import DandiAPIClient
from pynwb import NWBHDF5IO
from nlb_tools.nwb_interface import NWBDataset
from tqdm import tqdm
import numpy as np
from dnnlib.util import ToyDsetDynamics
from torch.utils.data import DataLoader
from scipy.io import loadmat
from skimage.transform import resize


def validate_metadata(data,metadata):
    """
    just checks metadata against data to ensure
    metadata has the number of trials per split as
    data
    """

    for split in data.keys():
        d = data[split]
        md = metadata[split]['behavior']
        for key in md.keys():
            print(f"checking {split}: {key}")
            assert len(md[key]) == len(d)
        md = metadata[split]['trial_info']
        for key in md.keys():
            print(f"checking {split}: {key}")
            assert len(md[key]) == len(d)

    return

def load_nwb_stream(id,filepath):

    with DandiAPIClient() as client:
        asset = client.get_dandiset(id, 'draft').get_asset_by_path(filepath)
        s3_url = asset.get_content_url(follow_redirects=1, strip_query=True)
    
        rem_file = remfile.File(s3_url)
        file = h5py.File(rem_file, "r")
        io_stream = NWBHDF5IO(file=file)
        nwbfile_stream = io_stream.read()

    return nwbfile_stream,io_stream

##### Loaders for Motor Cortex recordings -- Maze (center out reaching with barriers) task ######

def loader_mc_maze_with_behavior(spike_smooth_ms = 40, nForward = 1, num_workers = 1, batch_size = 128):
    """
    this dataset is very large!!  use caution when loading,
      it takes up all RAM on gungnir when spike smoothing
    """

    dandiset_id = '000128' ### name of the dataset on DANDI archive (should be in the notebook example)
    filepath= "sub-Jenkins/sub-Jenkins_ses-full_desc-train_behavior+ecephys.nwb" ### filepath on dandi archive
                                                            ### accessible by searching the dandi ID 
                                                            ### on DANDI archive https://dandiarchive.org/
    with DandiAPIClient() as client:
        asset = client.get_dandiset(dandiset_id, 'draft').get_asset_by_path(filepath)
        s3_url = asset.get_content_url(follow_redirects=1, strip_query=True)

        rem_file = remfile.File(s3_url)
        file = h5py.File(rem_file, "r")
        io_stream = NWBHDF5IO(file=file)
        nwbfile_stream = io_stream.read()
        dataset = NWBDataset(nwbfile_stream, split_heldout=True)

    dataset.smooth_spk(spike_smooth_ms, name='smth')
    trial_start_times = dataset.trial_info.start_time.dt.total_seconds().to_numpy()
    trial_end_times = dataset.trial_info.end_time.dt.total_seconds().to_numpy()
    smoothed_spikes = dataset.data.spikes_smth
    smoothed_spikes_numpy = smoothed_spikes.to_numpy()
    smoothed_spikes['time_stamp'] = smoothed_spikes.index.total_seconds()
    train_val_label = dataset.trial_info.split.to_numpy()
    splitted_data = {
        'train':[],
        'val':[],
    }
    data_timestamp_interval = (dataset.data.index[1] - dataset.data.index[0]).total_seconds()

    trial_info_names = ['trial_type', 'start_time', 'stop_time', 'trial_version', 'maze_id', 'success',
                        'target_on_time', 'go_cue_time', 'move_onset_time', 
                        'rt', 'delay', 'num_targets', 'target_pos',
                        'num_barriers', 'barrier_pos', 'active_target']
    behavior_names = ['cursor_pos', 'eye_pos', 'hand_pos', 'hand_vel']
    metadata = {'train':{
                        'trial_info':{name:[] for name in trial_info_names},
                        'behavior':{name:[] for name in behavior_names}
                        },
               'val':{
                        'trial_info':{name:[] for name in trial_info_names},
                        'behavior':{name:[] for name in behavior_names}
                        }}


    for trial_ind, (trial_start, trial_end, label) in enumerate(zip(trial_start_times, trial_end_times, train_val_label)):
        in_trial_index = (smoothed_spikes.time_stamp >= trial_start) & (smoothed_spikes.time_stamp <= trial_end)
        in_trial_data = smoothed_spikes_numpy[in_trial_index, :]
        isnan = np.isnan(in_trial_data[:, 0])

        lead_end = np.argmax(~isnan)

        tail_start_reverse = np.argmax(~isnan[::-1])
        tail_start = len(in_trial_data[:, 0]) - tail_start_reverse  
        nan_index_single_trial = [lead_end, tail_start]

        # cell_same = np.all(nan_index_single_trial == nan_index_single_trial[0], axis=1)
        # all_same = np.all(cell_same == 1)#just for checking of the nan ends and start at the same index in each trial
        
        excluded_nan_single_trial = in_trial_data[nan_index_single_trial[0]:nan_index_single_trial[1], :]
        #print(np.sum(np.isnan(excluded_nan_single_trial[0, :])) == 0, np.sum(np.isnan(excluded_nan_single_trial[-1, :])) == 0)#should be all false
        # only keeping the train and val data
        if label in splitted_data:
            splitted_data[label].append(excluded_nan_single_trial)
            # print(np.any(np.isnan(excluded_nan_single_trial)))
            for name in behavior_names:
                in_trial_behavior = dataset.data.loc[in_trial_index][name].to_numpy()
                metadata[label]['behavior'][name].append(in_trial_behavior[nan_index_single_trial[0]:nan_index_single_trial[1],:])
            for name in trial_info_names:
                if name == 'start_time':
                    metadata[label]['trial_info'][name].append(nwbfile_stream.trials[trial_ind][name].item() + data_timestamp_interval*lead_end)
                elif name == 'stop_time':
                    metadata[label]['trial_info'][name].append(nwbfile_stream.trials[trial_ind][name].item() - data_timestamp_interval*(tail_start_reverse-1))
                else:
                    metadata[label]['trial_info'][name].append(nwbfile_stream.trials[trial_ind][name].item())
        
        # print('old_start_time = ', nwbfile_stream.trials[trial_ind]['start_time'].item(), 'old_end_time = ', nwbfile_stream.trials[trial_ind]['stop_time'].item())
        # print('trial_length =', np.shape(in_trial_behavior[nan_index_single_trial[0]:nan_index_single_trial[1],:]), 'start_time = ', nwbfile_stream.trials[trial_ind]['start_time'].item() + data_timestamp_interval*lead_end, 
        #       'end_time = ', nwbfile_stream.trials[trial_ind]['stop_time'].item() - data_timestamp_interval*(tail_start_reverse-1),
        #       'nan_index = ', nan_index_single_trial)
        
    train_dataset = ToyDsetDynamics(splitted_data['train'], dt=1e-3, nForward=nForward)
    val_dataset = ToyDsetDynamics(splitted_data['val'], dt=1e-3, nForward=nForward)

    train_dataloader = DataLoader(train_dataset, batch_size=batch_size, num_workers=num_workers, shuffle=True)
    val_dataloader = DataLoader(val_dataset, batch_size=batch_size, num_workers=num_workers, shuffle=False)

    io_stream.close()

    return train_dataloader, val_dataloader, splitted_data['train'], splitted_data['val'], metadata


#### Loaders for Brodmann's area 2 recordings -- center-out reach with bump task #############

def loader_area2_bump_with_behavior(spike_smooth_ms = 40, nForward = 1, num_workers = 1, batch_size = 128):
    dandiset_id = '000127' ### name of the dataset on DANDI archive (should be in the notebook example)
    filepath= "sub-Han/sub-Han_desc-train_behavior+ecephys.nwb" ### filepath on dandi archive
                                                                ### accessible by searching the dandi ID 
                                                                ### on DANDI archive https://dandiarchive.org/
    with DandiAPIClient() as client:
        asset = client.get_dandiset(dandiset_id, 'draft').get_asset_by_path(filepath)
        s3_url = asset.get_content_url(follow_redirects=1, strip_query=True)

        rem_file = remfile.File(s3_url)
        file = h5py.File(rem_file, "r")
        io_stream = NWBHDF5IO(file=file)
        nwbfile_stream = io_stream.read()
        dataset = NWBDataset(nwbfile_stream, split_heldout=True)

    dataset.smooth_spk(spike_smooth_ms, name='smth')
    trial_start_times = dataset.trial_info.start_time.dt.total_seconds().to_numpy()
    trial_end_times = dataset.trial_info.end_time.dt.total_seconds().to_numpy()
    smoothed_spikes = dataset.data.spikes_smth
    smoothed_spikes_numpy = smoothed_spikes.to_numpy()
    smoothed_spikes['time_stamp'] = smoothed_spikes.index.total_seconds()
    train_val_label = dataset.trial_info.split.to_numpy()
    splitted_data = {
        'train':[],
        'val':[],
    }

    trial_info_names = ['result', 'ctr_hold', 'ctr_hold_bump', 'bump_dir',
                        'target_on_time', 'target_dir', 'go_cue_time', 
                        'bump_time', 'move_onset_time', 'cond_dir']
    behavior_names = ['force', 'hand_pos', 'hand_vel', 'joint_ang', 'joint_vel', 'muscle_len', 'muscle_vel']
    metadata = {'train':{
                        'trial_info':{name:[] for name in trial_info_names},
                        'behavior':{name:[] for name in behavior_names}
                        },
               'val':{
                        'trial_info':{name:[] for name in trial_info_names},
                        'behavior':{name:[] for name in behavior_names}
                        }}


    for trial_ind, (trial_start, trial_end, label) in enumerate(zip(trial_start_times, trial_end_times, train_val_label)):
        in_trial_index = (smoothed_spikes.time_stamp >= trial_start) & (smoothed_spikes.time_stamp <= trial_end)
        
        # only keeping the train and val data
        if label in splitted_data:
            splitted_data[label].append(smoothed_spikes_numpy[in_trial_index, :])
            for name in behavior_names:
                metadata[label]['behavior'][name].append(dataset.data.loc[in_trial_index][name].to_numpy())
            for name in trial_info_names:
                metadata[label]['trial_info'][name].append(nwbfile_stream.trials[trial_ind][name].item())
    
    train_dataset = ToyDsetDynamics(splitted_data['train'], dt=1e-3, nForward=nForward)
    val_dataset = ToyDsetDynamics(splitted_data['val'], dt=1e-3, nForward=nForward)

    train_dataloader = DataLoader(train_dataset, batch_size=batch_size, num_workers=num_workers, shuffle=True)
    val_dataloader = DataLoader(val_dataset, batch_size=batch_size, num_workers=num_workers, shuffle=False)

    io_stream.close()

    return train_dataloader, val_dataloader, splitted_data['train'], splitted_data['val'], metadata

############## Loaders for Dorsomedial Frontal Cortex recordings -- Ready-Set-Go task ################

def make_dmfc_rsg_loaders(batch_size=128,nForward=1,smooth_len_ms=8,num_workers=1,validate=False):

    dandiset_id = '000130'
    filepath = "sub-Haydn/sub-Haydn_desc-train_ecephys.nwb"
    
    nwbfile_stream,io_stream = load_nwb_stream(dandiset_id,filepath)

    #### create nwbdataset for sorting, smoothing data
    ds = NWBDataset(fpath=nwbfile_stream,split_heldout=True)
    print(f"done!") 
    print(f"smoothing spikes with {smooth_len_ms}ms gaussian window...")
    ds.smooth_spk(smooth_len_ms,name='smth_data',ignore_nans=True)
    print("done!")
    #### get start, stop, data split labels from stream ######
    nwbfile_stream.trials[:]#['split']
    start_times_s = nwbfile_stream.trials[:]['start_time'].to_numpy()
    end_times_s = nwbfile_stream.trials[:]['stop_time'].to_numpy()
    trial_labels = nwbfile_stream.trials[:]['split'].to_list()

    #### separate into trial types #####
    rates = ds.data.spikes_smth_data
    time_s = rates.index.seconds + rates.index.microseconds/1e6
    rates_vals = rates.to_numpy()
    data = {
        'train':[],
        'val':[]
    }

    #### set up metadata dictionary ####
    trial_info_names = ['start_time','stop_time','target_on_time','ready_time','set_time','go_time','reward_time','is_eye','ts','tp']
    behavior_names=[]
    metadata = {'train':{
                        'trial_info':{name:[] for name in trial_info_names},
                        'behavior':{name:[] for name in behavior_names}
                        },
               'val':{
                        'trial_info':{name:[] for name in trial_info_names},
                        'behavior':{name:[] for name in behavior_names}
                        }}

    #### get trial data ####
    for trial_ind,(label,onset,offset) in tqdm(enumerate(zip(trial_labels,start_times_s,end_times_s)),total=len(trial_labels),desc='separating into trials'):
    
        inds = (time_s >=onset) & (time_s < offset)
        if label != 'none':
            data[label].append(rates_vals[inds,:])
            
            for name in behavior_names:
                metadata[label]['behavior'][name].append(ds.data.loc[inds][name].to_numpy())
            for name in trial_info_names:
                metadata[label]['trial_info'][name].append(nwbfile_stream.trials[trial_ind][name].item())
    #### convert to things we can load data with ####
    if validate:
        validate_metadata(data,metadata)
    train_dataset = ToyDsetDynamics(data['train'],dt=1/1000,nForward=nForward)
    val_dset = ToyDsetDynamics(data['val'],dt=1/1000,nForward=nForward)

    train_loader = DataLoader(train_dataset,batch_size=batch_size,num_workers=num_workers,shuffle=True)
    val_loader = DataLoader(train_dataset,batch_size=batch_size,num_workers=num_workers,shuffle=False)

    io_stream.close()

    return train_loader,val_loader,data['train'],data['val'],metadata


############ Loaders for Motor Cortex recordings -- Reach to touch task ##############################

def make_mc_rtt_loaders(batch_size=128,nForward=1,smooth_len_ms=8,num_workers=1,validate=False):

    dandiset_id = '000129' ### name of the dataset on DANDI archive (should be in the notebook example)
    filepath= "sub-Indy/sub-Indy_desc-train_behavior+ecephys.nwb" ### filepath on dandi archive
                                                                ### accessible by searching the dandi ID 
                                                                ### on DANDI archive https://dandiarchive.org/
    ### load nwbfile using remfile
    print('loading mc_rtt data...')
    nwbfile_stream,io_stream = load_nwb_stream(dandiset_id,filepath)
    
    #### create nwbdataset for sorting, smoothing data
    ds = NWBDataset(fpath=nwbfile_stream,split_heldout=True)
    print(f"done!") 
    print(f"smoothing spikes with {smooth_len_ms}ms gaussian window...")
    ds.smooth_spk(smooth_len_ms,name='smth_data',ignore_nans=True)
    print("done!")
    #### get start, stop, data split labels from stream ######
    nwbfile_stream.trials[:]#['split']
    start_times_s = nwbfile_stream.trials[:]['start_time'].to_numpy()
    end_times_s = nwbfile_stream.trials[:]['stop_time'].to_numpy()
    trial_labels = nwbfile_stream.trials[:]['split'].to_list()

    #### separate into trial types #####
    rates = ds.data.spikes_smth_data
    time_s = rates.index.seconds + rates.index.microseconds/1e6
    rates_vals = rates.to_numpy()
    data = {
        'train':[],
        'val':[]
    }
    behavior_names = ['cursor_pos','finger_pos','finger_vel']
    trial_info_names = []
    metadata = {'train':{
                        'trial_info':{name:[] for name in trial_info_names},
                        'behavior':{name:[] for name in behavior_names}
                        },
               'val':{
                        'trial_info':{name:[] for name in trial_info_names},
                        'behavior':{name:[] for name in behavior_names}
                        }}
    for trial_ind,(label,onset,offset) in tqdm(enumerate(zip(trial_labels,start_times_s,end_times_s)),total=len(trial_labels),desc='separating into trials'):
    
        if label != 'none':
            inds = (time_s >=onset) & (time_s < offset)
            data[label].append(rates_vals[inds,:])
            
            for name in behavior_names:
                metadata[label]['behavior'][name].append(ds.data.loc[inds][name].to_numpy())
            for name in trial_info_names:
                metadata[label]['trial_info'][name].append(nwbfile_stream.trials[trial_ind][name].item())
                #metadata[label][name].append(ds.data.loc[inds][name].to_numpy())

    if validate:
        validate_metadata(data,metadata)
    train_dataset = ToyDsetDynamics(data['train'],dt=1/1000,nForward=nForward)
    val_dset = ToyDsetDynamics(data['val'],dt=1/1000,nForward=nForward)

    train_loader = DataLoader(train_dataset,batch_size=batch_size,num_workers=num_workers,shuffle=True)
    val_loader = DataLoader(train_dataset,batch_size=batch_size,num_workers=num_workers,shuffle=False)

    io_stream.close()

    return train_loader,val_loader,data['train'],data['val'],metadata


############ Loaders for Widefield Musall data  ##############################

def loader_musall_widefield(filepath, recon_trial_num, val_split = 0.2, frame_resize_ratio = 1.0, square = True,
                            nForward = 1, num_workers = 1, batch_size = 128):
    """
    Musall widefield dataset loader.
    Use either mSM30 or mSM43
    
    - The input file is the `Vc.mat` file for each mouse (stored on Isilon).
    Example: ~/isilon/All_Staff/mice/mSM30/10-Oct-2017/Vc.mat
    - This dataset is large: loading all trials for mouse `mSM43` may take 40 minutes.
    - Use `recon_trial_num` to limit loading to the first *k* trials for faster preprocessing.
    """
    # '''
    # U:spatial, needs to be pixel by components, the raw data is components by x by y
    # Vc: temporal, needs to be components by frames
    # Reconstruct widefield: needs to be Y by X by frame
    # '''
    splitted_data = {}
    print('Read Vc.mat file...')
    Vc_path = filepath
    
    with h5py.File(Vc_path, 'r') as f:
        U = np.asarray(f['U'], dtype=np.float32)            # (C, H, W)
        Vc = f['Vc']                                        # HDF5 dataset: (trials, frames, C)  (do NOT load)
        C, H, W = U.shape
    
        # target size
        new_H = max(1, int(round(H * frame_resize_ratio)))
        new_W = max(1, int(round(W * frame_resize_ratio)))
    
        # resize U once, then reconstruct directly at small size
        if frame_resize_ratio != 1.0:
            from skimage.transform import resize
            U_small = np.empty((C, new_H, new_W), dtype=np.float32)
            for k in range(C):
                U_small[k] = resize(U[k], (new_H, new_W), order=1,
                                    preserve_range=True, anti_aliasing=True).astype(np.float32)
        else:
            U_small = U
    
        total_trials = Vc.shape[0]
        trial_num = min(recon_trial_num, total_trials)
        F = Vc.shape[1]
        print(f'reconstruct {trial_num} trials...')
    
        # preallocate already-downscaled array
        recon_widefield = np.empty((trial_num, F, new_H, new_W), dtype=np.float32)
    
        for trial in range(trial_num):
            t = np.asarray(Vc[trial, :, :], dtype=np.float32)         # (F, C) – read just this trial
            recon_widefield[trial] = np.tensordot(t, U_small, axes=(1, 0))

    # recon_widefield = np.array(recon_widefield)
    # if frame_resize_ratio != 1.0:
    #     T = recon_widefield.shape[1]
    #     H = recon_widefield.shape[2]
    #     W = recon_widefield.shape[3]
    #     new_H = max(1, int(round(H * frame_resize_ratio)))
    #     new_W = max(1, int(round(W * frame_resize_ratio)))
    #     out = np.empty((recon_widefield.shape[0], T, new_H, new_W), dtype=np.float32)
    #     for i in range(recon_widefield.shape[0]):
    #         for j in range(T):
    #             out[i, j] = resize(recon_widefield[i, j], (new_H, new_W),
    #                                order=1, preserve_range=True,
    #                                anti_aliasing=True).astype(np.float32)
    #     recon_widefield = out
        
    print('processed widefield video shape: (trial, frame, x, y)', recon_widefield.shape)
    
    if square:
        print('pad each frame to a square..')
        # shape is (trial, frame, x, y) == (N, T, H, W)
        H = recon_widefield.shape[2]
        W = recon_widefield.shape[3]
        if H > W:
            d = H - W
            pad_height = (0, 0)
            pad_width  = (d // 2, d - d // 2)
        elif W > H:
            d = W - H
            pad_height = (d // 2, d - d // 2)
            pad_width  = (0, 0)
        else:
            pad_height = (0, 0)
            pad_width  = (0, 0)
        pad_trial = (0, 0)
        pad_frame = (0, 0)
        recon_widefield = np.pad(recon_widefield,
                                 (pad_trial, pad_frame, pad_height, pad_width),
                                 mode='constant', constant_values=0)


    # replace all nan (backgground) with 0
    recon_widefield = np.nan_to_num(recon_widefield, nan=0, copy=False)

    #train test split
    num_trials = len(recon_widefield)
    np.random.seed(440)
    indices = np.random.permutation(num_trials)

    if num_trials >= 2:
        val_size = max(1, min(num_trials - 1, int(np.ceil(val_split * num_trials))))
    else:
        val_size = 0  # only one trial; put it all in train

    train_indices = indices[val_size:]
    val_indices   = indices[:val_size]

    train_data = recon_widefield[train_indices]
    val_data   = recon_widefield[val_indices]

    splitted_data['train'] = train_data
    splitted_data['val'] = val_data
        
    train_dataset = ToyDsetDynamics(splitted_data['train'], dt=1/30, nForward=nForward)#The sampling rate of widefield is 30Hz bsed on their own code
    val_dataset = ToyDsetDynamics(splitted_data['val'], dt=1/30, nForward=nForward)

    train_dataloader = DataLoader(train_dataset, batch_size=batch_size, num_workers=num_workers, shuffle=True)
    val_dataloader = DataLoader(val_dataset, batch_size=batch_size, num_workers=num_workers, shuffle=False)

    print(f'finish loading {filepath}')
    return train_dataloader, val_dataloader, splitted_data['train'], splitted_data['val']


############ Loaders for Musall Behavior data  ##############################

def loder_musall_behavior(folderpath, start_V = 0, end_V = 89928, val_split_ratio = 0.2, 
                          frame_resize_ratio = 1.0, save_recon_video = False, square = True, square_padding = True,
                          nForward = 1, num_workers = 1, batch_size = 128):
    """
    Load a behavior video from the Musall dataset.

    Notes:
        - only mSM49 dataset has video we want.
        - To use this dataset, set `filepath` to:
          '~/isilon/All_Staff/mice/mSM49/SpatialDisc/30-Jul-2018/BehaviorVideo'

    Parameters:
        filepath (str): Path to the folder containing the SVD video `.mat` files.
        start_V (int, optional): The starting frame to load. Defaults to the first frame.
        end_V (int, optional): The ending frame to load. Defaults to the last frame.
        frame_resize_ratio (float, optional): Ratio to shrink the video frames.
            Lower values reduce file size but decrease video quality. Default is 1.0 (no resizing).
        square (boolean, optional): whether modify the video as a square
        square_padding (boolean, optional): only used with square = True, 
            if True, pad the video with 0 on the shorer side so x=y,
            if False, truncate the video on the longer side so x-y

    Returns:
        np.ndarray: Video frames as a NumPy array of shape (frames, height, width).
    """

    # cPath = '/home/sp645/isilon/All_Staff/mice'
    # Animal = 'mSM49/SpatialDisc'
    # Rec = '30-Jul-2018'
    # cPath = os.path.join(cPath, Animal, Rec, 'BehaviorVideo')

    used_V = end_V - start_V

    print(f'Reconstruct {used_V} frame of video..')
    frame_x = int(240*frame_resize_ratio)
    frame_y = int(320*frame_resize_ratio)
    segment_frame_x = int(60*frame_resize_ratio)
    segment_frame_y = int(80*frame_resize_ratio)
    recon_data = np.zeros((used_V, frame_x, frame_y), dtype=np.float32)

    # Loop over 16 segments
    for i in range(1, 17):
        name = f"SVD_Cam1-Seg{i}.mat"
        mat = loadmat(os.path.join(folderpath, name))
        
        # Load U and V
        U = mat['U']   # shape (components, pixel)
        V = mat['V']   # shape (frames, components)
        VU = V[start_V:end_V, :] @ U # shape (frame, pixel)
        #print('VU shape:', VU.shape )
        seg = VU.reshape((used_V, 60, 80),  order='F')
        
        if frame_resize_ratio < 1.0: 
            seg = resize(seg, (used_V, segment_frame_x, segment_frame_y), order=1, preserve_range=True, anti_aliasing=True)
        
        row_start = (i-1) % 4 * segment_frame_x
        row_end   = row_start + segment_frame_x
        col_start = (i-1) // 4 * segment_frame_y
        col_end   = col_start + segment_frame_y
        
        recon_data[:, row_start:row_end, col_start:col_end] = seg
        print(f"Finished reconstructing segment {i}")
    
    if save_recon_video:
        with h5py.File(os.path.join(folderpath, 'reconstructed_behavior_video.h5'), 'w') as f:
            f.create_dataset('video', data=recon_data)

    # subtract mean frame pixels
    sum_image = np.sum(recon_data, axis = 0)
    frame_count = recon_data.shape[0]
    mean_image = sum_image / frame_count
    mean_subtracted_recon_data = recon_data - mean_image
    print('processed video shape: (frame, x, y)', mean_subtracted_recon_data.shape)
    
    if square:
        print('make each frame a square..')
        if square_padding:
            print('padding each frame..')
            #padding the video shape to 320,320
            pad_height = (int(40*frame_resize_ratio), int(40*frame_resize_ratio))
            pad_width = (0, 0)
            pad_frame = (0, 0)
            mean_subtracted_recon_data = np.pad(mean_subtracted_recon_data, (pad_frame, pad_height, pad_width), 
                                                mode = 'constant', constant_values=0)
            mean_image = np.pad(mean_image, (pad_height, pad_width),  mode = 'constant', constant_values=0)
        else:   
            print('truncate each frame..')
            #truncate the top of the video
            size_truncate = mean_subtracted_recon_data.shape[2] - mean_subtracted_recon_data.shape[1]
            mean_subtracted_recon_data = mean_subtracted_recon_data[:, :, size_truncate:]

    # Rotate the video pixels
    mean_subtracted_recon_data = np.transpose(mean_subtracted_recon_data, (0, 2, 1))
    mean_image = np.transpose(mean_image, (1, 0))
    
    # split data and make loader
    train_val_split_index = int((1-val_split_ratio)*mean_subtracted_recon_data.shape[0])
    train_video = mean_subtracted_recon_data[:train_val_split_index]
    val_video = mean_subtracted_recon_data[train_val_split_index:]

    train_dataset = ToyDsetDynamics(train_video, dt=1/30, nForward=nForward)#The sampling rate of video is 30Hz
    val_dataset = ToyDsetDynamics(val_video, dt=1/30, nForward=nForward)

    train_dataloader = DataLoader(train_dataset, batch_size=batch_size, num_workers=num_workers, shuffle=True)
    val_dataloader = DataLoader(val_dataset, batch_size=batch_size, num_workers=num_workers, shuffle=False)

    print(f'finish loading videos')
    return train_dataloader, val_dataloader, train_video, val_video, mean_image
```
