## Supplementary material for "Dynamic Compression Flows for Neuroscience Data": Python code: util_v7.py

```
# This work is licensed under a Creative Commons
# Attribution-NonCommercial-ShareAlike 4.0 International License.
# You should have received a copy of the license along with this
# work. If not, see http://creativecommons.org/licenses/by-nc-sa/4.0/

"""Miscellaneous utility classes and functions."""

import ctypes
import fnmatch
import importlib
import inspect
import numpy as np
import os
import shutil
import sys
import types
import io
import pickle
import re
import requests
import html
import hashlib
import glob
import tempfile
import urllib
import urllib.request
import uuid
import torch 
from torch.utils.data.dataset import Dataset
from sklearn.decomposition import PCA
from abc import ABC,abstractmethod
from scipy.ndimage import gaussian_filter
import matplotlib.pyplot as plt 

from distutils.util import strtobool
from typing import Any, List, Tuple, Union, Optional
from torchdyn.core import NeuralODE

#------------------------------------------------------------------------------------------#
# Utils for VFM
#------------------------------------------------------------------------------------------#

#------------------------------------------------------------------------------------------
#Dynamic Toy dataset utils 

class ToyData(ABC):

    """
    generic base class for toy datasets. implements
    generate, which all classes need, and defines methods needed
    by all inheriting classes
    """


    def __init__(self):

        pass

    @abstractmethod
    def f(self,x,t):
        raise NotImplementedError

    @abstractmethod
    def g(self,x,t):
        raise NotImplementedError

    @abstractmethod
    def dW(self,dt):
        raise NotImplementedError 

    @abstractmethod
    def init_conditions(self):
        raise NotImplementedError

    def dx(self,x,t,dt,sigma):

        fx = self.f(x,t)
        gx = self.g(x,t,sigma)
        dw = self.dW(dt)

        return fx * dt + gx @ dw
    
    def generate(self,n,T,dt,sigma):

        trajectories = []
        t = np.arange(dt,T+dt/2,dt)
        for ii in range(n):

            xnot = self.init_conditions()
            x = [xnot]

            for jj in range(1,len(t)):

                xx = x[jj-1]
                tt = t[jj-1]

                dx = self.dx(xx,tt,dt,sigma)
                
                x.append(xx + dx)
            
            x = np.vstack(x)
            trajectories.append(x)

        return trajectories

#vanderpol oscillator 
class Vanderpol(ToyData):

    def __init__(self,coeffs=[2,15],seed=1234):

        super(Vanderpol,self).__init__()
        self.rho,self.tau = coeffs
        self.gen = np.random.default_rng(seed=seed)

    def f(self,x,t):
        dx1 = self.rho * self.tau * (x[0] - x[0]**3/3 - x[1])
        dx2 = self.tau/self.rho * x[0]
        return np.hstack([dx1,dx2])
    def g(self,x,t,sigma):
        return sigma*np.eye(2)
    
    def dW(self,dt):
        return self.gen.multivariate_normal(mean=np.zeros((2,)),cov=dt*np.eye(2))

    def init_conditions(self):

        return self.gen.multivariate_normal(mean=[1,1],cov=np.eye(2)*0.03)

#double circles data 
class DoubleCircles(ToyData):

    def __init__(self,coeffs=[3.5,4,2*np.pi],seed=1234):

        super(DoubleCircles,self).__init__()

        self.r0,self.a,self.omega = coeffs
        self.gen = np.random.default_rng(seed=seed)

    def f(self,x,t):
        print("f unused in double circles")
        pass

    def ft(self,theta,r,t):
        #dtheta = omega
        if r > self.r0:
            return self.omega
        else:
            return -self.omega
    
    def fr(self,r,t):
        
        # potential function: (r - r0)^4 - a(r - r0)^2
        return -(4 * (r - self.r0)**3 - 2*self.a * (r - self.r0))

    def g(self,x,t,sigma):
        return sigma*np.eye(2)

    def dW(self,dt):
        return self.gen.multivariate_normal(mean=np.zeros((2,)),cov = dt*np.eye(2))
    
    def _polar_to_cartesian(self,r,theta):
    
        return np.hstack([r*np.cos(theta),r*np.sin(theta)])
    
    def _cartesian_to_polar(self,xy):
        
        r = np.linalg.norm(xy)
        theta = np.arctan2(xy[1],xy[0])
        return r,theta
    
    def init_conditions(self):
        r0 = self.r0 + self.gen.normal(loc=0,scale=0.1)
        t0 = self.gen.uniform(0,2*np.pi)

        return self._polar_to_cartesian(r0,t0)

    def dx(self,x,t,dt,sigma):
        
        r,theta = self._cartesian_to_polar(x)
        dr = self.fr(r,t)
        dtheta = self.ft(theta,r,t)
        r += dr*dt 
        theta += dtheta*dt

        newX = self._polar_to_cartesian(r,theta)
        dw_xy = self.g(x,t,sigma) @ self.dW(dt)
        xy2 = newX + dw_xy
        
        return xy2 - x
    

#Lorenz attractor - 3D ODE sys 
class Lorenz63(ToyData):

    def __init__(self,coeffs=[10,28,8/3],seed=1234):

        super(Lorenz63,self).__init__()
        self.sigma,self.rho,self.beta = coeffs
        self.gen = np.random.default_rng(seed=seed)

    def f(self,x,t):
        
        dx = self.sigma * (x[1] - x[0]) #+ sample_dW[0]
        dy = (x[0] * (self.rho - x[2]) - x[1]) #+ sample_dW[1]
        dz = (x[0]*x[1]  - self.beta*x[2]) #+ sample_dW[2]
        return np.hstack([dx,dy,dz])
    
    def g(self,x,t,sigma):
        return sigma*np.eye(3)
    
    def dW(self,dt):
        return self.gen.multivariate_normal(mean=np.zeros((3,)),cov=dt*np.eye(3))
    
    def init_conditions(self):

        return self.gen.multivariate_normal(mean=np.zeros((3,)),cov=np.eye(3))

#Alternative Lorenz attractor 
class Lorenz96(ToyData):

    def __init__(self,coeffs=[8],d=10,seed=1234):

        super(Lorenz96,self).__init__()
        self.F = coeffs
        self.d = d
        self.gen = np.random.default_rng(seed=seed)

    def f(self,x,t):
        dx = np.zeros(self.d)
        # Loops over indices (with operations and Python underflow indexing handling edge cases)
        for i in range(self.d):
            dx[i] = (x[(i + 1) % self.d] - x[i - 2]) * x[i - 1] - x[i] + self.F
        return dx
    
    def g(self,x,t,sigma):
        return np.eye(self.d)*sigma
    
    def dW(self,dt):
        return self.gen.multivariate_normal(mean=np.zeros((self.d,)),cov=np.eye(self.d)*dt)
    
    def init_conditions(self):

        return self.gen.multivariate_normal(mean=np.zeros((self.d,)),cov=np.eye(self.d))

#Rossler attractor 
class Rossler(ToyData):

    def __init__(self,coeffs=[0.1,0.1,14],seed=1234):

        super(Rossler,self).__init__()
        self.a,self.b,self.c = coeffs
        self.gen = np.random.default_rng(seed=seed)

    def f(self,x,t):
        dx = -x[1] - x[2] #+ sample_dW[0]
        dy = x[0] + self.a*x[1] #+ sample_dW[1]
        dz = self.b + x[2]*(x[0] - self.c) #+ sample_dW[2]
        return np.hstack([dx,dy,dz])
    
    def g(self,x,t,sigma):
        return np.eye(3)*sigma
    
    def dW(self,dt):
        return self.gen.multivariate_normal(mean=np.zeros((3,)),cov=np.eye(3)*dt)
    
    def init_conditions(self):

        return self.gen.multivariate_normal(mean=[0,-9,0],cov=6*np.eye(3))

#Double SDE 
class DoubleSDE(ToyData):

    def __init__(self,coeffs=[np.pi,-np.pi,np.array([-0.5,0]),np.array([0.5,0])],seed=1234):

        super(DoubleSDE,self).__init__()
        self.omega1,self.omega2,self.center1,self.center2 = coeffs
        #self.center2 = -self.center1
        
        self.gen = np.random.default_rng(seed=seed)

    def f(self,x,t):
        print("f unused in diverging sdes")
        pass
    def ft(self,theta,x,t):
        if x[0] < 0:
            return self.omega1
        else:
            return self.omega2
        
    def g(self,x,t,sigma):
        return sigma * np.eye(2)#/(np.abs(x[0])+1/10)
    
    def dW(self,dt):
        return self.gen.multivariate_normal(mean=np.zeros((2,)),cov=np.eye(2)*dt)
    
    def _polar_to_cartesian(self,r,theta):
    
        return np.hstack([r*np.cos(theta),r*np.sin(theta)])
    
    def _cartesian_to_polar(self,xy):
    
        r = np.linalg.norm(xy)
        theta = np.arctan2(xy[1],xy[0])
        return r,theta
    
    def init_conditions(self):

        return self.gen.multivariate_normal(mean=[0,-0.5],cov=np.eye(2)*0.01)

    def dx(self,x,t,dt,sigma):

        if x[0] < 0:
            v = x - self.center1
            r,theta = self._cartesian_to_polar(v)
            dtheta = self.ft(theta,x,t)
            theta += dtheta*dt
            xx2 = self._polar_to_cartesian(r,theta) + self.center1
        else:
            v = x - self.center2
            r,theta = self._cartesian_to_polar(v)
            dtheta = self.ft(theta,x,t)
            theta += dtheta*dt
            xx2 = self._polar_to_cartesian(r,theta) + self.center2

        return xx2 + self.g(x,t,sigma)@self.dW(dt) - x
    
#Alternative double SDE 
class DoubleSDEOrbit(ToyData):

    def __init__(self,coeffs=[1.65*np.pi,np.array([-8,0]),np.array([8,0])],mass=7000,seed=1234):

        super(DoubleSDEOrbit,self).__init__()
        self.omega,self.center1,self.center2 = coeffs
        self.mass=mass
        self.gen = np.random.default_rng(seed=seed)

    def f(self,x,t):
        r1,theta1 = self._cartesian_to_polar(self.center1-x)
        r2,theta2 = self._cartesian_to_polar(self.center2-x)
        F1 = self.mass/r1**3
        F2 = self.mass/r2**3
        return F1 * (self.center1- x) + F2 * (self.center2-x)

    def g(self,x,t,sigma):
        return sigma * np.eye(2)#/(np.abs(x[0])+1/10)
    
    def dW(self,dt):
        return self.gen.multivariate_normal(mean=np.zeros((2,)),cov=np.eye(2)*dt)
    
    def _polar_to_cartesian(self,r,theta):
    
        return np.hstack([r*np.cos(theta),r*np.sin(theta)])
    
    def _cartesian_to_polar(self,xy):
    
        r = np.linalg.norm(xy)
        theta = np.arctan2(xy[1],xy[0])
        return r,theta

    def init_conditions(self):

        init_xy = self.gen.multivariate_normal(mean=[0,0],cov=np.array([[0,0],[0,2]]))
        if init_xy[0] < 0:
            r1,theta1 = self._cartesian_to_polar(init_xy - self.center1)
            omega = self.omega
        else:
            r1,theta1 = self._cartesian_to_polar(init_xy - self.center2)
            omega = -self.omega
        self.vel = np.array([-omega*r1 *np.sin(theta1), omega*r1*np.cos(theta1)])

        
        return init_xy

    def dx(self,x,t,dt,sigma):

        
        dv = self.f(x,t)*dt
        
        dx = self.vel * dt
        self.vel += dv
        
        return dx + self.g(x,t,sigma)@self.dW(dt)

#Moving balls dataset (toy movie)
class Balls(ToyData):

    def __init__(self,theta=180,seed=1234):

        #dx = -x - 3y
        #dy = y - 3x
        super(Balls,self).__init__()
        self.theta= theta
        self.coeffs = lambda dt: np.array([[np.cos(theta/(2*np.pi) *dt),-np.sin(theta/(2*np.pi)*dt)],\
									  [np.sin(theta/(2*np.pi)*dt),np.cos(theta/(2*np.pi)*dt)]])

        #self.center2 = -self.center1
        
        self.gen = np.random.default_rng(seed=seed)

    def f(self,x,t,dt):
    
        return self.coeffs(dt) @ x 
    
    def g(self,x,t,sigma):
    
        return sigma*np.eye(2)
    
    def dW(self,dt):
    
        return self.gen.multivariate_normal(mean=np.zeros((2,)),cov=dt*np.eye(2))
    
    def dx(self,x,t,dt,sigma):
        
        x2 = self.f(x,t,dt)
        gx = self.g(x,t,sigma)
        dw = self.dW(dt)
        
        x2 += gx @ dw
        return x2 - x

    def init_conditions(self):

        return self.gen.multivariate_normal(mean=[0,0],cov=np.eye(2)*0.25**2)
    
    def traj_to_movie(self,trajectories,image_shape,radius,blur=False):
        """
        converts a latent trajectory to a movie
        """


        movies = []
        gX,gY = np.meshgrid(np.linspace(-1,1,2*radius),np.linspace(-1,1,2*radius))
        ball = (gX**2 + gY**2 < 1)
        image_shape = np.array(image_shape)
        
        for traj in trajectories:
            traj -= np.amin(traj)
            traj /= np.amax(np.abs(traj))

            frames = np.zeros((len(traj),1,image_shape[0],image_shape[1]))
        
            for point in range(len(traj)):
        
                ballCenter = (traj[point,:] *(image_shape - 2*radius)).astype(int) + radius
                frames[point,0,ballCenter[0]-radius:ballCenter[0] + radius, ballCenter[1]-radius:ballCenter[1] + radius] += ball
                if blur:
                    frames[point,0,:,:] = gaussian_filter(frames[point,0,:,:],sigma=radius*4,truncate=0.05)

            movies.append(frames)

        return movies


class ToyDsetDynamics(Dataset):
    """
    dataloader for toy datasets with dynamics. expects data in the form of that created by
    my toy data creation methods -- in other words, a list of np.arrays.
    flattens all arrays and creates a set of valid indices of that array to sample from.
    This set of valid indices is also based on nForward: the number of steps forward in time
    that we want our model to predict. 
    When sampling, will return samples of length nForward + 2 (last index is dt)
    
    Now also handles time-varying covariates if provided.
    """

    def __init__(self, data, dt, nForward=1, cov_dynamic_data=None, cov_static_data=None,
                image_lag_source=None, lag_k=0):
        self.maxForward = nForward
        exampleInd = np.random.choice(len(data), 1)[0]
        self.exampleTraj = data[exampleInd]
        
        lens = list(map(len, data))
        lens2 = [0] + list(np.cumsum([l for l in lens][:-1]))
        sets = [np.vstack([np.arange(ii, l+ ii - self.maxForward) for ii in range(self.maxForward + 1)]).T for l in lens]
        sumSets = [p+l for p,l in zip(sets,lens2)]
        validInds = np.vstack(sumSets)
        
        self.data = np.vstack(data)
        self.data_inds = validInds 
        self.dt = dt
        self.length = len(validInds)

        self.cov_dynamic_data = None
        if cov_dynamic_data is not None:
            cov_lens = list(map(len, cov_dynamic_data))
            assert lens == cov_lens, "Dynamic covariate trajectories must have same lengths as data trajectories"
            self.cov_dynamic_data = np.vstack(cov_dynamic_data)

        self.is_image = (self.exampleTraj.ndim == 4)  # (T,C,H,W)
        if self.is_image:
            _, self.C, self.H, self.W = self.exampleTraj.shape
        else:
            self.C = self.H = self.W = None
        

        # Handle dynamic covariates
        self.cov_static_data = None
        self.cov_static_list = None   # NEW: for images we keep list, not vstack
        if cov_static_data is not None:
            if isinstance(cov_static_data, np.ndarray) and len(cov_static_data.shape) == 2:
                expanded_static = []
                for i, l in enumerate(lens):
                    expanded_static.append(np.tile(cov_static_data[i:i+1], (l, 1)))
                self.cov_static_data = np.vstack(expanded_static)
            else:
                # list of arrays [T, dim_static] — for images we keep it as a list
                if self.is_image:
                    self.cov_static_list = [np.asarray(a) for a in cov_static_data]  # NEW
                else:
                    cov_lens = list(map(len, cov_static_data))
                    assert lens == cov_lens, "Static covariate trajectories must have same lengths as data trajectories"
                    self.cov_static_data = np.vstack(cov_static_data)

        self.image_lag_source = image_lag_source if self.is_image else None
        self.lag_k = int(lag_k) if self.is_image else 0

        # NEW: keep trial offsets to map flat indices → (trial_id, local_t)
        self.trial_offsets = np.array([0] + list(np.cumsum(lens)[:-1]))
        self.trial_lengths = np.array(lens)

    def _flat_index_to_trial_t(self, flat_idx: int):
        # Find trial j such that trial_offsets[j] <= flat_idx < trial_offsets[j] + trial_lengths[j]
        j = int(np.searchsorted(self.trial_offsets[1:], flat_idx, side='right'))
        t_local = flat_idx - int(self.trial_offsets[j])
        return j, t_local

    def __len__(self):
        return self.length 
    
    def __getitem__(self, index):
        single_index = False
        result = []
        try:
            iterator = iter(index)
        except TypeError:
            index = [index]
            single_index = True

        for ii in index:
            inds = self.data_inds[ii]
            
            # Get data samples
            samples = [self.transform(self.data[ind]) for ind in inds]
            samples.append(self.dt)

            # Add dynamic covariate samples if available
            if self.cov_dynamic_data is not None:
                cov_dynamic_samples = [self.transform(self.cov_dynamic_data[ind]) for ind in inds]
                samples.extend(cov_dynamic_samples)

            # Add static covariate samples if available
            if self.is_image:
                # Build ONE row at the left endpoint (repeat across window)
                j_trial, t_trunc = self._flat_index_to_trial_t(int(inds[0]))
                # t_original = t_trunc + lag_k  (because you truncated T→T-k before creating this dataset)
                t_original = t_trunc + self.lag_k
                # slice original frames [t-k .. t] → concat on channel
                Xorig = self.image_lag_source[j_trial]  # (T, C, H, W) original
                # safety: bounds within trial
                t0 = t_original - self.lag_k
                t1 = t_original
                lag_blocks = [Xorig[t0 + s] for s in range(self.lag_k + 1)]  # [(C,H,W), ...]
                lag_stack = np.concatenate(lag_blocks, axis=0)               # ((k+1)·C, H, W)
                lag_row = lag_stack.reshape(-1)                               # ((k+1)·C·H·W,)

                # true static (time-varying) if provided
                if self.cov_static_list is not None:
                    static_row = self.cov_static_list[j_trial][t_trunc]       # (S,)
                    fused = np.concatenate([static_row, lag_row], axis=0)
                elif self.cov_static_data is not None:
                    # unlikely for images; kept for completeness
                    fused = self.cov_static_data[inds[0]]
                else:
                    fused = lag_row

                fused_t = self.transform(fused)
                samples.extend([fused_t for _ in inds])  # repeat across the window
            else:
                # vector path (unchanged)
                if self.cov_static_data is not None:
                    cov_static_samples = [self.transform(self.cov_static_data[ind]) for ind in inds[:1]]
                    samples.extend(cov_static_samples * len(inds))

            result.append(samples)

        if single_index:
            return result[0]
        return result
    
    def transform(self, data):
        return torch.from_numpy(data).type(torch.FloatTensor)

#projection helper 
class projection():

    """
    projects things to a higher dimensional space, plus some additional nonlinearity. 
    should save out everything 

    """

    def __init__(self,origDim,newDim, projType='linear',temp=1,seed=1234) -> None:
        self.gen = np.random.default_rng(seed=seed)
        self.d1 = origDim
        self.d2 = newDim
        self.projType=projType 
        self.W = self.gen.normal(loc=0.0,scale=1.5,size=(origDim,newDim))
        self.temp=temp
        if projType == 'linear':
            

            self.projection = lambda x: x @ self.W 
        
        elif projType == 'softmax':

            assert self.temp >0, print('Temperature should be a positive number')
            self.projection = lambda x: _softmax(x @ self.W,temp=self.temp)

        elif projType == 'combine':

            self.projection = lambda x: _combine_dims(x @ self.W)

        elif projType == 'sigmoid':

            assert self.temp >0, print('Temperature should be a positive number')
            self.projection = lambda x: _sigmoid(x @ self.W,temp=self.temp)

        elif projType == 'swish':

            assert self.temp >0, print("Temperature should be a positive number")
            self.projection = lambda x: _swish(x @ self.W,temp=self.temp)
        elif projType == 'double_swish':

            assert self.temp >0, print("Temperature should be a positive number")
            self.projection = lambda x: _double_swish(x @ self.W,temp=self.temp)


        else:
            print('Method must be softmax,combine,sigmoid,swish,double swish,or linear')
            raise NotImplementedError

    
    def project(
        self,
        data: np.array,
        noise: float= 0.
     ) -> np.array:
        
        r"""
        Return batch projectiojns.
        

        Args: 
            data: what we are embedding in a higher-d space
            
        """
        if noise:
            proj = self.projection(data)
            return  proj + noise * np.random.normal(size=proj.shape)
        else:
            return self.projection(data)
          
def _swish(x:np.array,temp=1):

    xout = x * _sigmoid(x,temp)

    return xout

def _double_swish(x:np.array,temp=1):

    xout = np.sign(x) * _swish(np.abs(x),temp=temp)

    return xout

def _softmax(x:np.array,temp=1):

    m = np.amax (x,axis=1,keepdims=True)
    e_x = np.exp((x - m)/temp)
    return e_x/np.sum(e_x,axis=1,keepdims=True) 

def _combine_dims(x:np.array):

    xOut = x 
    for ii in range(x.shape[-1]-1):
        xOut[:,ii] = x[:,ii]*x[:,ii+1]

    return xOut

def _sigmoid(x:np.array,temp=1):

    xout = 1/(1 + np.exp(-x*temp))

    return xout


#wrapper to get toy dynamics dset by name 
#added option to generate LS balls case, and its corresponds IS/video samples
def get_toy_dynamicdset(dset_name, n_trajs, T, dt, sigma, project=False, proj_specs=None, balls_dset_specs=None):
    """
    Wrapper to construct desired toy dynamic dset from name, params.
    Can handle projections too if desired.
    """
    DATASETS = {'vanderpol': "Vanderpol()", 
            'doublecircles': "DoubleCircles()", 
            'rossler': "Rossler()", 
            'lorenz63': "Lorenz63()",
            'lorenz96': "Lorenz96()", 
            'doublesde': "DoubleSDE()", 
            'doublesdeorbit': "DoubleSDEOrbit()", 
               'balls':"Balls()"}
    dset_name = dset_name.lower()
    try: 
        dset_gen_obj = eval(DATASETS[dset_name])
    except KeyError:
        raise ValueError(f"Unknown Dataset: {dset_name}")
    #sample deserired trajs from it 
    dset_samples = dset_gen_obj.generate(n=n_trajs, T=T, dt=dt, sigma=sigma)
    ls_dset_samples = None 
    #project data if desired 
    if project: 
        assert proj_specs != None, 'To project need projection specs!'
        orig_dim = dset_samples[0][0].shape[0]
        projection_obj = projection(orig_dim, proj_specs.project_to, \
                                    projType=proj_specs.proj_type, \
                                    temp=proj_specs.temp)
        proj_dset_samples = [[projection_obj.project(t) for t in traj] for traj in dset_samples]
        dset_samples = proj_dset_samples
    if dset_name == 'balls':
        #pass dset to movie format (from ls trajs)
        #std imgs too 
        ls_dset_samples = dset_samples
        dset_samples = dset_gen_obj.traj_to_movie(dset_samples, balls_dset_specs.img_shape, \
                                                radius=balls_dset_specs.radius, blur=balls_dset_specs.blur)
        dset_samples = np.array(dset_samples)
        dset_samples = ((dset_samples - np.mean(dset_samples))/np.std(dset_samples))
    
    dset_obj = ToyDsetDynamics(dset_samples, dt, nForward=1)
    return dset_obj, dset_samples, ls_dset_samples

#------------------------------------------------------------------------------#
# methods to run trajectory simulation
#------------------------------------------------------------------------------#

#for flow net 

class flow_torch_wrapper(torch.nn.Module):
    """
    Wraps model to torchdyn compatible format.
    
    This is for flow net mapping between DS and LS.
    
    """
    def __init__(self, model):
        super().__init__()
        self.model = model

    def forward(self, t, x, *args, **kwargs):
        t = t.repeat(x.shape[0]) #bs 
        out = self.model(x, t)        
        return out

def calc_flow_trajectories(model, init_samples, start_tau, end_tau, nt=100):
    """
    Computes ODE trajectories for plotting/checking
    model during training.
    
    Args
    -----
    model: torch.nn.Module. Instance of IFsCFMToyNet class. 
    dset_kwargs: dict. Contains original args for data specification.
    loss_fn: instance of IFsCFMToyLoss class. 
    n:int. Number of samples to simulate.
    nt: number of time pts to use for trajectory integration.
    device: instance of torch.device.
    """
    #setup node 
    node = NeuralODE(flow_torch_wrapper(model), solver='dopri5', \
                     sensitivity="adjoint", atol=1e-4, rtol=1e-4)
    #get ts 
    ts = torch.linspace(start_tau, end_tau, nt).to(init_samples.device)
    #now sim ODE 
    with torch.no_grad():
        traj = node.trajectory(init_samples, ts)
    return traj


#for dynamics net 

class dyn_torch_wrapper(torch.nn.Module):
    """
    Wraps model to torchdyn compatible format.
    Now handles concatenated input with optional x0_tau and covariates.
    Supports covariate interpolation during integration.
    """
    def __init__(self, model, tau, x0_tau=None, cov_start=None,
                 cov_delta=None, include_x0_tau=False,
                 cov_static=None):
    
        super().__init__()
        self.model = model
        self.tau = tau
        self.include_x0_tau = include_x0_tau
        self.x0_tau = x0_tau
        self.cov_start = cov_start
        self.cov_delta = cov_delta
        self.cov_static = cov_static
        
        self.is_conv_wrapper = (
            hasattr(model, 'base_model') or  # ConvVNetWrapper has base_model attribute
            (hasattr(model, '__class__') and 'ConvVNetWrapper' in model.__class__.__name__) or
            'ConvVNetWrapper' in str(type(model))
        )
        
    def forward(self, t, x, *args, **kwargs):
        B = x.shape[0]
        # time inputs
        taus  = torch.full((B,), float(self.tau), dtype=torch.float32, device=x.device)
        t_dyn = (t if torch.is_tensor(t) else torch.tensor(t, device=x.device, dtype=torch.float32))
        t_dyn = t_dyn.float().view(-1)
        if t_dyn.numel() != B: 
            t_dyn = t_dyn[:1].repeat(B)

        # interpolate dynamic covariates at the current ODE time (if provided)
        if (self.cov_start is not None) and (self.cov_delta is not None):
            cov_current = self.cov_start + t_dyn.view(B, 1) * self.cov_delta
        elif self.cov_start is not None:
            cov_current = self.cov_start
        else:
            cov_current = None

        if self.is_conv_wrapper:
            try:
                out = self.model(x, cov_current, self.cov_static, taus, t_dyn)
            except TypeError:
                out = self.model(x, cov_current, self.cov_static, taus)

        else:
            # MLP v-net expects three parts: (v_input, taus, t_dyn)
            parts = [x]
            # if self.include_x0_tau and (self.x0_tau is not None):
            #     parts.append(self.x0_tau)
            if cov_current is not None:
                parts.append(cov_current)
            if self.cov_static is not None:
                parts.append(self.cov_static)
            x_concat = torch.cat(parts, dim=-1) if len(parts) > 1 else x
            try:
                out = self.model(x_concat, taus, t_dyn)
            except TypeError:
                out = self.model(x_concat, taus)
        return out

def calc_dyn_trajectories(model, init_samples, tau,
                          x0_tau=None, covariates=None,
                          next_covariates=None,
                          include_x0_tau=False, nt=100,
                          covariates_static=None):
    """
    Computes dynamics net trajectories for a given set of initial
    samples and tau, now with support for interpolated covariates.
    
    """
    # If x0_tau not provided but needed, assume dynamics starts where flow started
    if x0_tau is None and include_x0_tau:
        x0_tau = init_samples.clone()
    
    # Compute covariate delta if covariates and next_covariates provided
    cov_delta = None
    if covariates is not None and next_covariates is not None:
        cov_delta = next_covariates - covariates
    
    # Setup node with covariate interpolation
    node = NeuralODE(
        dyn_torch_wrapper(
            model, tau, x0_tau,
            cov_start=covariates, cov_delta=cov_delta,
            include_x0_tau=include_x0_tau,
            cov_static=covariates_static
        ), 
        solver='dopri5', 
        sensitivity="adjoint", 
        atol=1e-4, 
        rtol=1e-4
    )
    
    # Get ts 
    ts = torch.linspace(0.0, 1.0, nt).to(init_samples.device)
    # Now sim ODE 
    with torch.no_grad():
        traj = node.trajectory(init_samples, ts)
    return traj


#method to simulate traj encoding 

def sim_encoded_trajs(gt_trajs, encoder_net, device):
    """
    Chooses one GT traj from training set at random
    and encodes it. 
    """
    traj_to_sim_idx = np.random.choice(np.arange(gt_trajs.shape[0]), size=1, replace=False)
    traj_to_sim = torch.from_numpy(gt_trajs[traj_to_sim_idx, :, :]).type(torch.float32).to(device).squeeze(0)
    with torch.no_grad():
        encoded_traj = encoder_net.rsample(traj_to_sim)
    return traj_to_sim.cpu().numpy(), encoded_traj.detach().cpu().numpy()


def plot_balls_traj(traj_to_plot):
    """
    Constructs simple 5x10 grid with 50 first
    steps of a given trajectory.
    """
    fig,axs = plt.subplots(nrows=5,ncols=10, figsize=(10,10))
    for i in range(5):
        for j in range(10):
            frame_idx = i*10 + j 
            frame_to_plot = traj_to_plot[frame_idx, :, :, :]
            axs[i, j].imshow(np.squeeze(frame_to_plot), cmap='gray')
    return fig 


def plot_encoded_trajs(gt_traj, encoded_traj, imgshape):
    """
    Construct GT vs. encoded trajectories figure
    which will be logged into TB.
    """
    gt_fig = plot_balls_traj(gt_traj.reshape(-1, imgshape, imgshape, 1))
    enc_fig = plot_balls_traj(encoded_traj.reshape(-1, imgshape, imgshape, 1))
    return gt_fig, enc_fig 

#------------------------------------------------------------------------------#

# Methods to sim dynamics trajectories using simple SDEs

def get_dyn_SDE_dx(dyn_net, curr_x, flow_time, x0_tau=None, covariates_dynamic=None,
                   next_covariates_dynamic=None, covariates_static=None,
                   include_x0_tau=True, sigma=0.1, dt=0.001, local_t=None):
    """
    One Euler–Maruyama step.
    - Dynamic covs: interpolate with local_t∈[0,1] if both ends given; otherwise hold constant.
    - Static covs: constant within each Δt interval.
    flow_time = τ (conditioning for the v-net), kept constant.
    """
    cov_delta = None
    if (covariates_dynamic is not None) and (next_covariates_dynamic is not None):
        cov_delta = next_covariates_dynamic - covariates_dynamic

    dyn_net_wrapper = dyn_torch_wrapper(
        dyn_net, flow_time,
        x0_tau=x0_tau,
        cov_start=covariates_dynamic, cov_delta=cov_delta,
        include_x0_tau=include_x0_tau,
        cov_static=covariates_static,
    )

    t_eval = 0.0 if local_t is None else float(local_t)
    f = dyn_net_wrapper(t_eval, curr_x)

    g = sigma * torch.eye(curr_x.shape[1], device=curr_x.device, dtype=torch.float32)
    dW = torch.randn_like(curr_x, dtype=torch.float32) * np.sqrt(dt)
    g_dW = torch.einsum('ij, bjk -> bik', g, dW.unsqueeze(-1)).squeeze(-1)
    dx = f * dt + g_dW
    return dx


def int_dyn_SDE(init_x, dyn_net, flow_time, x0_tau=None,
                covariates_dynamic=None, next_covariates_dynamic=None,
                covariates_static=None,
                include_x0_tau=True, sigma=0.1, start_time=0.0, end_time=1.0, dt=0.001):
    """
    Euler–Maruyama simulation on t ∈ [start_time, end_time] with step dt.
    Dynamic covs are interpolated w.r.t. normalized local time; static covs are held constant.
    """
    device = init_x.device
    times = torch.arange(start_time, end_time, step=dt, device=device)
    int_trajs = []
    curr_x = init_x
    int_trajs.append(init_x.cpu().numpy())
    T = max(float(end_time - start_time), 1e-12)
    for i in range(times.shape[0]):
        t_norm = (float(times[i]) - float(start_time)) / T
        dx = get_dyn_SDE_dx(
            dyn_net, curr_x, flow_time,
            x0_tau=x0_tau,
            covariates_dynamic=covariates_dynamic,
            next_covariates_dynamic=next_covariates_dynamic,
            covariates_static=covariates_static,
            include_x0_tau=include_x0_tau, sigma=sigma, dt=dt, local_t=t_norm,
        )
        curr_x = curr_x + dx
        int_trajs.append(curr_x.cpu().numpy())
    return int_trajs
    
#------------------------------------------------------------------------------#

# Util classes (EDM repo)

#-------------------------------------------------------------------------------#


class EasyDict(dict):
    """Convenience class that behaves like a dict but allows access with the attribute syntax."""

    def __getattr__(self, name: str) -> Any:
        try:
            return self[name]
        except KeyError:
            raise AttributeError(name)

    def __setattr__(self, name: str, value: Any) -> None:
        self[name] = value

    def __delattr__(self, name: str) -> None:
        del self[name]


class Logger(object):
    """Redirect stderr to stdout, optionally print stdout to a file, and optionally force flushing on both stdout and the file."""

    def __init__(self, file_name: Optional[str] = None, file_mode: str = "w", should_flush: bool = True):
        self.file = None

        if file_name is not None:
            self.file = open(file_name, file_mode)

        self.should_flush = should_flush
        self.stdout = sys.stdout
        self.stderr = sys.stderr

        sys.stdout = self
        sys.stderr = self

    def __enter__(self) -> "Logger":
        return self

    def __exit__(self, exc_type: Any, exc_value: Any, traceback: Any) -> None:
        self.close()

    def write(self, text: Union[str, bytes]) -> None:
        """Write text to stdout (and a file) and optionally flush."""
        if isinstance(text, bytes):
            text = text.decode()
        if len(text) == 0: # workaround for a bug in VSCode debugger: sys.stdout.write(''); sys.stdout.flush() => crash
            return

        if self.file is not None:
            self.file.write(text)

        self.stdout.write(text)

        if self.should_flush:
            self.flush()

    def flush(self) -> None:
        """Flush written text to both stdout and a file, if open."""
        if self.file is not None:
            self.file.flush()

        self.stdout.flush()

    def close(self) -> None:
        """Flush, close possible files, and remove stdout/stderr mirroring."""
        self.flush()

        # if using multiple loggers, prevent closing in wrong order
        if sys.stdout is self:
            sys.stdout = self.stdout
        if sys.stderr is self:
            sys.stderr = self.stderr

        if self.file is not None:
            self.file.close()
            self.file = None


# Cache directories
# ------------------------------------------------------------------------------------------#

_dnnlib_cache_dir = None

def set_cache_dir(path: str) -> None:
    global _dnnlib_cache_dir
    _dnnlib_cache_dir = path

def make_cache_dir_path(*paths: str) -> str:
    if _dnnlib_cache_dir is not None:
        return os.path.join(_dnnlib_cache_dir, *paths)
    if 'DNNLIB_CACHE_DIR' in os.environ:
        return os.path.join(os.environ['DNNLIB_CACHE_DIR'], *paths)
    if 'HOME' in os.environ:
        return os.path.join(os.environ['HOME'], '.cache', 'dnnlib', *paths)
    if 'USERPROFILE' in os.environ:
        return os.path.join(os.environ['USERPROFILE'], '.cache', 'dnnlib', *paths)
    return os.path.join(tempfile.gettempdir(), '.cache', 'dnnlib', *paths)

# Small util functions
# ------------------------------------------------------------------------------------------#


def format_time(seconds: Union[int, float]) -> str:
    """Convert the seconds to human readable string with days, hours, minutes and seconds."""
    s = int(np.rint(seconds))

    if s < 60:
        return "{0}s".format(s)
    elif s < 60 * 60:
        return "{0}m {1:02}s".format(s // 60, s % 60)
    elif s < 24 * 60 * 60:
        return "{0}h {1:02}m {2:02}s".format(s // (60 * 60), (s // 60) % 60, s % 60)
    else:
        return "{0}d {1:02}h {2:02}m".format(s // (24 * 60 * 60), (s // (60 * 60)) % 24, (s // 60) % 60)


def format_time_brief(seconds: Union[int, float]) -> str:
    """Convert the seconds to human readable string with days, hours, minutes and seconds."""
    s = int(np.rint(seconds))

    if s < 60:
        return "{0}s".format(s)
    elif s < 60 * 60:
        return "{0}m {1:02}s".format(s // 60, s % 60)
    elif s < 24 * 60 * 60:
        return "{0}h {1:02}m".format(s // (60 * 60), (s // 60) % 60)
    else:
        return "{0}d {1:02}h".format(s // (24 * 60 * 60), (s // (60 * 60)) % 24)


def ask_yes_no(question: str) -> bool:
    """Ask the user the question until the user inputs a valid answer."""
    while True:
        try:
            print("{0} [y/n]".format(question))
            return strtobool(input().lower())
        except ValueError:
            pass


def tuple_product(t: Tuple) -> Any:
    """Calculate the product of the tuple elements."""
    result = 1

    for v in t:
        result *= v

    return result


_str_to_ctype = {
    "uint8": ctypes.c_ubyte,
    "uint16": ctypes.c_uint16,
    "uint32": ctypes.c_uint32,
    "uint64": ctypes.c_uint64,
    "int8": ctypes.c_byte,
    "int16": ctypes.c_int16,
    "int32": ctypes.c_int32,
    "int64": ctypes.c_int64,
    "float32": ctypes.c_float,
    "float64": ctypes.c_double
}


def get_dtype_and_ctype(type_obj: Any) -> Tuple[np.dtype, Any]:
    """Given a type name string (or an object having a __name__ attribute), return matching Numpy and ctypes types that have the same size in bytes."""
    type_str = None

    if isinstance(type_obj, str):
        type_str = type_obj
    elif hasattr(type_obj, "__name__"):
        type_str = type_obj.__name__
    elif hasattr(type_obj, "name"):
        type_str = type_obj.name
    else:
        raise RuntimeError("Cannot infer type name from input")

    assert type_str in _str_to_ctype.keys()

    my_dtype = np.dtype(type_str)
    my_ctype = _str_to_ctype[type_str]

    assert my_dtype.itemsize == ctypes.sizeof(my_ctype)

    return my_dtype, my_ctype


def is_pickleable(obj: Any) -> bool:
    try:
        with io.BytesIO() as stream:
            pickle.dump(obj, stream)
        return True
    except:
        return False


# Functionality to import modules/objects by name, and call functions by name
# ------------------------------------------------------------------------------------------#

def get_module_from_obj_name(obj_name: str) -> Tuple[types.ModuleType, str]:
    """Searches for the underlying module behind the name to some python object.
    Returns the module and the object name (original name with module part removed)."""

    # allow convenience shorthands, substitute them by full names
    obj_name = re.sub("^np.", "numpy.", obj_name)
    obj_name = re.sub("^tf.", "tensorflow.", obj_name)

    # list alternatives for (module_name, local_obj_name)
    parts = obj_name.split(".")
    name_pairs = [(".".join(parts[:i]), ".".join(parts[i:])) for i in range(len(parts), 0, -1)]

    # try each alternative in turn
    for module_name, local_obj_name in name_pairs:
        try:
            module = importlib.import_module(module_name) # may raise ImportError
            get_obj_from_module(module, local_obj_name) # may raise AttributeError
            return module, local_obj_name
        except:
            pass

    # maybe some of the modules themselves contain errors?
    for module_name, _local_obj_name in name_pairs:
        try:
            importlib.import_module(module_name) # may raise ImportError
        except ImportError:
            if not str(sys.exc_info()[1]).startswith("No module named '" + module_name + "'"):
                raise

    # maybe the requested attribute is missing?
    for module_name, local_obj_name in name_pairs:
        try:
            module = importlib.import_module(module_name) # may raise ImportError
            get_obj_from_module(module, local_obj_name) # may raise AttributeError
        except ImportError:
            pass

    # we are out of luck, but we have no idea why
    raise ImportError(obj_name)


def get_obj_from_module(module: types.ModuleType, obj_name: str) -> Any:
    """Traverses the object name and returns the last (rightmost) python object."""
    if obj_name == '':
        return module
    obj = module
    for part in obj_name.split("."):
        obj = getattr(obj, part)
    return obj


def get_obj_by_name(name: str) -> Any:
    """Finds the python object with the given name."""
    module, obj_name = get_module_from_obj_name(name)
    return get_obj_from_module(module, obj_name)


def call_func_by_name(*args, func_name: str = None, **kwargs) -> Any:
    """Finds the python object with the given name and calls it as a function."""
    assert func_name is not None
    func_obj = get_obj_by_name(func_name)
    assert callable(func_obj)
    return func_obj(*args, **kwargs)


def construct_class_by_name(*args, class_name: str = None, **kwargs) -> Any:
    """Finds the python class with the given name and constructs it with the given arguments."""
    return call_func_by_name(*args, func_name=class_name, **kwargs)


def get_module_dir_by_obj_name(obj_name: str) -> str:
    """Get the directory path of the module containing the given object name."""
    module, _ = get_module_from_obj_name(obj_name)
    return os.path.dirname(inspect.getfile(module))


def is_top_level_function(obj: Any) -> bool:
    """Determine whether the given object is a top-level function, i.e., defined at module scope using 'def'."""
    return callable(obj) and obj.__name__ in sys.modules[obj.__module__].__dict__


def get_top_level_function_name(obj: Any) -> str:
    """Return the fully-qualified name of a top-level function."""
    assert is_top_level_function(obj)
    module = obj.__module__
    if module == '__main__':
        module = os.path.splitext(os.path.basename(sys.modules[module].__file__))[0]
    return module + "." + obj.__name__


# File system helpers
# ------------------------------------------------------------------------------------------#

def list_dir_recursively_with_ignore(dir_path: str, ignores: List[str] = None, add_base_to_relative: bool = False) -> List[Tuple[str, str]]:
    """List all files recursively in a given directory while ignoring given file and directory names.
    Returns list of tuples containing both absolute and relative paths."""
    assert os.path.isdir(dir_path)
    base_name = os.path.basename(os.path.normpath(dir_path))

    if ignores is None:
        ignores = []

    result = []

    for root, dirs, files in os.walk(dir_path, topdown=True):
        for ignore_ in ignores:
            dirs_to_remove = [d for d in dirs if fnmatch.fnmatch(d, ignore_)]

            # dirs need to be edited in-place
            for d in dirs_to_remove:
                dirs.remove(d)

            files = [f for f in files if not fnmatch.fnmatch(f, ignore_)]

        absolute_paths = [os.path.join(root, f) for f in files]
        relative_paths = [os.path.relpath(p, dir_path) for p in absolute_paths]

        if add_base_to_relative:
            relative_paths = [os.path.join(base_name, p) for p in relative_paths]

        assert len(absolute_paths) == len(relative_paths)
        result += zip(absolute_paths, relative_paths)

    return result


def copy_files_and_create_dirs(files: List[Tuple[str, str]]) -> None:
    """Takes in a list of tuples of (src, dst) paths and copies files.
    Will create all necessary directories."""
    for file in files:
        target_dir_name = os.path.dirname(file[1])

        # will create all intermediate-level directories
        if not os.path.exists(target_dir_name):
            os.makedirs(target_dir_name)

        shutil.copyfile(file[0], file[1])


# URL helpers
# ------------------------------------------------------------------------------------------#

def is_url(obj: Any, allow_file_urls: bool = False) -> bool:
    """Determine whether the given object is a valid URL string."""
    if not isinstance(obj, str) or not "://" in obj:
        return False
    if allow_file_urls and obj.startswith('file://'):
        return True
    try:
        res = requests.compat.urlparse(obj)
        if not res.scheme or not res.netloc or not "." in res.netloc:
            return False
        res = requests.compat.urlparse(requests.compat.urljoin(obj, "/"))
        if not res.scheme or not res.netloc or not "." in res.netloc:
            return False
    except:
        return False
    return True


def open_url(url: str, cache_dir: str = None, num_attempts: int = 10, verbose: bool = True, return_filename: bool = False, cache: bool = True) -> Any:
    """Download the given URL and return a binary-mode file object to access the data."""
    assert num_attempts >= 1
    assert not (return_filename and (not cache))

    # Doesn't look like an URL scheme so interpret it as a local filename.
    if not re.match('^[a-z]+://', url):
        return url if return_filename else open(url, "rb")

    # Handle file URLs.  This code handles unusual file:// patterns that
    # arise on Windows:
    #
    # file:///c:/foo.txt
    #
    # which would translate to a local '/c:/foo.txt' filename that's
    # invalid.  Drop the forward slash for such pathnames.
    #
    # If you touch this code path, you should test it on both Linux and
    # Windows.
    #
    # Some internet resources suggest using urllib.request.url2pathname() but
    # but that converts forward slashes to backslashes and this causes
    # its own set of problems.
    if url.startswith('file://'):
        filename = urllib.parse.urlparse(url).path
        if re.match(r'^/[a-zA-Z]:', filename):
            filename = filename[1:]
        return filename if return_filename else open(filename, "rb")

    assert is_url(url)

    # Lookup from cache.
    if cache_dir is None:
        cache_dir = make_cache_dir_path('downloads')

    url_md5 = hashlib.md5(url.encode("utf-8")).hexdigest()
    if cache:
        cache_files = glob.glob(os.path.join(cache_dir, url_md5 + "_*"))
        if len(cache_files) == 1:
            filename = cache_files[0]
            return filename if return_filename else open(filename, "rb")

    # Download.
    url_name = None
    url_data = None
    with requests.Session() as session:
        if verbose:
            print("Downloading %s ..." % url, end="", flush=True)
        for attempts_left in reversed(range(num_attempts)):
            try:
                with session.get(url) as res:
                    res.raise_for_status()
                    if len(res.content) == 0:
                        raise IOError("No data received")

                    if len(res.content) < 8192:
                        content_str = res.content.decode("utf-8")
                        if "download_warning" in res.headers.get("Set-Cookie", ""):
                            links = [html.unescape(link) for link in content_str.split('"') if "export=download" in link]
                            if len(links) == 1:
                                url = requests.compat.urljoin(url, links[0])
                                raise IOError("Google Drive virus checker nag")
                        if "Google Drive - Quota exceeded" in content_str:
                            raise IOError("Google Drive download quota exceeded -- please try again later")

                    match = re.search(r'filename="([^"]*)"', res.headers.get("Content-Disposition", ""))
                    url_name = match[1] if match else url
                    url_data = res.content
                    if verbose:
                        print(" done")
                    break
            except KeyboardInterrupt:
                raise
            except:
                if not attempts_left:
                    if verbose:
                        print(" failed")
                    raise
                if verbose:
                    print(".", end="", flush=True)

    # Save to cache.
    if cache:
        safe_name = re.sub(r"[^0-9a-zA-Z-._]", "_", url_name)
        safe_name = safe_name[:min(len(safe_name), 128)]
        cache_file = os.path.join(cache_dir, url_md5 + "_" + safe_name)
        temp_file = os.path.join(cache_dir, "tmp_" + uuid.uuid4().hex + "_" + url_md5 + "_" + safe_name)
        os.makedirs(cache_dir, exist_ok=True)
        with open(temp_file, "wb") as f:
            f.write(url_data)
        os.replace(temp_file, cache_file) # atomic
        if return_filename:
            return cache_file

    # Return data as file object.
    assert not return_filename
    return io.BytesIO(url_data)

# util_v4.py
import numpy as np

def split_cov_static_new(cov_static_new_list, img_ch, img_size, lag_k):
    """
    cov_static_new_list: list of arrays [T_i, Ss + (k+1)*C*H*W]
    returns:
      cov_static_list: list of arrays [T_i, Ss]  (may be Ss=0)
      lag_flat_list:   list of arrays [T_i, (k+1)*C*H*W]
    """
    C, H, W = img_ch, img_size, img_size
    lag_dim = (lag_k + 1) * C * H * W

    cov_static_list, lag_flat_list = [], []
    for arr in cov_static_new_list:
        arr = np.asarray(arr)
        T, total = arr.shape
        Ss = total - lag_dim
        if Ss < 0:
            raise ValueError(f"split_cov_static_new: got total={total} < lag_dim={lag_dim}")
        cov_static_list.append(arr[:, :Ss] if Ss > 0 else np.zeros((T, 0), dtype=arr.dtype))
        lag_flat_list.append(arr[:, Ss:])  # always (T, lag_dim)
    return cov_static_list, lag_flat_list

# --------------------------------------------------
# TB plot for general image...

def _get_enc_mod(model):
    base = model.module if hasattr(model, 'module') else model
    return getattr(base, 'encoder', base)

def _is_image_like(arr):
    if arr.ndim == 3 and arr.shape[0] in (1, 3):
        return True  # (C,H,W)
    if arr.ndim == 4 and arr.shape[1] in (1, 3):
        return True  # (T,C,H,W)
    return False


def _make_recon_figure(sample_np, recon_np):
    if plt is None:
        return None

    sample_np = np.asarray(sample_np)
    recon_np = np.asarray(recon_np)

    # Image-like case.
    if _is_image_like(sample_np):
        # Normalize to (T,C,H,W)
        if sample_np.ndim == 3 and sample_np.shape[0] in (1, 3):  # (C,H,W)
            sample_np = sample_np[None, ...]
            recon_np = recon_np[None, ...]
        elif sample_np.ndim == 3 and sample_np.shape[-1] in (1, 3):  # (H,W,C)
            sample_np = np.moveaxis(sample_np, -1, 0)[None, ...]
            recon_np = np.moveaxis(recon_np, -1, 0)[None, ...]
        elif sample_np.ndim == 4 and sample_np.shape[-1] in (1, 3):  # (T,H,W,C)
            sample_np = np.moveaxis(sample_np, -1, 1)
            recon_np = np.moveaxis(recon_np, -1, 1)

        # Now sample_np: (T,C,H,W)
        T = min(8, sample_np.shape[0])
        fig, axes = plt.subplots(2, T, figsize=(2 * T, 4))
        if T == 1:
            axes = np.array(axes).reshape(2, 1)

        for t in range(T):
            gt = sample_np[t]
            rc = recon_np[t]
            gt = np.moveaxis(gt, 0, -1)  # (H,W,C)
            rc = np.moveaxis(rc, 0, -1)

            axes[0, t].imshow(np.clip(gt, 0.0, 1.0), cmap=None if gt.shape[-1] > 1 else "gray")
            axes[0, t].axis("off")

            axes[1, t].imshow(np.clip(rc, 0.0, 1.0), cmap=None if rc.shape[-1] > 1 else "gray")
            axes[1, t].axis("off")

        axes[0, 0].set_ylabel("GT", fontsize=8)
        axes[1, 0].set_ylabel("REC", fontsize=8)
        fig.tight_layout()
        return fig

    # Vector-like case: treat as (T,D) or (D,)
    if sample_np.ndim == 1:
        sample_np = sample_np[:, None]
        recon_np = recon_np[:, None]
    elif sample_np.ndim > 2:
        # Flatten trailing dims to D
        T = sample_np.shape[0]
        sample_np = sample_np.reshape(T, -1)
        recon_np = recon_np.reshape(T, -1)

    T = sample_np.shape[0]
    D = sample_np.shape[1]
    n_plot = min(2, D)

    fig, axes = plt.subplots(n_plot, 1, figsize=(6, 3 * n_plot))
    if n_plot == 1:
        axes = [axes]

    t = np.arange(T)
    for d in range(n_plot):
        ax = axes[d]
        ax.plot(t, sample_np[:, d], label="gt")
        ax.plot(t, recon_np[:, d], label="rec", linestyle="--")
        ax.set_xlabel("t")
        ax.set_ylabel(f"dim {d}")
        ax.legend(fontsize=8)
    fig.tight_layout()
    return fig


def log_reconstruction_viz(writer, model, dset_samples, device, gs):
    if writer is None:
        return
    if dset_samples is None or len(dset_samples) == 0:
        return

    try:
        sample = dset_samples[0]
        sample_np = np.asarray(sample)

        # Build input batch for encoder: treat time/frames as batch dimension if present.
        if sample_np.ndim == 1:
            x_in = sample_np[None, :]  # (1,D)
        else:
            # Use all time steps / frames as separate batch items.
            T = sample_np.shape[0]
            x_in = sample_np.reshape(T, -1)

        enc = _get_enc_mod(model)
        with torch.no_grad():
            x_t = torch.from_numpy(x_in).to(device=device, dtype=torch.float32)
            out = enc.rsample(x_t)
            if isinstance(out, (tuple, list)):
                recon_flat = out[0]
            else:
                recon_flat = out

        recon_flat = recon_flat.detach().cpu().numpy()
        recon_np = recon_flat.reshape(sample_np.shape)

        fig = _make_recon_figure(sample_np, recon_np)
        if fig is not None:
            writer.add_figure("recon/example", fig, gs)
            writer.flush()
            plt.close(fig)
    except Exception as e:  # pragma: no cover
        # Never break training because of visualization.
        dist.print0(f"[viz] Skipped reconstruction viz due to error: {repr(e)}")
```
