## Supplementary material for "Dynamic Compression Flows for Neuroscience Data": Python code: vae.py

```
import torch
import os
from torch import nn
import numpy as np
import matplotlib.pyplot as plt
from torch.utils.tensorboard import SummaryWriter
from torch.distributions import LowRankMultivariateNormal
### importing toy datasets
#sys.path.insert(0,'/hdd/miles/velocity_cfm/dnnlib')

class AutoEncoder(nn.Module):

    def __init__(self,encoder,decoder,latent_model,save_dir='./autoencoder',loss_fn = nn.MSELoss(reduction='mean'),beta=1):

        super(AutoEncoder,self).__init__()
        self.encoder = encoder
        self.decoder = decoder
        self.latent_model=latent_model
        self.loss_fn = loss_fn
        self.save_dir=save_dir
        self.epoch=0
        self.beta=beta
        self.device = 'cuda' if torch.cuda.is_available() else 'cpu'

        self.to(self.device)
        self.writer = SummaryWriter(log_dir=os.path.join(save_dir,'runs'))
        

    def encode(self,x):

        #z = torch.vmap(self.encoder,in_dims=1,out_dims=1)(x)
        #print(x.shape)
        z = self.encoder(x)
        
        z,latent_reg = self.latent_model(z)

        return z,latent_reg

    def decode(self,z):

        return self.decoder(z)

    def forward(self,x):

        #print(x.shape)
        z,latent_reg = self.encode(x)

        xhat = self.decode(z)

        return xhat,latent_reg

    def train_epoch(self,loader,optimizer):

        tl = []
        for ii,batch in enumerate(loader,start=len(loader)*self.epoch):

            optimizer.zero_grad()
            
            data,dt= batch[:-1],batch[-1]
            data = torch.hstack(data).to(self.device)[:,None,:]
            #print(data.shape)
            xhat,latent_reg = self.forward(data)
            #print(xhat.shape)
            #print(data.shape)
            if xhat.shape[1] < data.shape[1]:
                recon_loss = self.loss_fn(xhat,data[:,1:,...])

            else:
                recon_loss = self.loss_fn(xhat,data)
                #print(recon_loss)
        
                
            loss = (self.beta*latent_reg + recon_loss)/len(data)
            #print(loss)
            #assert False
            loss.backward()
            optimizer.step()

            self.writer.add_scalar('Train/recon loss',recon_loss.item(),ii)
            self.writer.add_scalar('Train/latent regularization',latent_reg.item(),ii)
            tl.append(loss.item())
        self.epoch += 1

        return tl,optimizer

    def val_epoch(self,loader):

        vl = []
        reg = []
        recon = []
        with torch.no_grad():

            for batch in loader:
                data,dt= batch[:-1],batch[-1]
                
                #data = torch.stack(data,axis=1).to(self.device)
                data = torch.hstack(data).to(self.device)[:,None,:]

                xhat,latent_reg = self.forward(data)
                if xhat.shape[1] < data.shape[1]:
                    recon_loss = self.loss_fn(xhat,data[:,1:,...])
    
                else:
                    #print("lossin' it up")
                    recon_loss = self.loss_fn(xhat,data)
                loss = (self.beta*latent_reg + recon_loss)/len(data)
                vl.append(loss.item())
                reg.append(latent_reg.item())
                recon.append(recon_loss.item())
            self.writer.add_scalar('Val/recon loss',np.nanmean(recon),self.epoch)
            self.writer.add_scalar('Val/latent regularization',np.nanmean(reg),self.epoch)

        return vl

    def save(self,optimizer):

        state_dict_full = {'model_dict':self.state_dict(),'opt_dict':optimizer.state_dict(),'epoch':self.epoch}
        torch.save(state_dict_full,os.path.join(self.save_dir,f'checkpoint_{self.epoch}.tar'))

    def load(self,path,optimizer):

        state_dict = torch.load(path)
        optimizer.load_state_dict(state_dict['opt_dict'])
        self.load_state_dict(state_dict['model_dict'])
        self.epoch = state_dict['epoch']
        return optimizer

class LatentLSTM(nn.Module):
    
    def __init__(self,input_size,hidden_size,num_layers,loss_fn=nn.MSELoss(reduction='mean')):

        super().__init__()

        self.net = nn.LSTM(input_size=input_size,hidden_size=hidden_size,num_layers=num_layers,batch_first=True)
        self.output = nn.Linear(hidden_size,input_size)
        self.loss_fn = loss_fn

    def forward(self,x):

        x,y = x[:,:-1,...],x[:,1:,...]
        out,_ = self.net(x)
        yhat = self.output(out)
        return yhat,self.loss_fn(yhat,y)

class LatentVAE(nn.Module):
    
    def __init__(self,input_size,output_size,num_hidden,hidden_size=10):

        super().__init__()

        mu = [nn.Linear(input_size,hidden_size,bias=True), nn.ReLU()]
        u = [nn.Linear(input_size,hidden_size,bias=True),nn.ReLU()]
        d = [nn.Linear(input_size,hidden_size,bias=True),nn.ReLU()]
        for _ in range(num_hidden):
            mu.append(nn.Linear(hidden_size,hidden_size))
            mu.append(nn.ReLU())
            u.append(nn.Linear(hidden_size,hidden_size))
            u.append(nn.ReLU())
            d.append(nn.Linear(hidden_size,hidden_size))
            d.append(nn.ReLU())
        mu.append(nn.Linear(hidden_size,output_size))
        u.append(nn.Linear(hidden_size,output_size))
        d.append(nn.Linear(hidden_size,output_size))

        self.mu = nn.Sequential(*mu)
        self.u = nn.Sequential(*u)
        self.d = nn.Sequential(*d)

    def forward(self,x):

        mu,u,d = self.mu(x),self.u(x),self.d(x)
        u = u.unsqueeze(-1)
        d = torch.exp(d)
        
        latent_dist = LowRankMultivariateNormal(mu, u, d)
        z = latent_dist.rsample()
        #print(z.shape)
        #print(latent_dist.entropy().shape)
        
        kl_term = -0.5 * (torch.sum(torch.pow(z,2),axis=-1) + z.shape[-1] * np.log(2*np.pi))
        #assert False
        kl_term = kl_term.sum() + torch.sum(latent_dist.entropy())
        
        
        return z,-kl_term

class LatentAC(nn.Module):

        def __init__(self):
            super().__init__()

        def forward(self,x):
            return x,torch.tensor([0]).type(torch.FloatTensor).to(x.device)
```
