## Supplementary material for "Dynamic Compression Flows for Neuroscience Data": Python code: loss_v7.py

```
# This work is licensed under a Creative Commons
# Attribution-NonCommercial-ShareAlike 4.0 International License.
# You should have received a copy of the license along with this
# work. If not, see http://creativecommons.org/licenses/by-nc-sa/4.0/

"""
Loss functions used. 

Containts both original loss fns (from EDM paper)
and inflationary flow loss fns (for toy and image data).
"""

import torch
from torch_utils import persistence
from torch_cfm import conditional_flow_matching as cfm
import torch, torch.nn.functional as F
# import numpy as np 

# # this is remained for IS....
# LAMBDA_MIX = 0.8
# def _mix_from_scores_local(scores, K):
#     probs = scores / (scores.sum() + 1e-12)
#     return (1.0 - LAMBDA_MIX) / float(K) + LAMBDA_MIX * probs


# def g_only(old, new):
#     return (old - new).detach() + new
# e.g., g_only(w_tau.view(-1,1), w_tau_g.view(-1,1))

#####################################################################
#Legacy code from EDM repo
#Will be removed eventually

#----------------------------------------------------------------------------
# Loss function corresponding to the variance preserving (VP) formulation
# from the paper "Score-Based Generative Modeling through Stochastic
# Differential Equations".

@persistence.persistent_class
class VPLoss:
    def __init__(self, beta_d=19.9, beta_min=0.1, epsilon_t=1e-5):
        self.beta_d = beta_d
        self.beta_min = beta_min
        self.epsilon_t = epsilon_t

    def __call__(self, net, images, labels, augment_pipe=None):
        rnd_uniform = torch.rand([images.shape[0], 1, 1, 1], device=images.device)
        sigma = self.sigma(1 + rnd_uniform * (self.epsilon_t - 1))
        weight = 1 / sigma ** 2
        y, augment_labels = augment_pipe(images) if augment_pipe is not None else (images, None)
        n = torch.randn_like(y) * sigma
        D_yn = net(y + n, sigma, labels, augment_labels=augment_labels)
        loss = weight * ((D_yn - y) ** 2)
        return loss

    def sigma(self, t):
        t = torch.as_tensor(t)
        return ((0.5 * self.beta_d * (t ** 2) + self.beta_min * t).exp() - 1).sqrt()

#----------------------------------------------------------------------------
# Loss function corresponding to the variance exploding (VE) formulation
# from the paper "Score-Based Generative Modeling through Stochastic
# Differential Equations".

@persistence.persistent_class
class VELoss:
    def __init__(self, sigma_min=0.02, sigma_max=100):
        self.sigma_min = sigma_min
        self.sigma_max = sigma_max

    def __call__(self, net, images, labels, augment_pipe=None):
        rnd_uniform = torch.rand([images.shape[0], 1, 1, 1], device=images.device)
        sigma = self.sigma_min * ((self.sigma_max / self.sigma_min) ** rnd_uniform)
        weight = 1 / sigma ** 2
        y, augment_labels = augment_pipe(images) if augment_pipe is not None else (images, None)
        n = torch.randn_like(y) * sigma
        D_yn = net(y + n, sigma, labels, augment_labels=augment_labels)
        loss = weight * ((D_yn - y) ** 2)
        return loss

#----------------------------------------------------------------------------
# Improved loss function proposed in the paper "Elucidating the Design Space
# of Diffusion-Based Generative Models" (EDM).

@persistence.persistent_class
class EDMLoss:
    def __init__(self, P_mean=-1.2, P_std=1.2, sigma_data=0.5):
        self.P_mean = P_mean
        self.P_std = P_std
        self.sigma_data = sigma_data

    def __call__(self, net, images, labels=None, augment_pipe=None):
        rnd_normal = torch.randn([images.shape[0], 1, 1, 1], device=images.device)
        sigma = (rnd_normal * self.P_std + self.P_mean).exp()
        weight = (sigma ** 2 + self.sigma_data ** 2) / (sigma * self.sigma_data) ** 2
        y, augment_labels = augment_pipe(images) if augment_pipe is not None else (images, None)
        n = torch.randn_like(y) * sigma
        D_yn = net(y + n, sigma, labels, augment_labels=augment_labels)
        loss = weight * ((D_yn - y) ** 2)
        return loss
    


#############################################################################

#--------------------------------------------------------
# Loss fn for VFM models         
@persistence.persistent_class
class VFMToyLoss:
    def __init__(self, flow_matcher_type='regular',
                 sigma_dynamics=0.1, 
                 sigma_compression=0.0
                ):
        
        #get flow matcher for u 
        assert flow_matcher_type in ['exactot', 'regular', 'sinkhorn']
        self.flow_matcher_type = flow_matcher_type
        
        if flow_matcher_type=='regular': 
            self.tau_flowmatcher = cfm.ConditionalFlowMatcher(sigma=sigma_compression)
        elif flow_matcher_type=='exactot':
            self.tau_flowmatcher = cfm.ExactOptimalTransportConditionalFlowMatcher(sigma=sigma_compression)
        elif flow_matcher_type=='sinkhorn':
            self.tau_flowmatcher = cfm.GeneralOptimalTransportConditionalFlowMatcher(sigma=sigma_compression, OTmethod = 'sinkhorn')
        else: 
            raise NotImplementedError('Only ifs, exactot or sinkhorn fm  supported!')
        
        #get flow matcher for v 
        #this is always just regular flow matcher! 
        self.t_flowmatcher = cfm.ConditionalFlowMatcher(sigma=sigma_dynamics)
        self.last_unmasked_sum = None
        
    def __call__(self, net, x0_1, xdt_1, dt, 
                 cov_dynamic_0=None, cov_dynamic_dt=None, cov_static=None):

        
        #sample taus, ts
        B = x0_1.shape[0]
        device = x0_1.device
        dt = float(dt)

        taus = torch.rand(B, device=device)
        ts = torch.rand(B, device=device) * dt

        # ---------------------------
        # Forward pass through network
        # ---------------------------
        u0_tau, ut_tau, u, v, x0_0, xdt_0, x0_0_mask, xdt_0_mask = net(
            x0_1, xdt_1, taus, ts, dt,
            self.tau_flowmatcher, self.t_flowmatcher,
            cov_dynamic_0=cov_dynamic_0,
            cov_dynamic_dt=cov_dynamic_dt,
            cov_static=cov_static,
        )

        flow_loss = (u - u0_tau)**2 #bs, dim
        dyn_loss = (v - ut_tau) **2 #bs, dim
        
        enc_pt_loss  = (x0_1 - x0_0_mask)**2
        with torch.no_grad():
            self.last_unmasked_sum = float(((x0_1 - x0_0)**2).sum().detach())
        
        enc_lie_loss = (xdt_1 - x0_1 - xdt_0 + x0_0)**2

        return torch.cat([flow_loss.unsqueeze(0),
                          dyn_loss.unsqueeze(0), enc_pt_loss.unsqueeze(0), \
                          enc_lie_loss.unsqueeze(0)], dim=0) # 4, bs, dim 
        
```
