## Supplementary material for "Dynamic Compression Flows for Neuroscience Data": Python code: networks_v7.py

```
# This work is licensed under a Creative Commons
# Attribution-NonCommercial-ShareAlike 4.0 International License.
# You should have received a copy of the license along with this
# work. If not, see http://creativecommons.org/licenses/by-nc-sa/4.0/

"""

Model architectures and preconditioning schemes used.

Includes original work from EDM repository 
along with modifications needed to implement inflationary flows
pre-conditioning for toy and image datasets.

"""

import numpy as np
import torch
import torch.nn.functional as F
import torch.distributions as D
from torch_utils import persistence
from torch.nn.functional import silu
import torch.nn as nn
import os
from torch_cfm.models.unet.unet import UNetModelWrapper 
import torch.nn.functional as F
import math
import torch.nn as nn
import torch.distributions as D
from torch_utils import distributed as dist

def _center_crop_like(x, ref):
    # x, ref: (B, C, H, W) ; crop x to ref spatial size
    _, _, h, w   = x.shape
    _, _, hr, wr = ref.shape
    if (h == hr) and (w == wr):
        return x
    dh = (h - hr) // 2
    dw = (w - wr) // 2
    return x[:, :, dh:dh+hr, dw:dw+wr]

def _center_crop_or_resize(x, target):
    # x: (B, C, H, W)
    H, W = x.shape[-2], x.shape[-1]
    if torch.is_tensor(target):
        H_t = int(target.shape[-2])
        W_t = int(target.shape[-1])
    else:
        H_t = W_t = int(target)
    if H == H_t and W == W_t:
        return x
    if H >= H_t and W >= W_t:
        top = (H - H_t) // 2
        left = (W - W_t) // 2
        return x[:, :, top:top+H_t, left:left+W_t]

    return F.interpolate(x, size=(H_t, W_t), mode='bilinear', align_corners=False)

def _prefer_gn_groups(C, prefer):
    if C % prefer == 0:
        return prefer
    for g in (32, 16, 8, 4, 2, 1):
        if C % g == 0:
            return g
    return 1

def _xy_coords_like(x):
    # x: [B,C,H,W] -> coords: [B,2,H,W] with channels [X,Y] in [-1,1]
    B, _, H, W = x.shape
    device, dtype = x.device, x.dtype
    ys = torch.linspace(-1, 1, steps=H, device=device, dtype=dtype)
    xs = torch.linspace(-1, 1, steps=W, device=device, dtype=dtype)
    yy, xx = torch.meshgrid(ys, xs, indexing='ij') 
    coords = torch.stack([xx, yy], dim=0).unsqueeze(0).expand(B, -1, H, W)
    return coords
    
#----------------------------------------------------------------------------
# Unified routine for initializing weights and biases.

def weight_init(shape, mode, fan_in, fan_out):
    if mode == 'xavier_uniform': return np.sqrt(6 / (fan_in + fan_out)) * (torch.rand(*shape) * 2 - 1)
    if mode == 'xavier_normal':  return np.sqrt(2 / (fan_in + fan_out)) * torch.randn(*shape)
    if mode == 'kaiming_uniform': return np.sqrt(3 / fan_in) * (torch.rand(*shape) * 2 - 1)
    if mode == 'kaiming_normal':  return np.sqrt(1 / fan_in) * torch.randn(*shape)
    raise ValueError(f'Invalid init mode "{mode}"')

#----------------------------------------------------------------------------
# Fully-connected layer.

@persistence.persistent_class
class Linear(torch.nn.Module):
    def __init__(self, in_features, out_features, bias=True, init_mode='kaiming_normal', init_weight=1, init_bias=0):
        super().__init__()
        self.in_features = in_features
        self.out_features = out_features
        init_kwargs = dict(mode=init_mode, fan_in=in_features, fan_out=out_features)
        self.weight = torch.nn.Parameter(weight_init([out_features, in_features], **init_kwargs) * init_weight)
        self.bias = torch.nn.Parameter(weight_init([out_features], **init_kwargs) * init_bias) if bias else None

    def forward(self, x):
        x = x @ self.weight.to(x.dtype).t()
        if self.bias is not None:
            x = x.add_(self.bias.to(x.dtype))
        return x
    

#----------------------------------------------------------------------------
# Convolutional layer with optional up/downsampling.
# Added a stride arg here. Defaults to 2, which was original/cte val in EDM code.
# Shouldn't alter original code/archs but allows more flexibility for toys. 

@persistence.persistent_class
class Conv2d(torch.nn.Module):
    def __init__(self,
        in_channels, out_channels, kernel, stride=2, bias=True, up=False, down=False,
        resample_filter=[1,1], fused_resample=False, init_mode='kaiming_normal', init_weight=1, init_bias=0,
    ):
        assert not (up and down)
        super().__init__()
        self.in_channels = in_channels
        self.out_channels = out_channels
        self.stride=stride
        self.up = up
        self.down = down
        self.fused_resample = fused_resample
        init_kwargs = dict(mode=init_mode, fan_in=in_channels*kernel*kernel, fan_out=out_channels*kernel*kernel)
        self.weight = torch.nn.Parameter(weight_init([out_channels, in_channels, kernel, kernel], **init_kwargs) * init_weight) if kernel else None
        self.bias = torch.nn.Parameter(weight_init([out_channels], **init_kwargs) * init_bias) if kernel and bias else None
        f = torch.as_tensor(resample_filter, dtype=torch.float32)
        f = f.ger(f).unsqueeze(0).unsqueeze(1) / f.sum().square()
        self.register_buffer('resample_filter', f if up or down else None)

    def forward(self, x):
        w = self.weight.to(x.dtype) if self.weight is not None else None
        b = self.bias.to(x.dtype) if self.bias is not None else None
        f = self.resample_filter.to(x.dtype) if self.resample_filter is not None else None
        w_pad = w.shape[-1] // 2 if w is not None else 0
        f_pad = (f.shape[-1] - 1) // 2 if f is not None else 0

        if self.fused_resample and self.up and w is not None:
            x = torch.nn.functional.conv_transpose2d(x, f.mul(4).tile([self.in_channels, 1, 1, 1]), groups=self.in_channels, stride=self.stride, padding=max(f_pad - w_pad, 0))
            x = torch.nn.functional.conv2d(x, w, padding=max(w_pad - f_pad, 0))
        elif self.fused_resample and self.down and w is not None:
            x = torch.nn.functional.conv2d(x, w, padding=w_pad+f_pad)
            x = torch.nn.functional.conv2d(x, f.tile([self.out_channels, 1, 1, 1]), groups=self.out_channels, stride=self.stride)
        else:
            if self.up:
                x = torch.nn.functional.conv_transpose2d(x, f.mul(4).tile([self.in_channels, 1, 1, 1]), groups=self.in_channels, stride=self.stride, padding=f_pad)
            if self.down:
                x = torch.nn.functional.conv2d(x, f.tile([self.in_channels, 1, 1, 1]), groups=self.in_channels, stride=self.stride, padding=f_pad)
            if w is not None:
                x = torch.nn.functional.conv2d(x, w, padding=w_pad) #leave stride==1 (default!!)
        if b is not None:
            x = x.add_(b.reshape(1, -1, 1, 1))
        return x


#----------------------------------------------------------------------------#

@persistence.persistent_class
class GaussianFourierProjection(torch.nn.Module):
  """
  Gaussian random features for encoding time steps.
  This is similar to implementation from Karras.
  """  
  def __init__(self, embed_dim, scale=30.):
    super().__init__()
    # Randomly sample weights during initialization. These weights are fixed 
    # during optimization and are not trainable.
    self.W = torch.nn.Parameter(torch.randn(embed_dim // 2) * scale, requires_grad=False)
  def forward(self, x):
    x_proj = x[:, None] * self.W[None, :] * 2 * np.pi
    return torch.cat([torch.sin(x_proj), torch.cos(x_proj)], dim=-1)

#----------------------------------------------------------------------------#

@persistence.persistent_class
class Dense(torch.nn.Module):
  """
  A fully connected layer that reshapes outputs to feature maps.
  """
  def __init__(self, input_dim, output_dim):
    super().__init__()
    self.dense = torch.nn.Linear(input_dim, output_dim)
  def forward(self, x):
    return self.dense(x)[..., None, None]

#----------------------------------------------------------------------------#

#Toy Conv UNet for simple img data 
@persistence.persistent_class
class ToyConvUNet(torch.nn.Module):
    """A time-dependent model built upon U-Net architecture."""
    
    def __init__(self, channels=[32, 64, 128, 256], embed_dim=256,
                 img_size=28, img_ch=1, input_size=784, cov_dim = 0,
                 use_coord_input = True, reflect_pad_first = True):
        """
        Initialize a time-dependent flow network.
        
        Args:
          channels: The number of channels for feature maps of each resolution.
          embed_dim: The dimensionality of Gaussian random feature embeddings.
          
        """
        super().__init__()
        self.use_coord_input = use_coord_input
        self.reflect_pad_first = reflect_pad_first

        #set up basic shapes for img
        self.img_size=img_size
        self.img_ch=img_ch
        self.cov_dim = cov_dim
          
        # Gaussian random feature embedding layer for time
        self.embed = torch.nn.Sequential(GaussianFourierProjection(embed_dim=embed_dim),
                                         torch.nn.Linear(embed_dim, embed_dim))
        if cov_dim > 0:
            self.cov_embed = torch.nn.Linear(cov_dim, embed_dim)
        else:
            self.cov_embed = None
        
        # Encoding layers where the resolution decreases
        # self.conv1 = torch.nn.Conv2d(img_ch, channels[0], 3, stride=1, bias=False)
        in_ch1 = img_ch + (2 if self.use_coord_input else 0)
        self.conv1 = torch.nn.Conv2d(
            in_ch1, channels[0], 3, stride=1, padding=1,
            padding_mode=('reflect' if self.reflect_pad_first else 'zeros'),
            bias=False
        )
        self.dense1 = Dense(embed_dim, 2*channels[0])
        self.gnorm1 = torch.nn.GroupNorm(_prefer_gn_groups(channels[0], 4), num_channels=channels[0],  affine=False)
        self.conv2 = torch.nn.Conv2d(channels[0], channels[1], 3, stride=2, bias=False)
        self.dense2 = Dense(embed_dim, 2*channels[1])
        self.gnorm2 = torch.nn.GroupNorm(_prefer_gn_groups(channels[1], 32), num_channels=channels[1],  affine=False)
        self.conv3 = torch.nn.Conv2d(channels[1], channels[2], 3, stride=2, bias=False)
        self.dense3 = Dense(embed_dim, 2*channels[2])
        self.gnorm3 = torch.nn.GroupNorm(_prefer_gn_groups(channels[2], 32), num_channels=channels[2],  affine=False)
        self.conv4 = torch.nn.Conv2d(channels[2], channels[3], 3, stride=2, bias=False)
        self.dense4 = Dense(embed_dim, 2*channels[3])
        self.gnorm4 = torch.nn.GroupNorm(_prefer_gn_groups(channels[3], 32), num_channels=channels[3],  affine=False)    
        
        # Decoding layers where the resolution increases
        self.tconv4 = torch.nn.ConvTranspose2d(channels[3], channels[2], 3, stride=2, bias=False)
        self.dense5 = Dense(embed_dim, 2*channels[2])
        self.tgnorm4 = torch.nn.GroupNorm(_prefer_gn_groups(channels[2], 32), num_channels=channels[2],  affine=False)
        self.tconv3 = torch.nn.ConvTranspose2d(channels[2] + channels[2], channels[1], 3, stride=2, bias=False, output_padding=1)    
        self.dense6 = Dense(embed_dim, 2*channels[1])
        self.tgnorm3 = torch.nn.GroupNorm(_prefer_gn_groups(channels[1], 32), num_channels=channels[1],  affine=False)
        self.tconv2 = torch.nn.ConvTranspose2d(channels[1] + channels[1], channels[0], 3, stride=2, bias=False, output_padding=1)    
        self.dense7 = Dense(embed_dim, 2*channels[0])
        self.tgnorm2 = torch.nn.GroupNorm(_prefer_gn_groups(channels[0], 32), num_channels=channels[0],  affine=False)
        self.tconv1 = torch.nn.ConvTranspose2d(channels[0] + channels[0], img_ch, 3, stride=1)
        
        # The swish activation function
        #self.act = lambda x: x * torch.sigmoid(x)

    @staticmethod
    def _film(h, gb, *, B=None, C=None):
        # gb: (B, 2C) from Dense(embed_dim, 2*C)
        g, b = gb.chunk(2, dim=1)  # (B,C), (B,C)
        if B is None: B = h.size(0)
        if C is None: C = h.size(1)
        return h * (1 + g.view(B, C, 1, 1)) + b.view(B, C, 1, 1)
    
    def forward(self, x, t, c=None): 
        # reshape flattened array 
        x = x.reshape(-1, self.img_ch, self.img_size, self.img_size)
        if self.use_coord_input:
            x = torch.cat([x, _xy_coords_like(x)], dim=1)
        
        # Obtain the Gaussian random feature embedding for t   
        cond  = silu(self.embed(t))
        if (self.cov_embed is not None) and (c is not None):
            cond = cond + self.cov_embed(c) 
            
        # Encoding path
        h1 = self.conv1(x)
        h1 = self.gnorm1(h1)
        h1 = self._film(h1, self.dense1(cond))
        h1 = silu(h1)

        
        h2 = self.conv2(h1)
        h2 = self.gnorm2(h2)
        h2 = self._film(h2, self.dense2(cond))
        h2 = silu(h2)
        
        h3 = self.conv3(h2)
        h3 = self.gnorm3(h3)
        h3 = self._film(h3, self.dense3(cond))
        h3 = silu(h3)

        
        h4 = self.conv4(h3)
        h4 = self.gnorm4(h4)
        h4 = self._film(h4, self.dense4(cond))
        h4 = silu(h4)
        
        # Decoding path
        h  = self.tconv4(h4)
        h  = self.tgnorm4(h)
        h  = self._film(h,  self.dense5(cond))
        h  = silu(h)

        h3 = _center_crop_or_resize(h3, h)
        h  = self.tconv3(torch.cat([h, h3], 1))
        h  = self.tgnorm3(h)
        h  = self._film(h,  self.dense6(cond))
        h  = silu(h)

        h2 = _center_crop_or_resize(h2, h)
        h  = self.tconv2(torch.cat([h, h2], 1))
        h  = self.tgnorm2(h)
        h  = self._film(h,  self.dense7(cond))
        h  = silu(h)

        h1 = _center_crop_or_resize(h1, h)
        h  = self.tconv1(torch.cat([h, h1], 1))
        
        #flatten output once again before returning 
        h  = _center_crop_or_resize(h, self.img_size)
        # h = h.reshape(-1, self.img_ch*self.img_size**2)
          
        return h

#---------------------------------------------------------------------------#
#Equivalent ToyConvUNet adapted for non-image toys...

@persistence.persistent_class
class Adapted_ToyConvUNet(torch.nn.Module):
    
    def __init__(self, channels=[32, 64, 128, 256], embed_dim=256, img_size=8, img_ch=1, input_size=2, cov_dim = 0, use_coord_input = True, reflect_pad_first = True):
        super().__init__()
        self.use_coord_input = use_coord_input
        self.reflect_pad_first = reflect_pad_first
        
        #set up basic shapes for img
        self.img_size=img_size
        self.img_ch=img_ch
        self.latent_size = img_ch*(img_size**2)
        self.cov_dim = cov_dim
        
          
        # Gaussian random feature embedding layer for time
        self.embed = torch.nn.Sequential(GaussianFourierProjection(embed_dim=embed_dim),
             torch.nn.Linear(embed_dim, embed_dim))
        self.cov_embed = torch.nn.Linear(cov_dim, embed_dim) if cov_dim > 0 else None

        
        #Up/Down-sampling linear layers for non-image toys... 
        self.up_fc = torch.nn.Linear(input_size, self.latent_size, bias=True)
        self.down_fc = torch.nn.Linear(self.latent_size, input_size, bias=True)
        # Encoding layers where the resolution decreases
        # self.conv1 = torch.nn.Conv2d(img_ch, channels[0], 3, stride=1, bias=False)
        in_ch1 = img_ch + (2 if self.use_coord_input else 0)
        self.conv1 = torch.nn.Conv2d(
            in_ch1, channels[0], 3, stride=1, padding=1,
            padding_mode=('reflect' if self.reflect_pad_first else 'zeros'),
            bias=False
        )
        self.dense1 = Dense(embed_dim, 2*channels[0])
        self.gnorm1 = torch.nn.GroupNorm(_prefer_gn_groups(channels[0], 4), num_channels=channels[0])
        self.conv2 = torch.nn.Conv2d(channels[0], channels[1], 3, stride=1, bias=False)
        self.dense2 = Dense(embed_dim, 2*channels[1])
        self.gnorm2 = torch.nn.GroupNorm(_prefer_gn_groups(channels[1], 32), num_channels=channels[1])
        self.conv3 = torch.nn.Conv2d(channels[1], channels[2], 3, stride=1, bias=False)
        self.dense3 = Dense(embed_dim, 2*channels[2])
        self.gnorm3 = torch.nn.GroupNorm(_prefer_gn_groups(channels[2], 32), num_channels=channels[2])
        
        # Decoding layers where the resolution increases
        self.tconv3 = torch.nn.ConvTranspose2d(channels[2], channels[1], 3, stride=1, bias=False)    
        self.dense4 = Dense(embed_dim, 2*channels[1])
        self.tgnorm3 = torch.nn.GroupNorm(_prefer_gn_groups(channels[1], 32), num_channels=channels[1])
        self.tconv2 = torch.nn.ConvTranspose2d(channels[1] + channels[1], channels[0], 3, stride=1, bias=False)    
        self.dense5 = Dense(embed_dim, 2*channels[0])
        self.tgnorm2 = torch.nn.GroupNorm(_prefer_gn_groups(channels[0], 32), num_channels=channels[0])
        self.tconv1 = torch.nn.ConvTranspose2d(channels[0] + channels[0], img_ch, 3, stride=1)

    @staticmethod
    def _film(h, gb):
        # gb: (B, 2C)
        g, b = gb.chunk(2, dim=1)  # (B,C), (B,C)
        B, C = h.size(0), h.size(1)
        return h * (1 + g.view(B, C, 1, 1)) + b.view(B, C, 1, 1)

    
    def forward(self, x, t, c = None): 
        
        x = self.up_fc(x)
        x = x.reshape(-1, self.img_ch, self.img_size, self.img_size)
        if self.use_coord_input:
            x = torch.cat([x, _xy_coords_like(x)], dim=1)        

        cond = silu(self.embed(t))
        if (self.cov_embed is not None) and (c is not None):
            cond = cond + self.cov_embed(c)
        
        # Encoding path
        h1 = self.conv1(x)
        h1 = self.gnorm1(h1)
        h1 = self._film(h1, self.dense1(cond))
        h1 = silu(h1)  # 6x6

        h2 = self.conv2(h1)
        h2 = self.gnorm2(h2)
        h2 = self._film(h2, self.dense2(cond))
        h2 = silu(h2)  # 4x4

        h3 = self.conv3(h2)
        h3 = self.gnorm3(h3)
        h3 = self._film(h3, self.dense3(cond))
        h3 = silu(h3)  # 2x2

        # Decoding path
        h4 = self.tconv3(h3)          # 2->4
        h4 = self.tgnorm3(h4)
        h4 = self._film(h4, self.dense4(cond))
        h4 = silu(h4)

        h5 = self.tconv2(torch.cat([h4, h2], dim=1))  # ->6
        h5 = self.tgnorm2(h5)
        h5 = self._film(h5, self.dense5(cond))
        h5 = silu(h5)

        h6 = self.tconv1(torch.cat([h5, h1], dim=1))  # ->8
        
        #now re-shape and downsample
        h6 = h6.view(-1, self.latent_size)
        out = self.down_fc(h6)  # (B, input_size)
        return out

#---------------------------------------------------------------------------#
#Simple toy MLP arch (from CFM repo)
#This can be used for any type of toy! 

@persistence.persistent_class
class ToyMLP(torch.nn.Module):
    def __init__(self, dim, out_dim=None, n_hidden=2, w=64, time_varying=False):
        super().__init__()

        time_dims = int(time_varying) if isinstance(time_varying, bool) else int(time_varying)
        if out_dim is None:
            out_dim = dim
        net = [torch.nn.Linear(dim + time_dims, w), torch.nn.SELU()]
        
        for _ in range(n_hidden):
            net.append(torch.nn.Linear(w, w))
            net.append(torch.nn.SELU())
        net.append(torch.nn.Linear(w, out_dim))
        self.net = torch.nn.Sequential(*net)

    def forward(self, x, t, t2=None):
        t = t[:, None] if t.dim() == 1 else t
        if t2 is not None:
            t2 = t2[:, None] if t2.dim() == 1 else t2
            t = torch.cat([t, t2], dim=-1)
        return self.net(torch.cat([x, t], dim=-1))


#----------------------------------------------------------------------------
#MLP Encoder 
#Can be used for any toy type!!

#MLP VAE case is tested/works fine 
@persistence.persistent_class
class Latent_MLP_VAE(torch.nn.Module):

    @torch.no_grad()
    def orthonormalize_L(self, threshold = 1e-4):
        d_full = self.input_size
        K = int(min(self.k_max, d_full))
        if K <= 0:
            return

        W = self.L_head[:d_full, :K]               # [d_full, K]
        gram = W.T @ W                              # [K, K]
        I = torch.eye(K, device=W.device, dtype=W.dtype)
        max_err = (gram - I).abs().max()

        if max_err <= threshold:
            return

        # Exact non-pivoted QR: preserves prefix structure
        Q, R = torch.linalg.qr(W, mode='reduced')   # Q: [d_full, K], R: [K, K]

        # Canonical sign fix: diag(R) >= 0
        diag_R = torch.diagonal(R)
        sign = torch.sign(diag_R)
        sign[sign == 0] = 1.0
        Q = Q * sign.unsqueeze(0)

        # Write back only the head; tail [:, K:] untouched
        self.L_head[:d_full, :K].copy_(Q)

    
    def __init__(self, input_size, output_size, k_max,
                 num_hidden, hidden_size=10, d_min=1e-15):

        super().__init__()
        self.k_max = k_max
        self.ndrop_p = getattr(self, 'ndrop_p', 0.2)
        self.d_min = d_min
        self.input_size = input_size

        self.use_pop_stats = bool(getattr(self, 'use_pop_stats', True))
        self.register_buffer('mu_pop_mean', torch.zeros(self.k_max))
        self.register_buffer('mu_pop_var',  torch.ones(self.k_max))
        self.register_buffer('mu_pop_s2',   torch.ones(self.k_max))
        self.diag_gauge_eps = float(getattr(self, 'diag_gauge_eps', 1e-6))
        self.whiten_mom = float(getattr(self, 'whiten_mom', 0.9))

        self.mu_radial_R = float(getattr(self, 'mu_radial_R', 10))
        self.mu_radial_eps = float(getattr(self, 'mu_radial_eps', 1e-6))
        
        self.register_buffer('data_mean', torch.zeros(input_size))
        self.register_buffer('data_mean_n', torch.zeros((), dtype=torch.long))
                        
        d = output_size
        self.L_head = torch.nn.Parameter(torch.empty(d, self.k_max))
        with torch.no_grad():
            W0 = torch.randn(d, self.k_max)
            Q0, _ = torch.linalg.qr(W0, mode='reduced')  # [d, k_max], Q0^T Q0 = I
            self.L_head.copy_(Q0)

        keep_cap = 1e20
        self.register_buffer('d_max_diag', torch.full((1, output_size), keep_cap))
        self.register_buffer('d_min_diag', torch.full((1, output_size), d_min))
        
        d_vec = torch.full((output_size,), math.log(10.0 * d_min))
        K = int(min(self.k_max, output_size))
        if K > 0:
            d_vec[:K] = math.log(1e-8)
        self.d = torch.nn.Parameter(d_vec)
        
        mu = [torch.nn.Linear(input_size,hidden_size,bias=True), torch.nn.ReLU()] 
        for _ in range(num_hidden):
            mu.append(torch.nn.Linear(hidden_size,hidden_size))
            mu.append(torch.nn.ReLU())
        mu.append(torch.nn.Linear(hidden_size,output_size))

        self.mu = torch.nn.Sequential(*mu)
        
        self.register_buffer('_d_grad_mask', torch.zeros_like(self.d))
        self._d_grad_mask[: self.k_max] = 1.0
        self.d.register_hook(lambda g: g * self._d_grad_mask)
        self.register_buffer('last_K', torch.tensor(0.0))

    def encode(self, x, return_data_mean = True, update_stats=True):

        mu = self.mu(x) #bs, dim
        d = mu.shape[1]
        K = int(min(self.k_max, mu.shape[1]))
        if K > 0 and self.use_pop_stats:
            # raw head from encoder
            head_raw = mu[:, :K]

            # ---- radial cap on raw head (per-sample) ----
            R   = float(self.mu_radial_R)
            eps = self.mu_radial_eps
            r2 = (head_raw * head_raw).sum(dim=1, keepdim=True)
            r  = torch.sqrt(r2 + eps)
            scale = (R / r).clamp(max=1.0) # scale <= 1
            head = head_raw * scale  # [B, K], norm <= R

            if self.training and update_stats:
                with torch.no_grad():
                    Bsz = head.size(0)
                    if Bsz > 0:
                        sum_local   = head.sum(dim=0)
                        sumsq_local = (head * head).sum(dim=0)
                        if torch.distributed.is_available() and torch.distributed.is_initialized():
                            torch.distributed.all_reduce(sum_local)
                            torch.distributed.all_reduce(sumsq_local)
                            Btot_t = torch.tensor(
                                [Bsz], device=head.device, dtype=head.dtype
                            )
                            torch.distributed.all_reduce(Btot_t)
                            Btot = int(Btot_t.item())
                        else:
                            Btot = Bsz

                        if Btot > 0:
                            mean_b = sum_local[:K] / float(Btot)
                            s2_b   = sumsq_local[:K] / float(Btot)
                            mom = float(self.whiten_mom)
                            self.mu_pop_mean[:K].mul_(mom).add_(mean_b, alpha=1.0 - mom)
                            self.mu_pop_s2[:K].mul_(mom).add_(s2_b,   alpha=1.0 - mom)
                            var_ema = self.mu_pop_s2[:K] - self.mu_pop_mean[:K].pow(2)
                            var_ema.clamp_(min=0.0)
                            self.mu_pop_var[:K].copy_(var_ema)

            # EMA-normalization
            mean_use = self.mu_pop_mean[:K]
            var_use  = self.mu_pop_var[:K]

            # s = torch.ones_like(var_use)
            # scaling_use = var_use >= self.diag_gauge_eps
            # s[scaling_use] = torch.rsqrt(var_use[scaling_use])

            var_clamped = torch.clamp(var_use, min=self.diag_gauge_eps)
            s = torch.rsqrt(var_clamped)
            mu_head = (head - mean_use) * s

            if K < d:
                mu = torch.cat([mu_head, mu[:, K:]], dim=1)
            else:
                mu = mu_head
        
        K_max = int(min(getattr(self, 'k_max', mu.shape[1]), mu.shape[1]))
        if K_max < mu.shape[1]:
            mu[:, K_max:] = 0.0
            
        d_vec = torch.exp(self.d)  # [d]
        d_vec = torch.clamp(
            d_vec,
            min=self.d_min_diag.to(x.device).squeeze(0),
            max=self.d_max_diag.to(x.device).squeeze(0)
        )

        K = int(min(getattr(self, 'k_max', mu.shape[1]), mu.shape[1]))
        d_full = mu.shape[1]
        if K > 0:
            L = self.L_head[:d_full, :K]  # [d_full, K]
        else:
            L = torch.zeros(d_full, 0, device=x.device, dtype=x.dtype)

        D = torch.diag(d_vec)

        # bias term: running mean, change to EMA if online data
        data_mean = self.data_mean
        if self.training and update_stats:
            with torch.no_grad():
                sum_local = x.sum(dim=0)  # [d]
                n_local = torch.tensor([x.size(0)],
                                       device=sum_local.device,
                                       dtype=torch.long)
                if torch.distributed.is_available() and torch.distributed.is_initialized():
                    torch.distributed.all_reduce(sum_local)
                    torch.distributed.all_reduce(n_local)
        
                n_batch = int(n_local.item())
                if n_batch > 0:
                    m_batch = sum_local / float(n_batch)  # [d]
                    n_prev = int(self.data_mean_n.item())
                    n_new  = n_prev + n_batch
                    if n_new > 0:
                        w_old = float(n_prev)  / float(n_new)
                        w_new = float(n_batch) / float(n_new)
                        self.data_mean.mul_(w_old).add_(m_batch, alpha=w_new)
                        self.data_mean_n.fill_(n_new)
            data_mean = self.data_mean
        else:
            if self.data_mean_n.item() == 0:
                raise RuntimeError("data_mean not initialized...")

        return (mu, D, L, data_mean) if return_data_mean else (mu, D, L)
    
    def rsample(self, x, K = None, update_stats=True):
        mu, D, L, data_mean = self.encode(x, return_data_mean=True, update_stats=update_stats)
        d = mu.shape[1]
        kmax = int(min(getattr(self, 'k_max', d), d))
        if self.training:
            mu_unmask = mu.clone()
            p = float(getattr(self, 'ndrop_p', 0.0))
            if p > 0.0 and kmax > 0:
                if K is None:
                    # sample once if caller did not fix it
                    K = int(
                        torch.distributions.Geometric(
                            probs=torch.tensor(p, device=mu.device)
                        ).sample().clamp_(1, kmax).item()
                    )
                K = int(max(1, min(K, kmax)))
                self.last_K.fill_(float(K))
    
                mask = mu.new_zeros(d)
                mask[:K] = 1.0
                mu = mu * mask

        d_vec  = torch.diagonal(D)[:kmax]          # [kmax]
        d_sqrt = torch.sqrt(d_vec)                 # [kmax]
        L_scaled = L[:, :kmax] * d_sqrt.unsqueeze(0)  # [d, kmax]
        
        if self.training:
            mu_unmask_head = mu_unmask[:, :kmax]
            mu_head = mu[:, :kmax]
            z_unmask = torch.einsum('ij, bjk -> bik', L_scaled, mu_unmask_head.unsqueeze(-1)).squeeze(-1) + data_mean
            z_mask = torch.einsum('ij, bjk -> bik', L_scaled, mu_head.unsqueeze(-1)).squeeze(-1) + data_mean
            return z_unmask, z_mask
        else:
            mu_head = mu[:, :kmax]
            z = torch.einsum('ij, bjk -> bik', L_scaled, mu_head.unsqueeze(-1)).squeeze(-1) + data_mean
            return z
        
    def forward(self,x):
        if self.training:
            z_unmask, z_mask = self.rsample(x)
            zero = z_mask.new_tensor(0.0)
            return z_mask, zero
        else:
            z = self.rsample(x)
            zero = z.new_tensor(0.0)
            return z, zero
            
#---------------------------------------------------------------------------

#CNN Encoder Arches for Image
@persistence.persistent_class
class Latent_CNN_VAE(torch.nn.Module):

    @torch.no_grad()
    def orthonormalize_L(self, threshold = 1e-4):
        d_full = self.input_dim
        K = int(min(self.k_max, d_full))
        if K <= 0:
            return

        W = self.L_head[:d_full, :K]               # [d_full, K]
        gram = W.T @ W                              # [K, K]
        I = torch.eye(K, device=W.device, dtype=W.dtype)
        max_err = (gram - I).abs().max()

        if max_err <= threshold:
            return

        # Exact non-pivoted QR: preserves prefix structure
        Q, R = torch.linalg.qr(W, mode='reduced')   # Q: [d_full, K], R: [K, K]

        # Canonical sign fix: diag(R) >= 0
        diag_R = torch.diagonal(R)
        sign = torch.sign(diag_R)
        sign[sign == 0] = 1.0
        Q = Q * sign.unsqueeze(0)

        # Write back only the head; tail [:, K:] untouched
        self.L_head[:d_full, :K].copy_(Q)

    
    def __init__(self, img_resolution=28, img_ch=1, k_max=784,
                 d_min=1e-15, use_coord_input = True, reflect_pad_first = True):
        
        super().__init__()
        self.k_max = k_max
        self.input_dim = img_ch*img_resolution**2
        self.d_min = d_min

        self.use_pop_stats = bool(getattr(self, 'use_pop_stats', True))
        self.register_buffer('mu_pop_mean', torch.zeros(self.k_max))
        self.register_buffer('mu_pop_var',  torch.ones(self.k_max))
        self.register_buffer('mu_pop_s2',   torch.ones(self.k_max))
        self.diag_gauge_eps = float(getattr(self, 'diag_gauge_eps', 1e-6))
        self.whiten_mom = float(getattr(self, 'whiten_mom', 0.9))

        self.mu_radial_R = float(getattr(self, 'mu_radial_R', 10))
        self.mu_radial_eps = float(getattr(self, 'mu_radial_eps', 1e-6))

        self.register_buffer('data_mean', torch.zeros(self.input_dim))
        self.register_buffer('data_mean_n', torch.zeros((), dtype=torch.long))        

        self.use_coord_input = use_coord_input
        self.reflect_pad_first = reflect_pad_first

        self.latent_size = self.input_dim
        self.img_resolution= img_resolution
        self.img_ch = img_ch

        d = self.input_dim
        self.L_head = torch.nn.Parameter(torch.empty(d, self.k_max))
        with torch.no_grad():
            W0 = torch.randn(d, self.k_max)
            Q0, _ = torch.linalg.qr(W0, mode='reduced')  # [d, k_max], Q0^T Q0 = I
            self.L_head.copy_(Q0)
        
        keep_cap = 1e20
        self.register_buffer('d_max_diag', torch.full((1, self.input_dim), keep_cap))
        self.register_buffer('d_min_diag', torch.full((1, self.input_dim), d_min))
        
        # For mean encoder
        in_ch1 = img_ch + (2 if self.use_coord_input else 0)
        self.conv1 = torch.nn.Conv2d(
            in_ch1, 16, kernel_size=5, stride=2, padding=2,
            padding_mode=('reflect' if self.reflect_pad_first else 'zeros')
        )
        self.conv2 = torch.nn.Conv2d(16, 32, kernel_size=5, stride=2)
        with torch.no_grad():
            dummy = torch.zeros(
                1, self.conv1.in_channels, self.img_resolution, self.img_resolution,
                device=self.conv1.weight.device, dtype=self.conv1.weight.dtype
            )
            t = torch.nn.functional.relu(self.conv1(dummy))
            t = torch.nn.functional.relu(self.conv2(t))
            feat_dim = t.view(1, -1).size(1)
        self.linear1 = torch.nn.Linear(feat_dim, 300)
        self.mu = torch.nn.Linear(300, self.latent_size)
        
        d_vec = torch.full((self.input_dim,), math.log(10.0 * d_min))  # tail: near floor
        K = int(min(self.k_max, self.input_dim))
        if K > 0:
            d0 = 1.0
            d_vec[:K] = math.log(d0)
        self.d = torch.nn.Parameter(d_vec)

        self.register_buffer('_d_grad_mask', torch.zeros_like(self.d))
        self._d_grad_mask[: self.k_max] = 1.0
        self.d.register_hook(lambda g: g * self._d_grad_mask)
        self.register_buffer('last_K', torch.tensor(0.0))
    
    def encode(self, x, return_data_mean = True, update_stats=True):
        #get mu, u, d
        x_img = x.reshape(-1, self.img_ch, self.img_resolution, self.img_resolution)
        x_flat = x_img.view(x_img.size(0), -1)
        x = torch.cat([x_img, _xy_coords_like(x_img)], dim=1) if self.use_coord_input else x_img
        
        t = torch.nn.functional.relu(self.conv1(x))
        t = torch.nn.functional.relu(self.conv2(t))
        t = t.reshape((x.shape[0], -1))
        
        t = torch.nn.functional.relu(self.linear1(t))
        mu = self.mu(t)
        d = mu.shape[1]
        K = int(min(getattr(self, 'k_max', d), d))
        if K > 0 and self.use_pop_stats:
            # raw head from encoder
            head_raw = mu[:, :K]

            # ---- radial cap on raw head (per-sample) ----
            R   = float(self.mu_radial_R)
            eps = self.mu_radial_eps
            r2 = (head_raw * head_raw).sum(dim=1, keepdim=True)
            r  = torch.sqrt(r2 + eps)
            scale = (R / r).clamp(max=1.0) # scale <= 1
            head = head_raw * scale  # [B, K], norm <= R

            if self.training and update_stats:
                with torch.no_grad():
                    Bsz = head.size(0)
                    if Bsz > 0:
                        sum_local   = head.sum(dim=0)
                        sumsq_local = (head * head).sum(dim=0)
                        if torch.distributed.is_available() and torch.distributed.is_initialized():
                            torch.distributed.all_reduce(sum_local)
                            torch.distributed.all_reduce(sumsq_local)
                            Btot_t = torch.tensor(
                                [Bsz], device=head.device, dtype=head.dtype
                            )
                            torch.distributed.all_reduce(Btot_t)
                            Btot = int(Btot_t.item())
                        else:
                            Btot = Bsz

                        if Btot > 0:
                            mean_b = sum_local[:K] / float(Btot)
                            s2_b   = sumsq_local[:K] / float(Btot)
                            mom = float(self.whiten_mom)
                            self.mu_pop_mean[:K].mul_(mom).add_(mean_b, alpha=1.0 - mom)
                            self.mu_pop_s2[:K].mul_(mom).add_(s2_b,   alpha=1.0 - mom)
                            var_ema = self.mu_pop_s2[:K] - self.mu_pop_mean[:K].pow(2)
                            var_ema.clamp_(min=0.0)
                            self.mu_pop_var[:K].copy_(var_ema)

            # EMA-normalization
            mean_use = self.mu_pop_mean[:K]
            var_use  = self.mu_pop_var[:K]

            s = torch.ones_like(var_use)
            scaling_use = var_use >= self.diag_gauge_eps
            s[scaling_use] = torch.rsqrt(var_use[scaling_use])

            mu_head = (head - mean_use) * s

            if K < d:
                mu = torch.cat([mu_head, mu[:, K:]], dim=1)
            else:
                mu = mu_head

        K_max = int(min(getattr(self, 'k_max', mu.shape[1]), mu.shape[1]))
        if K_max < mu.shape[1]:
            mu[:, K_max:] = 0.0

        d_vec = torch.exp(self.d)  # [d]
        d_vec = torch.clamp(
            d_vec,
            min=self.d_min_diag.to(x.device).squeeze(0),
            max=self.d_max_diag.to(x.device).squeeze(0)
        )
        
        K = int(min(getattr(self, 'k_max', mu.shape[1]), mu.shape[1]))
        d_full = mu.shape[1]
        if K > 0:
            L = self.L_head[:d_full, :K]  # [d_full, K]
        else:
            L = torch.zeros(d_full, 0, device=x.device, dtype=x.dtype)
        
        D = torch.diag(d_vec)

        # bias term: running mean, change to EMA if online data
        data_mean = self.data_mean
        if self.training and update_stats:
            with torch.no_grad():
                sum_local = x_flat.sum(dim=0)  # [d]
                n_local = torch.tensor([x_flat.size(0)],
                                       device=sum_local.device,
                                       dtype=torch.long)
                if torch.distributed.is_available() and torch.distributed.is_initialized():
                    torch.distributed.all_reduce(sum_local)
                    torch.distributed.all_reduce(n_local)
        
                n_batch = int(n_local.item())
                if n_batch > 0:
                    m_batch = sum_local / float(n_batch)  # [d]
                    n_prev = int(self.data_mean_n.item())
                    n_new  = n_prev + n_batch
                    if n_new > 0:
                        w_old = float(n_prev)  / float(n_new)
                        w_new = float(n_batch) / float(n_new)
                        self.data_mean.mul_(w_old).add_(m_batch, alpha=w_new)
                        self.data_mean_n.fill_(n_new)
            data_mean = self.data_mean
        else:
            if self.data_mean_n.item() == 0:
                raise RuntimeError("data_mean not initialized...")

        return (mu, D, L, data_mean) if return_data_mean else (mu, D, L)
          
    def rsample(self, x, K = None, update_stats=True):
        mu, D, L, data_mean = self.encode(x, return_data_mean=True, update_stats=update_stats)
        d = mu.shape[1]
        kmax = int(min(getattr(self, 'k_max', d), d))
        if self.training:
            mu_unmask = mu.clone()
            p = float(getattr(self, 'ndrop_p', 0.0))
            if p > 0.0 and kmax > 0:
                if K is None:
                    # sample once if caller did not fix it
                    K = int(
                        torch.distributions.Geometric(
                            probs=torch.tensor(p, device=mu.device)
                        ).sample().clamp_(1, kmax).item()
                    )
                K = int(max(1, min(K, kmax)))
                self.last_K.fill_(float(K))
    
                mask = mu.new_zeros(d)
                mask[:K] = 1.0
                mu = mu * mask


        d_vec  = torch.diagonal(D)[:kmax]          # [kmax]
        d_sqrt = torch.sqrt(d_vec)                 # [kmax]
        L_scaled = L[:, :kmax] * d_sqrt.unsqueeze(0)  # [d, kmax]
        
        if self.training:
            mu_unmask_head = mu_unmask[:, :kmax]
            mu_head = mu[:, :kmax]
            z_unmask = torch.einsum('ij, bjk -> bik', L_scaled, mu_unmask_head.unsqueeze(-1)).squeeze(-1) + data_mean
            z_mask = torch.einsum('ij, bjk -> bik', L_scaled, mu_head.unsqueeze(-1)).squeeze(-1) + data_mean
            return z_unmask, z_mask
        else:
            mu_head = mu[:, :kmax]
            z = torch.einsum('ij, bjk -> bik', L_scaled, mu_head.unsqueeze(-1)).squeeze(-1) + data_mean
            return z
    
    def forward(self, x):
        if self.training:
            z_unmask, z_mask = self.rsample(x)
            zero = z_mask.new_tensor(0.0)
            return z_mask, zero
        else:
            z = self.rsample(x)
            zero = z.new_tensor(0.0)
            return z, zero

@persistence.persistent_class
class Latent_LargeCNN_VAE(torch.nn.Module):

    @torch.no_grad()
    def orthonormalize_L(self, threshold = 1e-4):
        d_full = self.input_dim
        K = int(min(self.k_max, d_full))
        if K <= 0:
            return

        W = self.L_head[:d_full, :K]               # [d_full, K]
        gram = W.T @ W                              # [K, K]
        I = torch.eye(K, device=W.device, dtype=W.dtype)
        max_err = (gram - I).abs().max()

        if max_err <= threshold:
            return

        # Exact non-pivoted QR: preserves prefix structure
        Q, R = torch.linalg.qr(W, mode='reduced')   # Q: [d_full, K], R: [K, K]

        # Canonical sign fix: diag(R) >= 0
        diag_R = torch.diagonal(R)
        sign = torch.sign(diag_R)
        sign[sign == 0] = 1.0
        Q = Q * sign.unsqueeze(0)

        # Write back only the head; tail [:, K:] untouched
        self.L_head[:d_full, :K].copy_(Q)
    
    def __init__(self, channels=[32, 64, 128, 256], img_size=28, img_ch=1,
                 k_max=784, d_min=1e-15,
                 use_coord_input = True, reflect_pad_first = True):
        super().__init__()
        self.use_coord_input = use_coord_input
        self.reflect_pad_first = reflect_pad_first
        self.k_max = k_max
        self.input_dim = img_ch*img_size**2
        self.d_min = d_min

        self.use_pop_stats = bool(getattr(self, 'use_pop_stats', True))
        self.register_buffer('mu_pop_mean', torch.zeros(self.k_max))
        self.register_buffer('mu_pop_var',  torch.ones(self.k_max))
        self.register_buffer('mu_pop_s2',   torch.ones(self.k_max))
        self.diag_gauge_eps = float(getattr(self, 'diag_gauge_eps', 1e-6))
        self.whiten_mom = float(getattr(self, 'whiten_mom', 0.9))

        self.mu_radial_R = float(getattr(self, 'mu_radial_R', 10))
        self.mu_radial_eps = float(getattr(self, 'mu_radial_eps', 1e-6))

        self.register_buffer('data_mean', torch.zeros(self.input_dim))
        self.register_buffer('data_mean_n', torch.zeros((), dtype=torch.long))
        
        #set up general attributes
        self.img_size=img_size
        self.img_ch=img_ch

        d = self.input_dim
        self.L_head = torch.nn.Parameter(torch.empty(d, self.k_max))
        
        with torch.no_grad():
            W0 = torch.randn(d, self.k_max)
            Q0, _ = torch.linalg.qr(W0, mode='reduced')  # [d, k_max], Q0^T Q0 = I
            self.L_head.copy_(Q0)
        
        #set d_max tensor
        keep_cap = 1e20 # no upper bound...
        self.register_buffer('d_max_diag', torch.full((1, self.input_dim), keep_cap))
        self.register_buffer('d_min_diag', torch.full((1, self.input_dim), d_min))
        
        # Conv layers with decreasing res 
        in_ch1 = img_ch + (2 if self.use_coord_input else 0)
        self.conv1 = torch.nn.Conv2d(
            in_ch1, channels[0], 3, stride=1, padding=1,
            padding_mode=('reflect' if self.reflect_pad_first else 'zeros'),
            bias=False)
        # self.conv1 = torch.nn.Conv2d(img_ch, channels[0], 3, stride=1, bias=False)
        self.gnorm1 = torch.nn.GroupNorm(_prefer_gn_groups(channels[0], 4), num_channels=channels[0])
        
        self.conv2 = torch.nn.Conv2d(channels[0], channels[1], 3, stride=2, bias=False)
        self.gnorm2 = torch.nn.GroupNorm(_prefer_gn_groups(channels[1], 32), num_channels=channels[1])
        
        self.conv3 = torch.nn.Conv2d(channels[1], channels[2], 3, stride=2, bias=False)
        self.gnorm3 = torch.nn.GroupNorm(_prefer_gn_groups(channels[2], 32), num_channels=channels[2])
        
        self.conv4 = torch.nn.Conv2d(channels[2], channels[3], 3, stride=2, bias=False)
        self.gnorm4 = torch.nn.GroupNorm(_prefer_gn_groups(channels[3], 32), num_channels=channels[3])    
        
        #Linear Layers to extract mus
        with torch.no_grad():
            dummy = torch.zeros(
                1, self.conv1.in_channels, img_size, img_size,
                device=self.conv1.weight.device, dtype=self.conv1.weight.dtype
            )
            h1 = F.silu(self.gnorm1(self.conv1(dummy)))
            h2 = F.silu(self.gnorm2(self.conv2(h1)))
            h3 = F.silu(self.gnorm3(self.conv3(h2)))
            h4 = F.silu(self.gnorm4(self.conv4(h3)))
            feat_dim = h4.view(1, -1).size(1)
        self.mu = torch.nn.Linear(feat_dim, self.input_dim)

        # self.U = torch.nn.Parameter(torch.randn(d, r))
        d_vec = torch.full((self.input_dim,), math.log(10.0 * d_min))  # tail: near floor
        K = int(min(self.k_max, self.input_dim))
        if K > 0:
            d0 = 1e-2
            d_vec[:K] = math.log(d0)
        self.d = torch.nn.Parameter(d_vec)
        
        self.register_buffer('_d_grad_mask', torch.zeros_like(self.d))
        self._d_grad_mask[: self.k_max] = 1.0
        self.d.register_hook(lambda g: g * self._d_grad_mask)
        self.register_buffer('last_K', torch.tensor(0.0))
        
    def encode(self, x, return_data_mean = True, update_stats=True):
        # Reshape flattened array 
        x_img = x.reshape(-1, self.img_ch, self.img_size, self.img_size)
        x_flat = x_img.view(x_img.size(0), -1)
        x = torch.cat([x_img, _xy_coords_like(x_img)], dim=1) if self.use_coord_input else x_img

        # Encoder Conv/Gnorm BLocks
        h1 = silu(self.gnorm1(self.conv1(x)))
        h2 = silu(self.gnorm2(self.conv2(h1)))
        h3 = silu(self.gnorm3(self.conv3(h2)))
        h4 = silu(self.gnorm4(self.conv4(h3)))
        
        #Extract mu, d, u
        mu = self.mu(h4.reshape(x.shape[0], -1))
        d = mu.shape[1]
        K = int(min(getattr(self, 'k_max', d), d))
        if K > 0 and self.use_pop_stats:
            # raw head from encoder
            head_raw = mu[:, :K]

            # ---- radial cap on raw head (per-sample) ----
            R   = float(self.mu_radial_R)
            eps = self.mu_radial_eps
            r2 = (head_raw * head_raw).sum(dim=1, keepdim=True)
            r  = torch.sqrt(r2 + eps)
            scale = (R / r).clamp(max=1.0) # scale <= 1
            head = head_raw * scale  # [B, K], norm <= R
            

            if self.training and update_stats:
                with torch.no_grad():
                    Bsz = head.size(0)
                    if Bsz > 0:
                        sum_local   = head.sum(dim=0)
                        sumsq_local = (head * head).sum(dim=0)
                        if torch.distributed.is_available() and torch.distributed.is_initialized():
                            torch.distributed.all_reduce(sum_local)
                            torch.distributed.all_reduce(sumsq_local)
                            Btot_t = torch.tensor(
                                [Bsz], device=head.device, dtype=head.dtype
                            )
                            torch.distributed.all_reduce(Btot_t)
                            Btot = int(Btot_t.item())
                        else:
                            Btot = Bsz

                        if Btot > 0:
                            mean_b = sum_local[:K] / float(Btot)
                            s2_b   = sumsq_local[:K] / float(Btot)
                            mom = float(self.whiten_mom)
                            self.mu_pop_mean[:K].mul_(mom).add_(mean_b, alpha=1.0 - mom)
                            self.mu_pop_s2[:K].mul_(mom).add_(s2_b,   alpha=1.0 - mom)
                            var_ema = self.mu_pop_s2[:K] - self.mu_pop_mean[:K].pow(2)
                            var_ema.clamp_(min=0.0)
                            self.mu_pop_var[:K].copy_(var_ema)

            # EMA-normalization
            mean_use = self.mu_pop_mean[:K]
            var_use  = self.mu_pop_var[:K]

            # s = torch.ones_like(var_use)
            # scaling_use = var_use >= self.diag_gauge_eps
            # s[scaling_use] = torch.rsqrt(var_use[scaling_use])

            var_clamped = torch.clamp(var_use, min=self.diag_gauge_eps)
            s = torch.rsqrt(var_clamped)
            mu_head = (head - mean_use) * s

            if K < d:
                mu = torch.cat([mu_head, mu[:, K:]], dim=1)
            else:
                mu = mu_head

        K_max = int(min(getattr(self, 'k_max', mu.shape[1]), mu.shape[1]))
        if K_max < mu.shape[1]:
            mu[:, K_max:] = 0.0

        d_vec = torch.exp(self.d)  # [d]
        d_vec = torch.clamp(
            d_vec,
            min=self.d_min_diag.to(x.device).squeeze(0),
            max=self.d_max_diag.to(x.device).squeeze(0)
        )
        
        K = int(min(getattr(self, 'k_max', mu.shape[1]), mu.shape[1]))
        d_full = mu.shape[1]
        
        if K > 0:
            L = self.L_head[:d_full, :K]  # [d_full, K]
        else:
            L = torch.zeros(d_full, 0, device=x.device, dtype=x.dtype)
        
        D = torch.diag(d_vec)

        # bias term: running mean, change to EMA if online data
        data_mean = self.data_mean
        if self.training and update_stats:
            with torch.no_grad():
                sum_local = x_flat.sum(dim=0)  # [d]
                n_local = torch.tensor([x_flat.size(0)],
                                       device=sum_local.device,
                                       dtype=torch.long)
                if torch.distributed.is_available() and torch.distributed.is_initialized():
                    torch.distributed.all_reduce(sum_local)
                    torch.distributed.all_reduce(n_local)
        
                n_batch = int(n_local.item())
                if n_batch > 0:
                    m_batch = sum_local / float(n_batch)  # [d]
                    n_prev = int(self.data_mean_n.item())
                    n_new  = n_prev + n_batch
                    if n_new > 0:
                        w_old = float(n_prev)  / float(n_new)
                        w_new = float(n_batch) / float(n_new)
                        self.data_mean.mul_(w_old).add_(m_batch, alpha=w_new)
                        self.data_mean_n.fill_(n_new)
            data_mean = self.data_mean
        else:
            if self.data_mean_n.item() == 0:
                raise RuntimeError("data_mean not initialized...")

        return (mu, D, L, data_mean) if return_data_mean else (mu, D, L)
    
    def rsample(self, x, K = None, update_stats=True):        
        mu, D, L, data_mean = self.encode(x, return_data_mean=True, update_stats=update_stats)
        d = mu.shape[1]
        kmax = int(min(getattr(self, 'k_max', d), d))
        if self.training:
            mu_unmask = mu.clone()
            p = float(getattr(self, 'ndrop_p', 0.0))
            if p > 0.0 and kmax > 0:
                if K is None:
                    K = int(
                        torch.distributions.Geometric(
                            probs=torch.tensor(p, device=mu.device)
                        ).sample().clamp_(1, kmax).item()
                    )
                K = int(max(1, min(K, kmax)))
                self.last_K.fill_(float(K))
    
                mask = mu.new_zeros(d)
                mask[:K] = 1.0
                mu = mu * mask

        # scale only first kmax columns of L by sqrt(D_ii)
        d_vec  = torch.diagonal(D)[:kmax]          # [kmax]
        d_sqrt = torch.sqrt(d_vec)                 # [kmax]
        L_scaled = L[:, :kmax] * d_sqrt.unsqueeze(0)  # [d, kmax]
        
        if self.training:
            mu_unmask_head = mu_unmask[:, :kmax]
            mu_head = mu[:, :kmax]
            z_unmask = torch.einsum('ik,bk->bi', L_scaled, mu_unmask_head) + data_mean
            z_mask = torch.einsum('ik,bk->bi', L_scaled, mu_head) + data_mean
            return z_unmask, z_mask
        else:
            mu_head = mu[:, :kmax]
            z = torch.einsum('ik,bk->bi', L_scaled, mu_head) + data_mean
            return z
    
    def forward(self, x):
        if self.training:
            z_unmask, z_mask = self.rsample(x)
            zero = z_mask.new_tensor(0.0)
            return z_mask, zero
        else:
            z = self.rsample(x)
            zero = z.new_tensor(0.0)
            return z, zero
        
#----------------------------------------------------------------------------

#version of LargeCNN Enc above for non-image toys
#needs testing on regular (non-image) toys ... 
@persistence.persistent_class
class Adapted_Latent_LargeCNN_VAE(torch.nn.Module):

    @torch.no_grad()
    def orthonormalize_L(self, threshold = 1e-4):
        d_full = self.input_size
        K = int(min(self.k_max, d_full))
        if K <= 0:
            return

        W = self.L_head[:d_full, :K]               # [d_full, K]
        gram = W.T @ W                              # [K, K]
        I = torch.eye(K, device=W.device, dtype=W.dtype)
        max_err = (gram - I).abs().max()

        if max_err <= threshold:
            return

        # Exact non-pivoted QR: preserves prefix structure
        Q, R = torch.linalg.qr(W, mode='reduced')   # Q: [d_full, K], R: [K, K]

        # Canonical sign fix: diag(R) >= 0
        diag_R = torch.diagonal(R)
        sign = torch.sign(diag_R)
        sign[sign == 0] = 1.0
        Q = Q * sign.unsqueeze(0)

        # Write back only the head; tail [:, K:] untouched
        self.L_head[:d_full, :K].copy_(Q)
    
    def __init__(self, channels=[32, 64, 128, 256], img_size=8, img_ch=1,
                 input_size=2, k_max=2, d_min=1e-15, 
                 use_coord_input = True, reflect_pad_first = True):
        super().__init__()

        self.use_coord_input = use_coord_input
        self.reflect_pad_first = reflect_pad_first
        self.k_max = k_max
        self.d_min = d_min

        #set up basic shapes for img
        self.img_size=img_size
        self.img_ch=img_ch
        self.latent_size = img_ch*img_size**2 
        self.input_size = input_size

        self.use_pop_stats = bool(getattr(self, 'use_pop_stats', True))
        self.register_buffer('mu_pop_mean', torch.zeros(self.k_max))
        self.register_buffer('mu_pop_var',  torch.ones(self.k_max))
        self.register_buffer('mu_pop_s2',   torch.ones(self.k_max))
        self.diag_gauge_eps = float(getattr(self, 'diag_gauge_eps', 1e-8))
        self.whiten_mom = float(getattr(self, 'whiten_mom', 0.9))

        self.mu_radial_R = float(getattr(self, 'mu_radial_R', 10))
        self.mu_radial_eps = float(getattr(self, 'mu_radial_eps', 1e-6))
        
        self.register_buffer('data_mean', torch.zeros(self.input_size))
        self.register_buffer('data_mean_n', torch.zeros((), dtype=torch.long))
        
        d = input_size
        self.L_head = torch.nn.Parameter(torch.empty(d, self.k_max))
        with torch.no_grad():
            W0 = torch.randn(d, self.k_max)
            Q0, _ = torch.linalg.qr(W0, mode='reduced')  # [d, k_max], Q0^T Q0 = I
            self.L_head.copy_(Q0)
            
        #set d_max tensor
        keep_cap = 1e20
        self.register_buffer('d_max_diag', torch.full((1, self.input_size), keep_cap))
        self.register_buffer('d_min_diag', torch.full((1, self.input_size), d_min))
        
        #Upsampling linear layers for non-image toys... 
        self.up_fc = torch.nn.Linear(input_size, self.latent_size, bias=True)
          
        # Encoding layers where the resolution decreases
        # self.conv1 = torch.nn.Conv2d(img_ch, channels[0], 3, stride=1, bias=False)
        in_ch1 = img_ch + (2 if self.use_coord_input else 0)
        self.conv1 = torch.nn.Conv2d(
            in_ch1, channels[0], 3, stride=1, padding=1,
            padding_mode=('reflect' if self.reflect_pad_first else 'zeros'),
            bias=False
        )
        
        self.gnorm1 = torch.nn.GroupNorm(_prefer_gn_groups(channels[0], 4), num_channels=channels[0])
        self.conv2 = torch.nn.Conv2d(channels[0], channels[1], 3, stride=1, bias=False)
        self.gnorm2 = torch.nn.GroupNorm(_prefer_gn_groups(channels[1], 32), num_channels=channels[1])
        self.conv3 = torch.nn.Conv2d(channels[1], channels[2], 3, stride=1, bias=False)
        self.gnorm3 = torch.nn.GroupNorm(_prefer_gn_groups(channels[2], 32), num_channels=channels[2])
          
        #general linear layer before extracting mu, d, u
        self.gap = torch.nn.AdaptiveAvgPool2d((2, 2))
        with torch.no_grad():
            dummy = torch.zeros(
                1, self.conv1.in_channels, img_size, img_size,
                device=self.conv1.weight.device, dtype=self.conv1.weight.dtype
            )
            d1 = F.silu(self.gnorm1(self.conv1(dummy)))
            d2 = F.silu(self.gnorm2(self.conv2(d1)))
            d3 = F.silu(self.gnorm3(self.conv3(d2)))
            d3 = self.gap(d3)
            feat_dim = d3.view(1, -1).size(1)
        self.fc1 = torch.nn.Linear(feat_dim, 100)
        self.mu = torch.nn.Linear(100, input_size)
        
        #now add global params d, u
        d_vec = torch.full((self.input_size,), math.log(10.0 * d_min))
        K = int(min(self.k_max, self.input_size))
        if K > 0:
            d0 = 1.0
            d_vec[:K] = math.log(d0)
        self.d = torch.nn.Parameter(d_vec)
        
        self.register_buffer('_d_grad_mask', torch.zeros_like(self.d))
        self._d_grad_mask[: self.k_max] = 1.0
        self.d.register_hook(lambda g: g * self._d_grad_mask)
        self.register_buffer('last_K', torch.tensor(0.0))
        
    def encode(self, x, return_data_mean = True, update_stats=True):
        
        x_in = x
        x = self.up_fc(x)
        x_img = x.reshape(-1, self.img_ch, self.img_size, self.img_size)
        x_flat = x_img.view(x_img.size(0), -1)
        x = torch.cat([x_img, _xy_coords_like(x_img)], dim=1) if self.use_coord_input else x_img

        # Feed through CNN Encoding path
        h1 = silu(self.gnorm1(self.conv1(x)))      
        h2 = silu(self.gnorm2(self.conv2(h1)))      
        h3 = silu(self.gnorm3(self.conv3(h2)))
        h3 = self.gap(h3)
        h4 = self.fc1(h3.reshape(x.shape[0], -1))
        
        mu = self.mu(h4)
        d = mu.shape[1]
        K = int(min(getattr(self, 'k_max', d), d))
        if K > 0 and self.use_pop_stats:
            # raw head from encoder
            head_raw = mu[:, :K]

            # ---- radial cap on raw head (per-sample) ----
            R   = float(self.mu_radial_R)
            eps = self.mu_radial_eps
            r2 = (head_raw * head_raw).sum(dim=1, keepdim=True)
            r  = torch.sqrt(r2 + eps)
            scale = (R / r).clamp(max=1.0) # scale <= 1
            head = head_raw * scale  # [B, K], norm <= R
            
            if self.training and update_stats:
                with torch.no_grad():
                    Bsz = head.size(0)
                    if Bsz > 0:
                        sum_local   = head.sum(dim=0)
                        sumsq_local = (head * head).sum(dim=0)
                        if torch.distributed.is_available() and torch.distributed.is_initialized():
                            torch.distributed.all_reduce(sum_local)
                            torch.distributed.all_reduce(sumsq_local)
                            Btot_t = torch.tensor(
                                [Bsz], device=head.device, dtype=head.dtype
                            )
                            torch.distributed.all_reduce(Btot_t)
                            Btot = int(Btot_t.item())
                        else:
                            Btot = Bsz

                        if Btot > 0:
                            mean_b = sum_local[:K] / float(Btot)
                            s2_b   = sumsq_local[:K] / float(Btot)
                            mom = float(self.whiten_mom)
                            self.mu_pop_mean[:K].mul_(mom).add_(mean_b, alpha=1.0 - mom)
                            self.mu_pop_s2[:K].mul_(mom).add_(s2_b,   alpha=1.0 - mom)
                            var_ema = self.mu_pop_s2[:K] - self.mu_pop_mean[:K].pow(2)
                            var_ema.clamp_(min=0.0)
                            self.mu_pop_var[:K].copy_(var_ema)

            # EMA-normalization
            mean_use = self.mu_pop_mean[:K]
            var_use  = self.mu_pop_var[:K]

            s = torch.ones_like(var_use)
            scaling_use = var_use >= self.diag_gauge_eps
            s[scaling_use] = torch.rsqrt(var_use[scaling_use])

            mu_head = (head - mean_use) * s

            if K < d:
                mu = torch.cat([mu_head, mu[:, K:]], dim=1)
            else:
                mu = mu_head

        K_max = int(min(getattr(self, 'k_max', mu.shape[1]), mu.shape[1]))
        if K_max < mu.shape[1]:
            mu[:, K_max:] = 0.0
            
        d_vec = torch.exp(self.d)  # [d]
        d_vec = torch.clamp(
            d_vec,
            min=self.d_min_diag.to(x.device).squeeze(0),
            max=self.d_max_diag.to(x.device).squeeze(0)
        )

        K = int(min(getattr(self, 'k_max', mu.shape[1]), mu.shape[1]))
        d_full = mu.shape[1]
        if K > 0:
            L = self.L_head[:d_full, :K]  # [d_full, K]
        else:
            L = torch.zeros(d_full, 0, device=x.device, dtype=x.dtype)

        D = torch.diag(d_vec)

        # bias term: running mean, change to EMA if online data
        data_mean = self.data_mean
        if self.training and update_stats:
            with torch.no_grad():
                sum_local = x_in.sum(dim=0)  # [d]
                n_local = torch.tensor([x_in.size(0)],
                                       device=sum_local.device,
                                       dtype=torch.long)
                if torch.distributed.is_available() and torch.distributed.is_initialized():
                    torch.distributed.all_reduce(sum_local)
                    torch.distributed.all_reduce(n_local)
        
                n_batch = int(n_local.item())
                if n_batch > 0:
                    m_batch = sum_local / float(n_batch)  # [d]
                    n_prev = int(self.data_mean_n.item())
                    n_new  = n_prev + n_batch
                    if n_new > 0:
                        w_old = float(n_prev)  / float(n_new)
                        w_new = float(n_batch) / float(n_new)
                        self.data_mean.mul_(w_old).add_(m_batch, alpha=w_new)
                        self.data_mean_n.fill_(n_new)
            data_mean = self.data_mean
        else:
            if self.data_mean_n.item() == 0:
                raise RuntimeError("data_mean not initialized...")
          
        return (mu, D, L, data_mean) if return_data_mean else (mu, D, L)
        
    def rsample(self, x, K = None, update_stats=True):
        mu, D, L, data_mean = self.encode(x, return_data_mean=True, update_stats=update_stats)
        d = mu.shape[1]
        kmax = int(min(getattr(self, 'k_max', d), d))
        if self.training:
            mu_unmask = mu.clone()
            p = float(getattr(self, 'ndrop_p', 0.0))
            if p > 0.0 and kmax > 0:
                if K is None:
                    # sample once if caller did not fix it
                    K = int(
                        torch.distributions.Geometric(
                            probs=torch.tensor(p, device=mu.device)
                        ).sample().clamp_(1, kmax).item()
                    )
                K = int(max(1, min(K, kmax)))
                self.last_K.fill_(float(K))
    
                mask = mu.new_zeros(d)
                mask[:K] = 1.0
                mu = mu * mask
                
        # scale only first kmax columns of L by sqrt(D_ii)
        d_vec  = torch.diagonal(D)[:kmax]          # [kmax]
        d_sqrt = torch.sqrt(d_vec)                 # [kmax]
        L_scaled = L[:, :kmax] * d_sqrt.unsqueeze(0)  # [d, kmax]

        if self.training:
            mu_unmask_head = mu_unmask[:, :kmax] 
            mu_head = mu[:, :kmax]
            z_unmask = mu_unmask_head @ L_scaled.T + data_mean
            z_mask = mu_head @ L_scaled.T + data_mean
            return z_unmask, z_mask
        else:
            mu_head = mu[:, :kmax]
            z = mu_head @ L_scaled.T + data_mean
            return z
    
    def forward(self, x):
        if self.training:
            z_unmask, z_mask = self.rsample(x)
            zero = z_mask.new_tensor(0.0)
            return z_mask, zero
        else:
            z = self.rsample(x)
            zero = z.new_tensor(0.0)
            return z, zero

#----------------------------------------------------------------------------
# Group normalization.

@persistence.persistent_class
class GroupNorm(torch.nn.Module):
    def __init__(self, num_channels, num_groups=32, min_channels_per_group=4, eps=1e-5):
        super().__init__()
        self.num_groups = min(num_groups, num_channels // min_channels_per_group)
        self.eps = eps
        self.weight = torch.nn.Parameter(torch.ones(num_channels))
        self.bias = torch.nn.Parameter(torch.zeros(num_channels))

    def forward(self, x):
        x = torch.nn.functional.group_norm(x, num_groups=self.num_groups, weight=self.weight.to(x.dtype), bias=self.bias.to(x.dtype), eps=self.eps)
        return x

#----------------------------------------------------------------------------
# Attention weight computation, i.e., softmax(Q^T * K).
# Performs all computation using FP32, but uses the original datatype for
# inputs/outputs/gradients to conserve memory.

class AttentionOp(torch.autograd.Function):
    @staticmethod
    def forward(ctx, q, k):
        w = torch.einsum('ncq,nck->nqk', q.to(torch.float32), (k / np.sqrt(k.shape[1])).to(torch.float32)).softmax(dim=2).to(q.dtype)
        ctx.save_for_backward(q, k, w)
        return w

    @staticmethod
    def backward(ctx, dw):
        q, k, w = ctx.saved_tensors
        db = torch._softmax_backward_data(grad_output=dw.to(torch.float32), output=w.to(torch.float32), dim=2, input_dtype=torch.float32)
        dq = torch.einsum('nck,nqk->ncq', k.to(torch.float32), db).to(q.dtype) / np.sqrt(k.shape[1])
        dk = torch.einsum('ncq,nqk->nck', q.to(torch.float32), db).to(k.dtype) / np.sqrt(k.shape[1])
        return dq, dk

#----------------------------------------------------------------------------
# Unified U-Net block with optional up/downsampling and self-attention.
# Represents the union of all features employed by the DDPM++, NCSN++, and
# ADM architectures.

@persistence.persistent_class
class UNetBlock(torch.nn.Module):
    def __init__(self,
        in_channels, out_channels, emb_channels, up=False, down=False, attention=False,
        num_heads=None, channels_per_head=64, dropout=0, skip_scale=1, eps=1e-5,
        resample_filter=[1,1], resample_proj=False, adaptive_scale=True,
        init=dict(), init_zero=dict(init_weight=0), init_attn=None,
    ):
        super().__init__()
        self.in_channels = in_channels
        self.out_channels = out_channels
        self.emb_channels = emb_channels
        self.num_heads = 0 if not attention else num_heads if num_heads is not None else out_channels // channels_per_head
        self.dropout = dropout
        self.skip_scale = skip_scale
        self.adaptive_scale = adaptive_scale

        self.norm0 = GroupNorm(num_channels=in_channels, eps=eps)
        self.conv0 = Conv2d(in_channels=in_channels, out_channels=out_channels, kernel=3, up=up, down=down, resample_filter=resample_filter, **init)
        self.affine = Linear(in_features=emb_channels, out_features=out_channels*(2 if adaptive_scale else 1), **init)
        self.norm1 = GroupNorm(num_channels=out_channels, eps=eps)
        self.conv1 = Conv2d(in_channels=out_channels, out_channels=out_channels, kernel=3, **init_zero)

        self.skip = None
        if out_channels != in_channels or up or down:
            kernel = 1 if resample_proj or out_channels!= in_channels else 0
            self.skip = Conv2d(in_channels=in_channels, out_channels=out_channels, kernel=kernel, up=up, down=down, resample_filter=resample_filter, **init)

        if self.num_heads:
            self.norm2 = GroupNorm(num_channels=out_channels, eps=eps)
            self.qkv = Conv2d(in_channels=out_channels, out_channels=out_channels*3, kernel=1, **(init_attn if init_attn is not None else init))
            self.proj = Conv2d(in_channels=out_channels, out_channels=out_channels, kernel=1, **init_zero)

    def forward(self, x, emb):
        orig = x
        x = self.conv0(silu(self.norm0(x)))

        params = self.affine(emb).unsqueeze(2).unsqueeze(3).to(x.dtype)
        if self.adaptive_scale:
            scale, shift = params.chunk(chunks=2, dim=1)
            x = silu(torch.addcmul(shift, self.norm1(x), scale + 1))
        else:
            x = silu(self.norm1(x.add_(params)))

        x = self.conv1(torch.nn.functional.dropout(x, p=self.dropout, training=self.training))
        x = x.add_(self.skip(orig) if self.skip is not None else orig)
        x = x * self.skip_scale

        if self.num_heads:
            q, k, v = self.qkv(self.norm2(x)).reshape(x.shape[0] * self.num_heads, x.shape[1] // self.num_heads, 3, -1).unbind(2)
            w = AttentionOp.apply(q, k)
            a = torch.einsum('nqk,nck->ncq', w, v)
            x = self.proj(a.reshape(*x.shape)).add_(x)
            x = x * self.skip_scale
        return x

#----------------------------------------------------------------------------
# Timestep embedding used in the DDPM++ and ADM architectures.

@persistence.persistent_class
class PositionalEmbedding(torch.nn.Module):
    def __init__(self, num_channels, max_positions=10000, endpoint=False):
        super().__init__()
        self.num_channels = num_channels
        self.max_positions = max_positions
        self.endpoint = endpoint

    def forward(self, x):
        freqs = torch.arange(start=0, end=self.num_channels//2, dtype=torch.float32, device=x.device)
        freqs = freqs / (self.num_channels // 2 - (1 if self.endpoint else 0))
        freqs = (1 / self.max_positions) ** freqs
        x = x.ger(freqs.to(x.dtype))
        x = torch.cat([x.cos(), x.sin()], dim=1)
        return x

#----------------------------------------------------------------------------
# Timestep embedding used in the NCSN++ architecture.

@persistence.persistent_class
class FourierEmbedding(torch.nn.Module):
    def __init__(self, num_channels, scale=16):
        super().__init__()
        self.register_buffer('freqs', torch.randn(num_channels // 2) * scale)

    def forward(self, x):
        x = x.ger((2 * np.pi * self.freqs).to(x.dtype))
        x = torch.cat([x.cos(), x.sin()], dim=1)
        return x

#----------------------------------------------------------------------------
# Reimplementation of the DDPM++ and NCSN++ architectures from the paper
# "Score-Based Generative Modeling through Stochastic Differential
# Equations". Equivalent to the original implementation by Song et al.,
# available at https://github.com/yang-song/score_sde_pytorch

@persistence.persistent_class
class SongUNet(torch.nn.Module):
    def __init__(self,
        img_resolution,                     # Image resolution at input/output.
        in_channels,                        # Number of color channels at input.
        out_channels,                       # Number of color channels at output.
        label_dim           = 0,            # Number of class labels, 0 = unconditional.
        augment_dim         = 0,            # Augmentation label dimensionality, 0 = no augmentation.

        model_channels      = 128,          # Base multiplier for the number of channels.
        channel_mult        = [1,2,2,2],    # Per-resolution multipliers for the number of channels.
        channel_mult_emb    = 4,            # Multiplier for the dimensionality of the embedding vector.
        num_blocks          = 4,            # Number of residual blocks per resolution.
        attn_resolutions    = [16],         # List of resolutions with self-attention.
        dropout             = 0.10,         # Dropout probability of intermediate activations.
        label_dropout       = 0,            # Dropout probability of class labels for classifier-free guidance.

        embedding_type      = 'positional', # Timestep embedding type: 'positional' for DDPM++, 'fourier' for NCSN++.
        channel_mult_noise  = 1,            # Timestep embedding size: 1 for DDPM++, 2 for NCSN++.
        encoder_type        = 'standard',   # Encoder architecture: 'standard' for DDPM++, 'residual' for NCSN++.
        decoder_type        = 'standard',   # Decoder architecture: 'standard' for both DDPM++ and NCSN++.
        resample_filter     = [1,1],        # Resampling filter: [1,1] for DDPM++, [1,3,3,1] for NCSN++.
    ):
        assert embedding_type in ['fourier', 'positional']
        assert encoder_type in ['standard', 'skip', 'residual']
        assert decoder_type in ['standard', 'skip']

        super().__init__()
        self.label_dropout = label_dropout
        emb_channels = model_channels * channel_mult_emb
        noise_channels = model_channels * channel_mult_noise
        init = dict(init_mode='xavier_uniform')
        init_zero = dict(init_mode='xavier_uniform', init_weight=1e-5)
        init_attn = dict(init_mode='xavier_uniform', init_weight=np.sqrt(0.2))
        block_kwargs = dict(
            emb_channels=emb_channels, num_heads=1, dropout=dropout, skip_scale=np.sqrt(0.5), eps=1e-6,
            resample_filter=resample_filter, resample_proj=True, adaptive_scale=False,
            init=init, init_zero=init_zero, init_attn=init_attn,
        )

        # Mapping.
        self.map_noise = PositionalEmbedding(num_channels=noise_channels, endpoint=True) if embedding_type == 'positional' else FourierEmbedding(num_channels=noise_channels)
        self.map_label = Linear(in_features=label_dim, out_features=noise_channels, **init) if label_dim else None
        self.map_augment = Linear(in_features=augment_dim, out_features=noise_channels, bias=False, **init) if augment_dim else None
        self.map_layer0 = Linear(in_features=noise_channels, out_features=emb_channels, **init)
        self.map_layer1 = Linear(in_features=emb_channels, out_features=emb_channels, **init)

        # Encoder.
        self.enc = torch.nn.ModuleDict()
        cout = in_channels
        caux = in_channels
        for level, mult in enumerate(channel_mult):
            res = img_resolution >> level
            if level == 0:
                cin = cout
                cout = model_channels
                self.enc[f'{res}x{res}_conv'] = Conv2d(in_channels=cin, out_channels=cout, kernel=3, **init)
            else:
                self.enc[f'{res}x{res}_down'] = UNetBlock(in_channels=cout, out_channels=cout, down=True, **block_kwargs)
                if encoder_type == 'skip':
                    self.enc[f'{res}x{res}_aux_down'] = Conv2d(in_channels=caux, out_channels=caux, kernel=0, down=True, resample_filter=resample_filter)
                    self.enc[f'{res}x{res}_aux_skip'] = Conv2d(in_channels=caux, out_channels=cout, kernel=1, **init)
                if encoder_type == 'residual':
                    self.enc[f'{res}x{res}_aux_residual'] = Conv2d(in_channels=caux, out_channels=cout, kernel=3, down=True, resample_filter=resample_filter, fused_resample=True, **init)
                    caux = cout
            for idx in range(num_blocks):
                cin = cout
                cout = model_channels * mult
                attn = (res in attn_resolutions)
                self.enc[f'{res}x{res}_block{idx}'] = UNetBlock(in_channels=cin, out_channels=cout, attention=attn, **block_kwargs)
        skips = [block.out_channels for name, block in self.enc.items() if 'aux' not in name]

        # Decoder.
        self.dec = torch.nn.ModuleDict()
        for level, mult in reversed(list(enumerate(channel_mult))):
            res = img_resolution >> level
            if level == len(channel_mult) - 1:
                self.dec[f'{res}x{res}_in0'] = UNetBlock(in_channels=cout, out_channels=cout, attention=True, **block_kwargs)
                self.dec[f'{res}x{res}_in1'] = UNetBlock(in_channels=cout, out_channels=cout, **block_kwargs)
            else:
                self.dec[f'{res}x{res}_up'] = UNetBlock(in_channels=cout, out_channels=cout, up=True, **block_kwargs)
            for idx in range(num_blocks + 1):
                cin = cout + skips.pop()
                cout = model_channels * mult
                attn = (idx == num_blocks and res in attn_resolutions)
                self.dec[f'{res}x{res}_block{idx}'] = UNetBlock(in_channels=cin, out_channels=cout, attention=attn, **block_kwargs)
            if decoder_type == 'skip' or level == 0:
                if decoder_type == 'skip' and level < len(channel_mult) - 1:
                    self.dec[f'{res}x{res}_aux_up'] = Conv2d(in_channels=out_channels, out_channels=out_channels, kernel=0, up=True, resample_filter=resample_filter)
                self.dec[f'{res}x{res}_aux_norm'] = GroupNorm(num_channels=cout, eps=1e-6)
                self.dec[f'{res}x{res}_aux_conv'] = Conv2d(in_channels=cout, out_channels=out_channels, kernel=3, **init_zero)

    def forward(self, x, noise_labels, class_labels, augment_labels=None):
        # Mapping.
        emb = self.map_noise(noise_labels)
        emb = emb.reshape(emb.shape[0], 2, -1).flip(1).reshape(*emb.shape) # swap sin/cos
        if self.map_label is not None:
            tmp = class_labels
            if self.training and self.label_dropout:
                tmp = tmp * (torch.rand([x.shape[0], 1], device=x.device) >= self.label_dropout).to(tmp.dtype)
            emb = emb + self.map_label(tmp * np.sqrt(self.map_label.in_features))
        if self.map_augment is not None and augment_labels is not None:
            emb = emb + self.map_augment(augment_labels)
        emb = silu(self.map_layer0(emb))
        emb = silu(self.map_layer1(emb))

        # Encoder.
        skips = []
        aux = x
        for name, block in self.enc.items():
            if 'aux_down' in name:
                aux = block(aux)
            elif 'aux_skip' in name:
                x = skips[-1] = x + block(aux)
            elif 'aux_residual' in name:
                x = skips[-1] = aux = (x + block(aux)) / np.sqrt(2)
            else:
                x = block(x, emb) if isinstance(block, UNetBlock) else block(x)
                skips.append(x)

        # Decoder.
        aux = None
        tmp = None
        for name, block in self.dec.items():
            if 'aux_up' in name:
                aux = block(aux)
            elif 'aux_norm' in name:
                tmp = block(x)
            elif 'aux_conv' in name:
                tmp = block(silu(tmp))
                aux = tmp if aux is None else tmp + aux
            else:
                if x.shape[1] != block.in_channels:
                    x = torch.cat([x, skips.pop()], dim=1)
                x = block(x, emb)
        return aux

#----------------------------------------------------------------------------
# Reimplementation of the ADM architecture from the paper
# "Diffusion Models Beat GANS on Image Synthesis". Equivalent to the
# original implementation by Dhariwal and Nichol, available at
# https://github.com/openai/guided-diffusion

@persistence.persistent_class
class DhariwalUNet(torch.nn.Module):
    def __init__(self,
        img_resolution,                     # Image resolution at input/output.
        in_channels,                        # Number of color channels at input.
        out_channels,                       # Number of color channels at output.
        label_dim           = 0,            # Number of class labels, 0 = unconditional.
        augment_dim         = 0,            # Augmentation label dimensionality, 0 = no augmentation.

        model_channels      = 192,          # Base multiplier for the number of channels.
        channel_mult        = [1,2,3,4],    # Per-resolution multipliers for the number of channels.
        channel_mult_emb    = 4,            # Multiplier for the dimensionality of the embedding vector.
        num_blocks          = 3,            # Number of residual blocks per resolution.
        attn_resolutions    = [32,16,8],    # List of resolutions with self-attention.
        dropout             = 0.10,         # List of resolutions with self-attention.
        label_dropout       = 0,            # Dropout probability of class labels for classifier-free guidance.
    ):
        super().__init__()
        self.label_dropout = label_dropout
        emb_channels = model_channels * channel_mult_emb
        init = dict(init_mode='kaiming_uniform', init_weight=np.sqrt(1/3), init_bias=np.sqrt(1/3))
        init_zero = dict(init_mode='kaiming_uniform', init_weight=0, init_bias=0)
        block_kwargs = dict(emb_channels=emb_channels, channels_per_head=64, dropout=dropout, init=init, init_zero=init_zero)

        # Mapping.
        self.map_noise = PositionalEmbedding(num_channels=model_channels)
        self.map_augment = Linear(in_features=augment_dim, out_features=model_channels, bias=False, **init_zero) if augment_dim else None
        self.map_layer0 = Linear(in_features=model_channels, out_features=emb_channels, **init)
        self.map_layer1 = Linear(in_features=emb_channels, out_features=emb_channels, **init)
        self.map_label = Linear(in_features=label_dim, out_features=emb_channels, bias=False, init_mode='kaiming_normal', init_weight=np.sqrt(label_dim)) if label_dim else None

        # Encoder.
        self.enc = torch.nn.ModuleDict()
        cout = in_channels
        for level, mult in enumerate(channel_mult):
            res = img_resolution >> level
            if level == 0:
                cin = cout
                cout = model_channels * mult
                self.enc[f'{res}x{res}_conv'] = Conv2d(in_channels=cin, out_channels=cout, kernel=3, **init)
            else:
                self.enc[f'{res}x{res}_down'] = UNetBlock(in_channels=cout, out_channels=cout, down=True, **block_kwargs)
            for idx in range(num_blocks):
                cin = cout
                cout = model_channels * mult
                self.enc[f'{res}x{res}_block{idx}'] = UNetBlock(in_channels=cin, out_channels=cout, attention=(res in attn_resolutions), **block_kwargs)
        skips = [block.out_channels for block in self.enc.values()]

        # Decoder.
        self.dec = torch.nn.ModuleDict()
        for level, mult in reversed(list(enumerate(channel_mult))):
            res = img_resolution >> level
            if level == len(channel_mult) - 1:
                self.dec[f'{res}x{res}_in0'] = UNetBlock(in_channels=cout, out_channels=cout, attention=True, **block_kwargs)
                self.dec[f'{res}x{res}_in1'] = UNetBlock(in_channels=cout, out_channels=cout, **block_kwargs)
            else:
                self.dec[f'{res}x{res}_up'] = UNetBlock(in_channels=cout, out_channels=cout, up=True, **block_kwargs)
            for idx in range(num_blocks + 1):
                cin = cout + skips.pop()
                cout = model_channels * mult
                self.dec[f'{res}x{res}_block{idx}'] = UNetBlock(in_channels=cin, out_channels=cout, attention=(res in attn_resolutions), **block_kwargs)
        self.out_norm = GroupNorm(num_channels=cout)
        self.out_conv = Conv2d(in_channels=cout, out_channels=out_channels, kernel=3, **init_zero)

    def forward(self, x, noise_labels, class_labels, augment_labels=None):
        # Mapping.
        emb = self.map_noise(noise_labels)
        if self.map_augment is not None and augment_labels is not None:
            emb = emb + self.map_augment(augment_labels)
        emb = silu(self.map_layer0(emb))
        emb = self.map_layer1(emb)
        if self.map_label is not None:
            tmp = class_labels
            if self.training and self.label_dropout:
                tmp = tmp * (torch.rand([x.shape[0], 1], device=x.device) >= self.label_dropout).to(tmp.dtype)
            emb = emb + self.map_label(tmp)
        emb = silu(emb)

        # Encoder.
        skips = []
        for block in self.enc.values():
            x = block(x, emb) if isinstance(block, UNetBlock) else block(x)
            skips.append(x)

        # Decoder.
        for block in self.dec.values():
            if x.shape[1] != block.in_channels:
                x = torch.cat([x, skips.pop()], dim=1)
            x = block(x, emb)
        x = self.out_conv(silu(self.out_norm(x)))
        return x

##########################################
# Preconditionings from EDM paper
# This is legacy code and will be removed! 

#----------------------------------------------------------------------------
# Preconditioning corresponding to the variance preserving (VP) formulation
# from the paper "Score-Based Generative Modeling through Stochastic
# Differential Equations".

@persistence.persistent_class
class VPPrecond(torch.nn.Module):
    def __init__(self,
        img_resolution,                 # Image resolution.
        img_channels,                   # Number of color channels.
        label_dim       = 0,            # Number of class labels, 0 = unconditional.
        use_fp16        = False,        # Execute the underlying model at FP16 precision?
        beta_d          = 19.9,         # Extent of the noise level schedule.
        beta_min        = 0.1,          # Initial slope of the noise level schedule.
        M               = 1000,         # Original number of timesteps in the DDPM formulation.
        epsilon_t       = 1e-5,         # Minimum t-value used during training.
        model_type      = 'SongUNet',   # Class name of the underlying model.
        **model_kwargs,                 # Keyword arguments for the underlying model.
    ):
        super().__init__()
        self.img_resolution = img_resolution
        self.img_channels = img_channels
        self.label_dim = label_dim
        self.use_fp16 = use_fp16
        self.beta_d = beta_d
        self.beta_min = beta_min
        self.M = M
        self.epsilon_t = epsilon_t
        self.sigma_min = float(self.sigma(epsilon_t))
        self.sigma_max = float(self.sigma(1))
        self.model = globals()[model_type](img_resolution=img_resolution, in_channels=img_channels, out_channels=img_channels, label_dim=label_dim, **model_kwargs)

    def forward(self, x, sigma, class_labels=None, force_fp32=False, **model_kwargs):
        x = x.to(torch.float32)
        sigma = sigma.to(torch.float32).reshape(-1, 1, 1, 1)
        class_labels = None if self.label_dim == 0 else torch.zeros([1, self.label_dim], device=x.device) if class_labels is None else class_labels.to(torch.float32).reshape(-1, self.label_dim)
        dtype = torch.float16 if (self.use_fp16 and not force_fp32 and x.device.type == 'cuda') else torch.float32

        c_skip = 1
        c_out = -sigma
        c_in = 1 / (sigma ** 2 + 1).sqrt()
        c_noise = (self.M - 1) * self.sigma_inv(sigma)

        F_x = self.model((c_in * x).to(dtype), c_noise.flatten(), class_labels=class_labels, **model_kwargs)
        assert F_x.dtype == dtype
        D_x = c_skip * x + c_out * F_x.to(torch.float32)
        return D_x

    def sigma(self, t):
        t = torch.as_tensor(t)
        return ((0.5 * self.beta_d * (t ** 2) + self.beta_min * t).exp() - 1).sqrt()

    def sigma_inv(self, sigma):
        sigma = torch.as_tensor(sigma)
        return ((self.beta_min ** 2 + 2 * self.beta_d * (1 + sigma ** 2).log()).sqrt() - self.beta_min) / self.beta_d

    def round_sigma(self, sigma):
        return torch.as_tensor(sigma)

#----------------------------------------------------------------------------
# Preconditioning corresponding to the variance exploding (VE) formulation
# from the paper "Score-Based Generative Modeling through Stochastic
# Differential Equations".

@persistence.persistent_class
class VEPrecond(torch.nn.Module):
    def __init__(self,
        img_resolution,                 # Image resolution.
        img_channels,                   # Number of color channels.
        label_dim       = 0,            # Number of class labels, 0 = unconditional.
        use_fp16        = False,        # Execute the underlying model at FP16 precision?
        sigma_min       = 0.02,         # Minimum supported noise level.
        sigma_max       = 100,          # Maximum supported noise level.
        model_type      = 'SongUNet',   # Class name of the underlying model.
        **model_kwargs,                 # Keyword arguments for the underlying model.
    ):
        super().__init__()
        self.img_resolution = img_resolution
        self.img_channels = img_channels
        self.label_dim = label_dim
        self.use_fp16 = use_fp16
        self.sigma_min = sigma_min
        self.sigma_max = sigma_max
        self.model = globals()[model_type](img_resolution=img_resolution, in_channels=img_channels, out_channels=img_channels, label_dim=label_dim, **model_kwargs)

    def forward(self, x, sigma, class_labels=None, force_fp32=False, **model_kwargs):
        x = x.to(torch.float32)
        sigma = sigma.to(torch.float32).reshape(-1, 1, 1, 1)
        class_labels = None if self.label_dim == 0 else torch.zeros([1, self.label_dim], device=x.device) if class_labels is None else class_labels.to(torch.float32).reshape(-1, self.label_dim)
        dtype = torch.float16 if (self.use_fp16 and not force_fp32 and x.device.type == 'cuda') else torch.float32

        c_skip = 1
        c_out = sigma
        c_in = 1
        c_noise = (0.5 * sigma).log()

        F_x = self.model((c_in * x).to(dtype), c_noise.flatten(), class_labels=class_labels, **model_kwargs)
        assert F_x.dtype == dtype
        D_x = c_skip * x + c_out * F_x.to(torch.float32)
        return D_x

    def round_sigma(self, sigma):
        return torch.as_tensor(sigma)

#----------------------------------------------------------------------------
# Preconditioning corresponding to improved DDPM (iDDPM) formulation from
# the paper "Improved Denoising Diffusion Probabilistic Models".

@persistence.persistent_class
class iDDPMPrecond(torch.nn.Module):
    def __init__(self,
        img_resolution,                     # Image resolution.
        img_channels,                       # Number of color channels.
        label_dim       = 0,                # Number of class labels, 0 = unconditional.
        use_fp16        = False,            # Execute the underlying model at FP16 precision?
        C_1             = 0.001,            # Timestep adjustment at low noise levels.
        C_2             = 0.008,            # Timestep adjustment at high noise levels.
        M               = 1000,             # Original number of timesteps in the DDPM formulation.
        model_type      = 'DhariwalUNet',   # Class name of the underlying model.
        **model_kwargs,                     # Keyword arguments for the underlying model.
    ):
        super().__init__()
        self.img_resolution = img_resolution
        self.img_channels = img_channels
        self.label_dim = label_dim
        self.use_fp16 = use_fp16
        self.C_1 = C_1
        self.C_2 = C_2
        self.M = M
        self.model = globals()[model_type](img_resolution=img_resolution, in_channels=img_channels, out_channels=img_channels*2, label_dim=label_dim, **model_kwargs)

        u = torch.zeros(M + 1)
        for j in range(M, 0, -1): # M, ..., 1
            u[j - 1] = ((u[j] ** 2 + 1) / (self.alpha_bar(j - 1) / self.alpha_bar(j)).clip(min=C_1) - 1).sqrt()
        self.register_buffer('u', u)
        self.sigma_min = float(u[M - 1])
        self.sigma_max = float(u[0])

    def forward(self, x, sigma, class_labels=None, force_fp32=False, **model_kwargs):
        x = x.to(torch.float32)
        sigma = sigma.to(torch.float32).reshape(-1, 1, 1, 1)
        class_labels = None if self.label_dim == 0 else torch.zeros([1, self.label_dim], device=x.device) if class_labels is None else class_labels.to(torch.float32).reshape(-1, self.label_dim)
        dtype = torch.float16 if (self.use_fp16 and not force_fp32 and x.device.type == 'cuda') else torch.float32

        c_skip = 1
        c_out = -sigma
        c_in = 1 / (sigma ** 2 + 1).sqrt()
        c_noise = self.M - 1 - self.round_sigma(sigma, return_index=True).to(torch.float32)

        F_x = self.model((c_in * x).to(dtype), c_noise.flatten(), class_labels=class_labels, **model_kwargs)
        assert F_x.dtype == dtype
        D_x = c_skip * x + c_out * F_x[:, :self.img_channels].to(torch.float32)
        return D_x

    def alpha_bar(self, j):
        j = torch.as_tensor(j)
        return (0.5 * np.pi * j / self.M / (self.C_2 + 1)).sin() ** 2

    def round_sigma(self, sigma, return_index=False):
        sigma = torch.as_tensor(sigma)
        index = torch.cdist(sigma.to(self.u.device).to(torch.float32).reshape(1, -1, 1), self.u.reshape(1, -1, 1)).argmin(2)
        result = index if return_index else self.u[index.flatten()].to(sigma.dtype)
        return result.reshape(sigma.shape).to(sigma.device)


#----------------------------------------------------------------------------
# Preconditioning corresponding to EDM (proposed) formulation from
# the paper "Improved Denoising Diffusion Probabilistic Models".
@persistence.persistent_class
class EDMPrecond(torch.nn.Module):
    def __init__(self,
        img_resolution,                     # Image resolution.
        img_channels,                       # Number of color channels.
        label_dim       = 0,                # Number of class labels, 0 = unconditional.
        use_fp16        = False,            # Execute the underlying model at FP16 precision?
        sigma_min       = 0,                # Minimum supported noise level.
        sigma_max       = float('inf'),     # Maximum supported noise level.
        sigma_data      = 0.5,              # Expected standard deviation of the training data.
        model_type      = 'DhariwalUNet',   # Class name of the underlying model.
        **model_kwargs,                     # Keyword arguments for the underlying model.
    ):
        super().__init__()
        self.img_resolution = img_resolution
        self.img_channels = img_channels
        self.label_dim = label_dim
        self.use_fp16 = use_fp16
        self.sigma_min = sigma_min
        self.sigma_max = sigma_max
        self.sigma_data = sigma_data
        self.model = globals()[model_type](img_resolution=img_resolution, in_channels=img_channels, out_channels=img_channels, label_dim=label_dim, **model_kwargs)

    def forward(self, x, sigma, class_labels=None, force_fp32=False, **model_kwargs):
        x = x.to(torch.float32)
        sigma = sigma.to(torch.float32).reshape(-1, 1, 1, 1)
        class_labels = None if self.label_dim == 0 else torch.zeros([1, self.label_dim], device=x.device) if class_labels is None else class_labels.to(torch.float32).reshape(-1, self.label_dim)
        dtype = torch.float16 if (self.use_fp16 and not force_fp32 and x.device.type == 'cuda') else torch.float32

        c_skip = self.sigma_data ** 2 / (sigma ** 2 + self.sigma_data ** 2)
        c_out = sigma * self.sigma_data / (sigma ** 2 + self.sigma_data ** 2).sqrt()
        c_in = 1 / (self.sigma_data ** 2 + sigma ** 2).sqrt()
        c_noise = sigma.log() / 4

        F_x = self.model((c_in * x).to(dtype), c_noise.flatten(), class_labels=class_labels, **model_kwargs)
        assert F_x.dtype == dtype
        D_x = c_skip * x + c_out * F_x.to(torch.float32)
        return D_x

    def round_sigma(self, sigma):
        return torch.as_tensor(sigma)

##################################################################################
@persistence.persistent_class
class ConvVNetWrapper(nn.Module):
    def __init__(self, base_model, img_ch, img_size, data_dim,
                 dim_cov_dynamic, dim_cov_static, use_t_dyn=True):
        super().__init__()
        self.base_model = base_model
        self.img_ch    = img_ch
        self.img_size  = img_size
        self.pixels    = img_ch * img_size * img_size
        self.dim_cov_dynamic = dim_cov_dynamic or 0
        self.dim_cov_static  = dim_cov_static  or 0
        self.use_t_dyn = bool(use_t_dyn)

        self.is_image_data = (data_dim == self.pixels)
        self.static_elems = 0
        self.lag_ch = 0
        self.lag_tempmix = None

        if self.is_image_data:
            self.static_elems = int(self.dim_cov_static) % self.pixels
            lag_flat_size = int(self.dim_cov_static) - self.static_elems
            if lag_flat_size > 0:
                assert lag_flat_size % self.pixels == 0
                self.lag_ch = lag_flat_size // (self.img_size * self.img_size)

            if self.lag_ch > 0:
                C = self.img_ch
                assert self.lag_ch % C == 0
                self.L_lag = self.lag_ch // C
                self.lag_tempmix = nn.Conv1d(
                    in_channels=C, out_channels=C,
                    kernel_size=3, padding=1, groups=C, bias=True
                )
                with torch.no_grad():
                    self.lag_tempmix.weight.zero_()
                    self.lag_tempmix.bias.zero_()
                    self.lag_tempmix.weight[:, 0, 1] = 1.0

            self.vec_cov_dim = self.dim_cov_dynamic + self.static_elems + (1 if self.use_t_dyn else 0)
            self.cov_norm = nn.LayerNorm(self.vec_cov_dim, elementwise_affine=False) if self.vec_cov_dim > 0 else None
            self.cov_film = nn.Linear(self.vec_cov_dim, self.img_ch * 2) if self.vec_cov_dim > 0 else None
            if self.cov_film is not None:
                with torch.no_grad():
                    self.cov_film.weight.zero_()
                    self.cov_film.bias.zero_()

            in_ch_total = self.img_ch + self.lag_ch
            self.in_proj = nn.Conv2d(in_ch_total, self.img_ch, kernel_size=1, bias=True)
        else:
            self.vec_cov_dim = self.dim_cov_dynamic + self.dim_cov_static + (1 if self.use_t_dyn else 0)
            self.cov_norm = nn.LayerNorm(self.vec_cov_dim, elementwise_affine=False) if self.vec_cov_dim > 0 else None
            self.cov_film = None
            self.in_proj = None
            self.linear_projection = nn.Linear(data_dim, data_dim)

    def _build_cov_vec(self, cov_dynamic, cov_static, t_dyn):
        parts = []
        if self.dim_cov_dynamic > 0 and cov_dynamic is not None:
            parts.append(cov_dynamic)
        if self.dim_cov_static > 0 and cov_static is not None:
            if self.is_image_data:
                if self.static_elems > 0:
                    parts.append(cov_static[..., :self.static_elems])
            else:
                parts.append(cov_static)
        if self.use_t_dyn and (t_dyn is not None):
            parts.append(t_dyn.view(-1, 1))
        if not parts:
            return None
        c = torch.cat(parts, dim=-1)
        if self.cov_norm is not None:
            c = self.cov_norm(c)
        return c

    def forward(self, xt_tau, cov_dynamic, cov_static, t, t_dyn=None):
        B = xt_tau.shape[0]
        c_vec = self._build_cov_vec(cov_dynamic, cov_static, t_dyn)

        if self.is_image_data:
            H = W = self.img_size
            xt_img = xt_tau.view(B, self.img_ch, H, W)

            lag_maps = None
            if self.dim_cov_static > 0 and self.lag_ch > 0 and cov_static is not None:
                lag_flat = cov_static[..., self.static_elems:]
                lag_maps = lag_flat.view(B, self.lag_ch, H, W)
                if self.lag_tempmix is not None:
                    C = self.img_ch
                    L = self.L_lag
                    x = lag_maps.view(B, C, L, H, W).permute(0, 3, 4, 1, 2).reshape(B * H * W, C, L)
                    x = self.lag_tempmix(x)
                    lag_maps = x.view(B, H, W, C, L).permute(0, 3, 4, 1, 2).reshape(B, self.lag_ch, H, W)

            if lag_maps is None and self.lag_ch > 0:
                lag_maps = torch.zeros(B, self.lag_ch, H, W, device=xt_tau.device, dtype=xt_tau.dtype)

            pieces = [xt_img]
            if lag_maps is not None:
                pieces.append(lag_maps)
            x_cat = torch.cat(pieces, dim=1)
            img_feat = self.in_proj(x_cat)

            if self.cov_film is not None and c_vec is not None:
                style = self.cov_film(c_vec).view(B, 2 * self.img_ch, 1, 1)
                scale, shift = style.chunk(2, dim=1)
                img_feat = img_feat * (1.0 + scale) + shift

            try:
                y = self.base_model(img_feat, t, c_vec)
            except TypeError:
                y = self.base_model(img_feat, t)

            if y.dim() == 2 and y.shape[1] == self.pixels:
                y = y.view(B, self.img_ch, H, W)
            elif y.dim() != 4:
                raise RuntimeError(f"Expected (B,C,H,W) or flat (B,{self.pixels}), got {tuple(y.shape)}")

            y = _center_crop_or_resize(y, self.img_size)
            return y.reshape(B, -1)

        projected = self.linear_projection(xt_tau)
        try:
            return self.base_model(projected, t, c_vec)
        except TypeError:
            if c_vec is not None:
                inp = torch.cat([projected, c_vec], dim=-1)
            else:
                inp = projected
            return self.base_model(inp, t)


#-------------------------------------------------------------------------------------
#Def network class for VFM

@persistence.persistent_class
class VFMToyNet(torch.nn.Module):
    """
    Wrapper for calling u (flow time) and v (dynamics time) nets.
    Supports either MLP or ToyConvUNet architectures.
    Now supports optional covariates for the dynamics network.
    """
    def _apply_nd_grad_mask(self, z):
        if (not self.training) or (not hasattr(self, "encoder")):
            return z
        enc = self.encoder
        d = z.shape[1]
        k_max = int(min(getattr(enc, "k_max", d), d))
        last_K = int(float(getattr(enc, "last_K", 0.0))) if hasattr(enc, "last_K") else 0
        K = k_max if last_K <= 0 else min(last_K, k_max)
        if K <= 0 or K >= d:
            return z
        mask = z.new_zeros(d)
        mask[:K] = 1.0
        return z * mask + z.detach() * (1.0 - mask)
    
    def __init__(self, 
                 channels = [32, 64, 128, 256],
                 conv_embed_dim = 256,
                 channels_cmp=None,            # list[int] for compression/flow UNet
                 channels_dyn=None,            # list[int] for dynamics UNet
                 channels_enc=None,            # list[int] for encoder CNN
                 conv_embed_dim_cmp=None,      # int for compression/flow UNet
                 conv_embed_dim_dyn=None,
                 data_dim=2,
                 k_max = 10,
                 dyn_model_type = "ToyMLP",
                 flow_model_type = "ToyMLP",
                 encoder_type = "Latent_MLP_VAE",
                 depth_encoder = 2,
                 width_encoder = 10,
                 depth_mlp_cmp = 2,
                 width_mlp_cmp = 64,
                 depth_mlp_dyn = 2,
                 width_mlp_dyn = 64,
                 img_size = 28,
                 in_ch=1,
                 d_min=1e-15,
                 mu_radial_R = 100.0,
                 save_dir='',
                 dim_cov_dynamic=0,  # dimension for dynamic covariates
                 dim_cov_static=0, # dimension for static covariates
                 use_t_dyn=True,
                 flow_detach=False,
                 ):
        super().__init__()
        self.img_size = img_size
        self.in_ch = in_ch
        self.mu_radial_R = mu_radial_R

        self.data_dim = data_dim
        self.k_max = k_max
        self.save_dir = save_dir
        self.dim_cov_dynamic = dim_cov_dynamic
        self.dim_cov_static = dim_cov_static
        self.use_t_dyn = bool(use_t_dyn)
        self.flow_detach = bool(flow_detach)      
        
        ch_cmp = channels if channels_cmp is None else channels_cmp
        ch_dyn = channels if channels_dyn is None else channels_dyn
        ch_enc = channels if channels_enc is None else channels_enc
        emb_cmp = conv_embed_dim if conv_embed_dim_cmp is None else conv_embed_dim_cmp
        emb_dyn = conv_embed_dim if conv_embed_dim_dyn is None else conv_embed_dim_dyn
        
        # Create u net (flow net) - input dimension stays the same
        if flow_model_type in ["ToyConvUNet", "Adapted_ToyConvUNet"]:
            self.unet_model = globals()[flow_model_type](channels=ch_cmp, embed_dim=emb_cmp, 
                                                         img_size=img_size, img_ch=in_ch,
                                                         input_size=data_dim)
        else:
            self.unet_model = globals()[flow_model_type](dim=data_dim, time_varying=True, 
                                                         n_hidden=depth_mlp_cmp, w=width_mlp_cmp)
        
        # Calculate input dimension for dynamics net
        vnet_input_dim = data_dim  # xt_tau
        if dim_cov_dynamic > 0:
            vnet_input_dim += dim_cov_dynamic  # dynamic covariates
        if dim_cov_static > 0:
            vnet_input_dim += dim_cov_static  # static covariates
        
        # Create vnet (dynamics net) with correct input dimension
        if dyn_model_type in ["ToyConvUNet", "Adapted_ToyConvUNet"]:
            pixels = in_ch * img_size * img_size
            static_elems = int(dim_cov_static or 0) % pixels
            cov_dim_for_vnet = (dim_cov_dynamic or 0) + static_elems + (1 if self.use_t_dyn else 0)
            base_vnet = globals()[dyn_model_type](
                channels=ch_dyn, embed_dim=emb_dyn,
                img_size=img_size, img_ch=in_ch,
                input_size=data_dim,
                cov_dim=cov_dim_for_vnet
            )
            self.vnet_model = ConvVNetWrapper(
                base_model=base_vnet,
                img_ch=in_ch,
                img_size=img_size,
                data_dim=data_dim,
                dim_cov_dynamic=dim_cov_dynamic,
                dim_cov_static=dim_cov_static,
                use_t_dyn=self.use_t_dyn
            )
        else:
            # For MLP architectures
            self.vnet_model = globals()[dyn_model_type](
                dim=vnet_input_dim, out_dim=data_dim,
                time_varying=(2 if self.use_t_dyn else True),
                n_hidden=depth_mlp_dyn, w=width_mlp_dyn
            )
        
        # Create encoder net
        if encoder_type == 'Latent_LargeCNN_VAE':
            self.encoder = Latent_LargeCNN_VAE(
                channels=ch_enc,
                img_size=img_size,
                img_ch=in_ch,
                k_max=k_max,
                d_min=d_min,
            )
        elif encoder_type == 'Adapted_Latent_LargeCNN_VAE':
            self.encoder = Adapted_Latent_LargeCNN_VAE(
                channels=ch_enc,
                img_size=img_size,
                img_ch=in_ch,
                input_size=data_dim,
                k_max=k_max,
                d_min=d_min,
            )
        elif encoder_type == "Latent_MLP_VAE":
            self.encoder = Latent_MLP_VAE(
                input_size=data_dim,
                output_size=data_dim,
                k_max=k_max,
                num_hidden=depth_encoder,
                hidden_size=width_encoder,
                d_min=d_min,
            )
        else:
            self.encoder = globals()[encoder_type](
                img_resolution=img_size,
                img_ch=in_ch,
                k_max=k_max,
                d_min=d_min,
            )

        if hasattr(self, "mu_radial_R") and hasattr(self.encoder, "mu_radial_R"):
            self.encoder.mu_radial_R = self.mu_radial_R
            
    def forward(self, x0_1, xdt_1, taus, ts, dt, tau_flowmatcher, t_flowmatcher, 
                cov_dynamic_0=None, cov_dynamic_dt=None, cov_static=None):

        x0_0_raw, x0_0_mask = self.encoder.rsample(x0_1)
        K_shared = int(float(getattr(self.encoder, "last_K", 0.0))) or None
        xdt_0_raw, xdt_0_mask = self.encoder.rsample(xdt_1, K=K_shared)

        x0_0 = self._apply_nd_grad_mask(x0_0_raw)
        xdt_0 = self._apply_nd_grad_mask(xdt_0_raw)

        if self.flow_detach:
            x0_0_flow = x0_0.detach()
            xdt_0_flow = xdt_0.detach()
        else:
            x0_0_flow = x0_0
            xdt_0_flow = xdt_0

        _, x0_tau, u0_tau = tau_flowmatcher.sample_location_and_conditional_flow(x0_0_flow, x0_1, t=taus)
        _, xdt_tau, _ = tau_flowmatcher.sample_location_and_conditional_flow(xdt_0_flow, xdt_1, t=taus)

        t_norm = (ts / dt).clamp(0, 1)
        _, xt_tau, ut_tau = t_flowmatcher.sample_location_and_conditional_flow(x0_tau,
                                                                               xdt_tau, t=t_norm)
        
        cov_dynamic_tau = None
        cov_static_tau = None

        if (cov_dynamic_0 is not None) and (cov_dynamic_dt is not None) and (self.dim_cov_dynamic > 0):
            t_interp = t_norm.unsqueeze(-1)
            cov_dynamic_tau = (1.0 - t_interp) * cov_dynamic_0 + t_interp * cov_dynamic_dt

        cov_static_tau = cov_static
        if (cov_static_tau is not None) and (self.dim_cov_static or 0) > 0:
            with torch.no_grad():
                base = self.in_ch * self.img_size * self.img_size
                is_image = (self.data_dim == base)
                P = base if is_image else int(self.data_dim)

                static_elems = int(self.dim_cov_static) % P
                lag_elems = int(self.dim_cov_static) - static_elems

                if lag_elems > 0 and cov_static_tau.shape[1] >= static_elems + lag_elems:
                    Bfull = cov_static_tau.shape[0]
                    lag_flat = cov_static_tau[:, static_elems: static_elems + lag_elems]

                    if is_image:
                        H = W = self.img_size
                        C = self.in_ch
                        assert lag_elems % (C * H * W) == 0, "lag_elems must be multiple of C*H*W."
                        L = lag_elems // (C * H * W)

                        lag_maps = lag_flat.view(Bfull, C * L, H, W)
                        X1_img = lag_maps.view(Bfull, L, C, H, W).permute(0, 2, 1, 3, 4).contiguous().view(
                            Bfull * L, C, H, W
                        )
                        X1_vec = X1_img.view(Bfull * L, -1)

                        enc_name = self.encoder.__class__.__name__
                        enc_is_cnn = enc_name in ('Latent_LargeCNN_VAE', 'Latent_CNN_VAE', 'Adapted_Latent_LargeCNN_VAE')
                        X0_vec, _ = self.encoder.rsample(X1_img if enc_is_cnn else X1_vec, update_stats=False)
                        if self.flow_detach:
                            X0_vec = X0_vec.detach()
                        else:
                            X0_vec = self._apply_nd_grad_mask(X0_vec)

                        taus_rep = taus.repeat_interleave(L, dim=0)  # (B*L,)
                        _, Xtau_vec, _ = tau_flowmatcher.sample_location_and_conditional_flow(
                            X0_vec, X1_vec, t=taus_rep
                        )

                        Xtau = Xtau_vec.view(Bfull, L, C, H, W).permute(0, 2, 1, 3, 4).contiguous()
                        lag_tau_flat = Xtau.view(Bfull, lag_elems)
                    else:
                        D = int(self.data_dim)
                        assert lag_elems % D == 0, "lag_elems must be multiple of D."
                        L = lag_elems // D
                        X1_vec = lag_flat.view(Bfull * L, D)

                        X0_vec, _ = self.encoder.rsample(X1_vec, update_stats=False)
                        if self.flow_detach:
                            X0_vec = X0_vec.detach()
                        else:
                            X0_vec = self._apply_nd_grad_mask(X0_vec)

                        taus_rep = taus.repeat_interleave(L, dim=0)
                        _, Xtau_vec, _ = tau_flowmatcher.sample_location_and_conditional_flow(
                            X0_vec, X1_vec, t=taus_rep
                        )
                        lag_tau_flat = Xtau_vec.view(Bfull, lag_elems)

                    if static_elems > 0:
                        cov_static_tau = torch.cat(
                            [cov_static_tau[:, :static_elems], lag_tau_flat], dim=1
                        )
                    else:
                        cov_static_tau = lag_tau_flat

        # ------ u_net (compression)
        u = self.unet_model(x0_tau, taus)
        if u.dim() == 4:
            u = _center_crop_or_resize(u, self.img_size).reshape(u.size(0), -1)

        # ------- v_net (dynamic)
        if hasattr(self.vnet_model, '__class__') and self.vnet_model.__class__.__name__ == 'ConvVNetWrapper':
            if self.use_t_dyn:
                v = self.vnet_model(xt_tau, cov_dynamic_tau, cov_static_tau, taus, t_norm)
            else:
                v = self.vnet_model(xt_tau, cov_dynamic_tau, cov_static_tau, taus)
        else:
            inputs = [xt_tau]
            if self.dim_cov_dynamic > 0 and cov_dynamic_tau is not None:
                inputs.append(cov_dynamic_tau)
            if self.dim_cov_static > 0 and cov_static_tau is not None:
                inputs.append(cov_static_tau)
            v_input = torch.cat(inputs, dim=-1)
            if self.use_t_dyn:
                v = self.vnet_model(v_input, taus, t_norm)
            else:
                v = self.vnet_model(v_input, taus)
        
        if v.dim() == 4:
            v = _center_crop_or_resize(v, self.img_size).reshape(v.size(0), -1)
        
        return u0_tau, ut_tau, u, v, x0_0, xdt_0, x0_0_mask, xdt_0_mask
            
            

    
    
```
