## Supplementary material for "Dynamic Compression Flows for Neuroscience Data": Python code: toy_training_loop_vfm_v7.py

```
"""
Actually does training for toy VFM models.

"""

import os
import re
import time
import copy
import json
import pickle
import numpy as np
import torch
import dnnlib
from torch_utils import distributed as dist
from torch_utils import training_stats
from torch_utils import misc
from torch.utils.tensorboard import SummaryWriter
from torch_cfm import conditional_flow_matching as cfm
import torch.serialization
import inspect
from torch_utils import persistence
import torch.nn as nn
import gc
import math

# ------------------------------- helper ----------------------------------------
def _get_enc_mod(model):
    base = model.module if hasattr(model, 'module') else model
    return getattr(base, 'encoder', base)

def _copy_pop_stats(src_enc, dst_enc):
    src_bufs = dict(src_enc.named_buffers())
    dst_bufs = dict(dst_enc.named_buffers())
    for n, s in src_bufs.items():
        if any(k in n for k in ('pop','ema','data_mean','running')) and n in dst_bufs:
            d = dst_bufs[n]
            if torch.is_tensor(s) and torch.is_tensor(d) and s.shape == d.shape:
                d.data.copy_(s.data)

# --------------------------- training loop ----------------------------------------
def training_loop(
    run_dir             = '.',      # Output directory.,
    seed                = 0,        # Global random seed.,
    device              = torch.device('cuda'),
    cudnn_benchmark     = True,     # Enable torch.backends.cudnn.benchmark?,
    
    kimg_per_tick       = 50,       # Interval of progress prints.,
    kimg_per_tick_print = None,
    snapshot_ticks      = 50,       # How often to save network snapshots, None = disable.,
    state_dump_ticks    = 500,      # How often to dump training state, None = disable.,
    
    total_kimg          = 200000,   # Training duration, in thousands of training images.,
    resume_pkl          = None,     # Start from the given network snapshot, None = random initialization.,
    resume_state_dump   = None,     # Start from the given training state, None = reset training state.,
    resume_kimg         = 0,        # Start from the given training progress.,
    
    use_ema             = True,     # Whether or not to apply EMA to model weights.,
    ema_halflife_kimg   = 500,      # Half-life of the exponential moving average (EMA) of model weights.,
    
    dataset_kwargs      = {},       # Options for training set.,
    data_loader_kwargs  = {},       # Options for constructing dataloader.,
    dataset_obj         = None,     # Pre-generated dataset object.,
    dset_samples        = None,     # Pre-generated dataset samples.,
    batch_size          = 512,      # Total batch size for one training iteration.,
    batch_gpu           = None,     # Limit batch size per GPU, None = no limit.,
    
    network_kwargs      = {},       # Options for model and preconditioning.,
    loss_kwargs         = {},       # Options for loss function.,
    optimizer_kwargs    = {},       # Options for optimizer.,
    loss_scaling        = 1,        # Loss scaling factor for reducing FP16 under/overflows.,
    grad_clip           = False,    # whether or no to apply grad norm clipping to model params,
    grad_clip_val       = None,     # val to clip model grad norms to.,
    d_penalty_type      = None,     # 'none' | 'ridge' | 'lasso' | 'horseshoe'
    d_penalty_lambda    = None,     # if None and type != 'none': adapt online
    d_hs_tau            = None,     # global scale τ for horseshoe
    
    alpha               = 1.0,      #scale for flow net loss,
    beta                = 1.0,      #scale for dynamics net loss,
    gamma               = 1.0,      #scale for Lie derivative loss,
    eta                 = 1.0,      #scale for encoder loss,
    eta_decay_start_kimg = 1e9,
    eta_decay_halflife_kimg = 0.0,
    eta_floor = 0.0,
    
    hist_noise_std      = 0.0,
    hist_lag_dim        = 0,
    hist_static_dim     = 0,

    k_target=None,
    ndrop_ramp_kimg     = 0.0,
    ndrop_p0            = 1.0,

):
    
    if kimg_per_tick_print is None:
        kimg_per_tick_print = kimg_per_tick
    
    np.random.seed((seed * dist.get_world_size() + dist.get_rank()) % (1 << 31))
    torch.manual_seed(np.random.randint(1 << 31))
    torch.backends.cudnn.benchmark = cudnn_benchmark
    torch.backends.cudnn.allow_tf32 = False
    torch.backends.cuda.matmul.allow_tf32 = False
    torch.backends.cuda.matmul.allow_fp16_reduced_precision_reduction = False

    ndrop_p_final = None
    use_ndrop_ramp = False
    ndrop_p0_local = ndrop_p0
    ndrop_ramp_kimg = float(ndrop_ramp_kimg or 0.0)

    ndrop_K0 = None
    ndrop_Kf = None
    

    batch_gpu_total = batch_size // dist.get_world_size()
    if batch_gpu is None or batch_gpu > batch_gpu_total:
        batch_gpu = batch_gpu_total
    num_accumulation_rounds = batch_gpu_total // batch_gpu
    assert batch_size == batch_gpu * num_accumulation_rounds * dist.get_world_size()

    # TB
    dist.print0('Creating TB dir and writer...')
    if dist.get_rank() == 0:
        writer_dir = os.path.join(run_dir, 'TB_logs')
        os.makedirs(writer_dir, exist_ok=True) 
        writer = SummaryWriter(log_dir = writer_dir)
    else:
        writer = None

    # data loader
    dist.print0('Using provided dataset...')
    dataset_sampler = misc.InfiniteSampler(dataset=dataset_obj, rank=dist.get_rank(), num_replicas=dist.get_world_size(), seed=seed)  
    dataset_iterator = iter(torch.utils.data.DataLoader(dataset=dataset_obj, sampler=dataset_sampler, \
                                                        batch_size=batch_gpu, **data_loader_kwargs))

    # construct model
    dist.print0('Constructing network...')
    net = dnnlib.util_v7.construct_class_by_name(**network_kwargs) # subclass of torch.nn.Module
    net.train().requires_grad_(True).to(device)
    enc_mod = _get_enc_mod(net)
    # 12/02/2025
    encoder_param_ids = {id(p) for p in enc_mod.parameters()}
    lambda_d = d_penalty_lambda
    
    if hasattr(enc_mod, 'd'):
        kmax = int(getattr(enc_mod, 'k_max', 0)) or enc_mod.d.numel()
        kmax = max(1, kmax)
        if k_target is not None:
            k_use = max(1, min(int(k_target), kmax))
            label = "k_target"
        else:
            k_use = max(1, kmax // 2)   # or kmax
            label = "half k_budget"
        enc_mod.ndrop_p = 1.0 / float(k_use)
        ndrop_p_final = float(enc_mod.ndrop_p)
        dist.print0(f"[ND] {label}={k_use} -> base ndrop_p={enc_mod.ndrop_p:.4f}")
    else:
        ndrop_p_final = None

    # print summary once on rank0
    if dist.get_rank() == 0:
        with torch.no_grad():
            images = torch.zeros([batch_gpu, net.data_dim], device=device)
            ts = torch.ones([batch_gpu], device=device)
            test_flow_matcher = cfm.ConditionalFlowMatcher(sigma=0.1)
            cov_dynamic_0 = cov_dynamic_dt = cov_static = None

            if getattr(net, 'dim_cov_dynamic', 0) > 0:
                cov_dynamic_0 = torch.zeros([batch_gpu, net.dim_cov_dynamic], device=device)
                cov_dynamic_dt = torch.zeros([batch_gpu, net.dim_cov_dynamic], device=device)
            if getattr(net, 'dim_cov_static', 0) > 0:
                cov_static = torch.zeros([batch_gpu, net.dim_cov_static], device=device)

            if (cov_dynamic_0 is not None) or (cov_static is not None):
                misc.print_module_summary(
                    net,
                    [images, images, ts, ts, 1e-3,
                     test_flow_matcher, test_flow_matcher,
                     cov_dynamic_0, cov_dynamic_dt, cov_static],
                    max_nesting=2,
                )
            else:
                misc.print_module_summary(
                    net,
                    [images, images, ts, ts, 1e-3,
                     test_flow_matcher, test_flow_matcher],
                    max_nesting=2,
                )

    gc.collect()
    torch.cuda.empty_cache()

    # boeadcast initial weights
    if torch.distributed.is_available() and torch.distributed.is_initialized():
        for t in net.state_dict().values():
            if torch.is_tensor(t):
                torch.distributed.broadcast(t, src=0)

    # loss function
    dist.print0('Setting up optimizer and loss fn...')
    loss_fn = dnnlib.util_v7.construct_class_by_name(**loss_kwargs)

    # optimizer + LR schedule
    abs_floor_train = float(optimizer_kwargs.pop('lr_floor_abs', 1e-4))
    lr_decay_kimg  = float(optimizer_kwargs.pop('lr_decay_kimg', 0.0))  # 0: disabled
    lr_gamma       = float(optimizer_kwargs.pop('lr_gamma', 0.5))       # step factor (e.g., 0.5)
    
    optimizer = dnnlib.util_v7.construct_class_by_name(
        params=[{'params': net.parameters(), 'weight_decay': 0.0}],
        **optimizer_kwargs
    )

    ddp = torch.nn.parallel.DistributedDataParallel(net, device_ids=[torch.cuda.current_device()],
                                                    broadcast_buffers=False,
                                                    find_unused_parameters=True)

    base_lr = float(optimizer_kwargs.get('lr', 1e-3))  # one LR from start to finish
    abs_floor = float(abs_floor_train)
    rel_floor = 1e-5
    lr_floor  = max(rel_floor * base_lr, abs_floor)

    # loss EMA for monitoring
    ema_beta_loss = 0.8   # keep loss EMA for logging only
    loss_ema = None
    
    # print accumulators
    print_steps = 0
    print_total_sum = 0.0
    print_cmp_sum = 0.0
    print_dyn_sum = 0.0
    print_con_unmasked_sum = 0.0
    print_K_sum = 0.0
    print_K_sqsum = 0.0
    print_K_cnt = 0

    # tick accumulators (per train tick)
    tick_steps = 0
    tick_total_sum = 0.0
    tick_cmp_sum = 0.0
    tick_dyn_sum = 0.0
    tick_con_unmasked_sum = 0.0
    tick_K_sum = 0.0
    tick_K_sqsum = 0.0
    tick_K_cnt = 0

    # EMA weights
    if use_ema: 
        ema = copy.deepcopy(net).eval().requires_grad_(False)
    else:
        ema = None

    # resume full training state if given
    # w_adapt_state = None
    if resume_state_dump:
        dist.print0(f'Loading training state from "{resume_state_dump}"...')
        all_nn_classes = [obj for name, obj in inspect.getmembers(nn) if inspect.isclass(obj) and obj.__module__.startswith('torch.nn')]
        torch.serialization.add_safe_globals(all_nn_classes + [persistence._reconstruct_persistent_obj])
        data = torch.load(resume_state_dump, map_location=torch.device('cpu'))

        misc.copy_params_and_buffers(src_module=data['net'], dst_module=net, require_all=False)
        optimizer.load_state_dict(data['optimizer_state'])
        
        _src_net = data['net']
        for _name in ('use_t_dyn', 'include_x0_tau', 'dim_cov_dynamic', 'dim_cov_static'):
            if hasattr(_src_net, _name) and hasattr(net, _name):
                setattr(net, _name, getattr(_src_net, _name))
        
        del data  # conserve memory
        dist.print0('[resume] Restored net + optimizer + forward flags from training-state dump.')
        
    # resume EMA from pkl if needed
    if resume_pkl is not None and use_ema:
        dist.print0(f'Loading EMA weights from "{resume_pkl}" (net untouched)...')
        with dnnlib.util.open_url(resume_pkl, verbose=(dist.get_rank() == 0)) as f:
            data = pickle.load(f)
        misc.copy_params_and_buffers(src_module=data['ema'], dst_module=ema, require_all=False)
        del data
        dist.print0('[resume] EMA set from .pkl.')

    # progress init
    dist.print0(f'Training for {total_kimg} kimg...')
    dist.print0()
    cur_nimg = int(resume_kimg) * 1000

    if (ndrop_p_final is not None) and (ndrop_ramp_kimg > 0.0) and (ndrop_p0_local > ndrop_p_final):
        use_ndrop_ramp = True

        # define expected K at start and end of ramp
        eps = 1e-8
        ndrop_K0 = 1.0 / max(float(ndrop_p0_local), eps)      # ~1 if p0=1
        ndrop_Kf = 1.0 / max(float(ndrop_p_final), eps)       # ~k_use

        cur_kimg = float(cur_nimg) / 1000.0
        if cur_kimg >= ndrop_ramp_kimg:
            K_des = ndrop_Kf
        else:
            u = max(0.0, min(1.0, cur_kimg / ndrop_ramp_kimg))
            K_des = ndrop_K0 + (ndrop_Kf - ndrop_K0) * u
        new_p = 1.0 / max(K_des, 1.0)

        enc_mod.ndrop_p = float(new_p)
        if dist.get_rank() == 0:
            dist.print0(
                f"[ND] Ramp enabled: K0≈{ndrop_K0:.2f}, Kf≈{ndrop_Kf:.2f}, "
                f"p0={ndrop_p0_local:.4f}, p_final={ndrop_p_final:.4f}, "
                f"ramp_kimg={ndrop_ramp_kimg:.1f}, start_p={enc_mod.ndrop_p:.4f}, "
                f"start_K_des≈{K_des:.2f} at kimg={cur_kimg:.1f}"
            )
    elif ndrop_p_final is not None and dist.get_rank() == 0:
        dist.print0(f"[ND] Ramp disabled, using fixed ndrop_p={enc_mod.ndrop_p:.4f}")

    # ---- state for D-penalty adaptation (per training_loop call) ----
    lambda_d_state = None       # current lamda_t
    d_L_main_ema   = None       # EMA of unmasked recon loss
    d_R_core_ema   = None       # EMA of core penalty R_core(D)
    hs_tau = None
    if (d_penalty_type or 'none').lower() == 'horseshoe':
        if d_hs_tau is not None:
            hs_tau = float(d_hs_tau)
            if dist.get_rank() == 0:
                dist.print0(f"[Horseshoe] Using fixed tau={hs_tau:.3e} from CLI.")
        else:
            if dist.get_rank() == 0:
                num_batches = 8
                sum_sq = 0.0
                n_samples = 0
                tmp_loader = torch.utils.data.DataLoader(
                    dataset=dataset_obj,
                    batch_size=min(batch_gpu, 256),
                    shuffle=True
                )
                for b_idx, batch in enumerate(tmp_loader):
                    if b_idx >= num_batches:
                        break
                    x0 = batch[0].float()      # [B, C, ...] or [B, D]
                    B = x0.shape[0]
                    x0 = x0.view(B, -1)        # [B, data_dim]
                    x0 = x0 - x0.mean(dim=0, keepdim=True)
                    sum_sq += float((x0 * x0).sum().item())
                    n_samples += B

                if n_samples > 0:
                    trace_cov = sum_sq / n_samples        # ≈ tr(Cov(x))
                    d_len = int(enc_mod.d.numel()) if hasattr(enc_mod, 'd') else 1
                    hs_tau = trace_cov / max(1, d_len)    # average variance per latent
                else:
                    hs_tau = 1.0

                dist.print0(
                    f"[Horseshoe] Data-based tau={hs_tau:.3e} "
                    f"(trace_cov≈{trace_cov:.3e}, D_len={d_len})"
                )

            # broadcast hs_tau from rank 0
            hs_tau_tensor = torch.tensor([0.0 if hs_tau is None else hs_tau],
                                         device=device, dtype=torch.float32)
            if torch.distributed.is_available() and torch.distributed.is_initialized():
                torch.distributed.broadcast(hs_tau_tensor, src=0)
            hs_tau = float(hs_tau_tensor.item())
    

    cur_tick = 0
    tick_start_nimg = cur_nimg
    tick_start_time = time.time()

    print_tick = 0
    print_tick_start_nimg = cur_nimg
    print_tick_start_time = tick_start_time
    
    dist.update_progress(cur_nimg // 1000, total_kimg)
    stats_jsonl = None
    gs = 0 
    loss_scales = torch.Tensor([alpha, beta, eta, gamma]).type(torch.float32).to(device)
    eta_base = eta
    
    # ----------------- training loop ----------------- #
    while True:

        optimizer.zero_grad(set_to_none=True)
        tot_scalar_loss = 0
        tot_separate_losses = torch.zeros(4).type(torch.float32).to(device)

        cur_kimg_eta = float(cur_nimg) / 1000.0
        eta_eff = eta_base
        if (eta_decay_halflife_kimg is not None) and (float(eta_decay_halflife_kimg) > 0.0) and (cur_kimg_eta >= float(eta_decay_start_kimg)):
            t = cur_kimg_eta - float(eta_decay_start_kimg)
            eta_eff = eta_base * (0.5 ** (t / float(eta_decay_halflife_kimg)))
            eta_eff = max(float(eta_floor), float(eta_eff))

        loss_scales[0] = alpha
        loss_scales[1] = beta
        loss_scales[2] = eta_eff
        loss_scales[3] = gamma
        if dist.get_rank() == 0:
            writer.add_scalar('cmp_train_loss/weight', float(alpha), gs)
            writer.add_scalar('dyn_train_loss/weight', float(beta),  gs)
            writer.add_scalar('enc_recon_loss/weight', float(eta_eff), gs)
            writer.add_scalar('lie_derivative_loss/weight', float(gamma), gs)
        

        # ----- forward/backward with grad accumulation ----- #
        step_unmask_sum = 0.0
        
        for round_idx in range(num_accumulation_rounds):
            last_round = (round_idx == num_accumulation_rounds - 1)
            with misc.ddp_sync(ddp, last_round): 
                batch_data = next(dataset_iterator)
                x0_1 = batch_data[0].type(torch.float32).reshape(batch_data[0].shape[0], -1).to(device)
                xdt_1 = batch_data[1].type(torch.float32).reshape(batch_data[1].shape[0], -1).to(device)

                idx = 3
                cov_dynamic_0 = None
                cov_dynamic_dt = None
                cov_static = None

                if hasattr(net, 'dim_cov_dynamic') and net.dim_cov_dynamic > 0:
                    cov_dynamic_0 = batch_data[idx].type(torch.float32).to(device)
                    cov_dynamic_dt = batch_data[idx+1].type(torch.float32).to(device)
                    idx += 2

                if hasattr(net, 'dim_cov_static') and net.dim_cov_static > 0:
                    cov_static = batch_data[idx].type(torch.float32).to(device)
                    if hist_noise_std > 0 and hist_lag_dim > 0:
                        start = hist_static_dim
                        cov_static[:, start:start+hist_lag_dim] += torch.randn_like(
                            cov_static[:, start:start+hist_lag_dim]
                        ) * float(hist_noise_std)


                loss = loss_fn(net=ddp, x0_1=x0_1, xdt_1=xdt_1, dt=dataset_kwargs.dt,
                               cov_dynamic_0=cov_dynamic_0,
                               cov_dynamic_dt=cov_dynamic_dt,
                               cov_static=cov_static)


                loss = loss * loss_scales[:, None, None] # Scale losses
                training_stats.report('Loss/loss', torch.sum(loss.detach(), dim=0)) # Log losses
                round_scalar_loss = loss.sum() * (loss_scaling / batch_gpu_total)


                sep = torch.sum(torch.sum(loss.detach(), dim=2), dim=1)
                tot_separate_losses += sep
                tot_scalar_loss += round_scalar_loss.item()
                round_scalar_loss.backward()

                # collect unmasked recon from loss fn if provided
                _u = getattr(loss_fn, 'last_unmasked_sum', None)
                if _u is not None:
                    step_unmask_sum += float(_u)
        
        # normalize per-component losses to per-sample scale for this step
        tot_separate_losses *= (loss_scaling / batch_gpu_total)
        
        # unmasked recon metric
        if step_unmask_sum > 0.0:
            step_con_unmasked = (loss_scaling / batch_gpu_total) * step_unmask_sum
        else:
            step_con_unmasked = float(tot_separate_losses[2].item())

        # ----- LR update ----- #
        if lr_decay_kimg > 0.0:
            # Simple decay based on total progress
            steps = max(0, int((cur_nimg) // int(lr_decay_kimg * 1000)))
            current_lr = max(lr_floor, base_lr * (lr_gamma ** steps))
        else:
            current_lr = base_lr

        # ----- ND ramp update ----- #
        if use_ndrop_ramp and (ndrop_p_final is not None) and (ndrop_K0 is not None) and (ndrop_Kf is not None):
            cur_kimg = float(cur_nimg) / 1000.0
            if cur_kimg >= ndrop_ramp_kimg:
                K_des = ndrop_Kf
            else:
                u = max(0.0, min(1.0, cur_kimg / ndrop_ramp_kimg))
                K_des = ndrop_K0 + (ndrop_Kf - ndrop_K0) * u
            new_p = 1.0 / max(K_des, 1.0)
            enc_mod.ndrop_p = float(new_p)
        
                    
        for g in optimizer.param_groups:
            g['lr'] = float(current_lr)

        with torch.no_grad():
            for p in net.parameters():
                if p.grad is not None:
                    p.grad.nan_to_num_(nan=0.0, posinf=1e5, neginf=-1e5)
            
        if grad_clip: 
            assert grad_clip_val != None, 'Need a value to clip grads to!'
            torch.nn.utils.clip_grad_norm_(net.parameters(), grad_clip_val)
        
        optimizer.step()

        # ------------------------------------------------------------------
        # Encoder D penalty (ridge / lasso / horseshoe), applied outside
        # autograd. Uses EMA of *unmasked* recon loss to keep penalty mild.
        # ------------------------------------------------------------------
        penalty_type = (d_penalty_type or 'none').lower()

        current_kimg = cur_nimg / 1000.0
        is_ramping = use_ndrop_ramp and (current_kimg < ndrop_ramp_kimg)
        
        if (not is_ramping) and penalty_type != 'none' and hasattr(enc_mod, 'd') and hasattr(enc_mod, 'last_K'):
            with torch.no_grad():
                d_param = enc_mod.d
                d_len = enc_mod.d.numel()
                K_max = int(min(int(getattr(enc_mod, 'k_max', d_len)), d_len))
                if K_max > 0:
                    d_min_attr = float(getattr(enc_mod, 'd_min', 1e-15))

                    # current D values and logs
                    d_log = d_param.data
                    region_log = d_log[:K_max]
                    region = region_log.exp()
                    region.clamp_(min=d_min_attr)

                    # main loss: unmasked reconstruction (per-sample)
                    L_main_t = float(step_con_unmasked)

                    # penalty "energy" R_core_t depending on type
                    if penalty_type == 'ridge':
                        R_core_t = float((region * region).mean().item())
                    elif penalty_type == 'lasso':
                        R_core_t = float(region.mean().item())
                    elif penalty_type == 'horseshoe':
                        tau = float(hs_tau)
                        tau2 = tau * tau
                        R_core_t = float(
                            torch.log1p(tau2 / (region * region + 1e-30)).mean().item())
                    else:
                        R_core_t = 0.0

                    # ---- update EMAs for L_main and R_core ----
                    beta_ema = 0.95
                    if d_L_main_ema is None:
                        d_L_main_ema = L_main_t
                        d_R_core_ema = R_core_t
                        lambda_d_state = (1e-4 if d_penalty_lambda is None
                                          else float(d_penalty_lambda))
                    else:
                        d_L_main_ema = beta_ema * d_L_main_ema + (1.0 - beta_ema) * L_main_t
                        d_R_core_ema = beta_ema * d_R_core_ema + (1.0 - beta_ema) * R_core_t

                    # ---- choose / adapt lambda_d ----
                    if d_penalty_lambda is None:
                        rho0 = 0.05
                        if d_R_core_ema > 0.0 and d_L_main_ema > 0.0:
                            target_lambda = rho0 * d_L_main_ema / (d_R_core_ema + 1e-12)
                            # smooth in log-space to avoid jumps
                            lam = lambda_d_state
                            lam = lam * math.exp(
                                0.1 * math.log(max(target_lambda, 1e-12) /
                                               max(lam, 1e-12)))
                            lambda_d_state = float(max(1e-8, min(lam, 1.0)))
                    else:
                        lambda_d_state = float(d_penalty_lambda)

                    lambda_d = lambda_d_state
                    if lambda_d > 0.0:
                        lr_here = float(current_lr)
                        t = lambda_d * lr_here
                        t = max(0.0, min(t, 0.1))   # gentle step

                        if t > 0.0:
                            if penalty_type == 'ridge':
                                region.mul_(1.0 / (1.0 + t))
                                region.clamp_(min=d_min_attr)
                                region_log.copy_(region.log())

                            elif penalty_type == 'lasso':
                                region.sub_(t)
                                region.clamp_(min=d_min_attr)
                                region_log.copy_(region.log())

                            elif penalty_type == 'horseshoe':
                                tau = float(hs_tau)
                                tau2 = tau * tau
                                shrink = 1.0 / (1.0 + (t * tau2 /
                                                       (region * region + 1e-30)))
                                region.mul_(shrink)
                                region.clamp_(min=d_min_attr)
                                region_log.copy_(region.log())
                                

        if hasattr(enc_mod, 'orthonormalize_L'):
            enc_mod.orthonormalize_L(threshold=1e-4)
        
        # EMA weights
        if use_ema: 
            ema_halflife_nimg = ema_halflife_kimg * 1000 
            ema_beta = 0.5 ** (batch_size / max(ema_halflife_nimg, 1e-8)) 
            for p_ema, p_net in zip(ema.parameters(), net.parameters()):
                if id(p_net) in encoder_param_ids:
                    p_ema.copy_(p_net.detach())
                else:
                    p_ema.copy_(p_net.detach().lerp(p_ema, ema_beta))
                # p_ema.copy_(p_net.detach().lerp(p_ema, ema_beta))           

        # TB scalars
        if writer is not None:
            writer.add_scalar('tot_train_loss', tot_scalar_loss, gs)
            writer.add_scalar('cmp_train_loss/value',  float(tot_separate_losses[0].item()), gs)
            writer.add_scalar('dyn_train_loss/value',  float(tot_separate_losses[1].item()), gs)
            writer.add_scalar('enc_recon_loss/value',  float(tot_separate_losses[2].item()), gs)
            writer.add_scalar('enc_recon_unmasked/value', float(step_con_unmasked), gs)
            writer.add_scalar('lie_derivative_loss/value', float(tot_separate_losses[3].item()), gs)
            writer.add_scalar('Kimgs', cur_nimg / 1000.0, gs)
            gs += 1

        # ----- accumulate into train-tick + print-window ----- #
        tick_recon = float(step_con_unmasked)
        tick_cmp   = float(tot_separate_losses[0].item())
        tick_dyn   = float(tot_separate_losses[1].item())

        # tick accumulators
        tick_steps += 1
        tick_total_sum += float(tot_scalar_loss)
        tick_cmp_sum += tick_cmp
        tick_dyn_sum += tick_dyn
        tick_con_unmasked_sum += tick_recon

        if hasattr(enc_mod, 'last_K'):
            k_val = float(enc_mod.last_K)
        else:
            k_val = None

        if (k_val is not None) and (k_val > 0.0):
            tick_K_sum += k_val
            tick_K_sqsum += k_val * k_val
            tick_K_cnt += 1
            print_K_sum += k_val
            print_K_sqsum += k_val * k_val
            print_K_cnt += 1

        # print-window accumulators
        print_steps += 1
        print_total_sum += float(tot_scalar_loss)
        print_cmp_sum += tick_cmp
        print_dyn_sum += tick_dyn
        print_con_unmasked_sum += tick_recon

        cur_nimg += batch_size
        done = (cur_nimg >= total_kimg * 1000)

        # ----- logging: print window (tick_print) ----- #
        print_end_time = time.time()
        do_print = ((cur_nimg - print_tick_start_nimg) >= kimg_per_tick_print * 1000) or done

        if do_print:
            denom = max(1, print_steps)
            cmp_log = print_cmp_sum / denom
            dyn_log = print_dyn_sum / denom
            tot_log = print_total_sum / denom
            con_unm = print_con_unmasked_sum / denom

            tick_val = training_stats.report0('Progress/tick', cur_tick)
            sec_print = print_end_time - print_tick_start_time
            sec_print_val = training_stats.report0('Timing/sec_per_print', sec_print)
            sec_kimg = (print_end_time - print_tick_start_time) / max(
                1, (cur_nimg - print_tick_start_nimg)
            ) * 1e3
            sec_kimg_val = training_stats.report0('Timing/sec_per_kimg', sec_kimg)

            if dist.get_rank() == 0:
                fields = []
                fields += [f"tick {tick_val:<5d}"]
                fields += [f"sec/print {sec_print_val:<7.1f}"]
                fields += [f"sec/kimg {sec_kimg_val:<7.2f}"]
                fields += [f"cmp_loss {cmp_log:<7.3f}"]
                fields += [f"dyn_loss {dyn_log:<7.3f}"]
                fields += [f"con_unmasked_avg {con_unm:<7.3f}"]
                fields += [f"total_loss {tot_log:<7.3f}"]

                if print_K_cnt > 0:
                    K_mean = print_K_sum / print_K_cnt
                    K_var = max(0.0, print_K_sqsum / print_K_cnt - K_mean * K_mean)
                    fields += [f"K-mean {K_mean:<4.2f}"]
                    fields += [f"K-std {K_var**0.5:<4.2f}"]

                torch.cuda.reset_peak_memory_stats()
                dist.print0(' '.join(fields))

            # reset print-window accumulators ON ALL RANKS
            print_steps = 0
            print_total_sum = 0.0
            print_cmp_sum = 0.0
            print_dyn_sum = 0.0
            print_con_unmasked_sum = 0.0
            print_K_sum = 0.0
            print_K_sqsum = 0.0
            print_K_cnt = 0
            print_tick_start_nimg = cur_nimg
            print_tick_start_time = print_end_time
            print_tick += 1
        
        # ----- abort / snapshots / state dumps ----- #
        if (not done) and dist.should_stop():
            done = True
            dist.print0()
            dist.print0('Aborting...')

        # ----- train-tick boundary check ----- #
        want_more_in_tick = (not done) and ((cur_nimg - tick_start_nimg) < kimg_per_tick * 1000)
        if torch.distributed.is_available() and torch.distributed.is_initialized():
            tflag = torch.tensor(
                [1 if want_more_in_tick else 0],
                device=device,
                dtype=torch.int32,
            )
            torch.distributed.all_reduce(tflag, op=torch.distributed.ReduceOp.SUM)
            want_more_in_tick = (tflag.item() > 0)

        if want_more_in_tick:
            dist.update_progress(cur_nimg // 1000, total_kimg)
            if done:
                break
            continue

        # ================= END OF TRAIN TICK =================
        # Compute means for EMAs & stage gating using tick_* accumulators.
        tick_total_mean = tick_total_sum / max(1, tick_steps)
        tick_recon_mean = tick_con_unmasked_sum / max(1, tick_steps)
        cmp_tick_mean   = tick_cmp_sum / max(1, tick_steps)
        dyn_tick_mean   = tick_dyn_sum / max(1, tick_steps)

        # Monitoring EMA
        metric_val = tick_total_mean
        metric_t = torch.tensor([metric_val], device=device, dtype=torch.float32)
        if torch.distributed.is_available() and torch.distributed.is_initialized():
            torch.distributed.all_reduce(metric_t, op=torch.distributed.ReduceOp.AVG)
        metric = float(metric_t.item())
        if loss_ema is None:
            loss_ema = metric
        else:
            loss_ema = ema_beta_loss * loss_ema + (1.0 - ema_beta_loss) * metric
        
        # reset tick accumulators
        tick_steps = 0
        tick_total_sum = 0.0
        tick_cmp_sum = 0.0
        tick_dyn_sum = 0.0
        tick_con_unmasked_sum = 0.0
        tick_K_sum = 0.0
        tick_K_sqsum = 0.0
        tick_K_cnt = 0
            
        # advance tick window
        tick_start_nimg = cur_nimg
        tick_start_time = time.time()
        cur_tick += 1
            
        # snapshots (use EMA or net)
        if (snapshot_ticks is not None) and (done or (cur_tick % snapshot_ticks == 0)):
            src_enc = _get_enc_mod(ddp)
            tgt_mod = ema if (use_ema and ema is not None) else net
            tgt_enc = _get_enc_mod(tgt_mod)
            _copy_pop_stats(src_enc, tgt_enc)

            if use_ema and ema is not None:
                snap_data = dict(ema=ema, loss_fn=loss_fn, dataset_kwargs=dict(dataset_kwargs))
            else:
                snap_data = dict(net=net, loss_fn=loss_fn, dataset_kwargs=dict(dataset_kwargs))
            for k, v in list(snap_data.items()):
                if isinstance(v, torch.nn.Module):
                    m = copy.deepcopy(v).eval().requires_grad_(False).to(device)
                    snap_data[k] = m.cpu()
                    del m
                else:
                    snap_data[k] = v

            if dist.get_rank() == 0:
                name = f'network-snapshot-{cur_nimg//1000:06d}.pkl'
                with open(os.path.join(run_dir, name), 'wb') as f:
                    pickle.dump(snap_data, f)
            del snap_data
            
        # training state dump 
        if dist.get_rank() == 0 and (
            done or (state_dump_ticks is not None and (cur_tick % state_dump_ticks == 0))
        ):
            tname = f"training-state-{cur_nimg//1000:06d}.pt"
            torch.save(
                {
                    "net": net,
                    "optimizer_state": optimizer.state_dict(),
                },
                os.path.join(run_dir, tname),
            )

        # # optional histograms / trajectory viz at state dumps
        # if (
        #     state_dump_ticks is not None
        #     and (done or (cur_tick % state_dump_ticks == 0))
        #     and dist.get_rank() == 0
        #     and writer is not None
        # ):
        #     for tag, value in net.named_parameters():
        #         if value.grad is not None:
        #             writer.add_histogram(tag + "/grad", value.grad.cpu(), gs)

        #     # Generic recon viz using EMA if available.
        #     model_viz = ema if (use_ema and ema is not None) else net
        #     dnnlib.util_v7.log_reconstruction_viz(writer, model_viz, dset_samples, device, gs)
        
        # stats.jsonl
        training_stats.default_collector.update()
        if dist.get_rank() == 0:
            if stats_jsonl is None:
                stats_jsonl = open(os.path.join(run_dir, 'stats.jsonl'), 'at')
            stats_jsonl.write(json.dumps(dict(training_stats.default_collector.as_dict(),
                                              timestamp=time.time())) + '\n')
            stats_jsonl.flush()

        dist.update_progress(cur_nimg // 1000, total_kimg)
        if done: break
    
    dist.print0()
    dist.print0('Exiting...')
```
