## Supplementary material for "Dynamic Compression Flows for Neuroscience Data": Python code: vfm_train_v7.py

```
#!/usr/bin/env python3
# -*- coding: utf-8 -*-
"""

Implements VFM model training 

"""


import os
import gc
import re
import json
import click
import torch
import dnnlib
import numpy as np
from torch_utils import distributed as dist
from training import toy_training_loop_vfm_v7
import warnings
warnings.filterwarnings('ignore', 'Grad strides do not match bucket view strides') # False warning printed by PyTorch 1.12.

def _to_list_of_arrays(x):
    """Normalize npz-loaded 'samples' into a Python list of arrays."""
    if isinstance(x, list):
        return [np.asarray(a) for a in x]
    x = np.asarray(x, dtype=object)
    if x.dtype == object:                  # object array of trials
        return [np.asarray(a) for a in x.tolist()]
    if x.ndim == 3:                        # (n_trial, T, D) or (n_trial, T, H, W)
        return [x[i] for i in range(x.shape[0])]
    if x.ndim == 5:                        # (n_trial, T, C, H, W)
        return [x[i] for i in range(x.shape[0])]
    raise RuntimeError(f"Unsupported dataset shape: {getattr(x,'shape',None)}")


def _is_movie_trials(trials):
    return all(arr.ndim in (3, 4) for arr in trials)   # (T,H,W) or (T,C,H,W)

def _ensure_TCHW(trials, dtype=np.float32):
    """Ensure each trial is (T, C, H, W) and dtype=float32. No value scaling."""
    out = []
    for a in trials:
        if a.ndim == 3:               # (T, H, W) -> add channel
            a = a[:, None, :, :]
        elif a.ndim == 4:             # already (T, C, H, W)
            pass
        else:
            raise RuntimeError(f"Expected image trial, got {a.shape}")
        out.append(a.astype(dtype, copy=False))
    return out


def _try_load_cov(path_npy, path_npz_key='samples'):
    if os.path.isfile(path_npy):
        arr = np.load(path_npy, allow_pickle=True)
        if isinstance(arr, np.ndarray) and arr.dtype == object:
            try:
                return arr.tolist()
            except:
                return arr
        return arr
    path_npz = os.path.splitext(path_npy)[0] + '.npz'
    if os.path.isfile(path_npz):
        arr = np.load(path_npz, allow_pickle=True)[path_npz_key]
        if isinstance(arr, np.ndarray) and arr.dtype == object:
            try:
                return arr.tolist()
            except:
                return arr
        return arr
    return None

def _make_lag_cov_list(trajs, k_):
    lag_list = []
    for X in trajs:
        X = np.asarray(X)
        if X.ndim != 2:
            raise click.ClickException("lag_k covariates are only supported for vector trials (T, D).")
        T, D = X.shape
        if k_ == 0:
            lag = X.copy()
        else:
            blocks = [X[k_-j : T-j] for j in range(k_ + 1)]
            lag = np.concatenate(blocks, axis=1)
        lag_list.append(lag)
    return lag_list

def _parse_channels_csv(s: str):
    """Parse channels from '32,64,128,256' or '32x64x128x256'. Returns list[int] or None."""
    if not s:
        return None
    import re
    toks = [t for t in re.split(r'[ ,x/]+', s) if t]
    return [int(t) for t in toks]

@click.command()

# Main Options (adapted for loading data)
@click.option('--data_path',               help='Path to the folder containing dataset_samples.npz', metavar='DIR', type=str, required=True)
@click.option('--data_name',               help='Name of the dataset', metavar='STR', type=str, required=True)
@click.option('--k_max',                   help='Max number of dimensions to use (budget)', metavar='INT', type=int, required=True)
@click.option('--k_target',
              help='Fix nested-dropout effective K (1..k_max); if omitted, use adaptive K.',
              metavar='INT', type=int, default=None, show_default=True)
@click.option('--dt',                      help='Time interval between successive pts in sampled trajectories', metavar='FLOAT', type=float, default=1e-3, show_default=True)

# Covariate options
@click.option('--use_dynamic_covariates/--no_use_dynamic_covariates',  help='Whether to use dynamic covariates (if available)', default=False, show_default=True)
@click.option('--use_static_covariates/--no_use_static_covariates',    help='Whether to use static covariates (if available)', default=False, show_default=True)

@click.option('--dim_cov_dynamic',         help='Dimension of dynamic covariates [default: inferred from data]', metavar='INT', type=int)
@click.option('--dim_cov_static',          help='Dimension of static covariates [default: inferred from data]', metavar='INT', type=int)
@click.option('--lag_k',                   help='Lag order k to build lag-k covariate matrix (>=0)', metavar='INT', type=click.IntRange(min=0), default=0, show_default=True)
@click.option('--hist_noise_std',          help='Std of Gaussian noise added to lag-history features during training (0 disables).', metavar='FLOAT', type=click.FloatRange(min=0), default=0.0, show_default=True)

# FM options
@click.option('--flow_matcher_type',       help='Flow matching implementation to use.', metavar='regular|exactot|sinkhorn', type=click.Choice(['regular', 'exactot', 'sinkhorn']), default='regular', show_default=True)
@click.option('--sigma_dyn_fm',            help='Sigma val for dynamics flow matcher class', metavar='FLOAT', type=float, default=0.1, show_default=True)
@click.option('--sigma_comp_fm',           help='Sigma val for compression flow matcher class', metavar='FLOAT', type=float, default=0.1, show_default=True)

# low-D encoder
@click.option('--d_min',                   help='Minimum variance for any/all dimensions in LS. Used to sample x0s.', metavar='FLOAT', type=float, default=1e-15, show_default=True)


# Arch Options
@click.option('--dyn_arch',                help='Dynamics net arch to use.', metavar='ToyConvUNet|ToyMLP|Adapted_ToyConvUNet', type=click.Choice(['ToyConvUNet', 'ToyMLP', 'Adapted_ToyConvUNet']), default='ToyMLP', show_default=True)
@click.option('--use_t_dyn/--no_use_t_dyn', help='Include normalized dynamics time t_dyn=ts/dt in v-net inputs', default=True, show_default=True)
@click.option('--flow_detach/--no_flow_detach',
              help='Detach encoder samples when forming FM targets.',
              default=True, show_default=True)

@click.option('--flow_arch',               help='Flow net arch to use.', metavar='ToyConvUNet|ToyMLP|Adapted_ToyConvUNet', type=click.Choice(['ToyConvUNet', 'ToyMLP', 'Adapted_ToyConvUNet']), default='ToyMLP', show_default=True)
@click.option('--encoder_arch',            help='Network architecture to use for encoder.', metavar='Latent_MLP_VAE|Latent_CNN_VAE|Latent_LargeCNN_VAE|Adapted_Latent_LargeCNN_VAE', type=click.Choice(['Latent_MLP_VAE', 'Latent_CNN_VAE', 'Latent_LargeCNN_VAE', 'Adapted_Latent_LargeCNN_VAE']), default='Latent_MLP_VAE', show_default=True)

@click.option('--encoder_depth',           help='Number of hidden layers in MLP encoder', metavar='INT', type=int, default=2, show_default=True)
@click.option('--encoder_width',           help='Width of each hidden layer in MLP encoder', metavar='INT', type=int, default=10, show_default=True)

@click.option('--conv_ch_cmp',  metavar='CSV', type=str, default='32,64,128,256', help='Channels for compression/flow UNet, e.g. "32,64,128,256" (default: 32,64,128,256)')
@click.option('--conv_embed_cmp', metavar='INT', type=int, default=256, help='Embedding dim for compression/flow UNet (default: 256)')
@click.option('--conv_ch_dyn',  metavar='CSV', type=str, default='32,64,128,256', help='Channels for dynamics UNet, e.g. "32,64,128,256" (default: 32,64,128,256)')
@click.option('--conv_embed_dyn', metavar='INT', type=int, default=256, help='Embedding dim for dynamics UNet (default: 256)')
@click.option('--conv_ch_enc',  metavar='CSV', type=str, default='32,64,128,256', help='Channels for CNN encoder, e.g. "32,64,128,256" (default: 32,64,128,256)')

@click.option('--mlp_depth_cmp',               help='Number of hidden layers in MLP, compression flow nets', metavar='INT', type=int, default=2, show_default=True)
@click.option('--mlp_width_cmp',               help='Width of each hidden layer in MLP, compression flow nets', metavar='INT', type=int, default=64, show_default=True)
@click.option('--mlp_depth_dyn',               help='Number of hidden layers in MLP, dynamic flow nets', metavar='INT', type=int, default=2, show_default=True)
@click.option('--mlp_width_dyn',               help='Width of each hidden layer in MLP, dynamic flow nets', metavar='INT', type=int, default=64, show_default=True)

# Training Hyperparameters.
@click.option('--duration',                help='Training duration', metavar='MIMG', type=click.FloatRange(min=0, min_open=True), default=7000, show_default=True)
@click.option('--batch',                   help='Total batch size', metavar='INT', type=click.IntRange(min=1), default=8192, show_default=True)
@click.option('--batch_gpu',               help='Limit batch size per GPU', metavar='INT', type=click.IntRange(min=1), default=1024, show_default=True)
@click.option('--lr',                      help='initial learning rate', metavar='FLOAT', type=click.FloatRange(min=0, min_open=True), default=1e-5, show_default=True)
@click.option('--lr_floor', help='Absolute LR floor for the adaptive scheduler',
              metavar='FLOAT', type=click.FloatRange(min=0), default=1e-4, show_default=True)
@click.option('--lr_decay_kimg',
              help='Interval in kimg between LR decay steps (0 disables decay)',
              metavar='FLOAT', type=float, default=0.0, show_default=True)

@click.option('--ndrop_ramp_kimg',
              help='Kimg over which nested-dropout p ramps from ndrop_p0 to final p (0 disables ramp)',
              metavar='FLOAT', type=float, default=1000, show_default=True)
@click.option('--ndrop_p0',
              help='Initial nested-dropout geometric probability p0 (default 1.0, expected K ≈ 1)',
              metavar='FLOAT', type=click.FloatRange(min=0.0, max=1.0), default=1.0, show_default=True)


@click.option(
    '--d_penalty_type',
    type=click.Choice(['none', 'ridge', 'lasso', 'horseshoe']),
    default='none',
    show_default=True,
    help="Penalty on encoder D: none | ridge | lasso | horseshoe."
)
@click.option(
    '--d_penalty_lambda',
    type=float,
    default=None,
    help="Global strength lambda for encoder-D penalty. If None and type != 'none', "
         "lambda is adapted online."
)
@click.option(
    '--d_hs_tau',
    type=float,
    default=None,
    show_default=True,
    help="Global scale tau for horseshoe penalty on encoder D. "
         "If None, tau is estimated once from the data scale."
)

@click.option('--use_ema/--no_use_ema',    help='Whether or not to apply EMA to model params', default=True, show_default=True)
@click.option('--ema',                     help='EMA half-life (if using EMA)', metavar='MIMG', type=click.FloatRange(min=0), default=0.5, show_default=True)
@click.option('--alpha',                   help='Scale for flow net component of loss', metavar='FLOAT', type=float, default=1.0, show_default=True)
@click.option('--beta',                    help='Scale for dynamics net component of loss', metavar='FLOAT', type=float, default=1.0, show_default=True)
@click.option('--gamma',                   help='Scale for Lie derivative component of loss', metavar='FLOAT', type=float, default=0.0, show_default=True)
@click.option('--eta',                     help='Scale for encoder reconstruction component of loss', metavar='FLOAT', type=float, default=1.0, show_default=True)
# @click.option('--eta_decay_start_kimg', help='Start (kimg) to begin exponential decay of eta (0 disables start offset).', metavar='FLOAT', type=float, default=0.0, show_default=True)
@click.option('--eta_decay_start_kimg',
              help='Start (kimg) to begin exponential decay of eta (default disables unless explicitly set).',
              metavar='FLOAT', type=float, default=1e9, show_default=True)

@click.option('--eta_decay_halflife_kimg', help='Halflife (kimg) for exponential decay of eta (0 disables).', metavar='FLOAT', type=float, default=0.0, show_default=True)
@click.option('--eta_floor', help='Minimum eta after decay.', metavar='FLOAT', type=float, default=0.0, show_default=True)


@click.option('--grad_clip/--no_grad_clip', help='Whether or not to clip model gradient norm.', default=False, show_default=True)
@click.option('--grad_clip_val',           help='Max value model gradients should be clipped to.', metavar='FLOAT', type=float, default=1.0, show_default=True)

# Performance-related.
@click.option('--ls',                      help='Loss scaling', metavar='FLOAT', type=click.FloatRange(min=0, min_open=True), default=1, show_default=True)
@click.option('--bench/--no_bench',        help='Enable cuDNN benchmarking', default=True, show_default=True)
@click.option('--workers',                 help='DataLoader worker processes', metavar='INT', type=click.IntRange(min=1), default=1, show_default=True)

# I/O-related.
@click.option('--outdir',                  help='Where to save the results', metavar='DIR', type=str, required=True)
@click.option('--desc',                    help='String to include in result dir name', metavar='STR', type=str)
@click.option('--nosubdir/--no_nosubdir',  help='Do not create a subdirectory for results', default=False, show_default=True)
@click.option('--tick_train', help='How often (kimg) to run schedule & gating ticks; default: --tick.',
              metavar='KIMG', type=click.IntRange(min=1))
@click.option('--tick_print', help='How often (kimg) to print logs; default: --tick_train.',
              metavar='KIMG', type=click.IntRange(min=1))
@click.option('--snap',                    help='How often to save snapshots', metavar='TICKS', type=click.IntRange(min=1), default=250, show_default=True)
@click.option('--dump',                    help='How often to dump state', metavar='TICKS', type=click.IntRange(min=1), default=250, show_default=True)
@click.option('--seed',                    help='Random seed  [default: random]', metavar='INT', type=int)
@click.option('--resume',                  help='Resume from previous training state', metavar='PT', type=str)
@click.option('-n', '--dry_run/--no_dry_run', help='Print training options and exit', default=False, show_default=True)


def main(**kwargs):
    
    """
    Set up and launch training for toy_training_loop_vfm_v7.
    """
    
    opts = dnnlib.EasyDict(kwargs)
    torch.multiprocessing.set_start_method('spawn')
    dist.init()

    # (1) Setup & init
    # Setup device
    # # local version
    # device_name = 'cuda' if torch.cuda.is_available() else 'cpu'
    # device = torch.device(device_name)
    
    # DCC version
    local_rank = int(os.environ.get("LOCAL_RANK", "0"))
    if torch.cuda.is_available():
        torch.cuda.set_device(local_rank)
        device = torch.device(f"cuda:{local_rank}")
    else:
        device = torch.device("cpu")
    dist.print0(f"[init] rank={dist.get_rank()} local_rank={local_rank} device={device}")
    
    

    # (2) Load data
    # Load main data
    dataset_samples = np.load(os.path.join(opts.data_path, 'dataset_samples.npz'), allow_pickle=True)
    dset_samples_raw = dataset_samples['samples']
    dset_samples = _to_list_of_arrays(dset_samples_raw)

    if _is_movie_trials(dset_samples):
        dset_samples = _ensure_TCHW(dset_samples)
    
    dset_samples_orig = [np.asarray(a) for a in dset_samples]

    # Load covariate data if available
    cov_dynamic_samples = None
    cov_static_samples = None
    
    # Try to load dynamic covariates
    if opts.use_dynamic_covariates:
        cov_dynamic_path = os.path.join(opts.data_path, 'cov_dynamic_samples.npy')
        cov_dynamic_samples = _try_load_cov(cov_dynamic_path)
    
    # Load static covariates only if flag is set
    if opts.use_static_covariates:
        cov_static_path = os.path.join(opts.data_path, 'cov_static_samples.npy')
        cov_static_samples = _try_load_cov(cov_static_path)


    # (3) Build lag-k covs, truncate, and concatenate
    k = int(max(0, opts.lag_k))
    min_T = min(np.asarray(tr).shape[0] for tr in dset_samples)
    if k >= min_T:
        raise click.ClickException(f'lag_k={k} must be < min trajectory length ({min_T}).')

    is_image = _is_movie_trials(dset_samples)
    if is_image:
        _T, _C, _H, _W = np.asarray(dset_samples[0]).shape
        opts.data_inch = _C
        opts.data_imgshape = _H
    else:
        # place holder
        opts.data_inch = 1
        opts.data_imgshape = 1
        
        

    
    # 3a) construct lag-k cov matrix from original samples
    if is_image:
        # dset_samples: list of (T, C, H, W) — build (T-k, C*(k+1), H, W)
        lag_cov_list = None
    else:
        # dset_samples: list of (T, D) — build (T-k, (k+1)*D)
        lag_cov_list = _make_lag_cov_list(dset_samples, k)

    
    # 3b) truncate dset_samples
    dset_samples_trunc = [np.asarray(X)[k:] for X in dset_samples]
    
    # 3c) truncate cov_dynamic
    cov_dynamic_trunc = None
    if cov_dynamic_samples is not None:
        if isinstance(cov_dynamic_samples, list):
            cov_dynamic_trunc = []
            for C in cov_dynamic_samples:
                C = np.asarray(C)        # (T, Sd) or (T, C, H, W) or (T,)
                Ck = C[k:]               # time-align
                if Ck.ndim >= 2:
                    Ck = Ck.reshape(Ck.shape[0], -1)  # (T-k, Sd_vec)
                else:
                    Ck = Ck[:, None]                   # (T-k,) -> (T-k,1)
                cov_dynamic_trunc.append(Ck)
        elif isinstance(cov_dynamic_samples, np.ndarray):
            # ndarray: either (n_trial, T, Sd) or (n_trial, T, C, H, W)
            if cov_dynamic_samples.ndim == 3:
                # vector covs
                cov_dynamic_trunc = [cov_dynamic_samples[i, k:]                   # (T-k, Sd)
                                     for i in range(cov_dynamic_samples.shape[0])]
            elif cov_dynamic_samples.ndim == 5:
                # image-like covs
                n_trial, T, C, H, W = cov_dynamic_samples.shape
                cov_dynamic_trunc = [cov_dynamic_samples[i, k:].reshape(T - k, -1)  # (T-k, C*H*W)
                                     for i in range(n_trial)]
            else:
                raise click.ClickException("Unsupported cov_dynamic array shape; expected 3D or 5D.")
        else:
            raise click.ClickException("Unsupported cov_dynamic type; expected list or ndarray.")

    # 3d) truncate cov_static if it is a list (time-indexed); if ndarray (per-traj), broadcast
    if cov_static_samples is None:
        # Case 1: lag + static=None
        cov_static_new = None
        cov_static_trunc_only = []
        if is_image:
            # stream per-trial: avoid materializing lag_cov_list
            for X in dset_samples:                      # X: (T, C, H, W)
                X = np.asarray(X)
                T = X.shape[0]
                cov_static_trunc_only.append(np.zeros((T - k, 0), dtype=X.dtype))
        else:
            cov_static_new = []
            cov_static_trunc_only = []
            for L in lag_cov_list:
                L_vec = np.asarray(L)                   # (T-k, (k+1)·D)
                cov_static_new.append(L_vec)            # keep legacy behavior for vectors
                cov_static_trunc_only.append(np.zeros((L_vec.shape[0], 0), dtype=L_vec.dtype))
    
    elif isinstance(cov_static_samples, list):
        # Case 2 & 3: time-indexed static; may be vector (T,S) or image-like (T,Cs,Hs,Ws)
        cov_static_new = None
        cov_static_trunc_only = []
        if is_image:
            for S, X in zip(cov_static_samples, dset_samples):
                S = np.asarray(S)
                # keep only time-aligned true static, flattened
                S_trunc = S[k:]
                if S_trunc.ndim >= 3: S_trunc = S_trunc.reshape(S_trunc.shape[0], -1)
                elif S_trunc.ndim == 1: S_trunc = S_trunc[:, None]
                cov_static_trunc_only.append(S_trunc)
        else:
            for S, L in zip(cov_static_samples, lag_cov_list):
                S = np.asarray(S)
                S_trunc = S[k:]
                if S_trunc.ndim == 1: S_trunc = S_trunc[:, None]
                cov_static_trunc_only.append(S_trunc)
                cov_static_new.append(np.concatenate([S_trunc, np.asarray(L)], axis=1))
        
    else:
        cov_static_new = None
        cov_static_trunc_only = []
        if is_image:
            for i, X in enumerate(dset_samples):
                X = np.asarray(X)
                T = X.shape[0]
                Prow = np.asarray(cov_static_samples[i])
                if Prow.ndim >= 2: Prow = Prow.reshape(-1)
                if Prow.ndim == 0: Prow = Prow[None]
                Tk = T - k
                P_b = np.repeat(Prow[None, :], Tk, axis=0)   # (T-k, Ss)
                cov_static_trunc_only.append(P_b)
        else:
            for i, L in enumerate(lag_cov_list):
                Prow = np.asarray(cov_static_samples[i])
                if Prow.ndim >= 2: Prow = Prow.reshape(-1)
                if Prow.ndim == 0: Prow = Prow[None]
                Tk = np.asarray(L).shape[0]
                P_b = np.repeat(Prow[None, :], Tk, axis=0)
                cov_static_trunc_only.append(P_b)
                cov_static_new.append(np.concatenate([P_b, np.asarray(L)], axis=1))

    if is_image:
        del lag_cov_list
        gc.collect()
        
    if dist.get_rank() == 0:
        data_outdir = os.path.join(opts.outdir, 'data')
        os.makedirs(data_outdir, exist_ok=True)
        np.savez(os.path.join(data_outdir, 'dataset_samples.npz'),
                 samples=np.array(dset_samples_trunc, dtype=object))
        if cov_dynamic_trunc is not None:
            np.savez(os.path.join(data_outdir, 'cov_dynamic_samples.npz'),
                     samples=np.array(cov_dynamic_trunc, dtype=object))
        if is_image:
            # Save only compact static (no lag) to avoid huge files / RAM spikes
            np.savez(os.path.join(data_outdir, 'cov_static_trunc_only.npz'),
                     samples=np.array(cov_static_trunc_only, dtype=object))
        else:
            np.savez(os.path.join(data_outdir, 'lag_cov_list.npz'),
                     samples=np.array(lag_cov_list, dtype=object))
            np.savez(os.path.join(data_outdir, 'cov_static_new.npz'),
                     samples=np.array(cov_static_new, dtype=object))
        dist.print0(f"Saved processed data to: {data_outdir}")
    
    dset_samples = dset_samples_trunc
    cov_dynamic_samples = cov_dynamic_trunc
    cov_static_samples = cov_static_new
    
    # (4) Infer data_dim from loaded data
    if is_image:
        # dset_samples_trunc element is (T, C, H, W)
        _, C, H, W = np.asarray(dset_samples_trunc[0]).shape
        opts.data_dim = int(C * H * W)
    else:
        # vectors: (T, D)
        opts.data_dim = int(np.asarray(dset_samples_trunc[0]).shape[1])

    if opts.dim_cov_dynamic is None:
        src_dyn = cov_dynamic_trunc
        opts.dim_cov_dynamic = (src_dyn[0].shape[1] if (isinstance(src_dyn, list) and len(src_dyn) > 0) else 0)


    if opts.dim_cov_static is None:
        if is_image:
            # static (time-varying) part we kept is cov_static_trunc_only; lag is on-the-fly
            static_dim = (cov_static_trunc_only[0].shape[1] if len(cov_static_trunc_only) > 0 else 0)
            _T, _C, _H, _W = np.asarray(dset_samples_trunc[0]).shape
            lag_dim = (k + 1) * _C * _H * _W
            opts.dim_cov_static = static_dim + lag_dim
        else:
            if cov_static_samples is not None:
                opts.dim_cov_static = cov_static_samples[0].shape[1]
            else:
                opts.dim_cov_static = 0

    if is_image:
        cov_static_samples_for_dataset = cov_static_trunc_only
    else:
        cov_static_samples_for_dataset = cov_static_samples

    
    # Infer n_trajs
    inferred_n_trajs = len(dset_samples)

    # (5) Build config dict. (c)
    # Initialize config dict.
    c = dnnlib.EasyDict()

    # (5a) Dataset & dataloader kwargs
    # Setup dataset kwargs
    c.dataset_kwargs = dnnlib.EasyDict(dset_name=opts.data_name, dt=opts.dt, balls_dset_specs=None)
    c.dataset_kwargs.n_trajs = inferred_n_trajs

    # (5b) Create dataset object with covariate support
    dataset_obj = dnnlib.util_v7.ToyDsetDynamics(
        data=dset_samples_trunc, dt=opts.dt, nForward=1,
        cov_dynamic_data=cov_dynamic_trunc,
        cov_static_data=cov_static_samples_for_dataset,
        image_lag_source=(dset_samples_orig if is_image else None),
        lag_k=(k if is_image else 0)
    )

    c.dataset_obj = dataset_obj
    c.dset_samples = dset_samples_trunc
    
    # Setup dataloader kwargs
    c.data_loader_kwargs = dnnlib.EasyDict(
        pin_memory=True, 
        num_workers=opts.workers, 
        prefetch_factor=2
    )
    
    # (5c) Setup optimizer kwargs
    c.optimizer_kwargs = dnnlib.EasyDict(class_name='torch.optim.AdamW', lr=opts.lr, betas=[0.9,0.999], eps=1e-8)
    c.optimizer_kwargs.lr_floor_abs = opts.lr_floor
    c.optimizer_kwargs.lr_decay_kimg  = opts.lr_decay_kimg

    c.update(k_target=opts.k_target,
             ndrop_ramp_kimg=opts.ndrop_ramp_kimg,
             ndrop_p0=opts.ndrop_p0,
             )
    
    # (5d) Setup loss kwargs
    c.loss_kwargs = dnnlib.EasyDict(flow_matcher_type=opts.flow_matcher_type, 
                                    sigma_dynamics=opts.sigma_dyn_fm, 
                                    sigma_compression=opts.sigma_comp_fm,
                                    class_name='training.loss_v7.VFMToyLoss')
    
    # (5e) Setup network kwargs
    c.network_kwargs = dnnlib.EasyDict(
        dyn_model_type=opts.dyn_arch, 
        use_t_dyn=opts.use_t_dyn,
        flow_model_type=opts.flow_arch, 
        encoder_type=opts.encoder_arch, 
        data_dim=opts.data_dim, 
        k_max=opts.k_max, 
        depth_mlp_cmp=opts.mlp_depth_cmp, 
        width_mlp_cmp=opts.mlp_width_cmp, 
        depth_mlp_dyn=opts.mlp_depth_dyn, 
        width_mlp_dyn=opts.mlp_width_dyn,
        depth_encoder=opts.encoder_depth, 
        width_encoder=opts.encoder_width,
        d_min=opts.d_min,
        img_size=opts.data_imgshape, 
        in_ch=opts.data_inch,
        dim_cov_dynamic=opts.dim_cov_dynamic,
        dim_cov_static=opts.dim_cov_static,
        flow_detach=opts.flow_detach,
        class_name='training.networks_v7.VFMToyNet'
    )

    ch_cmp = _parse_channels_csv(getattr(opts, 'conv_ch_cmp', None))
    ch_dyn = _parse_channels_csv(getattr(opts, 'conv_ch_dyn', None))
    ch_enc = _parse_channels_csv(getattr(opts, 'conv_ch_enc', None))
    if ch_cmp is not None: c.network_kwargs.channels_cmp = ch_cmp
    if ch_dyn is not None: c.network_kwargs.channels_dyn = ch_dyn
    if ch_enc is not None: c.network_kwargs.channels_enc = ch_enc
        
    if getattr(opts, 'conv_embed_cmp', None) is not None:
        c.network_kwargs.conv_embed_dim_cmp = int(opts.conv_embed_cmp)
    if getattr(opts, 'conv_embed_dyn', None) is not None:
        c.network_kwargs.conv_embed_dim_dyn = int(opts.conv_embed_dyn)
    
    # (5f) training options.
    c.total_kimg = max(int(opts.duration * 1000), 1)
    
    hist_lag_dim = 0
    hist_static_dim = 0
    if (getattr(opts, 'lag_k', 0) or 0) >= 0 and (getattr(opts, 'dim_cov_static', 0) or 0) > 0:
        if _is_movie_trials(dset_samples_trunc):
            _T, _C, _H, _W = np.asarray(dset_samples_trunc[0]).shape
            base = _C * _H * _W
        else:
            base = int(opts.data_dim)
        
        hist_lag_dim    = (k + 1) * base
        hist_static_dim = max(0, int(opts.dim_cov_static or 0) - hist_lag_dim)

        expected = int(opts.dim_cov_static or 0)
        actual   = hist_static_dim + hist_lag_dim
        if actual != expected:
            raise ValueError(f"dim_cov_static mismatch: expected {expected}, got {actual}")
    
    # Pass knobs to training loop (via **c)
    c.hist_noise_std = float(getattr(opts, 'hist_noise_std', 0.0) or 0.0)
    c.hist_lag_dim = int(hist_lag_dim)
    c.hist_static_dim = int(hist_static_dim)

    # (5g) ENA & logging
    kimg_per_tick_train = opts.tick_train
    kimg_per_tick_print = opts.tick_print or kimg_per_tick_train
    
    c.use_ema = opts.use_ema
    c.ema_halflife_kimg = int(opts.ema * 1000) # only used if use_ema==True 
    c.update(batch_size=opts.batch, batch_gpu=opts.batch_gpu)
    c.update(loss_scaling=opts.ls, cudnn_benchmark=opts.bench)
    c.update(kimg_per_tick=kimg_per_tick_train,
             kimg_per_tick_print=kimg_per_tick_print,
             snapshot_ticks=opts.snap, state_dump_ticks=opts.dump)

    c.update(alpha=opts.alpha, beta=opts.beta,
             gamma=opts.gamma, eta=opts.eta,
             eta_decay_start_kimg=opts.eta_decay_start_kimg,
             eta_decay_halflife_kimg=opts.eta_decay_halflife_kimg,
             eta_floor=opts.eta_floor,
            )
    c.update(grad_clip=opts.grad_clip, grad_clip_val=opts.grad_clip_val)
    # c.update(decay_lambda=opts.decay_lambda)
    c.update(
        d_penalty_type   = opts.d_penalty_type,
        d_penalty_lambda = opts.d_penalty_lambda,
        d_hs_tau         = opts.d_hs_tau,
    )
    
    
    # (5h) Random seed.
    if opts.seed is not None:
        c.seed = opts.seed
    else:
    
        # # local version
        # seed = torch.randint(1 << 31, size=[], device=device)
        # torch.distributed.broadcast(seed, src=0)
        # c.seed = int(seed)
        
        # DCC version
        seed_t = torch.randint(1 << 31, (1,), device="cpu")
        torch.distributed.broadcast(seed_t, src=0)
        c.seed = int(seed_t.item())

    # (5i) Resume learning
    if opts.resume is not None:
        match = re.fullmatch(r'training-state-(\d+).pt', os.path.basename(opts.resume))
        if not match or not os.path.isfile(opts.resume):
            raise click.ClickException('--resume must point to training-state-*.pt from a previous training run')
        c.resume_pkl = os.path.join(os.path.dirname(opts.resume), f'network-snapshot-{match.group(1)}.pkl')
        c.resume_kimg = int(match.group(1))
        c.resume_state_dump = opts.resume

    # (6) Description string.
    schedule_type_str = 'prp' if opts.data_dim == opts.k_max else 'prr' 
    desc = f'{opts.data_name}-{schedule_type_str}-uncond-{opts.flow_matcher_type}FM-gpus{dist.get_world_size():d}-batch{c.batch_size:d}-fp32'

    if opts.desc is not None:
        desc += f'-{opts.desc}'

    # (7) Pick output directory.
    if dist.get_rank() != 0:
        c.run_dir = None
    elif opts.nosubdir:
        c.run_dir = opts.outdir
    else:
        prev_run_dirs = []
        if os.path.isdir(opts.outdir):
            prev_run_dirs = [x for x in os.listdir(opts.outdir) if os.path.isdir(os.path.join(opts.outdir, x))]
        prev_run_ids = [re.match(r'^\d+', x) for x in prev_run_dirs]
        prev_run_ids = [int(x.group()) for x in prev_run_ids if x is not None]
        cur_run_id = max(prev_run_ids, default=-1) + 1
        c.run_dir = os.path.join(opts.outdir, f'{cur_run_id:05d}-{desc}')
        assert not os.path.exists(c.run_dir)

    # Define serializable version of c (exclude non-JSON-friendly keys)
    serial_c = {k: v for k, v in c.items() if k not in ['dataset_obj', 'dset_samples']}

    # (8) Print options.
    dist.print0()
    dist.print0('Training options:')
    dist.print0(json.dumps(serial_c, indent=2))
    dist.print0()
    dist.print0(f'Output directory:        {c.run_dir}')
    dist.print0(f'Dataset name:            {c.dataset_kwargs.dset_name}')
    dist.print0(f'Data dimension:          {opts.data_dim}')
    dist.print0(f'Dynamic covariate dimension:     {opts.dim_cov_dynamic}')
    dist.print0(f'use_t_dyn={opts.use_t_dyn}')
    dist.print0(f'Static covariate dimension:      {opts.dim_cov_static}')
    dist.print0(f'Lag k used:               {opts.lag_k}')
    dist.print0(f'Number of GPUs:          {dist.get_world_size()}')
    dist.print0(f'Batch size:              {c.batch_size}')
    dist.print0()
        
    # Dry run?
    if opts.dry_run:
        dist.print0('Dry run; exiting.')
        return

    # (9) Create output directory.
    dist.print0('Creating output directory...')
    if dist.get_rank() == 0:
        os.makedirs(c.run_dir, exist_ok=True)
        c.network_kwargs.update(save_dir=c.run_dir) # add save_dir to network kwargs 
        with open(os.path.join(c.run_dir, 'training_options.json'), 'wt') as f:
            json.dump(serial_c, f, indent=2)
        dnnlib.util_v7.Logger(file_name=os.path.join(c.run_dir, 'log.txt'), file_mode='a', should_flush=True)


    # (10) Train.
    toy_training_loop_vfm_v7.training_loop(**c)

#----------------------------------------------------------------------------

if __name__ == "__main__":
    main()

#----------------------------------------------------------------------------
```
